# Predicting Molecular Taste: Multi-Label and Multi-Class Classification

**DOI:** 10.1101/2025.05.02.651828

**Authors:** D.N. Siva Sathyaseelan, V. Ramanathan

## Abstract

Predicting the taste of chemical compounds is a complex task and has been a challenge for decades. This study explores the application of machine learning to predict taste profiles of chemical compounds using the **ChemTastesDB** dataset, comprising 2,944 tastants categorized into 44 taste labels and 9 taste classes. Addressing the challenges of label imbalance and correlation, the dataset was preprocessed using iterative stratified sampling and feature representations such as Mordred descriptors, Morgan fingerprints, and Daylight fingerprints. Baseline random forest models, along with binary relevance and classifier chains, were employed for multi-label classification, with evaluation metrics including micro-averaged F1 scores, precision, and recall. Results demonstrated that binary relevance models, particularly with Morgen fingerprints, achieved superior F1 scores, outperforming classifier chains likely due to random label ordering. Label correlation analysis via co-occurrence matrices and community detection revealed significant associations between taste labels, providing deeper insights into molecular taste interactions. Feature importance analysis highlighted structural elements influencing taste prediction. This work underscores the potential of computational models in advancing flavor science and paves the way for future exploration with deep learning and optimized label dependencies.

## INTRODUCTION

Taste perception plays a pivotal role in food science and flavor chemistry, influencing the development, selection, and acceptance of food products. Understanding the mechanisms by which humans perceive taste has been a long-standing interest, driven by advancements in computational chemistry and bioinformatics. The extraordinary developments in food informatics (computational food chemistry) and bioinformatics (computational biochemistry) have provided the necessary tools to study the receptor/ligand binding interaction. To achieve a particular taste, it is now understood that the structure of the receptors and the specific features of the tastant ligands to interact with receptors must be analyzed [1,2]. The five principal tastes are bitter, sweet, sour, umami, and salty, with each one being detected by specific receptors. Other tastes, such as fat taste, might be considered basic ones since they arise from the combination of somatosensory and gustation perceptions [3, 4]. In general, taste sensation relies on the affinity of specific biochemical compounds and their target taste receptors [5,6]. Small variations in tastants’ chemistry may result in a drastic change in perceived taste. Therefore, shedding light on the physio-chemical features of compounds in food is of primary importance to pinpoint the molecular bases and mechanisms determining the food taste and subsequent food consumption [7].

In recent years, several studies have developed machine-learning (ML) tools to predict the taste of specific compounds starting from their chemical structure [8]. However, many of these tools face limitations in accurately predicting multiple taste sensations and identifying the dominant taste of a chemical molecule based on its structure. To address these limitations, we propose a novel machine-learning framework that leverages advanced algorithms with the help of **ChemTastesDB**(Cristian Rojas, Davide Ballabio, Karen Pacheco Sarmiento, Elisa Pacheco Jaramillo, Mateo Mendoza, Fernando García) an extensive dataset of taste-related chemical structures.

Our approach focuses on improving the prediction of multiple taste sensations by incorporating comprehensive feature extraction techniques and multi-label classification models. Additionally, we introduce a methodology to determine the dominant taste of a molecule, ensuring a more accurate representation of its sensory profile using multi-class classification. This framework not only enhances predictive accuracy but also broadens the scope of taste prediction, providing a robust tool for researchers and industries involved in flavor science and food product development. This study leverages ChemTastesDB, a comprehensive dataset of 2,944 tastants categorized into 44 taste labels and 9 taste classes, to develop machine learning models for multi-label and multi-class taste prediction. The dataset was analyzed for label imbalance using metrics like MeanIR and label dependency through co-occurrence matrices and community detection algorithms. Molecular featurization was performed using Mordred descriptors, Morgan fingerprints, and Daylight fingerprints, with missing values imputed and labels encoded as binary vectors. Random forest classifiers served as the baseline model, with binary relevance and classifier chains employed for multi-label classification. Iterative stratified sampling ensured representative training and test splits, mitigating the impact of label imbalance. Model performance was evaluated using micro-averaged F1 scores, precision, and recall. Results revealed that binary relevance models trained on Morgan fingerprints yielded the best F1 scores, outperforming classifier chains, likely due to the random ordering of labels in the chains. The study also identified key molecular features influencing taste prediction and demonstrated that label correlations, while significant, did not consistently improve classifier chain performance. These findings provide a foundation for advancing computational taste prediction, with potential applications in food science, flavor chemistry, and bioinformatics.

## METHODS

### 1) Forming an integrated dataset

**ChemTastesDB** dataset consisting of 2944 tastants, including both organic and inorganic chemical compounds was used in this work. These tastants were categorized into various taste classes: the five basic tastes (sweet, bitter, umami, sour, and salty), as well as non-basic and other taste categories such as tasteless, non-sweet, multitaste, an miscellaneous. ChemTastesDB aims to support the scientific community by enhancing the understanding of taste molecules and facilitating research in the development of new tastants [9,10,11]. The database is freely accessible for public use, providing a valuable resource for food chemists, bioinformaticians, and other researchers interested in the molecular properties of taste. The database is freely available at the following URL: https://doi.org/10.5281/zenodo.5747393.

The data was compiled from various scientific sources, including 37 research papers, 3 books, and 53 book chapters, ensuring a broad spectrum of verified compounds and is shown in Table 1. Each entry in the dataset includes key information about the tastant, such as:

- **Name**
- **PubChem CID**
- **CAS Registry Number**
- **Canonical SMILES**
- **Taste Classification**
- **Scientific References**

**Table 1.** Random 5 entries in the dataset.

| ID | Name | PubChem CID | CAS number | canonical SMILES | Taste | Class taste | Reference_(cod)/[pp] |
| --- | --- | --- | --- | --- | --- | --- | --- |
| 0566 | Compound (R)-(+)-9 | * | * | COc1ccc(cc1O)C1OC2CC3CCC2(CS1)C3(C)C | [sweet] | sweetness | Bassoli2000_((+)-9) |
| 1221 | Artemisin | 65030 | 481-05-0 | CC1C2C(O)CC3(C)C=CC(=O)C(=C3C2OC1=O)C | [bitter] | bitterness | Dagan-Wiener2019_(463) |
| 1678 | L-Leucine | 6106 | 61-90-5 | CC(C)CC(N)C(O)=O | [bitter, slightly bitter, non-sweet] | bitterness | Belitz2009_[35]; Dagan-Wiener2019_(751); Glase... |
| 1578 | Hexethal sodium | 23690440 | 144-00-3 | [Na+].CCCCCCC1(CC)C(=O)NC(=NC1=O)[O-] | [bitter] | bitterness | Dagan-Wiener2019_(587) |
| 2832* | Sodium N-[3-chloro-5-(trifluoromethyl)phenyl]s... | * | * | [Na+].[O-]S(=O)(=O)Nc1cc(cc(c1)C(F)(F)F)Cl | [non-sweet] | non-sweetness | Spillane2009b_(51A) |

### 2) Exploring label imbalance and correlation

MLDs (Multi-Label Datasets), a comprehensive dataset of 2,944 tastants categorized into 44 taste labels and 9 taste classes, to develop machine learning models for multi-label and multi-class taste prediction. This dataset typically has a heavy label imbalance. To verify this a couple of label imbalance metrics by the name of MeanIR and IRLbl were computed and viewed as a histogram of taste labels to infer any remnant imbalances, if it existed. These are shown in Figure 1 and Figure 2.

**Fig 1.**
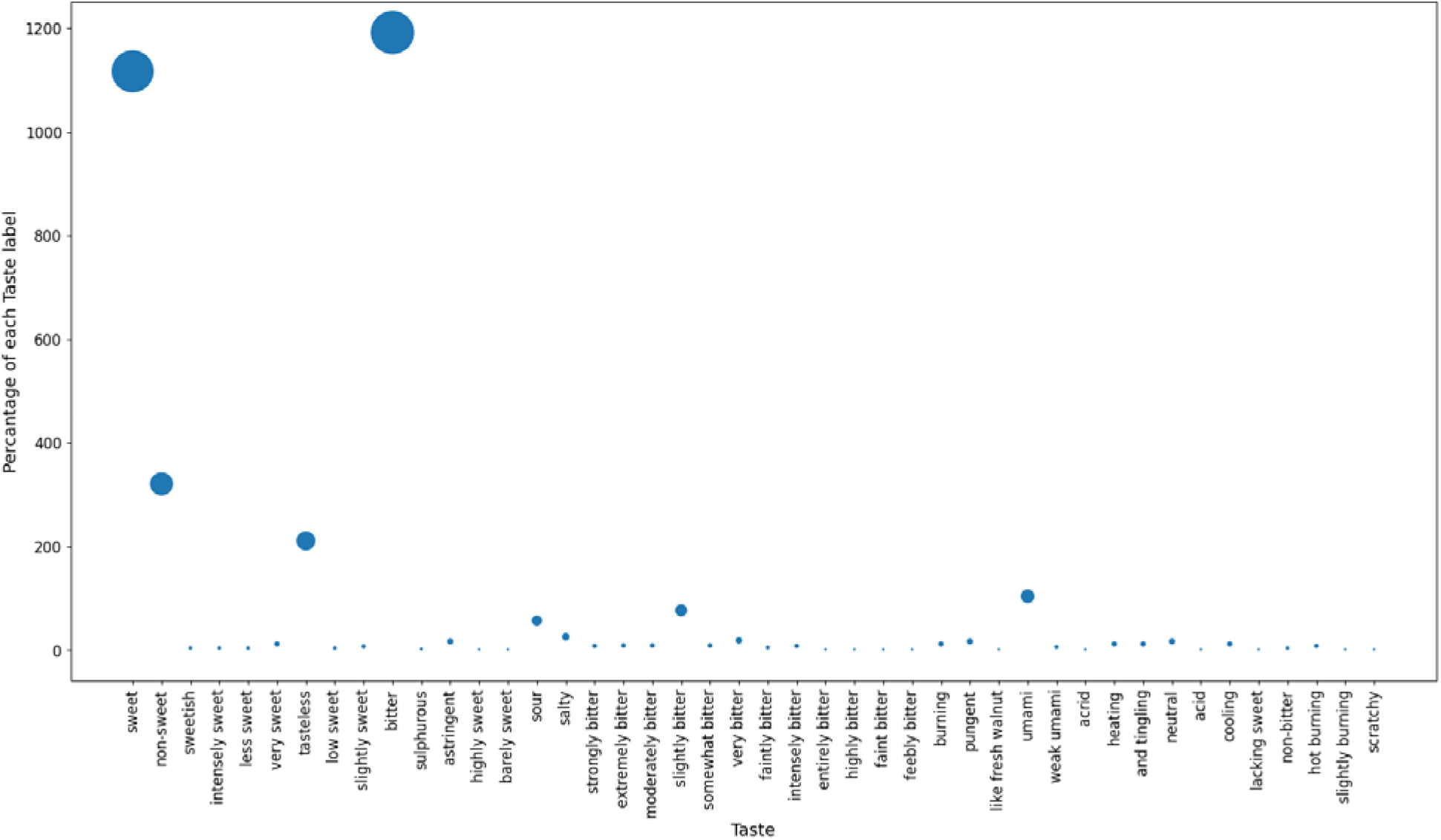
A scatter plot representing the number of samples in each of the 44 unique taste labels. The size of each point in the plot is proportional to the magnitude of the number of samples in that taste labels.

**Fig 2.**
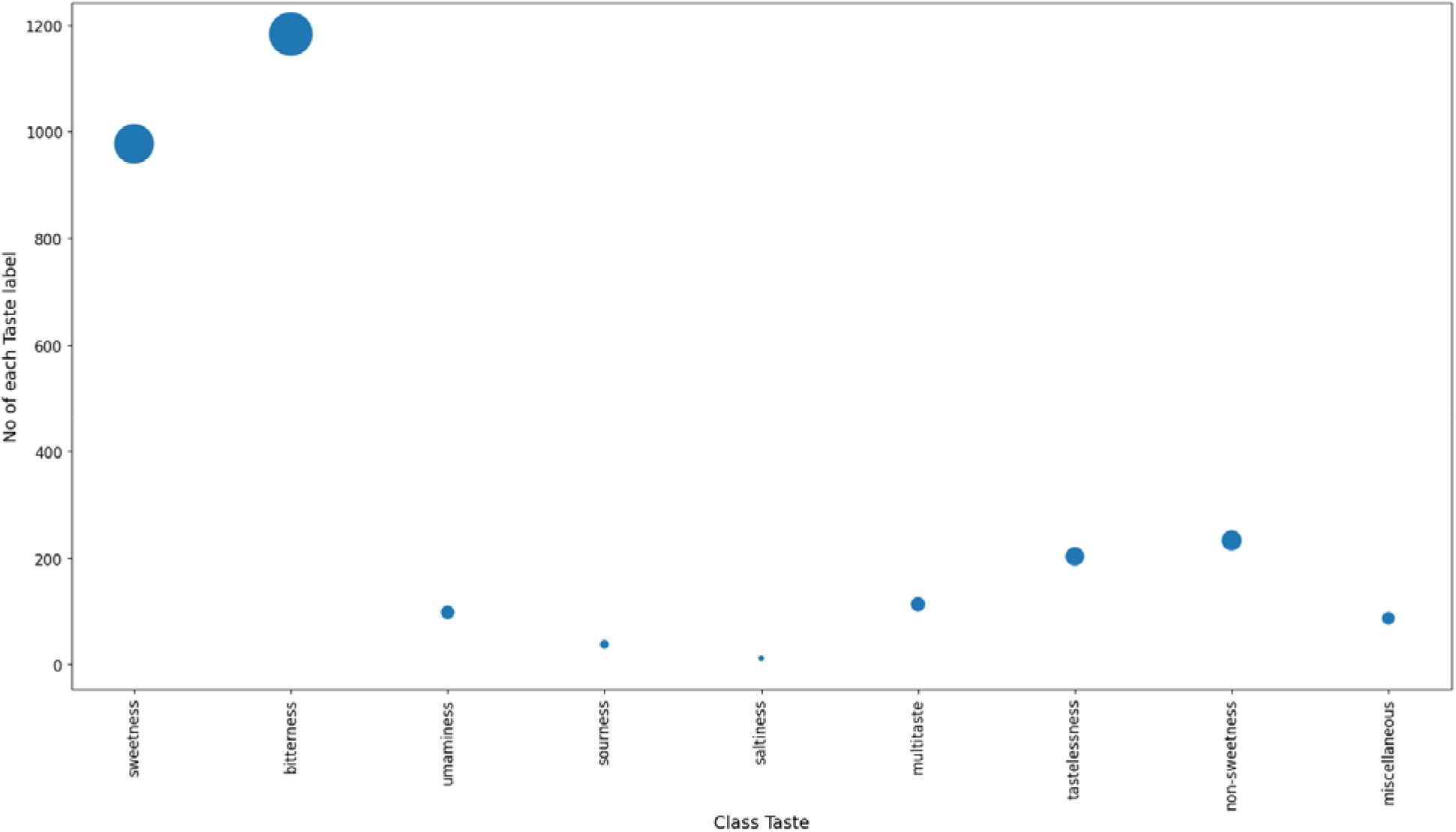
A scatter plot representing the number of samples in each of the 9 unique taste classes. The size of each point in the plot is proportional to the magnitude of the number of samples in that taste class.

Generally, complete independence between the labels is assumed. However, most researchers highlight the importance to take into account label dependency information [12]. For example, a ‘bitter’ label may be more likely to occur with the label ‘sweet’ based on linguistic similarity. A co-occurrence matrix (44×44, for 44 unique tastes in the integrated dataset) where i’th row and j’th column represents the frequency of the co-occurrence of the two labels was built to gauge label dependency up to second degree. To get a more global knowledge of any such label correlation, we ran the Louvain community detection algorithm on the taste labels which attempts to optimize modularity; a measure for the quality of partition between communities of nodes.

### 3) Featurizing molecules

To get a meaningful numerical representation of taste molecules for training our model, we used three traditional featurization techniques: - Mordred, Morgan, and daylight fingerprinting. The first of the three was generated using Mordred descriptor calculator [13] while Rdkit was used for the other two.

### 4) Pre-processing, training and evaluation metrics

Some columns from the feature space were dropped due to a large percentage of missing values. The remaining missing values were imputed using a KNN Imputer. Furthermore, each label set was converted into a 44-length bit vector; 1 denoting the presence of the label and 0 denoting its absence with the associated molecule.

A random forest classifier with multi-label support [14] in the scikit-learn library as the baseline model was used. Further, we use two different model approaches for chaining random forest models together: - Binary Relevance and Classifier chains [15].

To measure multi-label classifiers, the classes were subjected to the micro-averaging method where the individual true positives, false positives, and false negatives of the system for different label sets were averaged. The micro-averaged F1-Score represented the harmonic mean of micro averaged recall and micro averaged precision.

## Results and discussion

A preliminary histogram plot of Figure 3, Figure 4 and Table 2a, Table 2b, Table 3 revealed that “scratchy” was the least frequent label in the dataset, while “bitter” was the most frequent and “saltiness” was the least frequent class, while “bitterness” was most frequent. The distribution is highly skewed, indicating a significant label imbalance. This imbalance was further confirmed by calculating the MeanIR (425.37). The disparity in the absolute frequencies of the top 10 most and least frequently occurring taste labels, as shown in Table 2 and Table 3, highlights this skewness more clearly.

**Table 2.**
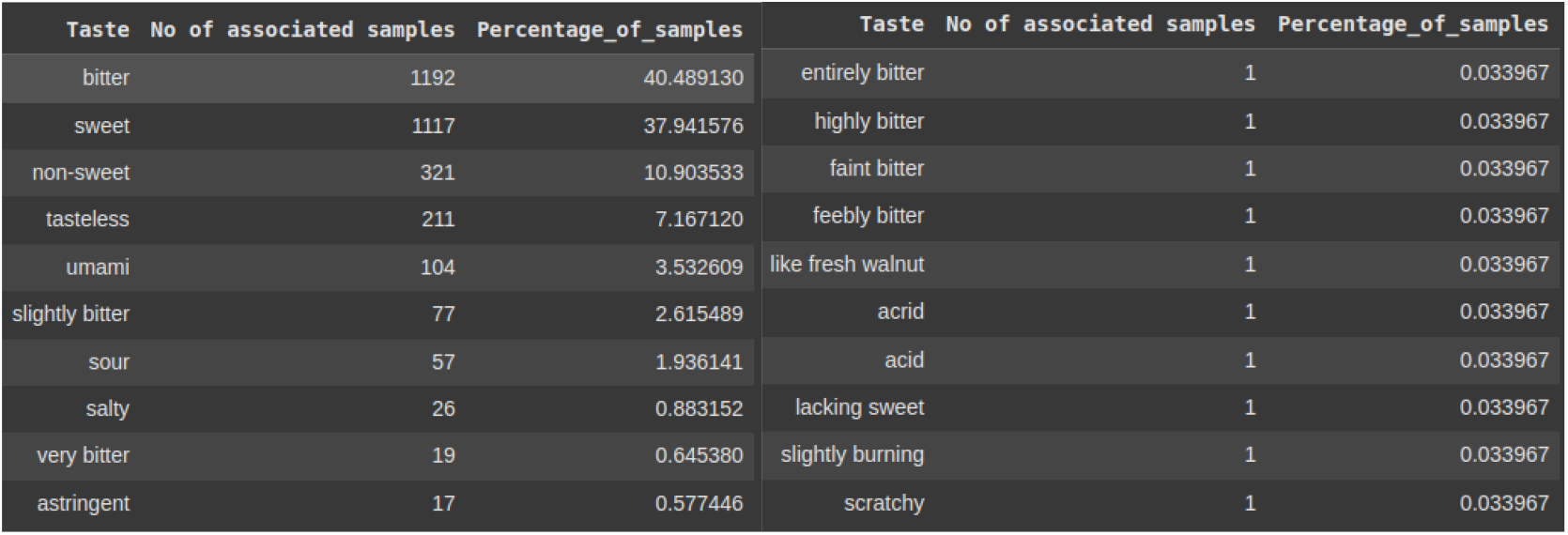
a and 2b: Top 10 most frequently and least frequently occurring taste labels respectively.

**Table 3.** Most frequently occurring Taste classes.

| Taste | No of associated samples | Percentage_of_samples |
| --- | --- | --- |
| bitterness | 1183 | 40.183424 |
| sweetness | 977 | 33.186141 |
| non-sweetness | 233 | 7.914402 |
| tastelessness | 203 | 6.895380 |
| multitaste | 113 | 3.838315 |
| umaminess | 98 | 3.328804 |
| miscellaneous | 87 | 2.955163 |
| sourness | 38 | 1.290761 |
| saltiness | 12 | 0.407609 |

**Fig 3.**
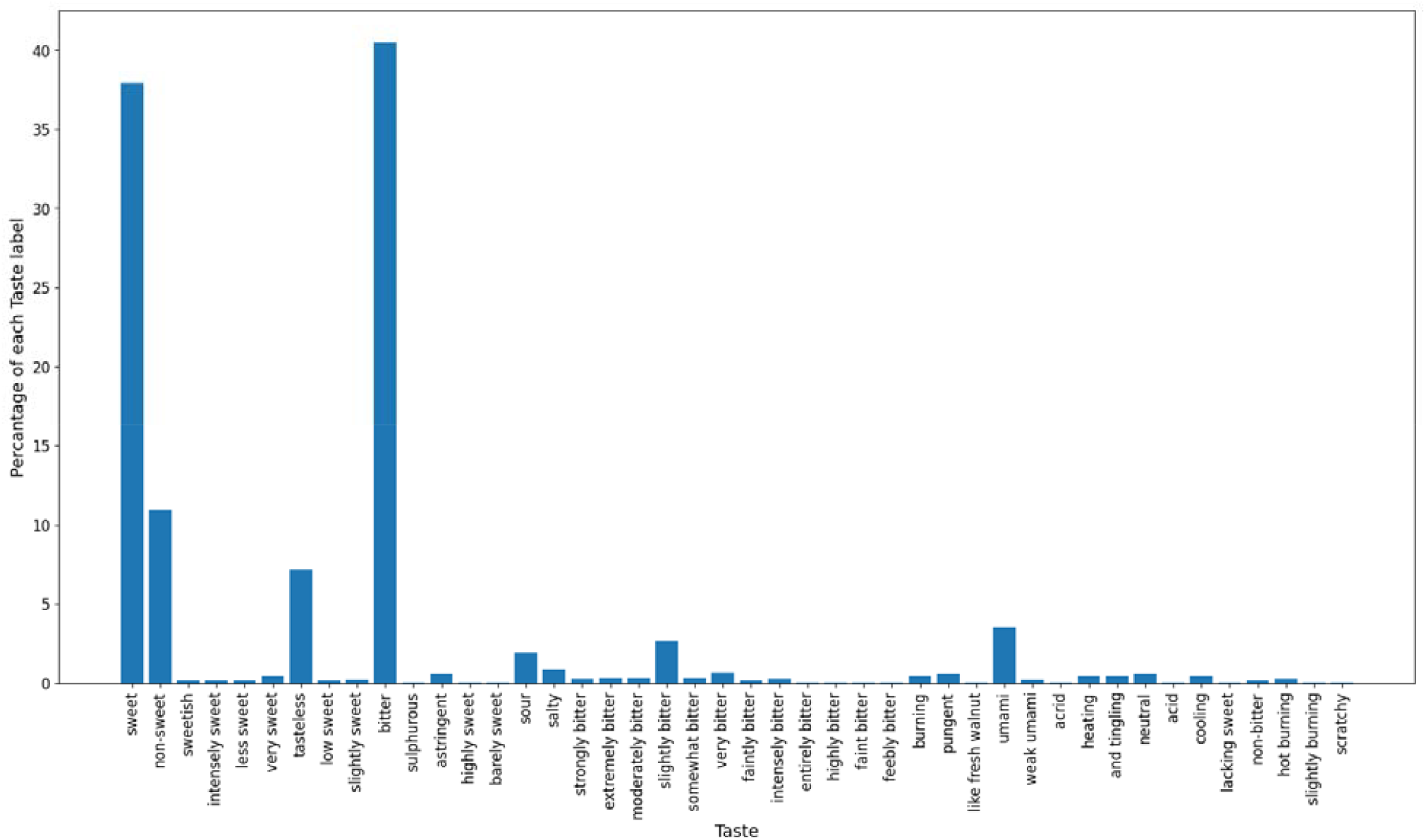
label imbalance with the bitter label occurring approximately 40 percent of the time as opposed to scratchy occurring less than 0.04 percent in **ChemTastesDB**.

**Fig 4.**
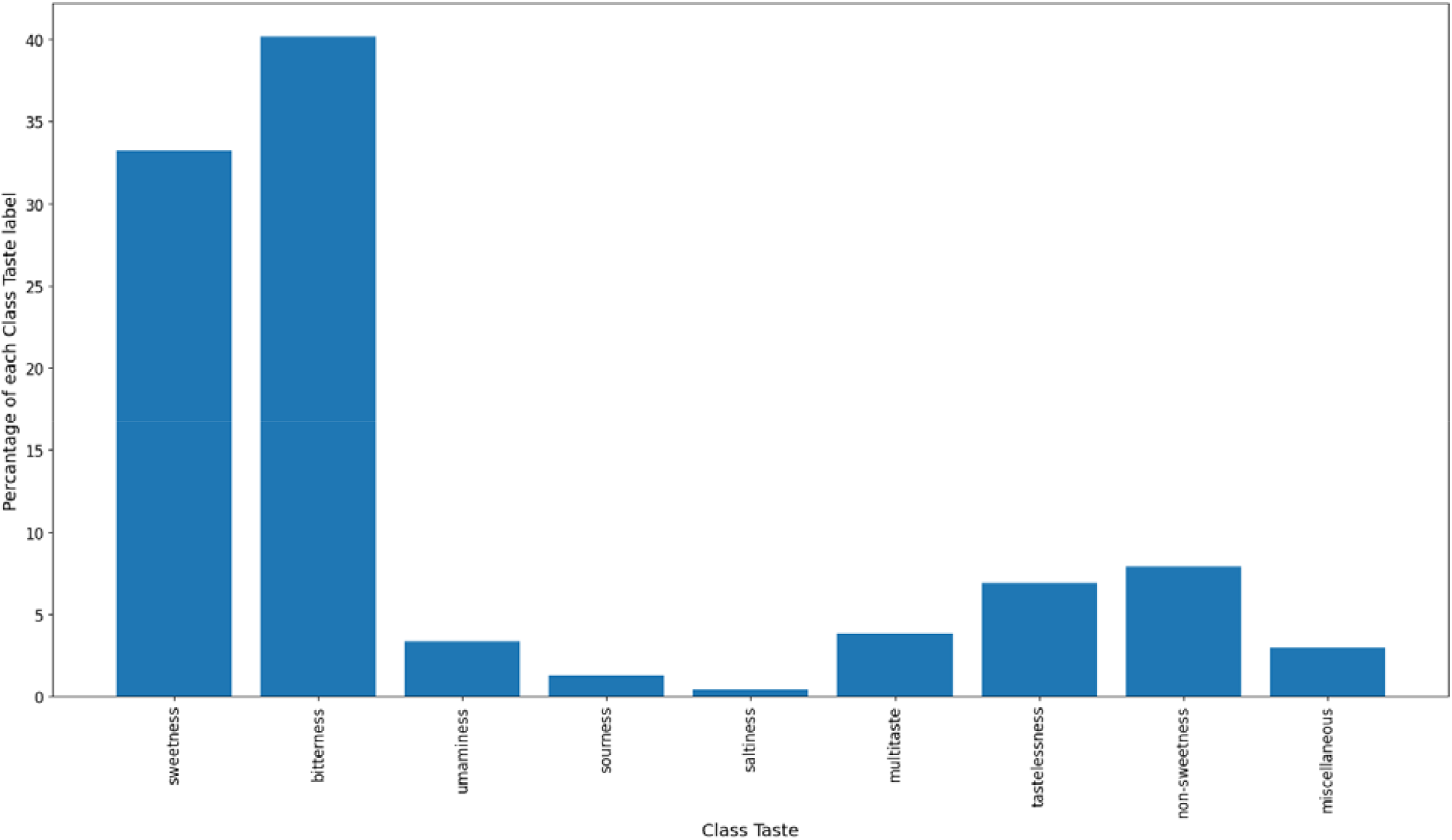
label imbalance with the bitterness class occurring approximately 40 percent of the time as opposed to saltness occurring less than 1 percent in **ChemTastesDB**.

To ensure the training and test datasets provide a general representation of the data, iterative stratified sampling was used instead of random sampling when splitting the dataset. This was achieved by utilizing the iterative_train_test_split class from the scikit-multilearn library. Further exacerbating the already skewed label imbalance. Figure 5a and Figure 5b shows the Random split versus a stratified split respectively on the data. The difference in the height of blue and orange bars represents the relative difference in the occurrence of a particular label in the test and training set. It is clearly visible that stratified sampling produces a much more representative training and test set.

**Fig 5a.**
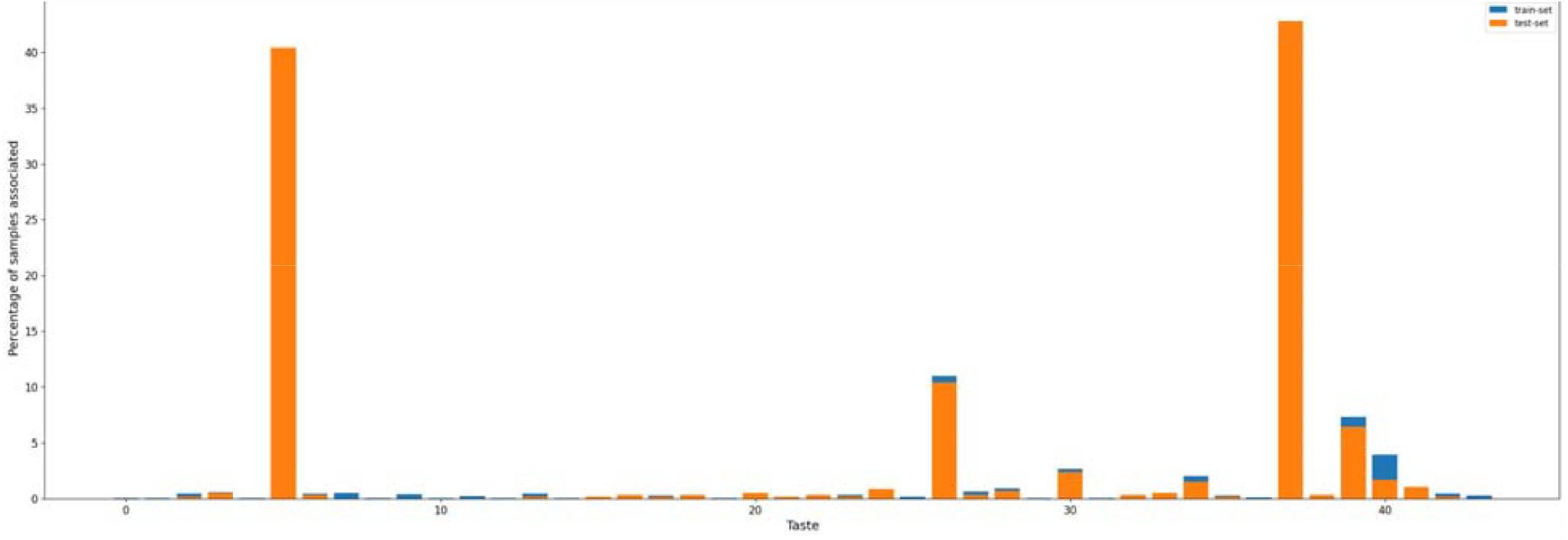
Label distribution while doing a random split

**Fig 5b.**
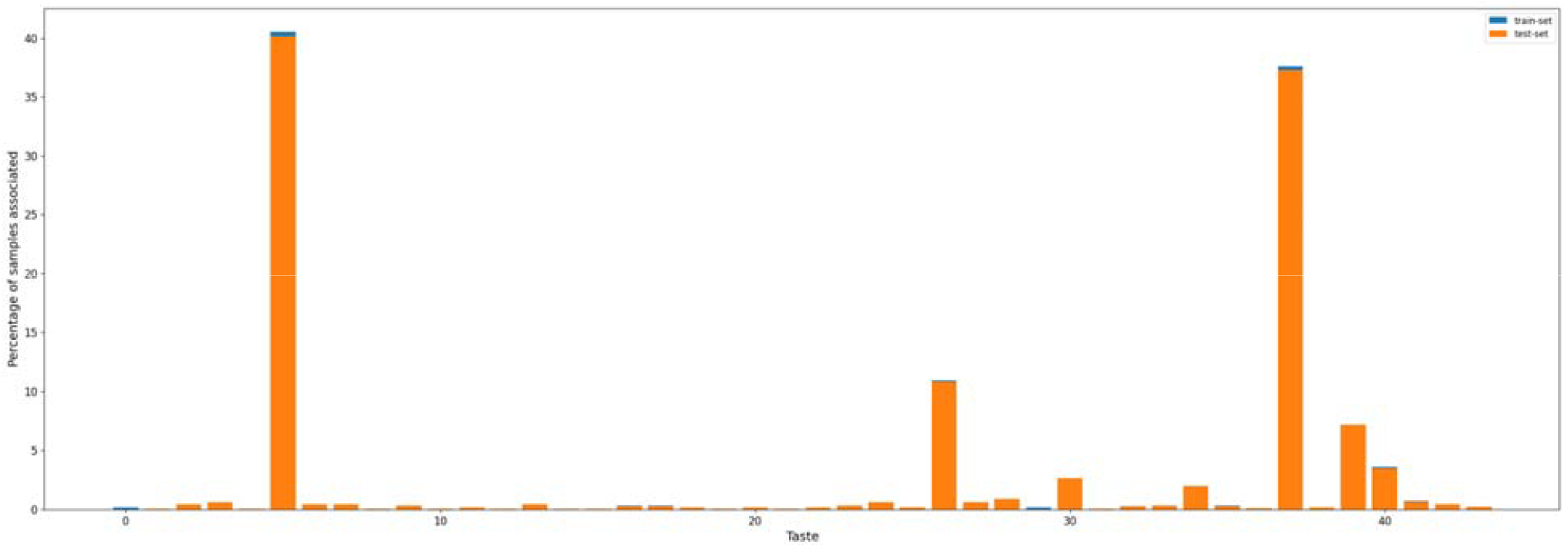
Label distribution while doing a stratified split

The co-occurrence matrix, represented in Figure 6a and Figure 6b, reveals that the labels “bitter” and “sweet” cooccur 113 times, while “sweet” and “non-sweet” co-occur 56 times, making them the two most frequently occurring label pairs (Table 4). This suggests that among molecules with a sweet taste, approximately 10.12% also exhibit a bitter taste. Conversely, around 9.48% of molecules with a bitter taste also have a sweet taste. Similarly, among molecules with a sweet taste, about 5.01% are also labeled as non-sweet. These findings align with typical patterns observed in multi-label datasets (MLDs), where correlations between labels frequently occur due to overlapping or complementary molecular properties.

**Table 4.**
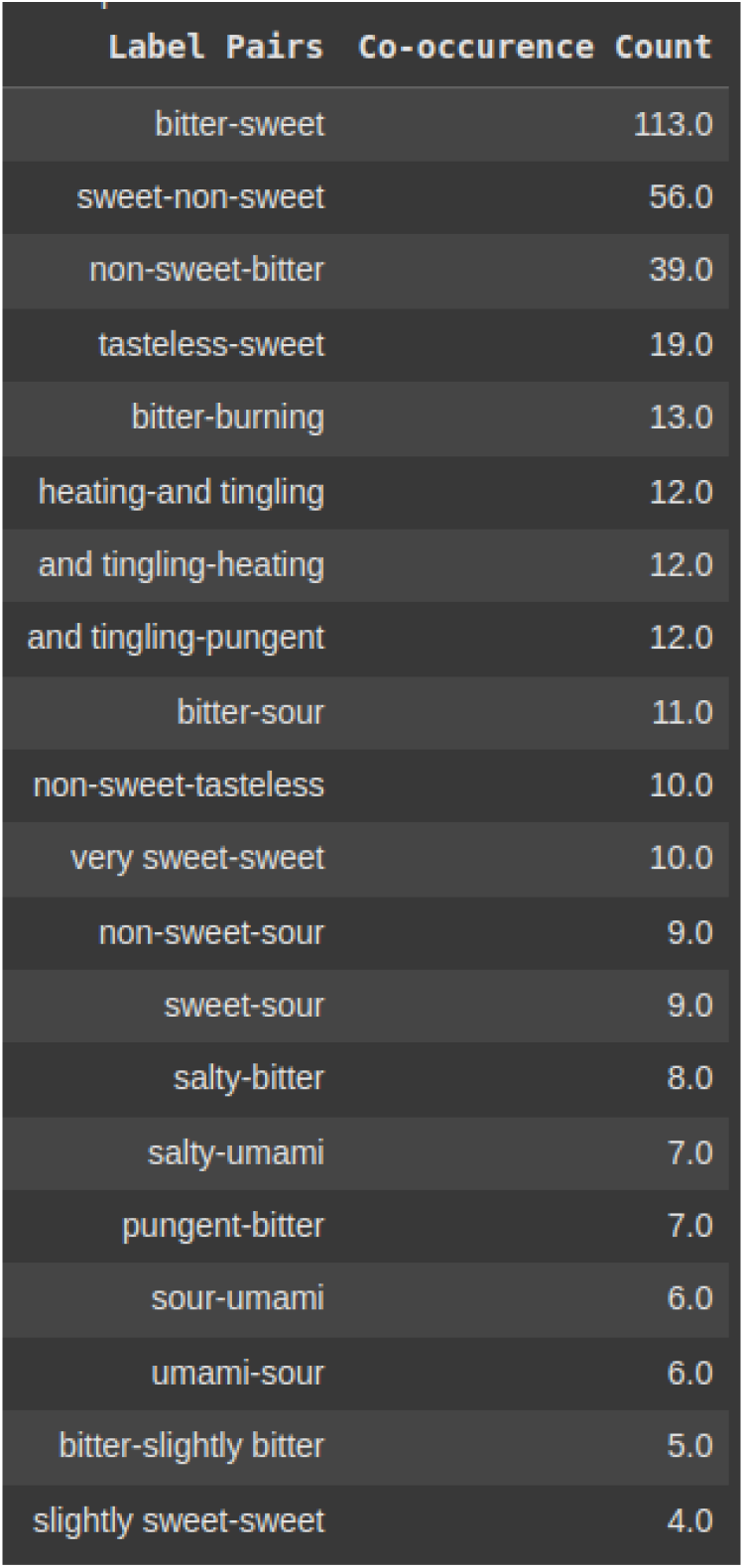
Top 20 taste associations.

| Label Pairs | Co-occurrence Count |
| --- | --- |
| bitter-sweet | 113.0 |
| sweet-non-sweet | 56.0 |
| non-sweet-bitter | 39.0 |
| tasteless-sweet | 19.0 |
| bitter-burning | 13.0 |
| heating-and tingling | 12.0 |
| and tingling-heating | 12.0 |
| and tingling-pungent | 12.0 |
| bitter-sour | 11.0 |
| non-sweet-tasteless | 10.0 |
| very sweet-sweet | 10.0 |
| non-sweet-sour | 9.0 |
| sweet-sour | 9.0 |
| salty-bitter | 8.0 |
| salty-umami | 7.0 |
| pungent-bitter | 7.0 |
| sour-umami | 6.0 |
| umami-sour | 6.0 |
| bitter-slightly bitter | 5.0 |
| slightly sweet-sweet | 4.0 |

**Fig 6a.**
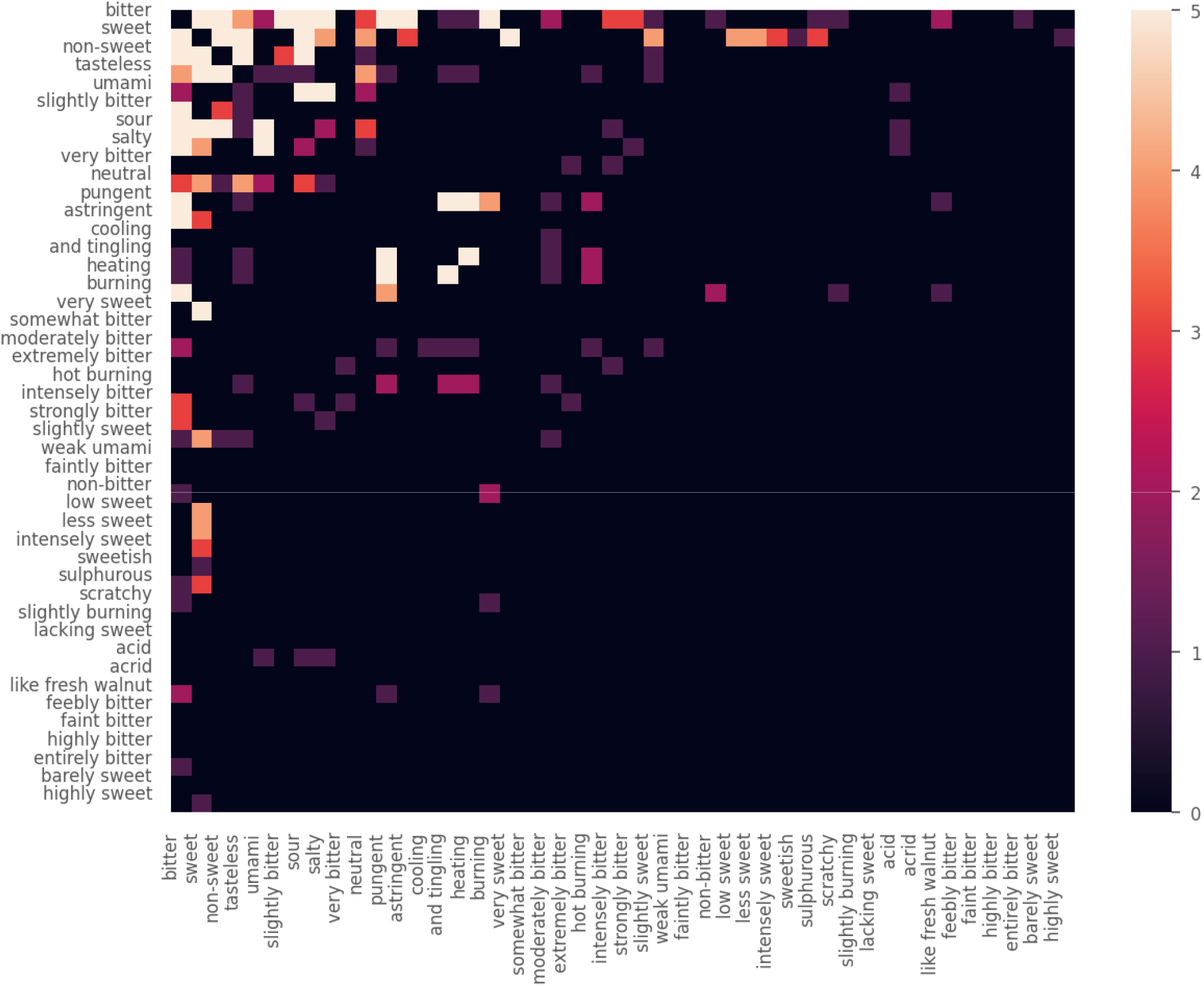
Heat map of our co-occurrence matrix truncated to top 44 taste labels. The darker the intensity of a square, the lesser the magnitude of co-occurrence count and correlation between the two labels.

**Fig 6b.**
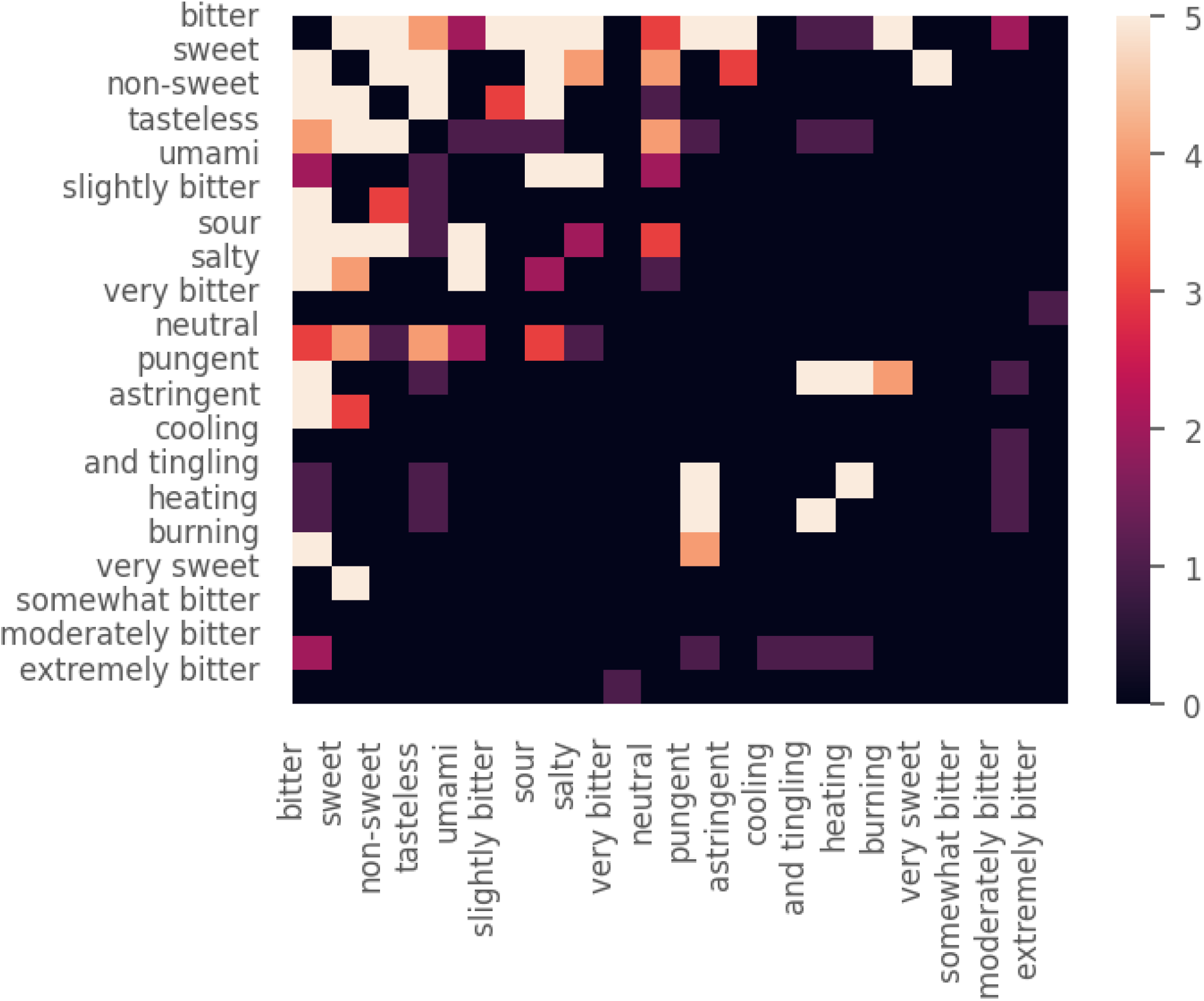
Heat map of our co-occurrence matrix truncated to top 20 taste labels. The darker the intensity of a square, the lesser the magnitude of co-occurrence count and correlation between the two labels.

Louvain community detection highlights correlations within larger label sets, and some groupings identified by the algorithm align with intuitive expectations. For instance, the five principal tastes: bitter, sweet, sour, umami, and salty, with each one being detected the most. The network graph further confirms the existence of global correlations between labels. As shown in Figure 7, the algorithm identified four clusters among 44 taste labels. This clustering approach is based on modularity, a measure that maximizes the difference between the actual number of edges within a community and the expected number of edges.

**Fig 7.**
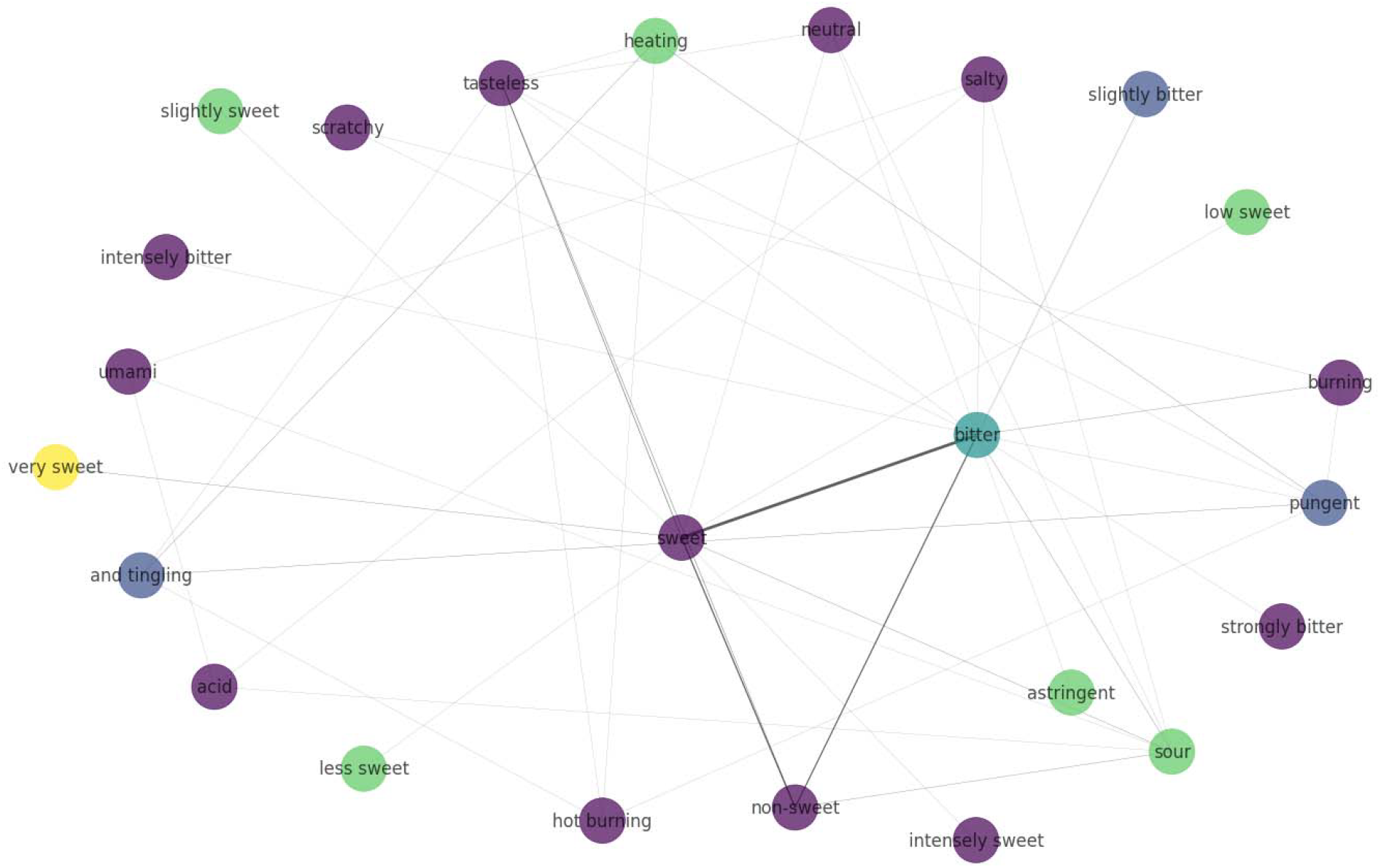
Graph constructed on the basis of Louvain method for community detection in networks. Nodes with the same colour belong to the same community.

The cardinality of our dataset (mean number of labels per sample) is 1.13, suggesting the multilabel-ness of our data is typical (most MLDs are in the {1, 5} interval) and the label density (cardinality divided by the number of labels) is 0.02576 (most MLDs have density values below 0.1). This value is useful to know how sparse are the label sets in the MLD. Higher density values denote label sets with more active labels than the lower ones.

Figure 8 shows the importance of features which was generated with random forests to get an idea of those features that contribute most significantly to the taste prediction task. Geary coefficient of lag 6 weighted by ionization potential (GATS6i) and centered Moreau-Broto autocorrelation of lag 1 weighted by Allred-Moscow EN (ATSC1are) are the top two Mordred features of importance, and further study into these might provide us with some insight into the kind of structural fragments or their relative spatial positions that result in imparting a particular taste to molecules.

**Fig 8.**
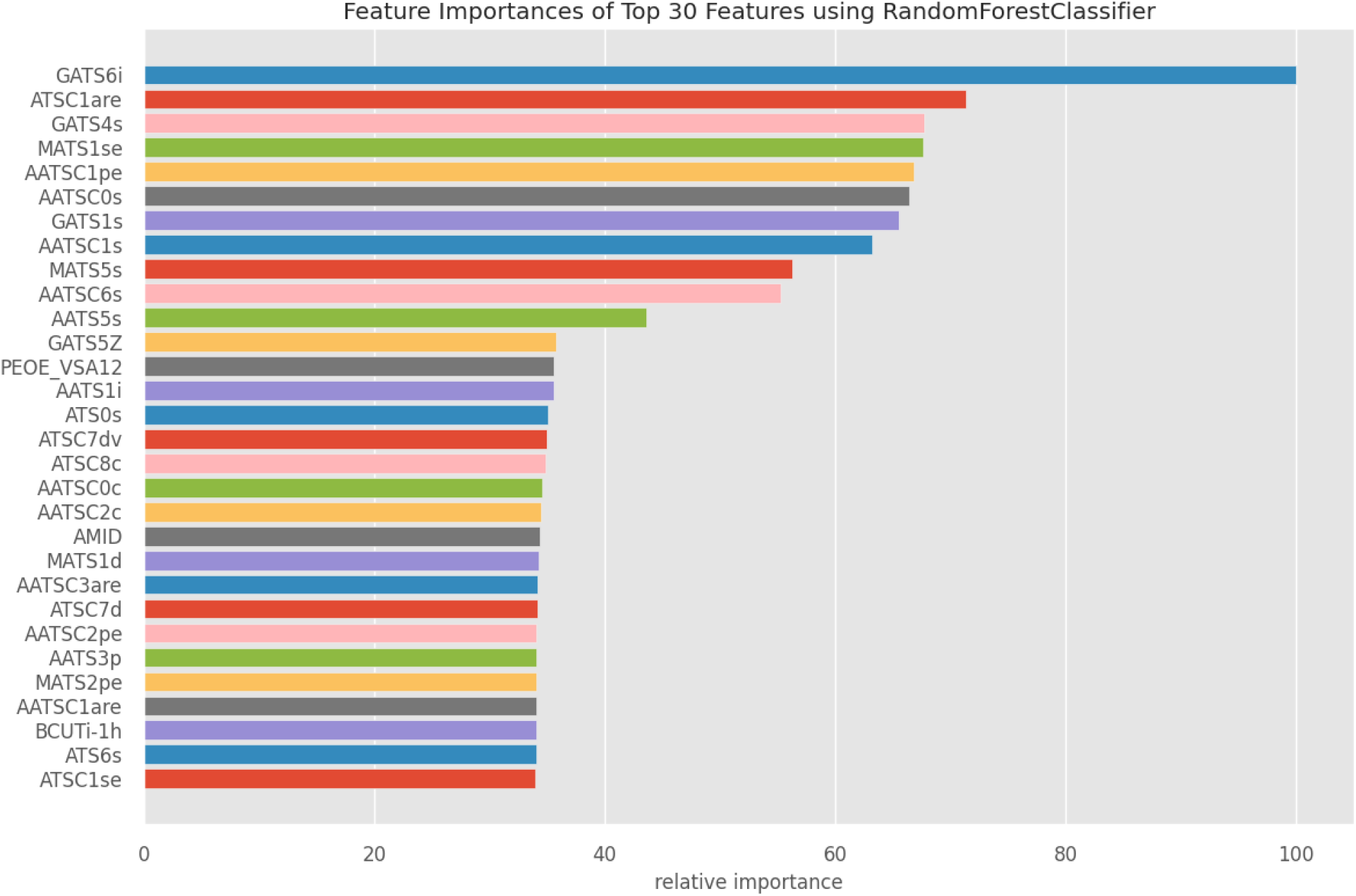
Feature importance based on closeness to the root node of decision trees in the forest computed on Mordred featurization

Both these descriptors are spatial autocorrelation descriptors which in general explain how the considered property is distributed along the topological structure of the molecule. Representing the molecule as a graph with atoms at the vertices and bonds as the edges, both the auto descriptors consider a certain molecular property (e.g. atomic masses, atomic van der Waals volumes, atomic Sanderson electronegativities, atomic polarizabilities, etc.) distribution between pairwise atoms in the molecule at a certain topological distance (smallest number of interconnecting bonds between the two atoms).

Performance of the model developed herein was evaluated on micro-averaged scores for each featurization revealed that the Random Forest model trained on Morgen Fingerprint yielded the best F1_score.

It is worthwhile noting that although there is a label correlation between our taste labels, we got better model performance from Random Forest, as evident from Table 5 and Table 6. And Morgan Fingerprint consistently outperforms Mordred Features and Daylight Fingerprint in terms of F1-score, Precision, and Recall across all model approaches. Daylight Fingerprint generally underperforms compared to Morgan Fingerprint and is comparable to Mordred Features in some cases.

**Table 5.** Showing micro averaged precision, recall, and F1 test scores for each featurization and model approach for multi-label classification.

|  | Random Forest |  |  | Binary Relevance |  |  | Classifier Chains |  |  |
| --- | --- | --- | --- | --- | --- | --- | --- | --- | --- |
|  | F1 score | Precision | Recall | F1 score | Precision | Recall | F1 score | Precision | Recall |
| Mordred Features | 0.57745<br>4886337<br>0049 | 0.76143<br>3868974<br>042 | 0.46508<br>1162702<br>9068 | 0.61842<br>3973362<br>9301 | 0.75053<br>8793103<br>4483 | 0.52585<br>8814647<br>0367 | 0.63200<br>4038364<br>4624 | 0.77860<br>6965174<br>1293 | 0.53186<br>0662701<br>7842 |
| Morgen Fingerprint | 0.62982<br>4561403<br>5087 | 0.75143<br>9037153<br>3228 | 0.54209<br>1355228<br>388 | 0.63346<br>2699439<br>4136 | 0.73856<br>2091503<br>268 | 0.55454<br>8886372<br>2159 | 0.64672<br>1833227<br>2437 | 0.73837<br>2093023<br>2558 | 0.57531<br>1438278<br>5957 |
| Daylight Fingerprint | 0.56477<br>1668219<br>9441 | 0.73767<br>4984783<br>9318 | 0.45753<br>1143827<br>8596 | 0.59441<br>4893617<br>0213 | 0.71980<br>6763285<br>0241 | 0.50622<br>8765571<br>9139 | 0.59320<br>1510775<br>3833 | 0.72084<br>2332613<br>3909 | 0.50396<br>3759909<br>3998 |

**Table 6.** Showing micro averaged precision, recall, and F1 test scores for each featurization and model approach for multi-class classification.

|  | Random Forest |  |  | Binary Relevance |  |  | Classifier Chains |  |  |
| --- | --- | --- | --- | --- | --- | --- | --- | --- | --- |
|  | F1 score | Precision | Recall | F1 score | Precision | Recall | F1 score | Precision | Recall |
| Mordred<br>Features | 0.62076<br>2155059<br>1327 | 0.81392<br>1433494<br>1419 | 0.50169<br>9235344<br>0951 | 0.62512<br>7681307<br>4566 | 0.78361<br>0755441<br>7413 | 0.51996<br>6015293<br>1181 | 0.63200<br>4038364<br>4624 | 0.77860<br>6965174<br>1293 | 0.53186<br>0662701<br>7842 |
| Morgen<br>Fingerpr<br>int | 0.65634<br>2182890<br>8554 | 0.77887<br>9813302<br>217 | 0.56711<br>9796091<br>7587 | 0.65953<br>3073929<br>9611 | 0.77133<br>1058020<br>4779 | 0.57604<br>0781648<br>2583 | 0.66650<br>9545133<br>1605 | 0.74854<br>4203282<br>1599 | 0.60067<br>9694137<br>6381 |
| Daylight<br>Fingerpr<br>int | 0.60157<br>0680628<br>2722 | 0.78376<br>5347885<br>4024 | 0.48810<br>5352591<br>3339 | 0.63042<br>3940149<br>6259 | 0.76328<br>5024154<br>5893 | 0.53695<br>8368734<br>0697 | 0.63636<br>3636363<br>6364 | 0.76614<br>8325358<br>8517 | 0.54418<br>0118946<br>4741 |

We also computed label-wise scores (Table 7) and class-wise score (Table 8), and observed that the top F1_scores generally belonged to labels that had a higher percentage of associated samples in our dataset, although there were exceptions to it like umami (3.532609 percentage).

**Table 7.**
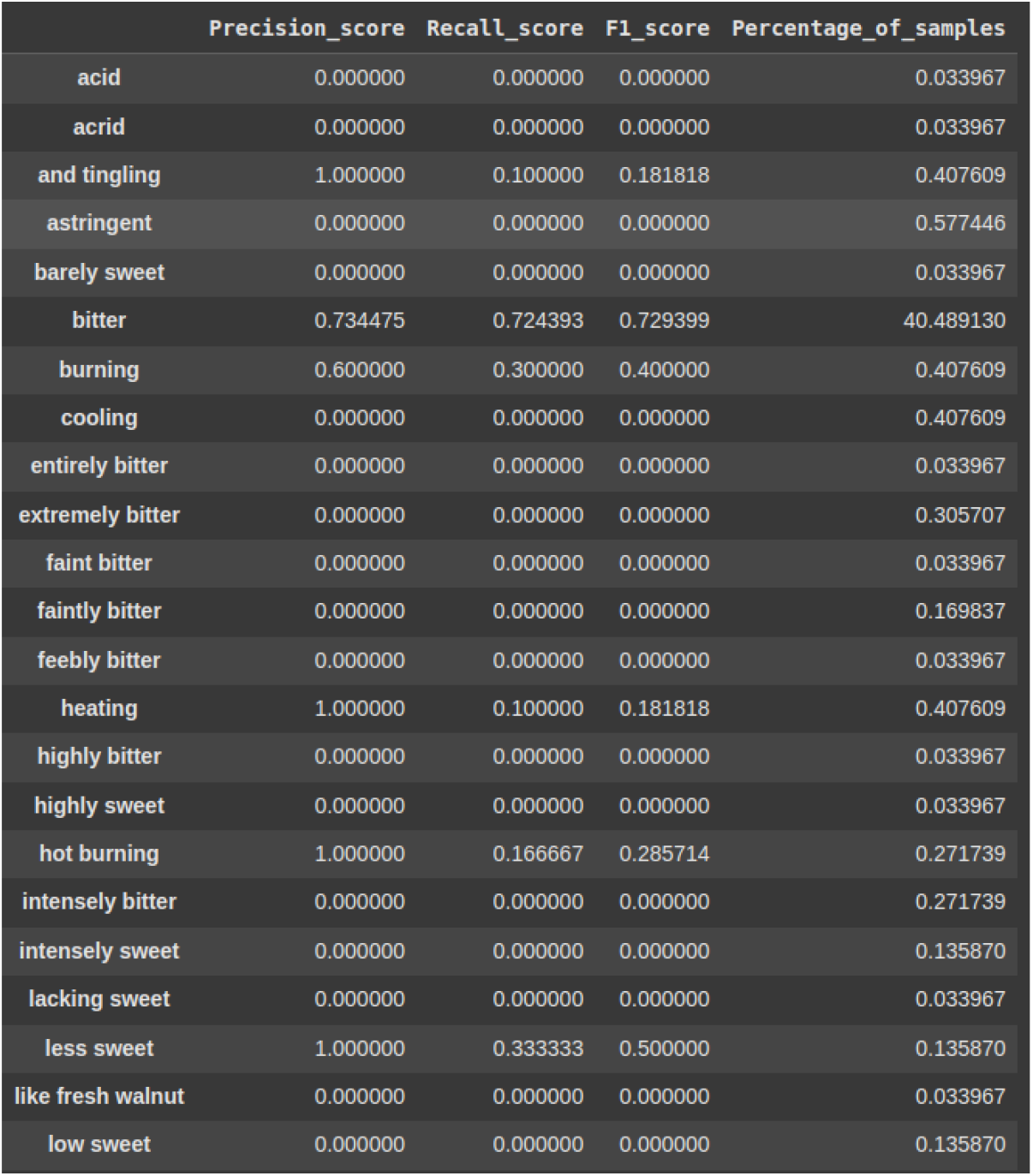
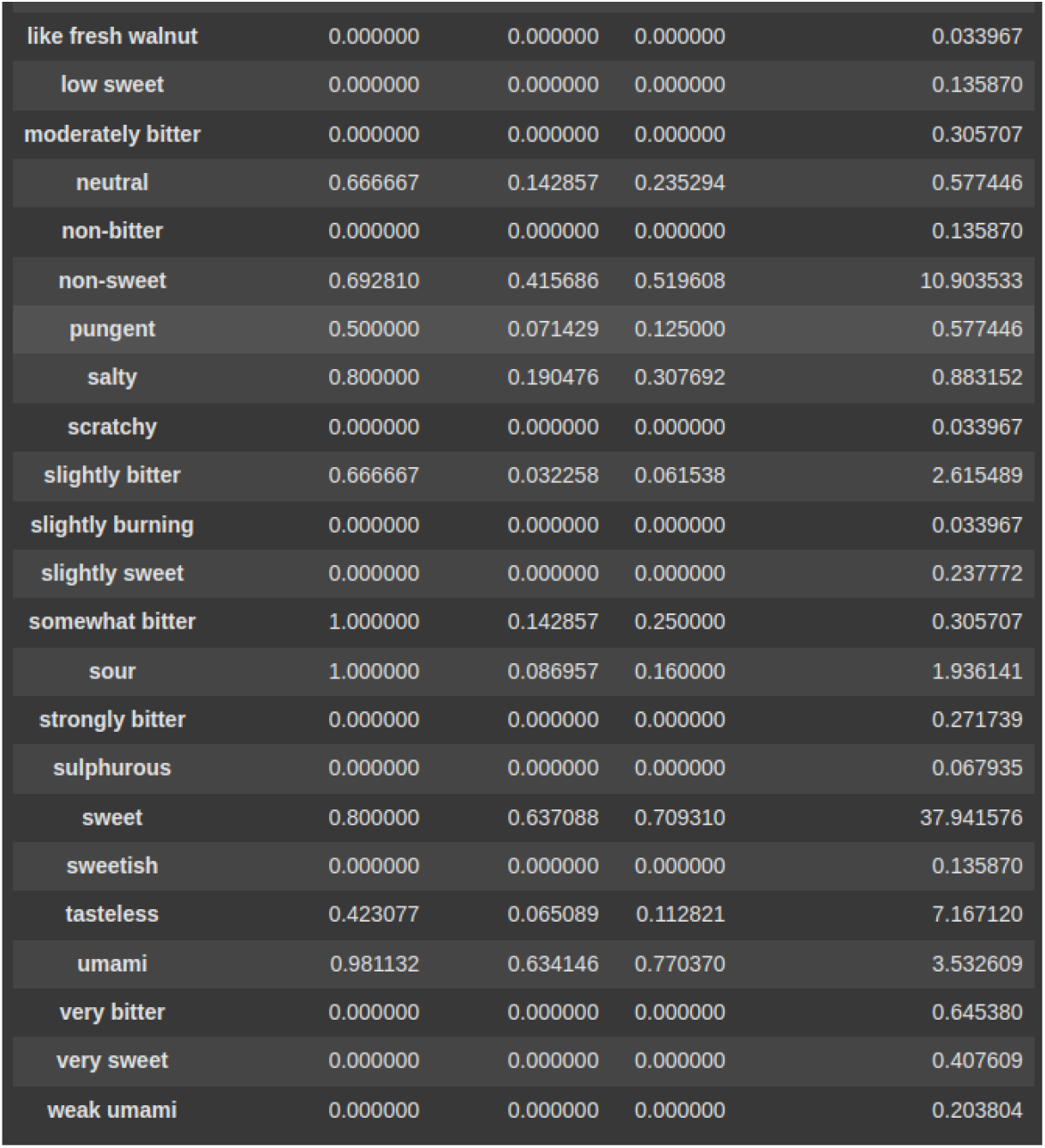
Label-wise F1_scores along with their precision and recall scores. Percentage of samples for labels was calculated on the entire dataset and not only the test set.

**Table 8.**
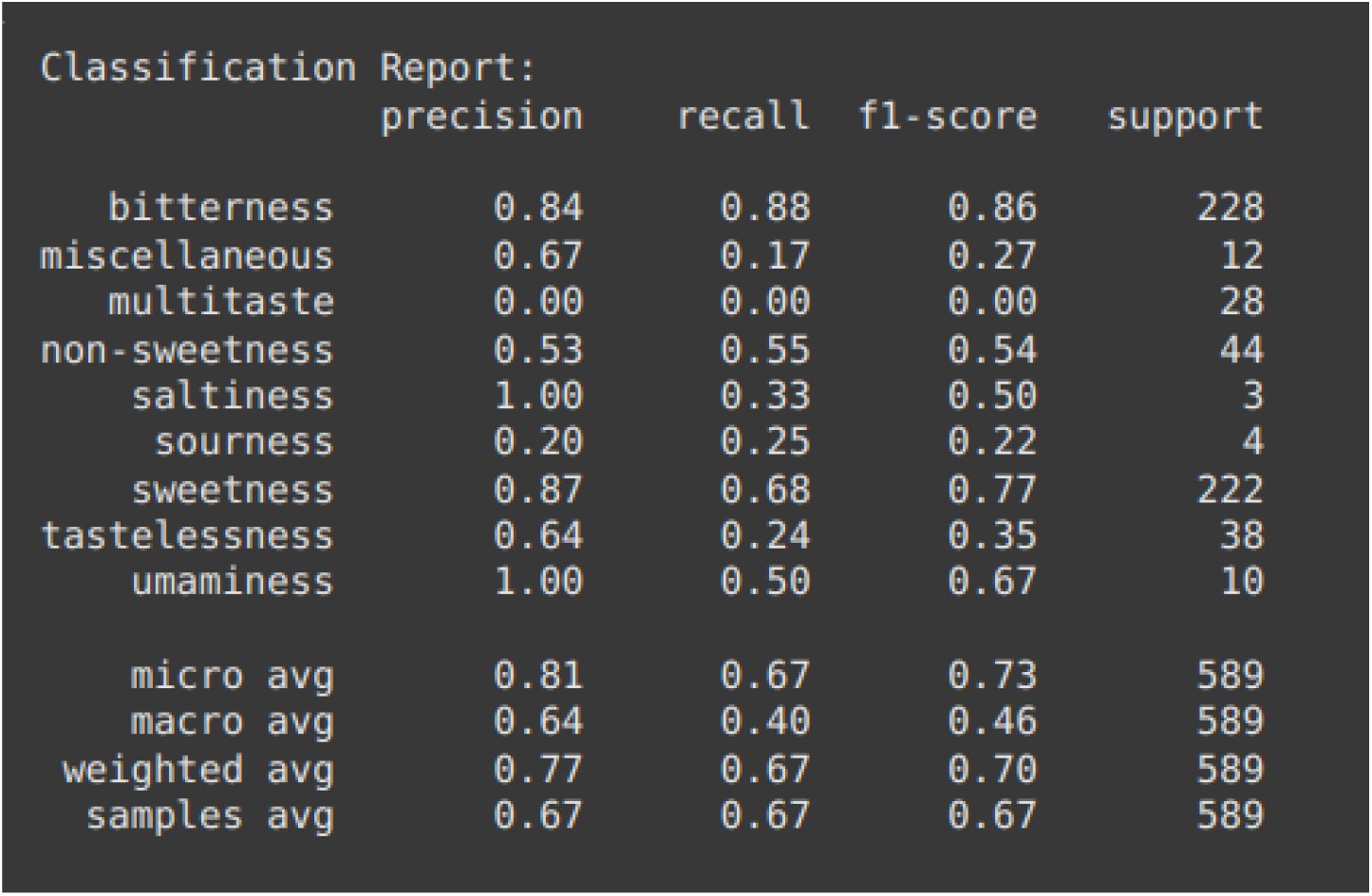
Classification report for multi-class classification.

## CONCLUSION

In this study several distinctive approaches to molecular taste prediction were successfully undertaken which sets it apart from previous works. Unlike traditional single-label classification models, our framework successfully handles both multi-label and multi-class taste prediction simultaneously, accommodating the complex reality that molecules can exhibit multiple taste sensations. The integration of Morgan fingerprints proved particularly effective, outperforming conventional Mordred descriptors and Daylight fingerprints across all evaluation metrics.

A key innovation of our approach lies in its comprehensive treatment of label correlations through co-occurrence matrices and community detection algorithms. This revealed previously unexplored relationships between taste labels, such as the significant co-occurrence between bitter and sweet tastes (∼10% overlap), providing valuable insights for molecular taste understanding. Our use of iterative stratified sampling to address label imbalance (MeanIR of 425.37) represents a methodological advancement in handling the inherent skewness of taste data.

The future scope of this research encompasses several promising directions for advancement. Deep learning integration, particularly through graph neural networks and transformer architectures, could significantly enhance molecular structure interpretation and feature extraction. The model could be improved by optimizing label ordering in classifier chains, developing hybrid fingerprinting techniques, and implementing multi-task learning approaches that simultaneously predict taste and other molecular properties. Practical applications could include developing user-friendly tools for food scientists and integration with existing molecular databases, while expanding the model to predict taste intensity and temporal characteristics would provide more comprehensive flavor profiles. Additionally, experimental validation through sensory panels and investigation of taste-odor interactions would strengthen the model’s reliability. The framework could also be extended to explore inverse molecular design, where specific taste profiles could be targeted in molecule development. These advancements would contribute to more sophisticated computational approaches in flavor science and molecular gastronomy, potentially revolutionizing food product development and taste prediction accuracy.

## Supporting information

code to run our model

database used for training our model

## Declarations

### Ethics approval and consent to participate

Not applicable

## Acknowledgment

We thank Cristian Rojas, Davide Ballabio, Karen Pacheco Sarmiento, Elisa Pacheco Jaramillo, Mateo Mendoza, Fernando García for their generosity in sharing their data for research use.

## Author Contributions

SSDN Methodology, software, data analysis, visualization, writing - original draft. Developed and implemented the machine learning models, performed data preprocessing and analysis, created visualizations, and wrote the initial manuscript draft.

VR Conceptualization, supervision, project administration. Conceived and designed the original idea and research direction. Provided critical feedback and guidance throughout the project. Supervised the research process and reviewed the final manuscript.

Both authors discussed the results and contributed to the final manuscript.

## Data Availability

The ChemTastesDB dataset used in this study is freely available at https://doi.org/10.5281/zenodo.5747393. All code used for analysis has been implemented using open-source libraries including Mordred (v1.2.0), RDkit (v2021.09.4), scikit-learn, and iterative-stratification (v0.1.7). The analysis scripts are available upon reasonable request to the corresponding author.

## SUPPORTING INFORMATION

Mordred featurization produced a total of 1521 features, out of which 911 features had at least one missing value. We dropped feature columns with more than 40 % missing values, which reduced the number of features to 1436. The remaining missing values were imputed using the mean strategy by a KNN imputer.

Reducing the number of Dimensions was attempted using PCA, but even while retaining 99.99 % variance, the model’s f1_score dropped by up to 0.4. This is a drastic drop given the model performance for the task was very low to begin with, so we opted not to transform our data using PCA.

Expanding the dataset was attempted using APIs of online chemical databases like PubChem, ChemSpider etc. But due to very poor quality of taste vocabulary and the rarity of taste description being available for molecules in the database, we stuck with our expertly labeled taste dataset.

Fingerprints were generated using the openly licensed RDkit tool, which is popularly used in cheminformatics. It was also used to visualize our Smile strings.

Iterative stratified splitting was done using the scikit-multilearn library, and further, a custom iterative stratified splitter was passed during cross-validation.

Networkx and scikit-multilearn libraries were used to visualize our graph communities during community detection.

**Fig 1.**
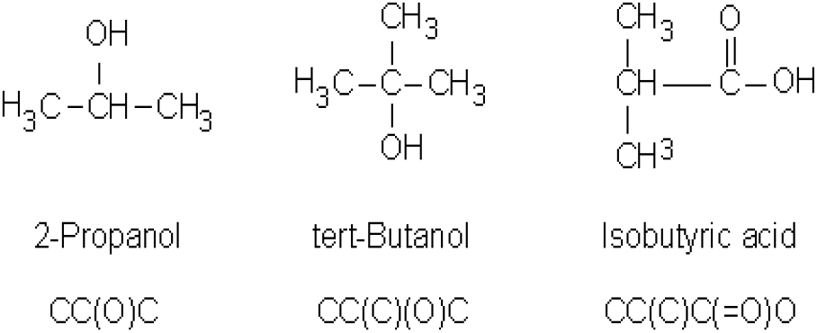
Smile notation for branched chains with their structures

**Fig 2.**
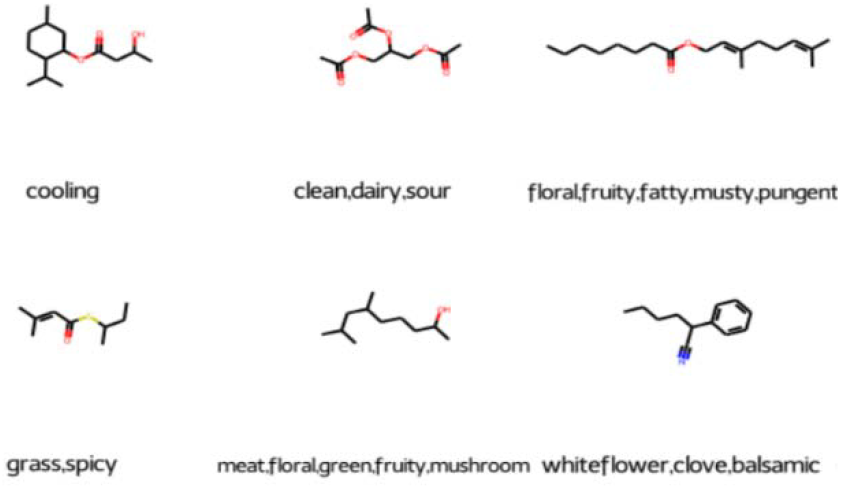
Molecules generated and visualized from smile strings in the dataset

SMILE (simplified molecular-input line-entry system) format, which is a specification in the form of a line notation for describing the structure of chemical species using short ASCII strings. This encoding into a string of characters is done to make the data machine-readable. A unique molecular structure can be reproduced from each smile string. For ease of data visualization and exploration, we chose only to retain Firmenich’s vocabulary, thus dropping labels from the PMP database which were non-intersecting.

### Mordred Featurization

A molecular descriptor is defined as the final result of a logical and mathematical procedure, which transforms chemical information encoded within a symbolic representation of a molecule into a useful number or the result of some standardized experiment. Various molecular-descriptor-calculation software programs have been developed-Mordred, Dragon, and ChemoPy, to name a few.. It computes thousands of handcrafted features like number of carbon atoms, number of hydrogen atoms, ring count, etc, all of which transcribe the information about our molecule.

### Fingerprinting

The molecular fingerprint is just another way of numerically representing a molecule. The bit-like patterns generated by the fingerprint indicate the absence or presence of certain substructures/fragments within a molecule. A molecular fingerprint characterizes the pattern, but the meaning of any particular bit is not well defined.

Molecular fingerprints represent a set of features derived from the structure of a molecule. The particular features calculated from the structure can be quite arbitrary and depend on the topology of the chemical graph or even a 3D conformation. Different fingerprint schemes emphasize different molecular attributes according to the design philosophy of the fingerprint system. The fundamental idea is to encapsulate certain properties directly or indirectly in the fingerprint and then use the fingerprint as a surrogate for the chemical structure.

The fingerprinting algorithm examines the molecule and generates fragments iteratively. Each fragment serves as a seed to a pseudo-random number generator (it is “hashed”), the output of which is a set of bits (0’s and 1’s, typically 4 or 5 bits per fragment); the set of bits thus produced is added (with a logical OR) to the fingerprint.

Types of fingerprints are differentiated by the manner of fragment generation: -

A. Path-Based Fingerprinting(daylight)-Fragments of the molecule are generated by following a path (usually linear) up to a certain number of bonds within the molecule.
B. Morgan Fingerprinting(circular) - Instead of linearly searching along each bond to generate fragments, morgan fingerprints generate fragments radially while varying the radius in each iteration.

**Fig.**
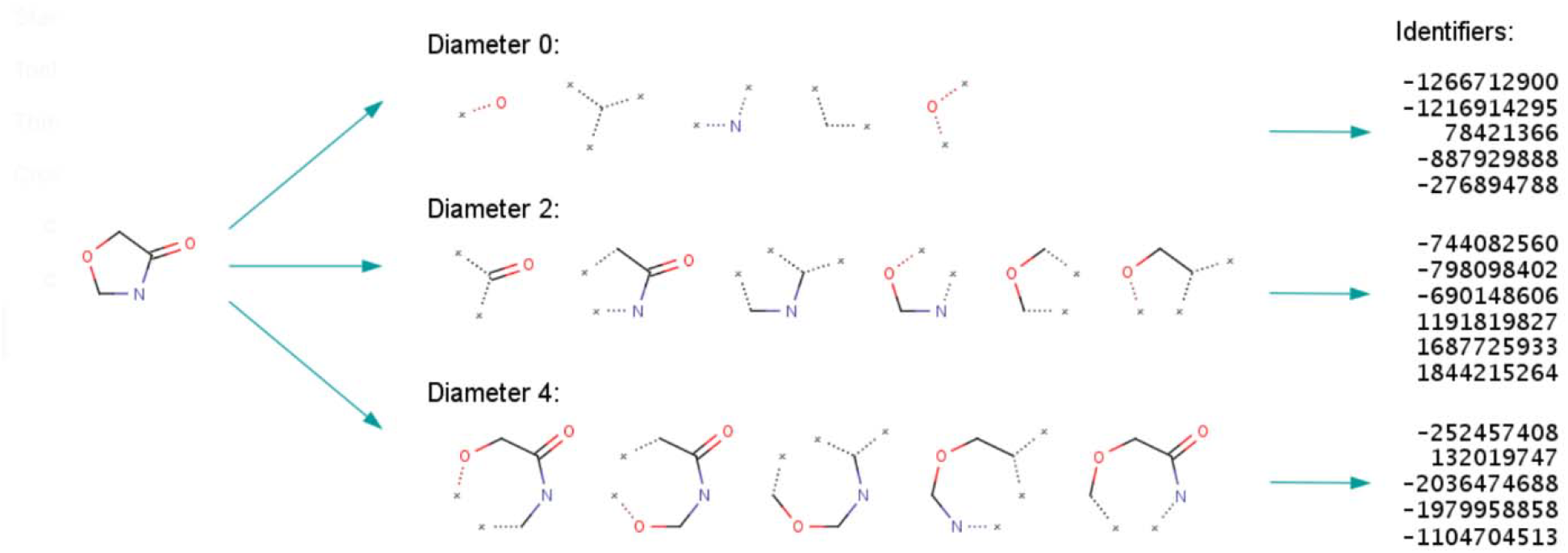
Morgan fingerprinting scheme in action with a varying radius of 0,1,2. All configurations of the 0^th^ radius are fed to a pseudo number generator which outputs the block of identifiers. These are further hashed to give a bit string. This is done for every radius, and the output is concatenated to the fingerprint

Multi-Label classification has proven to be very useful in fields like text categorization, Analysing protein properties and gene expression, and labeling of multimedia resources. The main difference between traditional and multi-label classification is in the output expected from trained models. Where a traditional classifier will return only one value, a multi-label one has to produce a vector of output values.

Problem transformation-based multi-label classification algorithms aim to convert the original dataset into one or more simpler datasets that can be delivered to traditional classification algorithms, while algorithmic adaptation techniques rely on modifying existing classification algorithms. The changes that must be introduced in the algorithms can be quite simple or really difficult, depending on the nature of the original method and also our data.

With the IRLbl metric, it is possible to know the imbalance level of one specific label. This is computed as the proportion between the number of appearances of the most common label and the considered label. Usually, a global assessment of the imbalance in the MLD is desired. This metric, named MeanIR, is calculated by averaging the IRLbl of all labels

#### List of abbreviations-

MLD’s: Multi-Label Datasets
SMILE: simplified molecular-input line-entry system

## Data and Software Availability

### DATASET

The dataset used here is **ChemTastesDB**(Cristian Rojas, Davide Ballabio, Karen Pacheco Sarmiento, Elisa Pacheco Jaramillo, Mateo Mendoza, Fernando García) which consists of a comprehensive collection of 2944 tastants, including both organic and inorganic chemical compounds. These tastants are categorized into various taste classes: the five basic tastes (sweet, bitter, umami, sour, and salty), as well as non-basic and other taste categories such as tasteless, non-sweet, multitaste, and miscellaneous.

### Software access

Mordred molecular descriptor calculator was used for generating handcrafted Mordred features for our molecules(https://pypi.org/project/mordred/)[version 1.2.0]

Morgan and path-based fingerprints were generated using Rdkit Toolkit (https://pypi.org/project/rdkit-pypi/) [version 2021.09.4] and this was also used for visualizing smile strings.

Fuzzywuzzy package[0.18.0] (https://pypi.org/project/fuzzywuzzy/) was used to find similar taste labels from two separate taste datasets so as to establish a consistent taste vocabulary

All machine learning models used during the course of this project have been imported from the scikit-learn open-source library(https://github.com/scikit-learn/scikit-learn)

Iterative-stratification package was used for stratified splitting into training and test set(https://pypi.org/project/iterative-stratification/) [version 0.1.7]

