## Supplementary material for "Predicting Molecular Taste: Multi-Label and Multi-Class Classification": code to run our model

### bxo7jbis3

March 25, 2025

#### 1 Run

[ ]:
```
!pip install scikit-multilearn==0.1.0
```

```
Collecting scikit-multilearn==0.1.0
  Downloading scikit_multilearn-0.1.0-py3-none-any.whl.metadata (6.0 kB)
Downloading scikit_multilearn-0.1.0-py3-none-any.whl (70 kB)
                         70.7/70.7 kB
3.4 MB/s eta 0:00:00
Installing collected packages: scikit-multilearn
Successfully installed scikit-multilearn-0.1.0
```

[ ]:
```
!pip install rdkit-pypi==2021.09.4
```

```
Collecting rdkit-pypi==2021.09.4
  Downloading rdkit_pypi-2021.9.4-cp310-cp310-manylinux_2_17_x86_64.manylinux201
4_x86_64.whl.metadata (2.6 kB)
Requirement already satisfied: numpy>=1.19 in /usr/local/lib/python3.10/dist-
packages (from rdkit-pypi==2021.09.4) (1.26.4)
Requirement already satisfied: Pillow in /usr/local/lib/python3.10/dist-packages
(from rdkit-pypi==2021.09.4) (11.0.0)
Downloading
rdkit_pypi-2021.9.4-cp310-cp310-manylinux_2_17_x86_64.manylinux2014_x86_64.whl
(20.8 MB)
                         20.8/20.8 MB
33.2 MB/s eta 0:00:00
Installing collected packages: rdkit-pypi
Successfully installed rdkit-pypi-2021.9.4
```

[ ]:
```
pip install pubchempy
```

```
Collecting pubchempy
  Downloading PubChemPy-1.0.4.tar.gz (29 kB)
  Preparing metadata (setup.py) … done
Building wheels for collected packages: pubchempy
  Building wheel for pubchempy (setup.py) … done
  Created wheel for pubchempy: filename=PubChemPy-1.0.4-py3-none-any.whl
size=13819
```

```
sha256=4996560fa5bb68d4038d4221f9472c85e169fac4eaf11cae8432ed4497cb9a4e
  Stored in directory: /root/.cache/pip/wheels/90/7c/45/18a0671e3c3316966ef7ed9a
d2b3f3300a7e41d3421a44e799
Successfully built pubchempy
Installing collected packages: pubchempy
Successfully installed pubchempy-1.0.4
```

[ ]:
```
!python --version
```

```
Python 3.10.12
```

[ ]:
```
import pandas as pd
import numpy as np
import os
import string
import sklearn
# to make this notebook's output stable across runs
np.random.seed(42)

# To plot pretty figures
%matplotlib inline
import matplotlib as mpl
import matplotlib.pyplot as plt
mpl.rc('axes', labelsize=14)
mpl.rc('xtick', labelsize=12)
mpl.rc('ytick', labelsize=12)

import sys
import os
import requests
import subprocess
import shutil
# sys.path.append("/content/drive/MyDrive/Summer_Project_Learning_To_Smell!/
↪Packages/scikit-multilearn-master/scikit-multilearn-master")
import skmultilearn
```

[ ]:
```
data=pd.read_csv("./taste_database.csv")

# Function to remove text in parentheses, convert taste string to lowercase,␣
↪and return a list
def convert_to_list_and_lowercase(taste):
    if pd.isna(taste):  # Check if the value is NaN
        return []  # Return an empty list if NaN

    # Remove text within parentheses
    taste = remove_parentheses(taste)
```

```
    # Convert to lowercase and split
    return [item.strip().lower() for item in str(taste).replace(',', ';').
 ↪replace('/', ';').split(';')]

# Function to remove text within parentheses
def remove_parentheses(s):
    while '(' in s and ')' in s:
        start = s.find('(')
        end = s.find(')', start)
        if end != -1:
            s = s[:start] + s[end + 1:]  # Slice out the part including␣
 ↪parentheses
        else:
            break
    return s.strip()

# Apply the function to the "Taste" column to create a list
data['Taste'] = data['Taste'].apply(convert_to_list_and_lowercase)

# Convert "Class taste" to lowercase and handle NaN if necessary
data['Class taste'] = data['Class taste'].apply(lambda x: x.lower() if␣
 ↪isinstance(x, str) else "unknownclass")

# Remove rows where "canonical SMILES" is not a string
data = data[data['canonical SMILES'].apply(lambda x: isinstance(x, str))]

# Display the first 100 rows of the DataFrame
data.head(100)
```

[ ]:
```
       ID                                               Name PubChem CID  \
0    0001                                     (-)-Haematoxylin      320930
1    0002                             (+)-4 -hydroxyhernandulcin      126862
2    0003                           (+)-Dihydroquercetin 3-acetate      442540
3    0004                                     (+)-Haematoxylin      442514
4    0005                                    (±)-chiro-inositol         892
..    ...                                                  ...         ...
95   0096  2-Cyanopyrid-5-yl guanidineacetic acid derivat…           *
96   0097  2-Cyanopyrid-5-yl guanidineacetic acid derivat…           *
97   0098  2-Cyanopyrid-5-yl guanidineacetic acid derivat…           *
98  0099^        2-Cyclopentyl-2-phenylsulfonylethanoic acid    82933891
99   0100                                     2-Deoxy-fructose    22858201

     CAS number                             canonical SMILES  \
0      517-28-2          Oc1cc2c(cc1O)C1c3ccc(c(c3OCC1(O)C2)O)O
1   145385-64-4                CC(C)=CCCC(C)(O)C1CC(O)C(=CC1=O)C
2    78834-97-6   CC(=O)OC1C(Oc2cc(cc(c2C1=O)O)O)c1ccc(c(c1)O)O
3      517-28-2          Oc1cc2c(cc1O)C1c3ccc(c(c3OCC1(O)C2)O)O
```

```
4     643-12-9                           OC1C(O)C(O)C(O)C(O)C1O
..         …                                                  …
95           *           OC(=O)CNC(NC1CCCCCCCC1)=Nc1ccc(nc1)C#N
96           *            OC(=O)CNC(NC1CCCCCCC1)=Nc1ccc(nc1)C#N
97           *  OC(=O)CNC(NC(C1=CC=CO1)c1ccccc1)=Nc1ccc(nc1)C#N
98           *                OC(=O)C(C1CCCC1)S(=O)(=O)c1ccccc1
99           *                                 OCC1OCC(O)C(O)C1O

                Taste Class taste  \
0              [sweet]   sweetness
1              [sweet]   sweetness
2              [sweet]   sweetness
3              [sweet]   sweetness
4              [sweet]   sweetness
..                 …         …
95             [sweet]   sweetness
96             [sweet]   sweetness
97             [sweet]   sweetness
98             [sweet]   sweetness
99  [low sweet, sweet]   sweetness

                              Reference_(cod)/[pp]
0             Arnoldi1995_((-)-1); Bassoli2001_(39)
1   Kinghorn1998_(2); Kinghorn2002_(7); Kinghorn20…
2   Bouysset2020_(175); Kinghorn2002_(26); Shallen…
3   Arnoldi1995_((+)-1); Arnoldi1996_(12); Bassoli…
4                            Shallenberger1993_[149]
..                                                …
95                               Nofre1993_(2)/[225]
96          Nofre1993_(5)/[225]; Nofre1993_(10)/[225]
97                              Nofre1993_(12)/[225]
98           Polański1993_(7)/[188]; Polański1997_(6)
99  Lichtenthaler1993_(57)/[45]; Shallenberger1993…

[100 rows x 8 columns]
```

[ ]:
```
train, valid, test = np.split(data.sample(frac=1), [int(0.6*len(data)), int(0.
↪8*len(data))])
```

```
/usr/local/lib/python3.10/dist-packages/numpy/core/fromnumeric.py:59:
FutureWarning: 'DataFrame.swapaxes' is deprecated and will be removed in a
future version. Please use 'DataFrame.transpose' instead.
  return bound(*args, **kwds)
```

[ ]:
```
train.head()
```

[ ]:
```
           ID                                               Name PubChem CID  \
565    0566                                 Compound (R)-(+)-9           *
1220   1221                                          Artemisin       65030
1677   1678                                          L-Leucine        6106
1577   1578                                     Hexethal sodium    23690440
2831   2832*  Sodium N-[3-chloro-5-(trifluoromethyl)phenyl]s…           *

     CAS number                            canonical SMILES  \
565           *        COc1ccc(cc1O)C1OC2CC3CCC2(CS1)C3(C)C
1220   481-05-0       CC1C2C(O)CC3(C)C=CC(=O)C(=C3C2OC1=O)C
1677    61-90-5                              CC(C)CC(N)C(O)=O
1577   144-00-3       [Na+].CCCCCCC1(CC)C(=O)NC(=NC1=O)[O-]
2831          *  [Na+].[O-]S(=O)(=O)Nc1cc(cc(c1)C(F)(F)F)Cl

                                   Taste    Class taste  \
565                              [sweet]      sweetness
1220                            [bitter]     bitterness
1677  [bitter, slightly bitter, non-sweet]     bitterness
1577                            [bitter]     bitterness
2831                         [non-sweet]  non-sweetness

                               Reference_(cod)/[pp]
565                               Bassoli2000_((+)-9)
1220                            Dagan-Wiener2019_(463)
1677  Belitz2009_[35]; Dagan-Wiener2019_(751); Glase…
1577                            Dagan-Wiener2019_(587)
2831                               Spillane2009b_(51A)
```

[ ]:
```python
valid.head()
```

[ ]:
```
           ID                               Name PubChem CID CAS number  \
1111   1112            4-Ethoxyphenylthiourea      853569   880-29-5
2800   2801*  Sodium N-(4-propylphenyl)sulfamate           *          *
1441   1442                            Diazepam        3016   439-14-5
2635   2636          1',6'-Di-O-acetate sucrose           *          *
2033   2034                        Sulfisoxazole        5344   127-69-5

                                   canonical SMILES        Taste  \
1111                            CCOc1ccc(cc1)NC(N)=S     [bitter]
2800                  [Na+].CCCc1ccc(cc1)NS([O-])(=O)=O  [non-sweet]
1441                CN1C(=O)CN=C(c2ccccc2)c2cc(ccc12)Cl     [bitter]
2635  CC(=O)OCC1OC(COC(C)=O)(OC2OC(CO)C(O)C(O)C2O)C(…  [non-sweet]
2033                  CC1=NOC(=C1C)NS(=O)(=O)c1ccc(cc1)N     [bitter]

        Class taste     Reference_(cod)/[pp]
1111     bitterness  Shallenberger1993_[278]
2800  non-sweetness       Spillane2009b_(33A)
```

```
1441     bitterness  Dagan-Wiener2019_(1244)
2635  non-sweetness           Hough1993a_[92]
2033     bitterness   Dagan-Wiener2019_(494)
```

[ ]:
```
test.head()
```

[ ]:
```
        ID                        Name PubChem CID  CAS number  \
933   0934               Superaspartame    18606782           *
928   0929  Suosan (N-glycine homolog)    19974425  67513-13-7
823   0824               Rebaudioside W           *           *
1531  1532      gamma,gamma'-dipyridyl       11107    553-26-4
1214  1215                     Arbutin      440936    497-76-7

                                       canonical SMILES     Taste Class taste  \
933   COC(=O)C(Cc1ccccc1)NC(=O)C(CC(O)=O)NC(=O)Nc1cc…   [sweet]   sweetness
928                     OC(=O)CNC(=O)Nc1ccc(cc1)N(=O)=O    [sweet]   sweetness
823   CC1OC(OC2C(OC(CO)C(O)C2OC2OC(CO)C(O)C(O)C2O)OC…   [sweet]   sweetness
1531                              c1cc(ccn1)-c1ccncc1  [bitter]  bitterness
1214                     OCC1OC(Oc2ccc(cc2)O)C(O)C(O)C1O  [bitter]  bitterness

                                  Reference_(cod)/[pp]
933   Belitz2009_[37442]; Bouysset2020_(312); DuBois…
928   Belitz2009_[439]; Shallenberger1993_[244]; Sha…
823                                 Soejarto2019_(26)
1531                           Dagan-Wiener2019_(192)
1214                           Dagan-Wiener2019_(103)
```

[ ]:
```
print(data["Class taste"])
print(data["Taste"])
```

```
0           sweetness
1           sweetness
2           sweetness
3           sweetness
4           sweetness
              …
2939    miscellaneous
2940    miscellaneous
2941    miscellaneous
2942    miscellaneous
2943    miscellaneous
Name: Class taste, Length: 2944, dtype: object
0                                 [sweet]
1                                 [sweet]
2                                 [sweet]
3                                 [sweet]
4                                 [sweet]
```

```
                            …
2939    [heating, pungent, and tingling]
2940                        [astringent]
2941    [heating, pungent, and tingling]
2942                           [cooling]
2943                           [cooling]
Name: Taste, Length: 2944, dtype: object
```

[ ]:
```
#Let's get the vocabulary for our odor descriptors
def get_vocab_taste(data):
  vocab=set()
  for x in data:
    for y in x:
      vocab.add(y)
  return vocab
```

[ ]:
```
#So we have a total of 109 unique classes of odors for our classification task
vocabt=get_vocab_taste(data["Taste"])
print(sorted(vocabt))
print(len(vocabt))
```

```
['acid', 'acrid', 'and tingling', 'astringent', 'barely sweet', 'bitter',
'burning', 'cooling', 'entirely bitter', 'extremely bitter', 'faint bitter',
'faintly bitter', 'feebly bitter', 'heating', 'highly bitter', 'highly sweet',
'hot burning', 'intensely bitter', 'intensely sweet', 'lacking sweet', 'less
sweet', 'like fresh walnut', 'low sweet', 'moderately bitter', 'neutral', 'non-
bitter', 'non-sweet', 'pungent', 'salty', 'scratchy', 'slightly bitter',
'slightly burning', 'slightly sweet', 'somewhat bitter', 'sour', 'strongly
bitter', 'sulphurous', 'sweet', 'sweetish', 'tasteless', 'umami', 'very bitter',
'very sweet', 'weak umami']
44
```

[ ]:
```
#Let's get the vocabulary for our odor descriptors
def get_vocab_classtaste(data):
  vocab=set()
  for x in data:
    vocab.add(x)
  return vocab
```

[ ]:
```
vocabct=get_vocab_classtaste(data["Class taste"])
print(sorted(vocabct))
print(len(vocabct))
```

```
['bitterness', 'miscellaneous', 'multitaste', 'non-sweetness', 'saltiness',
'sourness', 'sweetness', 'tastelessness', 'umaminess']
9
```

[ ]:
```
#Getting first 5 smile strings from our pandas table
List=[x for x in data["canonical SMILES"][:10]]
```

[ ]:
```
# Rdkit is an Open source toolkit for cheminformatics and machine learning with loads of functionalities
# We'll be using it here to render a 2d image of the molecule.
import rdkit
import rdkit.Chem as Chem
from rdkit.Chem import rdFMCS
from matplotlib import colors
from rdkit.Chem import Draw
m=[Chem.MolFromSmiles(x)for x in List]
leg=["".join(x) for x in data["Class taste"][::10][::-1]]
Draw.MolsToGridImage(m,molsPerRow=5,subImgSize=(200,200),legends=leg)
```

[ ]:

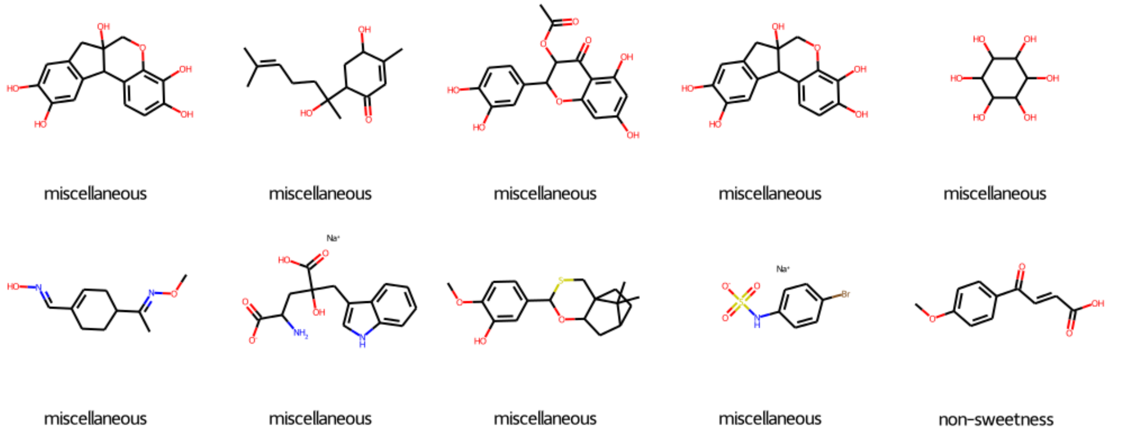


[ ]:
```
# Simply accessing the SMILES string as a regular string
smiles_string = data["canonical SMILES"][96]
print(smiles_string)   # This will print: OC(=O)CNC(NC1CCCCCCC1)=Nc1ccc(nc1)C#N
```

```
OC(=O)CNC(NC1CCCCCCC1)=Nc1ccc(nc1)C#N
```

[ ]:
```
from rdkit import Chem

# Convert SMILES string to a molecular object
smiles_string = data["canonical SMILES"][96]
print(type(smiles_string))
molecule = Chem.MolFromSmiles(smiles_string)

# Display some properties of the molecule (for example, its molecular weight)
from rdkit.Chem import Descriptors
print("Molecular Weight:", Descriptors.MolWt(molecule))
```

<class 'str'>
Molecular Weight: 329.40399999999994

```
[ ]: from collections import defaultdict
new_data=defaultdict(list) #leff dataset processed to fit the schema of
#fermenich dataset while filtering duplicate odorant molecules
for s in data.index:
  # if(data["smiles"][s] not in list(data["SMILES"])):
    new_data[data["canonical SMILES"][s]]=data["Class taste"][s]
```

```
[ ]: new_data.items()
```

```
[ ]: dict_items([('Oc1cc2c(cc1O)C1c3ccc(c(c3OCC1(O)C2)O)O', 'sweetness'),
('CC(C)=CCCC(C)(O)C1CC(O)C(=CC1=O)C', 'sweetness'),
('CC(=O)OC1C(Oc2cc(cc(c2C1=O)O)O)c1ccc(c(c1)O)O', 'sweetness'),
('OC1C(O)C(O)C(O)C(O)C1O', 'sweetness'), ('CON=C(C)C1CCC(=CC1)C=NO',
'sweetness'), ('[Na+].NC(CC(O)(CC1=CNc2ccccc12)C(O)=O)C([O-])=O', 'sweetness'),
('COc1ccc(cc1O)C1OC2CC3CCC2(CS1)C3(C)C', 'tastelessness'),
('[Na+].[O-]S(=O)(=O)Nc1ccc(cc1)Br', 'sweetness'),
('COc1ccc(cc1)C(=O)C=CC(O)=O', 'sweetness'), ('CC1=CCC(=CC1)C=NO', 'sweetness'),
('COC(C)C1=CC=C(CC1)C=NO', 'sweetness'), ('COC(C)C1=CCC(=CC1)C=NO',
'sweetness'), ('COc1c(cc2c(c1O)C(=O)C(OC(C)=O)C(O2)c1ccc(cc1)O)O', 'sweetness'),
('CCC(O)CO', 'sweetness'), ('CC(O)CO', 'sweetness'),
('COc1ccc(cc1O)C1OCc2c(cccc2O1)O', 'sweetness'), ('COc1ccc(cc1O)C1OCc2ccccc2O1',
'sweetness'), ('CC1=CC=C(C)CC1', 'sweetness'), ('[Na+].[O-]S(=O)(=O)NC1CSCSC1',
'sweetness'), ('OCC1OC(OC2(CCl)OC(CCl)C(Cl)C2O)C(O)C(O)C1O', 'sweetness'),
('OCC(O)C1OCC(O)C1O', 'sweetness'), ('OCC1OCC(O)C1O', 'sweetness'),
('COc1ccc(cc1O)C1COc2ccccc2O1', 'sweetness'),
('OCC1OC(OC2(CCl)OC(CO)C(Cl)C2O)C(O)C(O)C1O', 'sweetness'),
('OCC1OCC(O)C(O)C1O', 'sweetness'),
('[Na+].CC1C(NS([O-])(=O)=O)C(=O)N(N1C)c1ccccc1', 'sweetness'),
('OCC1OC(OC2(CCl)OC(CCl)C(O)C2O)C(O)C(O)C1O', 'sweetness'),
('OC1C(O)C(O)(CCl)OC1CCl', 'sweetness'),
('COCC1OC(COC)(OC2OC(CO)C(O)C(O)C2O)C(O)C1O', 'sweetness'),
('CSCC1OC(O)(CSC)C(O)C1O', 'sweetness'),
('CC1(CCCC2C1C(C(O)=O)c1ccccc21)C(O)=O', 'sweetness'), ('CC(=O)C1CCC(=CC1)C=NO',
'sweetness'), ('NC1(CCC1)C(O)=O', 'sweetness'),
('OCC1OC(OC2(CCl)OC(CO)C(O)C2O)C(O)C(O)C1O', 'sweetness'), ('ON=CC1=CCCCC1',
'sweetness'), ('ON=CC1=CCCC1', 'sweetness'),
('CC1(OC(CO)C(O)C1O)OC1OC(CO)C(O)C(O)C1O', 'sweetness'),
('OC(=O)CNC(NC1CCCCCCC1)=Nc1ccc2c(c1)NN=C2', 'sweetness'),
('CC(NC(NCC(O)=O)=Nc1cnc2c(c1)NN=C2)c1ccccc1', 'sweetness'), ('CC1=CC=CCC1',
'sweetness'), ('CC1=CCCCC1', 'sweetness'),
('COCC1(OC(CO)C(O)C1O)OC1OC(CO)C(O)C(O)C1O', 'sweetness'),
('CC1CC(=C(C)CC1(C(O)=O)S(=O)(=O)c1ccccc1)C', 'sweetness'),
('CC1=C(C)CC(CC1)(C(O)=O)S(=O)(=O)c1ccccc1', 'sweetness'),
('CC1CC(=CCC1(C(O)=O)S(=O)(=O)c1ccccc1)C', 'sweetness'),
```

('CC1=CCC(CC1)(C(O)=O)S(=O)(=O)c1ccccc1', 'sweetness'),
('OC(=O)c1ccccc1C(=O)c1ccc(c(c1)O)O', 'sweetness'),
('COc1c(cc2c(c1O)C(=O)C(O)C(O2)c1ccc(c(c1)O)O)O', 'sweetness'),
('COc1c(cc2c(c1O)C(=O)C(OC(C)=O)C(O2)c1ccc(c(c1)O)O)O', 'sweetness'),
('COc1ccc(cc1O)C(=O)c1ccccc1C(O)=O', 'sweetness'),
('COc1ccc(cc1O)OCc1ccccc1C(O)=O', 'sweetness'),
('COc1ccc(cc1O)C1OC(=O)c2ccccc2O1', 'sweetness'), ('COc1ccc(cc1O)C1Oc2ccccc2O1',
'sweetness'), ('COc1ccc(cc1O)C1OC(=O)c2ccccc2S1', 'sweetness'),
('COc1ccc(cc1O)C1Oc2ccccc2S1', 'sweetness'), ('COc1ccc(cc1O)C1CSc2ccccc2S1',
'sweetness'), ('COc1ccc(cc1O)C1CSc2cc(ccc2O1)O', 'sweetness'),
('COc1ccc(cc1O)C1CSc2ccccc2O1', 'sweetness'),
('COc1ccc(cc1O)C1CS(=O)(=O)c2ccccc2O1', 'sweetness'),
('COc1ccc(cc1O)C1SCc2c(cccc2S1)O', 'sweetness'),
('COc1ccc(cc1O)C1OCc2c(cccc2S1)O', 'sweetness'), ('COc1ccc(cc1O)C1CCc2ccccc2S1',
'sweetness'), ('COc1ccc(cc1O)C1SCc2ccccc2S1', 'tastelessness'),
('COc1ccc(cc1O)C1Sc2ccccc2S1', 'sweetness'), ('COc1ccc(cc1O)C1Oc2ccccc2CS1',
'sweetness'), ('COc1ccc(cc1S)C1OCc2ccccc2S1', 'sweetness'),
('OC(=O)c1ccccc1C(=O)c1ccc(cc1)Cl', 'sweetness'),
('CCOc1ccc(cc1)C(=O)c1ccccc1C(O)=O', 'sweetness'),
('CCc1ccc(cc1)S(=O)(=O)C(C(C)C)C(O)=O', 'sweetness'),
('OC(=O)c1ccccc1C(=O)c1ccc(cc1)O', 'sweetness'),
('COc1ccc(cc1)C(=O)c1ccccc1C(O)=O', 'sweetness'),
('COc1ccc(cc1)S(=O)c1ccccc1C(O)=O', 'sweetness'),
('Cc1ccc(cc1)C(=O)c1ccccc1C(O)=O', 'sweetness'),
('CC(C)C(C(O)=O)S(=O)(=O)c1ccc(cc1)C', 'sweetness'),
('CCC(C)C(C(O)=O)S(=O)(=O)c1ccccc1', 'sweetness'),
('CC1(C)CC1(C(O)=O)S(=O)(=O)c1ccc(cc1)O', 'sweetness'),
('CCOc1ccc(cc1O)CCC(=O)c1ccccc1O', 'sweetness'),
('COc1ccc(cc1O)CCC(=O)c1ccccc1O', 'sweetness'),
('COc1ccc(cc1O)CCC(=O)c1c(cc(c(c1O)C(O)=O)O)O', 'sweetness'),
('[Na+].[O-]S(=O)(=O)Nc1cc(nc(c1)C(F)(F)F)C(F)(F)F', 'sweetness'),
('CC(C)(OCC(O)=O)C1CCC(=CC1)C=NO', 'sweetness'),
('Oc1ccc(cc1OC(=O)Oc1ccc(cc1)N(=O)=O)C1OCc2ccccc2S1', 'sweetness'),
('COc1ccc(cc1OC(=O)Nc1ccc(cc1)N(=O)=O)C1OCc2ccccc2O1', 'sweetness'),
('COc1ccc(cc1OC(=O)Nc1ccc(cc1)N(=O)=O)C1OCc2ccccc2S1', 'sweetness'),
('O=C(COc1cccc(c1)C1OCc2ccccc2S1)c1ccc(cc1)N(=O)=O', 'sweetness'),
('CN(C)c1ccc(cc1)C(=O)c1ccccc1C(O)=O', 'sweetness'),
('CC(C)OC(=O)C(C)NC(=O)C(N)C(O)=O', 'tastelessness'), ('Nc1ccccc1C(O)=O',
'sweetness'), ('OC(=O)CNC(NC1CCCCCCC1)=Nc1ccc(nc1)Cl', 'sweetness'),
('OC(=O)CNC(NC1CCCCCCCC1)=Nc1ccc(nc1)C#N', 'sweetness'),
('OC(=O)CNC(NC1CCCCCCC1)=Nc1ccc(nc1)C#N', 'sweetness'),
('OC(=O)CNC(NC(C1=CC=CO1)c1ccccc1)=Nc1ccc(nc1)C#N', 'sweetness'),
('OC(=O)C(C1CCCC1)S(=O)(=O)c1ccccc1', 'sweetness'), ('CCOc1ccc(cc1O)CCc1ccccc1',
'sweetness'), ('COc1ccc(cc1O)CC(=O)c1ccccc1O', 'sweetness'),
('COc1ccc(cc1O)CCCc1ccccc1', 'sweetness'), ('COc1ccc(cc1O)CSc1ccccc1',
'sweetness'), ('COc1ccc(cc1O)CCc1ccc2c(c1)OCO2', 'sweetness'),
('COc1ccc(cc1O)CCc1ccccc1OC', 'sweetness'), ('COc1ccc(cc1O)CCc1ccccc1C',

'sweetness'), ('COc1cccc(c1)CCc1ccc(c(c1)O)OC', 'sweetness'),
('COc1ccc(cc1O)CCc1cccc(c1)C', 'sweetness'), ('CC=C(C)C', 'sweetness'),
('CC(NC(NCC(O)=O)=Nc1ccc2ccccc2c1)c1ccccc1', 'sweetness'),
('CC(C)C(C)(C(O)=O)S(=O)(=O)c1ccccc1', 'sweetness'),
('CC(C)C(C(O)=O)S(=O)(=O)c1ccccc1', 'sweetness'),
('CCC(C(O)=O)S(=O)(=O)c1ccccc1', 'sweetness'),
('CCOC(=O)c1ccc(cc1)NC(=S)NC(CC(O)=O)C(=O)NC(Cc1ccccc1)C(=O)OC', 'sweetness'),
('CCOC(=O)c1ccc(cc1)NC(=O)NC(CC(O)=O)C(=O)NC(Cc1ccccc1)C(=O)OC', 'sweetness'),
('CC(NC(=O)C(N)CC(O)=O)NC(=O)C12CC3CC(CC(C3)C1)C2', 'sweetness'),
('COc1ccc(c(c1O)C=Cc1ccc(cc1)N(=O)=O)C1OCc2ccccc2S1', 'sweetness'),
('OC(=O)CC(NC(=O)Nc1ccc(cc1)C#N)c1ccc2c(c1)OCO2', 'sweetness'),
('NC(=O)c1c(cc(c(c1Br)CCC(O)=O)Br)Br', 'sweetness'),
('COc1ccc(cc1O)C1Cc2ccccc2CO1', 'sweetness'),
('COc1ccc(cc1O)CCC(=O)c1cccc(c1)C(O)=O', 'sweetness'),
('COc1ccc(cc1O)C1COc2ccccc2S1', 'sweetness'), ('COc1ccc(cc1O)C1COc2ccccc2C1',
'sweetness'), ('NC(CC(=O)c1ccc(cc1N)Cl)C(O)=O', 'sweetness'),
('Oc1cccc(c1)C1OCc2ccccc2S1', 'sweetness'), ('[Na+].CC(C)(C)CCNS([O-])(=O)=O',
'sweetness'), ('Oc1ccc(cc1O)C1OCc2ccccc2S1', 'sweetness'),
('CC(NC(NCC(O)=O)=Nc1ccc2c(c1)OCO2)c1ccccc1', 'sweetness'),
('[Na+].[O-]S(=O)(=O)Nc1ncc(cc1Br)Br', 'sweetness'),
('CC(NC(NCC(O)=O)=Nc1cc(cc(c1)Cl)Cl)c1ccccc1', 'sweetness'),
('OC(=O)CNC(NCc1ccccc1)=Nc1cc(cc(c1)Cl)Cl', 'sweetness'),
('CC(NC(NCC(O)=O)=Nc1cc(cc(c1)Cl)Cl)C1CCCCC1', 'sweetness'),
('OC(=O)CNC(NC1CCCCCCC1)=Nc1cc(cc(c1)Cl)Cl', 'sweetness'),
('OC(=O)CNC(NCC1CCCCC1)=Nc1cc(cc(c1)Cl)Cl', 'sweetness'),
('OC(=O)CNC(Nc1cccc2ccccc12)=Nc1cc(cc(c1)Cl)Cl', 'sweetness'),
('OC(=O)CNC(NC1CCCCCC1)=Nc1cc(cc(c1)Cl)Cl', 'sweetness'),
('CCCCC(C)NC(NCC(O)=O)=Nc1cc(cc(c1)Cl)Cl', 'sweetness'),
('OC(=O)CNC(Nc1cc(cc(c1)Cl)Cl)=Nc1cc(cc(c1)Cl)Cl', 'sweetness'),
('CCOc1ccc(cc1O)C1CC(=O)c2c(cccc2O1)O', 'sweetness'),
('COc1ccc(cc1O)C1CC(=O)c2c(cccc2O1)O', 'sweetness'),
('CCCOc1ccc(cc1O)C1CC(=O)c2c(cccc2O1)O', 'sweetness'),
('CCOC(=O)C(NC(=O)C(CC(O)=O)NC(=O)Nc1ccc(cc1)N(=O)=O)C(=O)OCC', 'sweetness'),
('COC(=O)C(NC(=O)C(CC(O)=O)NC(=O)Nc1ccc(cc1)N(=O)=O)C(=O)OC', 'sweetness'),
('COC(=O)C(NC(=O)C(CC(O)=O)NC(=O)Nc1ccc(cc1)N(=O)=O)C1CCCCC1', 'sweetness'),
('CCOC(=O)C(C)NC(=O)C(CC(O)=O)NC(=O)Nc1ccc(cc1)N(=O)=O', 'sweetness'),
('COC(=O)C(Cc1ccccc1)NC(=O)C(CC(O)=O)NC(=O)Nc1ccc(cc1)C(=O)OC', 'sweetness'),
('CNS(=O)(=O)c1ccc(cc1)NC(=O)NC(CC(O)=O)C(=O)NC(Cc1ccccc1)C(=O)OC',
'sweetness'), ('COC(=O)C(Cc1ccccc1)NC(=O)C(CC(O)=O)NC(=S)Nc1ccc(cc1)N(=O)=O',
'sweetness'), ('COC(=O)C(Cc1ccccc1)NC(=O)C(CC(O)=O)NC(=O)Nc1ccc(cc1)N(=O)=O',
'sweetness'), ('CCCCC(NC(=O)C(CC(O)=O)NC(=O)Nc1ccc(cc1)N(=O)=O)C(=O)OC',
'sweetness'), ('CCCC(NC(=O)C(CC(O)=O)NC(=O)Nc1ccc(cc1)N(=O)=O)C(=O)OC',
'sweetness'), ('COC(=O)C(C)NC(=O)C(CC(O)=O)NC(=O)Nc1ccc(cc1)N(=O)=O',
'sweetness'), ('COC(=O)C(CC1CCCCC1)NC(=O)C(CC(O)=O)NC(=O)Nc1ccc(cc1)N(=O)=O',
'sweetness'), ('OC(=O)CCNC(=O)Oc1ccc(cc1)N(=O)=O', 'sweetness'),
('COC(=O)C(Cc1ccccc1)NC(=O)C(CC(O)=O)NC(=O)Nc1ccc(cc1)C(C)=O', 'sweetness'),
('NC(=O)c1ccc(cc1)NC(=O)NC(CC(O)=O)c1cccc(c1)O', 'sweetness'),

```
('NC(=O)c1ccc(cc1)NC(=O)NC(CC(O)=O)c1cccnc1', 'sweetness'),
('CC(NC(=O)C(CC(O)=O)NC(=S)Nc1ccc(cc1)C#N)C(=O)NC(C1CC1)C1CC1', 'sweetness'),
('CC(NC(=O)C(CC(O)=O)NC(=S)Nc1ccc(cc1)C#N)c1ccccc1', 'sweetness'),
('OC(=O)CC(NC(=S)Nc1ccc(cc1)C#N)c1ccccc1', 'sweetness'),
('CC(NC(=O)C(CC(O)=O)NC(=O)Nc1ccc(cc1)C#N)C(=O)NC(C1CC1)C1CC1', 'sweetness'),
('OC(=O)CC(NC(=O)Nc1ccc(cc1)C#N)c1ccc2c(c1)CCC2', 'sweetness'),
('OC(=O)CC(NC(=O)Nc1ccc(cc1)C#N)c1ccccc1N(=O)=O', 'sweetness'),
('OC(=O)CC(NC(=O)Nc1ccc(cc1)C#N)c1cccc(c1)O', 'sweetness'),
('COc1cccc(c1)C(CC(O)=O)NC(=O)Nc1ccc(cc1)C#N', 'sweetness'),
('OC(=O)CC(NC(=O)Nc1ccc(cc1)C#N)c1cccc(c1)N(=O)=O', 'sweetness'),
('CCc1ccc(cc1)C(CC(O)=O)NC(=O)Nc1ccc(cc1)C#N', 'sweetness'),
('OC(=O)CC(NC(=O)Nc1ccc(cc1)C#N)c1ccc(cc1)O', 'sweetness'),
('COc1ccc(cc1)C(CC(O)=O)NC(=O)Nc1ccc(cc1)C#N', 'sweetness'),
('OC(=O)CC(NC(=O)Nc1ccc(cc1)C#N)c1ccc(cc1)N(=O)=O', 'sweetness'),
('OC(=O)CC(NC(=O)Nc1ccc(cc1)C#N)c1ccc2ccccc2c1', 'sweetness'),
('OC(=O)CC(NC(=O)Nc1ccc(cc1)C#N)c1cnc2ccccc2c1', 'sweetness'),
('COC(=O)C(NC(=O)C(CC(O)=O)NC(=O)Nc1ccc(cc1)C#N)C(F)(F)F', 'sweetness'),
('COC(=O)C(C)NC(=O)C(CC(O)=O)NC(=O)Nc1ccc(cc1)C#N', 'sweetness'),
('CC(NC(=O)C(CC(O)=O)NC(=O)Nc1ccc(cc1)C#N)c1ccccc1', 'sweetness'),
('COC(=O)C(NC(=O)C(CC(O)=O)NC(=O)Nc1ccc(cc1)C#N)c1ccccc1', 'sweetness'),
('COC(=O)C(CO)NC(=O)C(CC(O)=O)NC(=O)Nc1ccc(cc1)C#N', 'sweetness'),
('CCCOC(=O)C(CO)NC(=O)C(CC(O)=O)NC(=O)Nc1ccc(cc1)C#N', 'sweetness'),
('COC(=O)C(NC(=O)C(CC(O)=O)NC(=O)Nc1ccc(cc1)C#N)C(=O)OC1C(C)(C)C2CCC1(C)C2',
'sweetness'), ('OC(=O)CC(Cc1ccccc1)NC(=O)Nc1ccc(cc1)C#N', 'sweetness'),
('OC(=O)CC(CCc1ccccc1)NC(=O)Nc1ccc(cc1)C#N', 'sweetness'),
('COC(=O)C(Cc1ccccc1)NC(=O)C(CC(O)=O)NC(=O)Nc1ccc(cc1)F', 'sweetness'),
('CCCOC(=O)C(C)NC(=O)C(CC(O)=O)NC(=O)Nc1ccc(cc1)N(=O)=O', 'sweetness'),
('NC(=O)c1ccc(cn1)NC(=O)NC(CC(O)=O)c1cccnc1', 'sweetness'),
('NC(=O)c1ccc(cn1)NC(=O)NC(CC(O)=O)c1ccccc1', 'sweetness'),
('OC(=O)CC(NC(=O)Nc1ccc(nc1)C#N)c1cccnc1', 'sweetness'),
('CC(O)C(COC(C)=O)NC(=O)C(N)CC(O)=O', 'sweetness'),
('CC(=O)OCC(CO)NC(=O)C(N)CC(O)=O', 'sweetness'),
('CC(C)C(COC(C)=O)NC(=O)C(N)CC(O)=O', 'sweetness'),
('CCCC(COC(C)=O)NC(=O)C(N)CC(O)=O', 'sweetness'),
('CC(COC(C)=O)NC(=O)C(N)CC(O)=O', 'sweetness'), ('CC(=O)OCCNC(=O)C(N)CC(O)=O',
'sweetness'), ('COC(=O)C(COC(C)=O)NC(=O)C(N)CC(O)=O', 'sweetness'),
('CC(NC(=O)C(N)CC(O)=O)C(=O)NC(C)(C1CC1)C1CC1', 'sweetness'),
('CC(NC(=O)C(N)CC(O)=O)C(=O)NC(C)(C(C)(C)C)C(C)(C)C', 'sweetness'),
('CC(NC(=O)C(N)CC(O)=O)C(=O)NC1C(C)(C)S(=O)(=O)C1(C)C', 'sweetness'),
('CC(NC(=O)C(N)CC(O)=O)C(=O)NC1C(C)(C)S(=O)C1(C)C', 'sweetness'),
('CC(NC(=O)C(N)CC(O)=O)C(=O)NC1C(C)(C)CC1(C)C', 'sweetness'),
('CC(NC(=O)C(N)CC(O)=O)C(=O)NC1C(C)(C)CCCC1(C)C', 'sweetness'),
('CC(COC(=O)C(C)(C)C)NC(=O)C(N)CC(O)=O', 'sweetness'),
('CC(C)C(C)(NC(=O)C(C)NC(=O)C(N)CC(O)=O)C(C)C', 'sweetness'),
('CC(NC(=O)C(N)CC(O)=O)NC(=O)C1C(C)CCCC1C', 'sweetness'),
('COC(=O)C(NC(=O)C(N)CC(O)=O)C(=O)OC1C(C)CCCC1C', 'sweetness'),
('CC(NC(=O)C(N)CC(O)=O)C(=O)NC(C)(C1CC1)C(C)(C)C', 'sweetness'),
```

```
('CC(NC(=O)C(N)CC(O)=O)C(=O)NC(C)(C)C1CC1', 'sweetness'),
('CCC(CC)C(=O)OCC(C)NC(=O)C(N)CC(O)=O', 'sweetness'),
('CCC(C)C(=O)OCC(C)NC(=O)C(N)CC(O)=O', 'sweetness'),
('CC(COC(=O)C(C)=C)NC(=O)C(N)CC(O)=O', 'sweetness'),
('CCC(COC(=O)C(C)C)NC(=O)C(N)CC(O)=O', 'sweetness'),
('CC(C)C(=O)OCC(C)NC(=O)C(N)CC(O)=O', 'sweetness'),
('CC(NC(=O)C(N)CC(O)=O)C(=O)NC1C(C)(C)C(O)C1(C)C', 'sweetness'),
('CC(C)CC(=O)OCC(C)NC(=O)C(N)CC(O)=O', 'sweetness'),
('CCC(C)(CC)NC(=O)C(C)NC(=O)C(N)CC(O)=O', 'sweetness'),
('CCOC(=O)OC(=O)C(NC(=O)C(N)CC(O)=O)C(=O)OC', 'sweetness'),
('CCOC(=O)C(NC(=O)C(N)CC(O)=O)C(=O)OC1CCCCC1C', 'sweetness'),
('CCC(C)C(=O)OCC(CO)NC(=O)C(N)CC(O)=O', 'sweetness'),
('CC(C)C(=O)OCC(CO)NC(=O)C(N)CC(O)=O', 'sweetness'),
('CC(C)CC(=O)OCC(CO)NC(=O)C(N)CC(O)=O', 'sweetness'),
('COC(=O)C(NC(=O)C(N)CC(O)=O)C(=O)OC1CC(C)CC(C)(C)C1', 'sweetness'),
('COC(=O)C(NC(=O)C(N)CC(O)=O)C(=O)OC1CCCCC1C', 'sweetness'),
('COC(=O)C(COC(=O)C(C)C)NC(=O)C(N)CC(O)=O', 'sweetness'),
('COC(=O)C(NC(=O)C(N)CC(O)=O)C(=O)OC1CCCC(C)C1', 'sweetness'),
('COC(=O)C(NC(=O)C(N)CC(O)=O)C(=O)OC1CCC(C)CC1', 'sweetness'),
('CC(C)C(=O)OCCNC(=O)C(N)CC(O)=O', 'sweetness'),
('CC(C)CC(=O)OCCNC(=O)C(N)CC(O)=O', 'sweetness'),
('COC(=O)C(COC(=O)C(C)(C)C)NC(=O)C(N)CC(O)=O', 'sweetness'),
('CC(C)C(COC(=O)C(C)C)NC(=O)C(N)CC(O)=O', 'sweetness'),
('COC(=O)CCC(NC(=O)C(N)CC(O)=O)C(=O)OC', 'sweetness'),
('COC(=O)C(C)CC(NC(=O)C(N)CC(O)=O)C(=O)OC', 'sweetness'),
('CCOC(=O)C(NC(=O)C(N)CC(O)=O)C(=O)OC(=O)OC(C)(C)C', 'sweetness'),
('CCCCOC(=O)C(C)NC(=O)C(N)CC(O)=O', 'sweetness'),
('CCCCOC(=O)C(CO)NC(=O)C(N)CC(O)=O', 'sweetness'),
('CC(CC1CCCCC1)NC(=O)C(N)CC(O)=O', 'sweetness'),
('CCCCC(NC(=O)C(N)CC(O)=O)C(=O)OCC', 'sweetness'),
('CCOC(=O)C(NC(=O)C(N)CC(O)=O)C(C)O', 'sweetness'),
('CCOC(=O)C(CO)NC(=O)C(N)CC(O)=O', 'sweetness'),
('CC(NC(=O)C(N)CC(O)=O)C(O)c1ccccc1', 'sweetness'),
('COC(=O)C(NC(=O)C(N)CC(O)=O)C(C)c1ccccc1', 'sweetness'),
('CCC(NC(=O)C(N)CC(O)=O)C(=O)OC', 'sweetness'),
('CCCCCC(NC(=O)C(N)CC(O)=O)C(=O)OC', 'sweetness'),
('CCCCC(NC(=O)C(N)CC(O)=O)C(=O)OC', 'sweetness'),
('CCCCCCC(NC(=O)C(N)CC(O)=O)C(=O)OC', 'sweetness'),
('CCCC(NC(=O)C(N)CC(O)=O)C(=O)OC', 'sweetness'),
('COC(=O)C(CCC(C)C)NC(=O)C(N)CC(O)=O', 'sweetness'),
('CCCOC(=O)C(CC)NC(=O)C(N)CC(O)=O', 'sweetness'),
('CCCOC(=O)C(CCC)NC(=O)C(N)CC(O)=O', 'sweetness'),
('CC(COc1ccccc1)NC(=O)C(N)CC(O)=O', 'sweetness'),
('CC(C)(Cc1ccccc1)NC(=O)C(N)CC(O)=O', 'sweetness'),
('CCCOC(=O)CNC(=O)C(N)CC(O)=O', 'sweetness'),
('COC(=O)C(NC(=O)C(N)CC(O)=O)C(C)O', 'sweetness'),
('CCCC(O)C(NC(=O)C(N)CC(O)=O)C(=O)OC', 'sweetness'),
```

('COC(=O)C(CO)NC(=O)C(N)CC(O)=O', 'sweetness'),
('CCCOC(=O)C(NC(=O)C(N)CC(O)=O)C(C)O', 'sweetness'),
('CCCOC(=O)C(CO)NC(=O)C(N)CC(O)=O', 'sweetness'),
('CCCCCC(C)C(C)NC(=O)C(N)CC(O)=O', 'sweetness'),
('CCOC(=O)CC(NC(=O)C(N)CC(O)=O)C(=O)OC', 'sweetness'),
('CCOC(=O)CC(C)NC(=O)C(N)CC(O)=O', 'sweetness'), ('CCOCCC(C)NC(=O)C(N)CC(O)=O', 'sweetness'), ('COC(=O)CC(C)NC(=O)C(N)CC(O)=O', 'sweetness'),
('COC(=O)C(CCCNC(C)=O)NC(=O)C(N)CC(O)=O', 'sweetness'),
('CC(C)CCC(C)NC(=O)C(N)CC(O)=O', 'sweetness'),
('COC(=O)C(CCCCNC(C)=O)NC(=O)C(N)CC(O)=O', 'sweetness'),
('CCCCCC(C)NC(=O)C(N)CC(O)=O', 'sweetness'),
('CC(NC(=O)C(N)CC(O)=O)NC(=O)C1CC2CCC1C2', 'sweetness'),
('CC(NC(=O)C(N)CC(O)=O)NC(=O)C(C1CC1)C1CC1', 'sweetness'),
('CC(NC(=O)C(N)CC(O)=O)NC(=O)C(C)(C)C', 'sweetness'),
('CC(COC(=O)C1CCC1C)NC(=O)C(N)CC(O)=O', 'sweetness'),
('CC(COC(=O)C1CC1C)NC(=O)C(N)CC(O)=O', 'sweetness'),
('CC(Cc1ccc(cc1)O)NC(=O)C(N)CC(O)=O', 'sweetness'),
('CCCC(=O)OCC(CO)NC(=O)C(N)CC(O)=O', 'sweetness'),
('CCCC(=O)OCC(NC(=O)C(N)CC(O)=O)C(C)C', 'sweetness'),
('CCCC(=O)OCC(C)NC(=O)C(N)CC(O)=O', 'sweetness'),
('NC(CC(O)=O)C(=O)NC(CO)COC(=O)C1CCC1', 'sweetness'),
('CC(C)C(COC(=O)C1CCC1)NC(=O)C(N)CC(O)=O', 'sweetness'),
('CCC(COC(=O)C1CCC1)NC(=O)C(N)CC(O)=O', 'sweetness'),
('CC(COC(=O)C1CCC1)NC(=O)C(N)CC(O)=O', 'sweetness'),
('CC(NC(=O)C(N)CC(O)=O)C(=O)NC1CCCCC1', 'sweetness'),
('CC(NC(=O)C(N)CC(O)=O)NC(=O)C1CCCCC1', 'sweetness'),
('CCOC(=O)C(NC(=O)C(N)CC(O)=O)C(=O)OC1CCCCC1', 'sweetness'),
('NC(CC(O)=O)C(=O)NC(CO)C(=O)OC1CCCCC1', 'sweetness'),
('COC(=O)C(NC(=O)C(N)CC(O)=O)C(=O)OC1CCCCC1', 'sweetness'),
('NC(CC(O)=O)C(=O)NC(CO)COC(=O)C1CCCC1', 'sweetness'),
('CC(COC(=O)C1CCCC1)NC(=O)C(N)CC(O)=O', 'sweetness'),
('CC(NC(=O)C(N)CC(O)=O)NC(=O)C1CCCC1', 'sweetness'),
('CCOC(=O)C(NC(=O)C(N)CC(O)=O)C(=O)OC1CCCC1', 'sweetness'),
('COC(=O)C(NC(=O)C(N)CC(O)=O)C(=O)OC1CCCC1', 'sweetness'),
('NC(CC(O)=O)C(=O)NC(CO)COC(=O)C1CC1', 'sweetness'),
('CC(C)C(COC(=O)C1CC1)NC(=O)C(N)CC(O)=O', 'sweetness'),
('CCC(COC(=O)C1CC1)NC(=O)C(N)CC(O)=O', 'sweetness'),
('CC(COC(=O)C1CC1)NC(=O)C(N)CC(O)=O', 'sweetness'),
('CCCCC(=O)OCC(C)NC(=O)C(N)CC(O)=O', 'sweetness'),
('CC(NC(=O)C(N)CC(O)=O)NC(=O)c1ccccc1', 'sweetness'),
('CC(COC(=O)C=C)NC(=O)C(N)CC(O)=O', 'sweetness'),
('CCC(COC(=O)CC)NC(=O)C(N)CC(O)=O', 'sweetness'),
('CCC(=O)OCC(C)NC(=O)C(N)CC(O)=O', 'sweetness'),
('CCOC(=O)C(NC(=O)C(N)CC(O)=O)C(=O)OC(C)(C)C', 'sweetness'),
('COC(=O)C(NC(=O)C(N)CC(O)=O)C(=O)OC(C)(C)C', 'sweetness'),
('CCOC(=O)C(NC(=O)C(N)CC(O)=O)C(=O)OC1C(C)(C)C2CCC1(C)C2', 'sweetness'),
('NC(CC(O)=O)C(=O)NC(CO)Cc1ccc(cc1)O', 'sweetness'),

('CCC(=O)OCC(CO)NC(=O)C(N)CC(O)=O', 'sweetness'),
('COC(=O)C(NC(=O)C(N)CC(O)=O)C(=O)OC1CC2CCC1(C)C2(C)C', 'sweetness'),
('COC(=O)C(CSC(C)C)NC(=O)C(N)CC(O)=O', 'sweetness'),
('CCC(=O)OCC(NC(=O)C(N)CC(O)=O)C(=O)OC', 'sweetness'),
('CCCSCC(NC(=O)C(N)CC(O)=O)C(=O)OC', 'sweetness'),
('CCCCCOC(=O)C(C)NC(=O)C(N)CC(O)=O', 'sweetness'),
('CCC(NC(=O)C(N)CC(O)=O)C(=O)OC(C)C', 'sweetness'),
('CCCC(NC(=O)C(N)CC(O)=O)C(=O)OC(C)C', 'sweetness'),
('CC(C)OC(=O)C(C)NC(=O)C(N)CC(O)=O', 'sweetness'),
('CCCC(=O)OCCNC(=O)C(N)CC(O)=O', 'sweetness'),
('NC(CC(O)=O)C(=O)NCCOC(=O)C1CCC1', 'sweetness'),
('NC(CC(O)=O)C(=O)NCC(=O)OC1CCCCC1', 'sweetness'),
('NC(CC(O)=O)C(=O)NCCOC(=O)C1CC1', 'sweetness'), ('CCC(=O)OCCNC(=O)C(N)CC(O)=O', 'sweetness'), ('CC(C)OC(=O)C(C)CNC(=O)C(N)CC(O)=O', 'sweetness'),
('CCC(C)OC(=O)CCNC(=O)C(N)CC(O)=O', 'sweetness'),
('CCCC(=O)OCC(NC(=O)C(N)CC(O)=O)C(=O)OC', 'sweetness'),
('CC(CNC(=O)C(N)CC(O)=O)C(=O)OC1CCCCC1', 'sweetness'),
('NC(CC(O)=O)C(=O)NCCC(=O)OC1CCCCC1', 'sweetness'),
('CC(CNC(=O)C(N)CC(O)=O)C(=O)OC1CCCC1', 'sweetness'),
('NC(CC(O)=O)C(=O)NCCC(=O)OC1CCCC1', 'sweetness'),
('CCSCC(NC(=O)C(N)CC(O)=O)C(=O)OC', 'sweetness'),
('NC(CC(O)=O)C(=O)NCCCC1=CC=CO1', 'sweetness'),
('COC(=O)C(COC(C)(C)C)NC(=O)C(N)CC(O)=O', 'sweetness'),
('CC(CNC(=O)C(N)CC(O)=O)C(=O)OC(C)(C)C', 'sweetness'),
('CC(C)(C)OC(=O)CCNC(=O)C(N)CC(O)=O', 'sweetness'),
('COC(=O)C(CSC(C)(C)C)NC(=O)C(N)CC(O)=O', 'sweetness'),
('CC(C)COC(=O)C(NC(=O)C(N)CC(O)=O)C(C)O', 'sweetness'),
('CC(C)COC(=O)C(CO)NC(=O)C(N)CC(O)=O', 'sweetness'),
('CCC(=O)OCC(NC(=O)C(N)CC(O)=O)C(C)O', 'sweetness'),
('CC(C)OC(=O)C(NC(=O)C(N)CC(O)=O)C(C)O', 'sweetness'),
('CC(C)OC(=O)C(CO)NC(=O)C(N)CC(O)=O', 'sweetness'),
('CCC(=O)OCC(NC(=O)C(N)CC(O)=O)C(C)C', 'sweetness'),
('CC(C)OC(=O)C(NC(=O)C(N)CC(O)=O)C(C)C', 'sweetness'),
('CCC(C)C(NC(=O)C(N)CC(O)=O)C(=O)OC(C)C', 'sweetness'),
('CCCC(C)OC(=O)CCNC(=O)C(N)CC(O)=O', 'sweetness'),
('CCC(CC)OC(=O)CCNC(=O)C(N)CC(O)=O', 'sweetness'),
('CC(C)OC(=O)CCNC(=O)C(N)CC(O)=O', 'sweetness'),
('CCC(C)OC(=O)CC(C)NC(=O)C(N)CC(O)=O', 'sweetness'),
('CC(CC(=O)OC1CCCCC1)NC(=O)C(N)CC(O)=O', 'sweetness'),
('COC(=O)C(CC(=O)OC(C)(C)C)NC(=O)C(N)CC(O)=O', 'sweetness'),
('CC(CC(=O)OC(C)(C)C)NC(=O)C(N)CC(O)=O', 'sweetness'),
('CC(C)OC(=O)CC(C)NC(=O)C(N)CC(O)=O', 'sweetness'),
('NC(CC(O)=O)C(=O)NCCC1CCCCC1', 'sweetness'), ('CCCCCCCNC(=O)C(N)CC(O)=O', 'sweetness'), ('CCCCCCNC(=O)C(N)CC(O)=O', 'sweetness'),
('CCOC(=O)C(C)NC(=O)C(N)CC(O)=O', 'sweetness'),
('NC(CC(O)=O)C(=O)NC(CO)Cc1ccccc1', 'sweetness'), ('COC(=O)CNC(=O)C(N)CC(O)=O', 'sweetness'), ('COC(=O)C(CC1CCCCC1)NC(=O)C(N)CC(O)=O', 'sweetness'),

('COC(=O)CCNC(=O)C(N)CC(O)=O', 'sweetness'), ('COC(=O)CCCNC(=O)C(N)CC(O)=O', 'sweetness'), ('CC(NC(=O)C(N)CC(O)=O)C(=O)NC1CCCC1', 'sweetness'), ('COC(=O)C(Cc1ccc(cc1)O)NC(=O)C(N)CC(O)=O', 'sweetness'), ('CC(Cc1ccc(cc1)F)NC(=O)C(N)CC(O)=O', 'sweetness'), ('NC(CC(O)=O)C(=O)NCCc1ccc(cc1)F', 'sweetness'), ('NC(CC(O)=O)C(=O)NCCc1ccc(cc1)O', 'sweetness'), ('CC(CCC1=CC=CO1)NC(=O)C(N)CC(O)=O', 'sweetness'), ('Nc1cc(ccc1Br)C#N', 'sweetness'), ('Nc1cc(ccc1Cl)C#N', 'sweetness'), ('CCOc1ccc(cc1N)C#N', 'sweetness'), ('COc1ccc(cc1N)C#N', 'sweetness'), ('CC(Cc1ccccc1)NC(=O)C(N)CC(O)=O', 'sweetness'), ('NC(CC(O)=O)C(=O)NCCOc1ccccc1', 'sweetness'), ('CC(CCc1ccccc1)NC(=O)C(N)CC(O)=O', 'sweetness'), ('CCCOc1ccc(cc1N)C#N', 'sweetness'), ('Nc1cccc(c1)C(O)=O', 'sweetness'), ('Nc1cccc(c1)C#N', 'sweetness'),
('CC(=C(C)C1=Nc2c(cc(c3ccccc23)S([O-])(=O)=O)N1Nc1ccccc1)c1ccccc1', 'sweetness'), ('NC(=O)c1c(cc(c(c1Br)C=CC(O)=O)Br)Br', 'sweetness'), ('COc1ccc(cc1O)COc1ccccc1', 'sweetness'), ('COc1ccc(cc1O)COC(=O)c1ccccc1O', 'sweetness'), ('OCC1OC(OC2(CO)OC(CO)C(O)C2O)C(O)C(=O)C1O', 'sweetness'), ('COc1cc2c(cc1O)C1CCC3(C)CCCC3C1CC2', 'sweetness'), ('CC1(C)C(CCC2(C)C1CCC1(C)C2 C(=O)C=C2C3CC(C)(CCC3(C)CCC12C)C(O)=O)OC1OCC(O)C(O)C1O', 'sweetness'), ('OC(=O)CNC(NCc1ccccc1)=Nc1cccnc1', 'sweetness'), ('CC(NC(NCC(O)=O)=Nc1cnc2ccccc2c1)c1ccccc1', 'sweetness'), ('OCC1OC(OC2(CO)OC(CO)C(Cl)C2O)C(O)C(O)C1O', 'sweetness'), ('COC(C)C1=CC=C(C)CC1', 'sweetness'), ('COC(C)C1=CCC(=CC1)C', 'sweetness'), ('COC(C)C1CCC(=CC1)C', 'sweetness'), ('[Na+].CCN(CC)c1ccc(cc1)NS([O-])(=O)=O', 'sweetness'), ('OCC1OC(OC2(CBr)OC(CBr)C(Br)C2O)C(O)C(O)C1Br', 'sweetness'), ('OCC1OC(OC2(CCl)OC(CCl)C(Cl)C2O)C(O)C(O)C1Cl', 'sweetness'), ('OCC1OC(OC2(CCl)OC(CO)C(Cl)C2O)C(O)C(O)C1Cl', 'sweetness'), ('OCC1OC(OC2(CBr)OC(CBr)C(O)C2O)C(O)C(O)C1Br', 'sweetness'), ('OCC1OC(OC2(CCl)OC(CCl)C(Br)C2O)C(O)C(O)C1Cl', 'sweetness'), ('OCC1OC(OC2(CCl)OC(CCl)C(I)C2O)C(O)C(O)C1Cl', 'sweetness'), ('OCC1OC(OC2(CF)OC(CF)C(O)C2O)C(O)C(O)C1F', 'sweetness'), ('OCC1OC(OC2(CI)OC(CI)C(O)C2O)C(O)C(O)C1I', 'sweetness'), ('OCC1OC(CCl)(OC2OC(CO)C(Cl)C(O)C2O)C(O)C1O', 'sweetness'), ('OCC1OC(OC2(CO)OC(CCl)C(Cl)C2O)C(O)C(O)C1Cl', 'sweetness'), ('OC1C(O)C(OC(CCl)C1Cl)OC1(CCl)OC(CCl)C(O)C1O', 'sweetness'), ('OCC1OC(OC2(CO)OC(CCl)C(O)C2O)C(O)C(O)C1Cl', 'sweetness'), ('[Na+].Cc1cc(nc(c1)NS([O-])(=O)=O)C', 'sweetness'), ('COCC1OC(CO)(OC2OC(CO)C(OC)C(O)C2O)C(O)C1O', 'sweetness'), ('COCC1OC(OC2(CO)OC(CO)C(O)C2O)C(O)C(O)C1OC', 'sweetness'), ('OCC1OC(OC2(CCl)OC(CCl)C(O)C2O)C(O)C(O)C1Cl', 'sweetness'), ('OCC1OC(CO)(OC2OC(CO)C(Cl)C(O)C2O)C(O)C1O', 'sweetness'), ('OCC1OC(OC2(CBr)OC(CBr)C(Br)C2O)C(O)C(O)C1Cl', 'sweetness'), ('Clc1cccc2c1C(=O)NS2(=O)=O', 'sweetness'), ('OC(=O)CNC(NC1CCCCCC1)=Nc1ccc(cc1)C#N', 'sweetness'), ('OC(=O)CNC(NC1CCCCCCCC1)=Nc1ccc(cc1)C#N', 'sweetness'), ('OC(=O)CNC(NC(c1ccccc1)c1ccccc1)=Nc1ccc(cc1)C#N', 'sweetness'),

('OC(=O)CNC(NC1CCCCCC1)=Nc1ccc(cc1)C#N', 'sweetness'),
('OC(=O)CNC(Nc1cccc2ccccc12)=Nc1ccc(cc1)C#N', 'sweetness'),
('OC(=O)CNC(NC(C1=CC=CO1)c1ccccc1)=Nc1ccc(cc1)C#N', 'sweetness'),
('CC(NC(NCC(O)=O)=Nc1ccc(cc1)C#N)C1CCCCC1', 'sweetness'),
('OC(=O)CNC(NCC1CCCCC1)=Nc1ccc(cc1)C#N', 'sweetness'),
('OC(=O)CNC(NCc1ccccc1)=Nc1ccc(cc1)C#N', 'sweetness'),
('CC(NC(NCC(O)=O)=Nc1ccc(cc1)C#N)c1ccccc1', 'sweetness'),
('COCC(NC(NCC(O)=O)=Nc1ccc(cc1)C#N)c1ccccc1', 'sweetness'),
('OC(=O)CNC(NCC12CC3CC(CC(C3)C1)C2)=Nc1ccc(cc1)C#N', 'sweetness'),
('CN(NC(NCC(O)=O)=Nc1ccc(cc1)C#N)c1ccccc1', 'sweetness'),
('OC(=O)CNC(NC1CCCCC1)=Nc1ccc(cc1)C#N', 'sweetness'),
('OC(=O)CNC(Nc1cccc(c1)Cl)=Nc1ccc(cc1)C#N', 'sweetness'),
('Cc1cccc(c1)NC(NCC(O)=O)=Nc1ccc(cc1)C#N', 'sweetness'),
('OC(=O)CNC(NCCc1ccccc1)=Nc1ccc(cc1)C#N', 'sweetness'),
('Cc1ccc(cc1)NC(NCC(O)=O)=Nc1ccc(cc1)C#N', 'sweetness'),
('CCCCC(C)NC(NCC(O)=O)=Nc1ccc(cc1)C#N', 'sweetness'),
('CCCCCCNC(NCC(O)=O)=Nc1ccc(cc1)C#N', 'sweetness'),
('Cc1ccccc1NC(NCC(O)=O)=Nc1ccc(cc1)C#N', 'sweetness'),
('OCc1ccc(cc1CO)NC(NCC(O)=O)=Nc1ccc(cc1)C#N', 'sweetness'),
('OC(=O)CNC(NC1CCc2ccccc12)=Nc1ccc(cc1)C#N', 'sweetness'),
('OC(=O)CNC(Nc1ccccc1)=Nc1ccc(cc1)C#N', 'sweetness'),
('OC(=O)CNC(NC12CC3CC(CC(C3)C1)C2)=Nc1ccc(cc1)C#N', 'sweetness'),
('NC(NCC(O)=O)=Nc1ccc(cc1)C#N', 'sweetness'), ('CCNC(NCC(O)=O)=Nc1ccc(cc1)C#N',
'sweetness'), ('OC(=O)CNC(NS(=O)(=O)c1ccccc1)=Nc1ccc(cc1)C#N', 'sweetness'),
('OC(=O)CNC(NS(=O)(=O)Cc1ccccc1)=Nc1ccc(cc1)C#N', 'sweetness'),
('OC(=O)CNC(NS(=O)(=O)C1CCCCCC1)=Nc1ccc(cc1)C#N', 'sweetness'),
('CCCCS(=O)(=O)NC(NCC(O)=O)=Nc1ccc(cc1)C#N', 'sweetness'),
('CCCCCS(=O)(=O)NC(NCC(O)=O)=Nc1ccc(cc1)C#N', 'sweetness'),
('OC(=O)CNC(NS(=O)(=O)C1CCCCC1)=Nc1ccc(cc1)C#N', 'sweetness'),
('CS(=O)(=O)NC(NCC(O)=O)=Nc1ccc(cc1)C#N', 'sweetness'),
('OCC1OC(OC2(CBr)OC(CBr)C(O)C2O)C(O)C(O)C1Cl', 'sweetness'),
('CC1(OC(CCl)C(F)C1O)OC1OC(CO)C(Cl)C(O)C1O', 'sweetness'),
('OCC1OC(OC2(CO)OC(CF)C(O)C2O)C(O)C(O)C1F', 'sweetness'),
('OCC1OC(OC2(CCl)OC(CCl)C(O)C2O)C(O)C(O)C1Br', 'sweetness'),
('OCC1CC(O)C(O)C(O1)OC1(CO)OC(CO)C(O)C1O', 'sweetness'), ('CC1=CCC(CC1)C=C',
'sweetness'), ('OCC1OC(OC2(CBr)OC(CBr)C(Br)C2O)C(O)C(O)C1F', 'sweetness'),
('OCC1OC(OC2(CCl)OC(CCl)C(Cl)C2O)C(O)C(O)C1F', 'sweetness'),
('OCC1OC(OC2(CCl)OC(CCl)C(F)C2O)C(O)C(O)C1Cl', 'sweetness'),
('OCC1OC(OC2C(O)OC(CO)C(OC3OC(CO)C(O)C(O)C3O)C2O)C(O)C(O)C1O', 'sweetness'),
('COCC1=CCC(=CC1)C', 'sweetness'), ('COCC1CCC(=CC1)C', 'sweetness'),
('COc1ccc(cc1)NC(N)=O', 'sweetness'), ('[Na+].CC1CCC(CC1)NS([O-])(=O)=O',
'sweetness'), ('NC(NCC(O)=O)=Nc1ccc(cc1)N(=O)=O', 'sweetness'),
('COC1C(O)C(O)C(OC1CO)OC1(CO)OC(CO)C(O)C1O', 'sweetness'),
('CCCOc1ccc(cc1O)CCC(=O)c1c(cc(cc1O)OC1OC(CO)C(O)C(O)C1OC1OC(C)C(O)C(O)C1O)O',
'sweetness'), ('CCCOc1ccc(cc1)NC(N)=O', 'sweetness'),
('CC1(CCCC2(C)C1C(C(O)=O)c1ccccc21)C(O)=O', 'sweetness'),
('COc1ccc(cc1OC(=O)Nc1ccccc1)C1OCc2ccccc2S1', 'sweetness'),

('COc1ccc(cc1O)CCC1CCCCC1', 'sweetness'), ('COc1ccc(cc1O)C1CCc2ccccc2O1', 'sweetness'), ('OC(=O)CCC(NC(=O)C(F)(F)F)C(=O)Nc1ccc(cc1)C#N', 'sweetness'), ('COc1ccc(cc1O)C1OCc2ccccc2S1', 'tastelessness'), ('COc1ccc(cc1O)NCc1ccccc1', 'sweetness'), ('CC1=C(C)C(=O)NS(=O)(=O)O1', 'sweetness'), ('COc1ccc(cc1O)CCc1ccccc1O', 'sweetness'), ('COc1ccc(cc1O)CCc1ccc(cc1)F', 'sweetness'), ('COc1ccc(cc1O)CCc1ccc(cc1)O', 'sweetness'), ('COc1ccc(cc1OC(C)=O)C1Cc2cccc(c2C(=O)O1)OC(C)=O', 'sweetness'), ('COc1ccc(cc1O)C(O)Oc1ccccc1', 'sweetness'), ('COc1ccc(cc1O)CCc1ccccc1CO', 'sweetness'), ('Brc1cnc(nc1)C#N', 'sweetness'), ('OCC1(O)OCCC(O)C1O', 'sweetness'), ('CCC1=C(C)OS(=O)(=O)NC1=O', 'sweetness'), ('CC(NC(NCC(O)=O)=Nc1ccc2c(c1)CCC2)c1ccccc1', 'sweetness'), ('[Na+].CC1=NN=C(NS([O-])(=O)=O)S1', 'sweetness'), ('CC1=COS(=O)(=O)NC1=O', 'sweetness'), ('CCCc1ccc(cc1N)N(=O)=O', 'sweetness'), ('OCC1SC(O)C(O)C(O)C1O', 'sweetness'), ('OC1C(O)C(CCl)OC(OC2(CCl)OC(CCl)C(O)C2O)C1O', 'sweetness'), ('COCC1OC(OC2(CO)OC(COC)C(O)C2O)C(O)C(O)C1O', 'sweetness'), ('OCC1OC(OC2(CO)OC(CCl)C(O)C2O)C(O)C(O)C1O', 'sweetness'), ('OCC1(O)OC(CCl)C(O)C1O', 'sweetness'), ('NC(CC1=CNc2cc(ccc12)Cl)C(O)=O', 'miscellaneous'), ('CC1OC(OC2(CO)OC(CO)C(O)C2O)C(O)C(O)C1O', 'sweetness'), ('CCC1=C(C)C(=O)NS(=O)(=O)O1', 'sweetness'), ('CCC1=CC(=O)NS(=O)(=O)O1', 'sweetness'), ('Fc1ccc2c(c1)S(=O)(=O)NC2=O', 'sweetness'), ('Oc1ccc2c(c1)S(=O)(=O)NC2=O', 'sweetness'), ('Cc1ccc2c(c1)S(=O)(=O)NC2=O', 'sweetness'), ('CC(=O)C1OC(OC2(CO)OC(CO)C(O)C2O)C(O)C(O)C1O', 'sweetness'), ('COCC1OC(CO)(OC2OC(CO)C(O)C(O)C2O)C(O)C1O', 'sweetness'), ('COCC1OC(OC2(CO)OC(CO)C(O)C2O)C(O)C(O)C1O', 'sweetness'), ('OC(=O)CNC(NC1CCCCCCC1)=Nc1ccc2ccccc2n1', 'sweetness'), ('CC(NC(NCC(O)=O)=Nc1ccc2ccccc2n1)c1ccccc1', 'sweetness'), ('OCC1(O)SCC(O)C(O)C1O', 'sweetness'), ('NC(CC1=CNc2cc(ccc12)C(F)(F)F)C(O)=O', 'sweetness'), ('Nc1cccc2c1S(=O)(=O)NC2=O', 'sweetness'), ('COc1ccc(cc1O)C1Cc2ccccc2C(=O)O1', 'sweetness'), ('COc1ccc(cc1O)C1Cc2cccc(c2CO1)O', 'sweetness'), ('COc1ccc(cc1O)C1Cc2cccc(c2C(=O)N1)O', 'sweetness'), ('COc1cc2c(cc1O)C1c3ccccc3OCC1(O)C2', 'sweetness'), ('CC(C1CC=C(C)C(=O)O1)C1CCC2(C)C3CCC4C(C)(C(CCC54CC35CCC12C)OC1OC(CO)C(O)C(O)C1O)C(O)=O', 'sweetness'), ('COC(=O)C1OC(OC2CCC34CC54CCC4(C)C(CCC4(C)C5CCC3C2(C)C(O)=O)C(C)C2CC=C(C)C(=O)O2)C(OC2OC(CO)C(O)C(O)C2O)C(O)C1O', 'sweetness'), ('CC(C1CC=C(C)C(=O)O1)C1CCC2(C)C3CCC4C(C)(C(CCC54CC35CCC12C)OC1OC(CO)C(O)C(O)C1OC1OC(CO)C(O)C(O)C1O)C(O)=O', 'sweetness'), ('CC(C1CC=C(C)C(=O)O1)C1CCC2(C)C3CCC4C(C)(C(CCC54CC35CCC12C)OC1OC(C(O)C(O)C1OC1OC(CO)C(O)C(O)C1O)C(O)=O)C(O)=O', 'sweetness'), ('CC(C1CC=C(C)C(=O)O1)C1CCC2(C)C3CCC4C(C)(C(CCC54CC35CCC12C)OC1OC(CO)C(O)C(O)C1OC1OC(C(O)C(O)C1O)C(O)=O)C(O)=O', 'sweetness'), ('COC(=O)C1OC(OC2C(O)C(O)C(CO)OC2OC2CCC34CC54CCC4(C)C(CCC4(C)C5CCC3C2(C)C(O)=O)C(C)C2CC=C(C)C(=O)O2)C(O)C(O)C1O', 'sweetness'), ('CC1=CC(=O)NS(=O)(=O)O1', 'sweetness'), ('[K+].CC1=CC(=O)[N-]S(=O)(=O)O1', 'sweetness'), ('CCC1=COS(=O)(=O)NC1=O', 'sweetness'), ('CCCC1=C(C)OS(=O)(=O)NC1=O', 'sweetness'), ('O=C1NS(=O)(=O)OC=C1', 'sweetness'), ('CC=NO', 'sweetness'), ('COC(=O)C(Cc1ccccc1)NC(=O)C(CC(O)=O)NCCCc1ccc(c(c1)O)OC', 'sweetness'), ('CC1(C)CC2C3=CCC4C5(C)CCC(OC6OC(COC7OCC(O)C(O)C7O)C(O)C(O)C6OC6OC(CO)C(O)C(O)C6O)C(C)(

```
C)C5CCC4(C)C3(C)CC(O)C32CC1OC3=O', 'sweetness'),
('CC(NC(=O)C(N)CC(O)=O)C(=O)NC1C(C)(C)SC1(C)C', 'sweetness'),
('Nc1cc(ccc1OC=C)N(=O)=O', 'sweetness'), ('[NH4+].CC1(C)C(CCC2(C)C1CCC1(C)C2C(=O
)C=C2C3CC(C)(CCC3(C)CCC12C)C(O)=O)OC1OC(C(O)C(O)C1OC1OC(C(O)C(O)C1O)C(O)=O)C([O-
])=O', 'sweetness'), ('ON=CC1=CC=C(Cc2ccccc2)O1', 'sweetness'),
('ON=CC1=CC=CO1', 'sweetness'), ('OCC(O)C(O)C(O)C=O', 'sweetness'),
('CC1(C)CC1(C(O)=O)S(=O)(=O)c1ccccc1', 'sweetness'),
('CC1CCC(CC1C)(C(O)=O)S(=O)(=O)c1ccccc1', 'sweetness'),
('CC1CC=CCC1(C(O)=O)S(=O)(=O)c1ccccc1', 'sweetness'),
('COC(=O)C(Cc1ccccc1)NC(=O)C(N)CC(O)=O', 'bitterness'),
('COC(=O)C(Cc1ccccc1)NC(=O)C([NH3+])CC(O)=O.CC1=CC(=O)[N-]S(=O)(=O)O1',
'sweetness'), ('COC(=O)C(NC(=O)C(N)C(O)=O)C(=O)OC1C(C)(C)C2CCC1(C)C2',
'sweetness'), ('OC(=O)CCNC(=O)Nc1ccc(cc1)N(=O)=O', 'sweetness'),
('CCCCOC(=O)C(Cc1ccccc1)NC(=O)C(N)CC(O)=O', 'sweetness'),
('CCOC(=O)C(Cc1ccccc1)NC(=O)C(N)CC(O)=O', 'sweetness'),
('CCCOC(=O)C(Cc1ccccc1)NC(=O)C(N)CC(O)=O', 'sweetness'),
('CC1=C(CCC2=COC=C2)C2(C)CCC(OC3OC(CO)C(O)C(O)C3OC3OCC(O)C(O)C3O)C(C)(C)C2CC1',
'sweetness'), ('OC(=O)CNC(NC1CCCCCCC1)=NC1=CC2=NON=C2C=C1', 'sweetness'),
('Cc1cccc(c1)CNC(NCC(O)=O)=Nc1ccc(cc1)C#N', 'sweetness'), ('[Be+2].[Cl-].[Cl-]',
'sweetness'), ('Nc1cc(ccc1Br)N(=O)=O', 'sweetness'),
('CCC(C)CC(C)NC(=O)C(N)CC(O)=O', 'sweetness'), ('CCCCC(CC)NC(=O)C(N)CC(O)=O',
'sweetness'), ('CCCCCCC(C)NC(=O)C(N)CC(O)=O', 'sweetness'),
('CCCCC(C)NC(=O)C(N)CC(O)=O', 'sweetness'), ('CCCCOS(N)(=O)=O', 'sweetness'),
('[Ca+2].[O-]S(=O)(=O)NC1CCCCC1.[O-]S(=O)(=O)NC1CCCCC1', 'sweetness'),
('[Ca+2].O=C1[N-]S(=O)(=O)c2ccccc12.O=C1[N-]S(=O)(=O)c2ccccc12', 'sweetness'),
('OC(=O)CN=C(NC(c1ccccc1)c1ccccc1)Nc1cc(cc(c1)Cl)Cl', 'sweetness'),
('OCC1OC(OC2C(O)C(O)C(O)OC2CO)C(O)C(O)C1O', 'sweetness'), ('ClC(Cl)Cl',
'sweetness'), ('Nc1cc(ccc1Cl)N(=O)=O', 'sweetness'),
('COC(=O)C(CC(=O)CNC(=O)Nc1ccc(cc1)C#N)Cc1ccccc1', 'sweetness'),
('OC(=O)CCNC(=O)Nc1ccc(cc1)C#N', 'sweetness'), ('OS(=O)(=O)NC1CCCCC1',
'sweetness'), ('CC1CC2C(C)(C)C(CCC2(C)C2CC(OC3OC(C)C(O)C(O)C3O)C3C(CCC3(C)C12)C1
(C)CCC(O1)C(C)(C)O)OC1OC(COC(C)=O)C(O)C1O', 'sweetness'),
('[Na+].[O-]S(=O)(=O)NC1CCCCCC1', 'sweetness'),
('[Na+].[O-]S(=O)(=O)NC1CCCCCCC1', 'sweetness'), ('[Na+].[O-]S(=O)(=O)NC1CCCC1',
'sweetness'), ('COC(=O)C(Cc1ccccc1)NC(=O)C(CC(O)=O)NC(=O)C(C)N', 'non-
sweetness'), ('CC(N)C(O)=O', 'sweetness'), ('OCC(O)C(O)C(O)CO', 'sweetness'),
('NC(CC(N)=O)C(O)=O', 'miscellaneous'), ('[Na+].[O-]S(=O)(=O)NN1CCC2CCCCC2C1',
'sweetness'), ('OCC1OC(OC2(CBr)OC(CBr)C(O)C2O)C(O)C(O)C1O', 'sweetness'),
('OCC1OC(O)C(O)C(O)C1O', 'bitterness'), ('NC(CCC(N)=O)C(O)=O', 'miscellaneous'),
('NC(CC1=CN=CN1)C(O)=O', 'bitterness'),
('COC(=O)C(Cc1ccccc1)NC(=O)C(CC(O)=O)NC(=S)Nc1cnc(nc1)C#N', 'sweetness'),
('COc1ccc(cc1O)C1Oc2cc(cc(c2C(=O)C1OC(C)=O)O)O', 'sweetness'), ('OCC(=O)CO',
'sweetness'), ('CN(C)C(N)=O', 'sweetness'), ('[Na+].[Na+].CC1(C)C(CCC2(C)C1CCC1(
C)C2C(=O)C=C2C3CC(C)(CCC3(C)CCC12C)C(O)=O)OC1OC(C(O)C(O)C1OC1OC(C(O)C(O)C1O)C([O
-])=O)C([O-])=O', 'sweetness'), ('CCC(C)C(N)C(O)=O', 'bitterness'),
('OCC1OC(OC2C(CO)OC(O)(CO)C2O)C(O)C(O)C1O', 'sweetness'), ('CC(C)CC(N)C(O)=O',
'bitterness'), ('OCC1OC(OC2COC(O)(CO)C(O)C2O)C(O)C(O)C1O', 'sweetness'),
```

('NCCCCC(N)C(O)=O', 'bitterness'),
('OCC1OC(OC2C(O)C(O)C(OC2CO)OC2C(O)C(O)C(O)OC2CO)C(O)C(O)C1O', 'sweetness'),
('OCC(O)C(O)C(O)C(O)CO', 'sweetness'), ('CSCCC(N)C(O)=O', 'multitaste'),
('CCCC(N)C(O)=O', 'bitterness'), ('NC(Cc1ccccc1)C(O)=O', 'bitterness'),
('CC1OC(O)C(O)C(O)C1O', 'sweetness'), ('NC(CO)C(O)=O', 'sweetness'),
('OCC1(O)OCC(O)C(O)C1O', 'bitterness'), ('CC(O)C(N)C(O)=O', 'sweetness'),
('NC(CC1=CNc2ccccc12)C(O)=O', 'bitterness'), ('NC(Cc1ccc(cc1)O)C(O)=O',
'bitterness'), ('CCOc1ccc(cc1)NC(N)=O', 'sweetness'), ('CC1OC(OC2C(O)C(O)C(CO)OC
2OC23CCC4C5(C)CCCC(C)(C5CCC4(CC2=C)C3)C(=O)OC2OC(CO)C(O)C(O)C2O)C(O)C(O)C1O',
'sweetness'), ('CC(C)C(N)C(O)=O', 'bitterness'), ('OC1COC(O)C(O)C1O',
'sweetness'), ('CCC(N)C(O)=O', 'sweetness'), ('OCC(O)C(O)CO', 'sweetness'),
('CCOc1ccc(cc1N)N(=O)=O', 'sweetness'), ('OCCO', 'sweetness'),
('[Na+].[O-]S(=O)(=O)NC1CC2CCC1C2', 'sweetness'), ('Nc1cc(ccc1F)N(=O)=O',
'sweetness'), ('O=CC1=CC=CO1', 'sweetness'),
('OCC1OC(OC2(CO)OC(CO)C(O)C2O)C(O)C(O)C1O', 'tastelessness'),
('CC(CCC1=C(CC(OC2OCC(O)C(O)C2O)C2C(C)(CO)CCCC12C)C=O)=CCO', 'sweetness'),
('OCC1OC(CO)(OC2(OC(CO)C(O)C(O)C2O)C2(O)OC(CO)C(O)C(O)C2O)C(O)C1O',
'sweetness'), ('OCC(O)CO', 'sweetness'), ('NCC(O)=O', 'sweetness'),
('OC(=O)CNC(=O)Nc1ccc(cc1)C#N', 'sweetness'), ('CC1(C)C(CCC2(C)C1CCC1(C)C2C(=O)C
=C2C3CC(C)(CCC3(C)CCC12C)C(O)=O)OC1OC(C(O)C(O)C1OC1OC(C(O)C(O)C1O)C(O)=O)C(O)=O'
, 'sweetness'), ('CC1(C)C(CCC2(C)C1CCC1(C)C2C(=O)C=C2C3CC(C)(CCC3(C)CCC12C)C(O)=
O)OC1OC(C(O)C(O)C1OC1CC(C(O)C(O)C1O)C(O)=O)C(O)=O', 'sweetness'), ('NC(N)=N',
'sweetness'), ('N#Cc1ccc(cc1)NC(NCC1=NN=NN1)=NC1CCCCCCC1', 'sweetness'),
('CC(N=C(NCC(O)=O)Nc1cc(cc(c1)Cl)Cl)c1ccccc1', 'sweetness'),
('OC(=O)CNC(Nc1ccc(cc1)C#N)=NC1CCCCCCC1', 'sweetness'),
('CN1N=NN=C1NC(NC1CCCCC1)=Nc1ccc(cc1)C#N', 'sweetness'),
('CN1N=NN=C1NC(NC1CCCCCCC1)=Nc1ccc(cc1)C#N', 'sweetness'),
('CC(NC(NCC(O)=O)Nc1ccc(cc1)C#N)c1ccccc1', 'sweetness'),
('CCCCCCCNC(CC(O)=O)C(=O)Nc1ccc(cc1)C#N', 'sweetness'),
('CC(C)=CCCC(C)(O)C1CCC(=CC1=O)C', 'non-sweetness'),
('COc1ccc(cc1O)CCC(=O)c1c(cc(cc1O)O)O', 'sweetness'),
('COc1ccc(cc1O)CCC(=O)c1c(cc(cc1O)OC1OCC(O)C(O)C1O)O', 'sweetness'),
('COc1ccc(cc1O)CCC(=O)c1c(cc(cc1O)OC1OC(CO)C(O)C(O)C1O)O', 'sweetness'),
('CC1OC(OC2C(O)C(O)C(CO)OC2Oc2cc(c(c(c2)O)C(=O)CCc2cccc(c2)O)O)C(O)C(O)C1O',
'sweetness'), ('OCC1OC(Oc2cc(c(c(c2)O)C(=O)CCc2ccc(c(c2)O)O)O)C(O)C(O)C1O',
'sweetness'), ('[Na+].[O-]S(=O)(=O)NC1CCCCSC1', 'sweetness'),
('[Na+].[O-]S(=O)(=O)NC1CCCSCC1', 'sweetness'),
('CCOc1ccc(cc1O)CCC(=O)c1c(cc(cc1O)OC1OC(CO)C(O)C(O)C1OC1OC(C)C(O)C(O)C1O)O',
'sweetness'), ('OCC(O)C(O)C(OC1OC(CO)C(OC2OC(CO)C(O)C(O)C2O)C(O)C1O)C(O)CO',
'sweetness'), ('Nc1cc(ccc1O)N(=O)=O', 'sweetness'),
('OCC(O)C(O)C(O)C(O)C=O.OCC(O)C(O)C(O)C(=O)CO', 'sweetness'),
('Nc1cc(ccc1I)N(=O)=O', 'sweetness'), ('CC(C)Oc1ccc(cc1N)N(=O)=O', 'sweetness'),
('OCC(O)C(O)C(O)C(O)COC1OC(CO)C(O)C(O)C1O', 'sweetness'), ('CC(CCC(OC1OC(COC2OC(
CO)C(O)C(O)C2O)C(O)C(O)C1OC1OC(CO)C(O)C(O)C1O)C(C)(C)O)C1CCC2(C)C3CC=C4C(CCC(OC5
OC(CO)C(OC6OC(CO)C(O)C(O)C6O)C(O)C5O)C4(C)C)C3(C)C(O)CC12C', 'sweetness'),
('Oc1ccc2c(c1O)OCC1(O)Oc3cc(c(cc3C21)O)O', 'sweetness'),
('COc1ccc2c(c1)OC1(O)COc3ccccc3C21', 'sweetness'),

('OCC1OC(O)C(OC2OC(CO)C(O)C(O)C2O)C(O)C1O', 'sweetness'),
('OCC(O)C(O)C(OC1OC(CO)C(O)C(O)C1O)C(O)CO', 'sweetness'),
('OCC1OC(OC2C(O)C(O)C(OC2CO)OC2(CO)OC(CO)C(O)C2O)C(O)C(O)C1O', 'sweetness'),
('OCC1OC(OC2C(CO)OC(CO)(OC3OC(CO)C(O)C(O)C3O)C2O)C(O)C(O)C1O', 'sweetness'),
('Cl.CC(N)C(=O)OC(C)(C)C', 'sweetness'), ('CC(N)C(=O)OC(C)(C)C', 'bitterness'),
('CCCCOC(=O)C(NC(=O)C(N)CC(O)=O)C(C)O', 'sweetness'),
('CC(O)C(NC(=O)C(N)CC(O)=O)C(=O)OC1CCCCC1', 'sweetness'),
('COC(=O)C(C)NC(=O)C(N)CC(O)=O', 'bitterness'),
('CC(NC(=O)C(N)CC(O)=O)C(=O)NC1C(C)(C)CCC1(C)C', 'bitterness'),
('COC(=O)C(NC(=O)C(N)CC(O)=O)C(=O)OC1C(C)(C)C2CCC1(C)C2', 'sweetness'),
('CCCOC(=O)C(C)NC(=O)C(N)CC(O)=O', 'sweetness'),
('COC(=O)C(CS(C)(C)C)NC(=O)C(N)CC(O)=O', 'sweetness'), ('OCC1OC(O)(CO)C(O)C1O',
'sweetness'), ('OC1CNC(C1)C(O)=O', 'tastelessness'), ('CC(=C)C1CCC(=CC1)C',
'sweetness'), ('OCC1OC(Oc2ccc(cc2)C2CC(=O)c3ccc(cc3O2)O)C(O)C(O)C1O',
'sweetness'), ('OC(=O)C1CCCN1', 'miscellaneous'),
('CC(NC(=O)C(N)CC(O)=O)NC(=O)C1C(C)(C)CCC1(C)C', 'sweetness'),
('OC(=O)CNC(Nc1ccc(cc1)C#N)=NCc1cccc2c1OCO2', 'sweetness'),
('OCC1OC(OC2C(O)COC(O)(CO)C2O)C(O)C(O)C1O', 'sweetness'),
('ClCS(=O)(=O)NCc1ccccc1', 'sweetness'), ('COc1ccc(cc1N)N(=O)=O', 'sweetness'),
('COC1CC(O)C(O)C(CO)O1', 'sweetness'), ('COC1OC(CO)C(O)CC1O', 'sweetness'),
('COC1OC(CO)CC(O)C1O', 'sweetness'), ('COC1OC(C)C(O)C(O)C1O', 'sweetness'),
('CC(=O)C(C)(O)O', 'sweetness'), ('COC1OC(CO)C(O)C(O)C1O', 'multitaste'),
('COC1(CO)OCC(O)C(O)C1O', 'sweetness'), ('Cc1ccc(cc1N)N(=O)=O', 'sweetness'),
('COC1OCC(O)C(O)C1O', 'sweetness'),
('CC(CCC(O)C(C)(C)O)C1CCC2(C)C3CC=C4C(CCC(O)C4(C)C)C3(C)C(O)CC12C',
'sweetness'), ('CC(CCC(OC1OC(CO)C(O)C(O)C1O)C(C)(C)O)C1CCC2(C)C3CC=C4C(CCC(OC5OC
(CO)C(O)C(O)C5O)C4(C)C)C3(C)C(O)CC12C', 'sweetness'), ('CC(CCC(OC1OC(CO)C(O)C(O)
C1OC1OC(CO)C(O)C(O)C1O)C(C)(C)O)C1CCC2(C)C3CC=C4C(CCC(OC5OC(COC6OC(CO)C(O)C(O)C6
O)C(O)C(O)C5O)C4(C)C)C3(C)C(O)CC12C', 'sweetness'), ('CC(CCC(OC1OC(COC2OC(CO)C(O
)C(O)C2O)C(O)C(O)C1OC1OC(CO)C(O)C(O)C1O)C(C)(C)O)C1CCC2(C)C3CC=C4C(CCC(OC5OC(COC
6OC(CO)C(O)C(O)C6O)C(O)C(O)C5O)C4(C)C)C3(C)C(O)CC12C', 'sweetness'),
('NC(CC(O)(CC1=CNc2ccccc12)C(O)=O)C(O)=O', 'sweetness'), ('CC1(C)C(CCC2(C)C1CCC1
(C)C2C(=O)C=C2C3CC(C)(CCC3(C)CCC12C)C(O)=O)OC1OC(C(O)C(O)C1O)C(O)=O',
'sweetness'), ('CC(=O)CO', 'sweetness'), ('CC1OC(OC2C(O)C(CO)OC(OCC=C(C)CCC=C(C)
CCC=C(C)COC3OC(CO)C(O)C(OC4OC(C)C(O)C(O)C4O)C3OC3OC(C)C(O)C(O)C3O)C2OC2OC(C)C(O)
C(O)C2O)C(O)C(O)C1O', 'sweetness'), ('[Na+].[O-]S(=O)(=O)NC1CCSCC1',
'sweetness'), ('[Na+].[O-]S(=O)(=O)NC1CCSC1', 'sweetness'),
('CCC(Cc1ccccc1)NC(=O)C(N)CC(O)=O', 'sweetness'), ('CC=C(C)C=NO', 'sweetness'),
('[Na+].CCC(C)CNS([O-])(=O)=O', 'sweetness'), ('COc1ccc(cc1O)CNc1ccccc1',
'sweetness'), ('ON=CC1=COCCC1', 'sweetness'),
('OC(=O)CC(NC(=O)Nc1ccc(cc1)C#N)c1cccnc1', 'sweetness'),
('OC(=O)CC(NC(=O)Nc1ccc(cc1)C#N)c1ccncc1', 'sweetness'),
('OC(=O)CC(NC(=O)Nc1ccc(cc1)C#N)c1ccccc1', 'sweetness'), ('ON=CC1=CCC(CC1)C=C',
'sweetness'), ('CCOC1CCC(=CC1)C=NO', 'sweetness'), ('COC1CCC(=CC1)C=NO',
'sweetness'), ('CC1=CC=C(CC1)C=NO', 'sweetness'), ('ON=CC1=CC=CCC1',
'sweetness'), ('ON=CC1=CCC=CC1', 'sweetness'), ('COC(C)C1CCC(=CC1)C=NO',
'sweetness'), ('CCC(COC(C)=O)NC(=O)C(N)CC(O)=O', 'sweetness'),

('CC1(CO1)C1CCC(=CC1)C=NO', 'sweetness'), ('COCC1CCC(=CC1)C=NO', 'sweetness'),
('COCC1CC=C(C1)C=NO', 'sweetness'), ('CC(=O)Nc1ccccc1C(=O)CC(N)C(O)=O',
'sweetness'), ('CC(=O)NC1C(O)OC(CO)C(O)C1O', 'sweetness'),
('CC1OC(OC2C(O)C(O)C(CO)OC2Oc2cc(c(c(c2)O)C(=O)CCc2ccc(cc2)O)O)C(O)C(O)C1O',
'sweetness'), ('CCCCOc1ccc(cc1N)N(=O)=O', 'sweetness'),
('[Na+].CCCCNS([O-])(=O)=O', 'sweetness'),
('CC1OC(OC2C(Oc3cc(cc(c3C2=O)O)O)c2ccc(c(c2)O)O)C(O)C(O)C1O', 'non-sweetness'),
('CC1OC(OC2C(O)C(O)C(CO)OC2Oc2cc(c(c(c2)O)C(=O)CCc2ccc(c(c2)O)O)O)C(O)C(O)C1O',
'sweetness'),
('CC(C)Oc1ccc(cc1O)CCC(=O)c1c(cc(cc1O)OC1OC(CO)C(O)C(O)C1OC1OC(C)C(O)C(O)C1O)O',
'sweetness'),
('COc1cccc(c1O)CCC(=O)c1c(cc(cc1O)OC1OC(CO)C(O)C(O)C1OC1OC(C)C(O)C(O)C1O)O',
'sweetness'),
('COc1ccc(cc1O)CCC(=O)c1c(cc(cc1O)OC1OC(CO)C(O)C(O)C1OC1OC(C)C(O)C(O)C1O)O',
'sweetness'), ('COC(=O)C(Cc1ccccc1)NC(=O)C(CC(O)=O)NCCC(C)(C)C', 'sweetness'),
('NC(CC(=O)c1ccccc1NC=O)C(O)=O', 'sweetness'), ('[Na+].CC(C)CNS([O-])(=O)=O',
'sweetness'), ('[Na+].CC(C)CCNS([O-])(=O)=O', 'sweetness'),
('Nc1cccc(c1)N(=O)=O', 'sweetness'), ('O=N(=O)c1ccccc1', 'sweetness'),
('[Na+].CCCNS([O-])(=O)=O', 'sweetness'), ('CC1CCC(OC1OC1OC(C)C(O)C(O)C1O)C(C)C1
CCC2C3CC(=O)C4CC(CCC4(C)C3CCC12C)OC1OC(CO)C(O)C(O)C1OC1OC(C)C(O)C(O)C1O',
'sweetness'), ('CCCOc1ccc(cc1N)N(=O)=O', 'sweetness'),
('OCC1OC(OCC2OC(O)(CO)C(O)C2O)C(O)C(O)C1O', 'sweetness'),
('Nc1cccc2c1C(=O)NS2(=O)=O', 'sweetness'), ('Cc1ccc(cc1)NC(N)=O', 'sweetness'),
('CC1(C)C(CCC2(C=O)C1CCC1(C)C2CCC2C3=CC(C)(CCC3(C)CCC12C)C(O)=O)OC1OC(C(O)C(O)C1
OC1OC(C(O)C(O)C1O)C(O)=O)C(O)=O', 'sweetness'), ('CC1(C)C(CCC2(C=O)C1CCC1(C)C2CC
=C2C3CC(C)(CCC3(C)CCC12C)C(O)=O)OC1OC(C(O)C(O)C1OC1OC(C(O)C(O)C1O)C(O)=O)C(O)=O'
, 'sweetness'), ('CC1(C)C(CCC2(CO)C1CCC1(C)C2CCC2C3=CC(C)(CCC3(C)CCC12C)C(O)=O)O
C1OC(C(O)C(O)C1OC1OC(C(O)C(O)C1O)C(O)=O)C(O)=O', 'sweetness'), ('CC1(C)C(CCC2(CO
)C1CCC1(C)C2CC=C2C3CC(C)(CCC3(C)CCC12C)C(O)=O)OC1OC(C(O)C(O)C1OC1OC(C(O)C(O)C1O)
C(O)=O)C(O)=O', 'sweetness'), ('CC1(C)C(CCC2(C=O)C1CCC1(C)C2CCC2C3=CC(C)(CCC3(C)
CCC12C)C(O)=O)OC1OC(C(O)C(O)C1OC1OCC(O)C(O)C1O)C(O)=O', 'sweetness'),
('CC(=C)C1CCC(=CC1)C=O', 'sweetness'), ('CC(=C)C1CCC(=CC1)C=NO', 'sweetness'),
('COc1ccc(cc1O)C(=O)Oc1ccccc1', 'sweetness'),
('OC(=O)CNC(NCC1CCCCC1)=Nc1ccc(cc1)N(=O)=O', 'sweetness'),
('OC(=O)CNC(NCC1CCCCC1)=Nc1ccccc1', 'sweetness'),
('OC(=O)CNC(NS(=O)(=O)c1ccccc1)=Nc1cccc(c1)Cl', 'sweetness'),
('OC(=O)CNC(NS(=O)(=O)c1ccccc1)=Nc1ccccc1', 'sweetness'),
('OC(=O)CNC(NC1CCCCC1)=Nc1ccccc1', 'sweetness'),
('OC(=O)CNC(Nc1ccccc1)=Nc1ccccc1', 'sweetness'),
('CN(C1CCCCCCC1)C(NCC(O)=O)=Nc1ccc(cc1)C#N', 'sweetness'),
('CN(Cc1ccccc1)C(NCC(O)=O)=Nc1cc(cc(c1)Cl)Cl', 'sweetness'),
('CN(Cc1ccccc1)C(NCC(O)=O)=Nc1ccc(cc1)C#N', 'sweetness'),
('CCN(Cc1ccccc1)C(NCC(O)=O)=Nc1ccc(cc1)C#N', 'sweetness'),
('CC(NC(NCC(O)=O)=Nc1ccc(cc1)N(=O)=O)c1ccccc1', 'sweetness'),
('CC(NC(NCC(O)=O)=Nc1ccccc1)c1ccccc1', 'sweetness'),
('CC(NC(NCC(O)=O)=Nc1cccc(c1)Cl)c1ccccc1', 'sweetness'),
('CC(NC(NCC(O)=O)=Nc1cccc(c1)Br)c1ccccc1', 'sweetness'),

```
('CC(NC(NCC(O)=O)=Nc1cccc(c1)C)c1ccccc1', 'sweetness'),
('CC(NC(NCC(O)=O)=Nc1cccc(c1)C(F)(F)F)c1ccccc1', 'sweetness'),
('CC(NC(NCC(O)=O)=Nc1cccc(c1)N(=O)=O)c1ccccc1', 'sweetness'),
('CC(NC(NCC(O)=O)=Nc1cccc(c1)C#N)c1ccccc1', 'sweetness'),
('CSc1cccc(c1)N=C(NCC(O)=O)NC(C)c1ccccc1', 'sweetness'),
('CC(NC(NCC(O)=O)=Nc1cc(cc(c1)C)C)c1ccccc1', 'sweetness'),
('CC(NC(NCC(O)=O)=Nc1cc(cc(c1)F)F)c1ccccc1', 'sweetness'),
('CC(NC(NCC(O)=O)=Nc1ccc(c(c1)C)C#N)c1ccccc1', 'sweetness'),
('CC(NC(NCC(O)=O)=Nc1cc(c(c(c1)C)C#N)C)c1ccccc1', 'sweetness'),
('CC(NC(NCC(O)=O)=Nc1cc(c(c(c1)Cl)Cl)Cl)c1ccccc1', 'sweetness'),
('OCC1OC(Oc2cc(cc(c2C(=O)CCc2ccc(cc2)O)O)O)C(O)C(O)C1O', 'sweetness'),
('Oc1cc(cc(c1)O)O', 'sweetness'), ('COc1ccc(cc1)OC(=O)c1ccccc1C(O)=O',
'sweetness'), ('COc1ccc(cc1O)C1Cc2cccc(c2C(=O)O1)O', 'tastelessness'), ('CC1CCC(
OC1OC1OC(C)C(O)C(O)C1O)C(C)C1CCC2C3=CC(=O)C4CC(CCC4(C)C3CCC12C)OC1OC(CO)C(O)C(O)
C1OC1OC(C)C(O)C(O)C1O', 'sweetness'), ('CC1CCC(OC1OC1OC(C)C(O)C(O)C1O)C(C)C1CCC2
C3=CC(=O)C4CC(CCC4(C)C3CCC12C)OC1OC(CO)C(O)C(O)C1O', 'sweetness'),
('COc1ccc(cc1)CCC(=O)c1c(cc(cc1O)OC1OC(CO)C(O)C(O)C1OC1OC(C)C(O)C(O)C1O)O',
'sweetness'), ('[K+].COc1ccc(cc1O)CCC(=O)c1c(cc(cc1O)OCCCP(O)([O-])=O)O',
'sweetness'), ('[K+].COc1ccc(cc1O)CCC(=O)c1c(cc(cc1O)OCCCS([O-])(=O)=O)O',
'sweetness'), ('[K+].CC1(C)C(CCC2(C)C1CCC1(C)C2C(=O)C=C2C3CC(C)(CCC3(C)CCC12C)C(
O)=O)OC1OC(C(O)C(O)C1OC1OC(C(O)C(O)C1O)C(O)=O)C([O-])=O', 'sweetness'),
('[K+].O=C1[N-]S(=O)(=O)c2ccccc12', 'sweetness'),
('OCC1OC(Oc2cc(c(c(c2)O)C(=O)CCc2ccc(cc2)O)O)C(O)C(O)C1O', 'sweetness'),
('OCC1(O)CCC(O)C(O)C1O', 'sweetness'), ('OCC1CC(O)C(O)C(O)C1O', 'sweetness'), ('
CC1OC(OC2CC3C(C)(CCC(O)=O)C(CCC3(C)C3(C)CCC(C23)C(C)(O)CC=CC(C)(C)O)C(C)=C)C(O)C
(O)C1O', 'sweetness'), ('CC(=C)C1CCC2(C)C(CC(OC3OCC(O)C(O)C3O)C3C(CCC23C)C(C)(O)
CC=CC(C)(C)O)C1(C)CCC(O)=O', 'sweetness'),
('OCC1OC(OCC2OC(OC3(CO)OC(CO)C(O)C3O)C(O)C(O)C2O)C(O)C(O)C1O', 'sweetness'), ('C
C12CCCC(C)(C2CCC23CC(=C)C(CCC12)(C3)OC1OC(CO)C(O)C(OC2OC(CO)C(O)C(O)C2O)C1OC1OC(
CO)C(O)C(O)C1O)C(=O)OC1OC(CO)C(O)C(O)C1O', 'sweetness'), ('[Na+].CC12CCCC(C)(C2C
CC23CC(=C)C(CCC12)(C3)OC1OC(CO)C(O)C(O)C1OC1OC(CO)C(O)C(O)C1O)C(=O)OCCCS([O-])(=
O)=O', 'sweetness'), ('CC12CCCC(C)(C2CCC23CC(=C)C(CCC12)(C3)OC1OC(CO)C(O)C(OC2OC
(CO)C(O)C(O)C2O)C1OC1OC(CO)C(O)C(O)C1O)C(O)=O', 'sweetness'), ('CC1OC(OC2C(OC(CO
)C(O)C2OC2OC(CO)C(O)C(O)C2O)OC23CCC4C5(C)CCCC(C)(C5CCC4(CC2=C)C3)C(=O)OC2OC(CO)C
(O)C(O)C2O)C(O)C(O)C1O', 'sweetness'), ('CC12CCCC(C)(C2CCC23CC(=C)C(CCC12)(C3)OC
1OC(CO)C(O)C(OC2OC(CO)C(O)C(O)C2O)C1OC1OC(CO)C(O)C(O)C1O)C(=O)OC1OC(CO)C(O)C(O)C
1OC1OC(CO)C(O)C(O)C1O', 'sweetness'), ('CC12CCCC(C)(C2CCC23CC(=C)C(CCC12)(C3)OC1
OC(CO)C(O)C(O)C1OC1OC(CO)C(O)C(O)C1O)C(=O)OC1OC(CO)C(O)C(O)C1OC1OC(CO)C(O)C(O)C1
O', 'sweetness'), ('CC12CCCC(C)(C2CCC23CC(=C)C(CCC12)(C3)OC1OC(CO)C(O)C(OC2OC(CO
)C(O)C(O)C2O)C1OC1OCC(O)C(O)C1O)C(=O)OC1OC(CO)C(O)C(O)C1O', 'sweetness'), ('CC12
CCCC(C)(C2CCC23CC(=C)C(CCC12)(C3)OC1OC(CO)C(O)C(OC2OC(CO)C(O)C(O)C2O)C1O)C(=O)OC
1OC(CO)C(O)C(O)C1O', 'sweetness'), ('CC1OC(OC2C(OC(CO)C(O)C2OC2OC(CO)C(O)C(O)C2O
)OC23CCC4C5(C)CCCC(C)(C5CCC4(CC2=C)C3)C(=O)OC2OC(CO)C(O)C(O)C2O)C(O)C(OC2OC(CO)C
(O)C(O)C2O)C1O', 'sweetness'), ('CC12CCCC(C)(C2CCC23CC(=C)C(CCC12)(C3)OC1OC(CO)C
(O)C(OC2OC(CO)C(O)C(O)C2O)C1OC1OC(CO)C(O)C(O)C1O)C(=O)OC1OC(CO)C(O)C(OC2OC(CO)C(
O)C(O)C2O)C1O', 'sweetness'), ('CC1OC(OC2C(O)C(O)C(CO)OC2OC(=O)C2(C)CCCC3(C)C2CC
C24CC(=C)C(CCC32)(C4)OC2OC(CO)C(O)C(OC3OC(CO)C(O)C(O)C3O)C2OC2OC(CO)C(O)C(O)C2O)
```

C(O)C(O)C1O', 'sweetness'), ('CC1OC(OC2C(OC(CO)C(O)C2OC2OC(CO)C(O)C(O)C2O)OC23CC
C4C5(C)CCCC(C)(C5CCC4(CC2=C)C3)C(=O)OC2OC(CO)C(O)C(O)C2OC2OC(CO)C(O)C(O)C2O)C(O)
C(O)C1O', 'sweetness'), ('CC12CCCC(C)(C2CCC23CC(=C)C(CCC12)(C3)OC1OC(CO)C(O)C(OC
2OC(CO)C(O)C(O)C2O)C1OC1OC(COC2OC(CO)C(O)C(O)C2O)C(O)C(O)C1O)C(=O)OC1OC(CO)C(O)C
(O)C1O', 'sweetness'), ('CC12CCCC(C)(C2CCC23CC(=C)C(CCC12)(C3)OC1OC(CO)C(O)C(OC2
OC(CO)C(O)C(O)C2O)C1OC1OC(CO)C(O)C(O)C1O)C(=O)OC1OC(CO)C(O)C(OC2OC(CO)C(O)C(O)C2
O)C1OC1OC(CO)C(O)C(O)C1O', 'sweetness'), ('CC1OC(OC2C(OC(CO)C(O)C2OC2OC(CO)C(O)C
(O)C2O)OC(=O)C2(C)CCCC3(C)C2CCC24CC(=C)C(CCC32)(C4)OC2OC(CO)C(O)C(OC3OC(CO)C(O)C
(O)C3O)C2OC2OC(CO)C(O)C(O)C2O)C(O)C(O)C1O', 'sweetness'), ('CC1OC(OC2C(OC(CO)C(O
)C2OC2OC(CO)C(O)C(O)C2O)OC(=O)C2(C)CCCC3(C)C2CCC24CC(=C)C(CCC32)(C4)OC2OC(CO)C(O
)C(OC3OC(CO)C(O)C(O)C3O)C2OC2OC(CO)C(O)C(O)C2O)C(O)C(OC2OC(CO)C(O)C(O)C2O)C1O',
'sweetness'), ('CC12CCCC(C)(C2CCC23CC(=C)C(CCC12)(C3)OC1OCC(O)C(OC2OC(CO)C(O)C(O
)C2O)C1OC1OC(CO)C(O)C(O)C1O)C(=O)OC1OC(CO)C(O)C(O)C1O', 'sweetness'), ('CC1OC(OC
2(O)C(O)C(O)C(CO)OC2OC(=O)C2(C)CCCC3(C)C2CCC24CC(=C)C(CCC32)(C4)OC2OC(CO)C(O)C(O
)C2OC2OC(CO)C(O)C(O)C2O)C(O)C(O)C1O', 'sweetness'), ('CC12CCCC(C)(C2CCC23CC(=C)C
(CCC12)(C3)OC1OC(CO)C(O)C(OC2OC(CO)C(O)C(O)C2O)C1OC1OC(CO)C(O)C(O)C1O)C(=O)OC1OC
(CO)C(O)C(OC2OC(CO)C(O)C(O)C2O)C1OC1OCC(O)C(O)C1O', 'sweetness'), ('CC12CCCC(C)(
C2CCC23CC(=C)C(CCC12)(C3)OC1OC(CO)C(O)C(OC2OC(CO)C(O)C(O)C2O)C1OC1OC(CO)C(O)C(O)
C1O)C(=O)OC1OC(COC2OCC(O)C(O)C2O)C(O)C(O)C1O', 'sweetness'), ('CC12CCCC(C)(C2CCC
23CC(=C)C(CCC12)(C3)OC1OC(CO)C(O)C(OC2OC(CO)C(O)C(O)C2O)C1OC1OCC(O)C(O)C1O)C(=O)
OC1OC(CO)C(O)C(O)C1OC1OC(CO)C(O)C(O)C1O', 'sweetness'), ('CC1OC(OC2C(OC(CO)C(O)C
2OC2OC(CO)C(O)C(O)C2O)OC(=O)C2(C)CCCC3(C)C2CCC24CC(=C)C(CCC32)(C4)OC2OC(CO)C(O)C
(OC3OC(CO)C(O)C(O)C3O)C2OC2OCC(O)C(O)C2O)C(O)C(O)C1O', 'sweetness'), ('CC12CCCC(
C)(C2CCC23CC(=C)C(CCC12)(C3)OC1OC(CO)C(O)C(O)C1OC1OC(CO)C(O)C(O)C1O)C(=O)OC1OC(C
OC2OC(CO)C(O)C(O)C2O)C(O)C(O)C1O', 'sweetness'), ('CC12CCCC(C)(C2CCC23CC(=C)C(CC
C12)(C3)OC1OC(COC2OC(CO)C(O)C(O)C2O)C(O)C(O)C1OC1OC(CO)C(O)C(O)C1O)C(=O)OC1OC(CO
)C(O)C(O)C1O', 'sweetness'), ('Oc1cccc(c1)O', 'sweetness'), ('CC12CCCC(C)(C2CCC2
3CC(=C)C(CCC12)(C3)OC1OC(CO)C(O)C(O)C1O)C(=O)OC1OC(CO)C(O)C(O)C1O',
'sweetness'), ('O=C1NS(=O)(=O)c2ccccc12', 'sweetness'),
('OCC12OCC(O1)C(O)C(O)C2O', 'sweetness'), ('OC1Cc2c(cc(c(c2OC1c1ccc(cc1)O)C1C(O)
C(Oc2c1c(cc1c2C2C(O)C(Oc3cc(cc(c23)O)O)(O1)c1ccc(cc1)O)O)c1ccc(cc1)O)O)O',
'miscellaneous'), ('CC(CCC(OC1OC(COC2OC(CO)C(O)C(O)C2O)C(O)C(O)C1OC1OC(CO)C(O)C(
O)C1O)C(C)(C)O)C1CCC2(C)C3CC=C4C(CCC(OC5OC(CO)C(O)C(O)C5O)C4(C)C)C3(C)C(O)CC12C'
, 'sweetness'), ('[Na+].CC(O)CNS([O-])(=O)=O', 'sweetness'),
('[Na+].O=C1[N-]S(=O)(=O)OC=C1', 'sweetness'), ('[Na+].OCCCCCNS([O-])(=O)=O',
'sweetness'), ('[Na+].CC1=C(C)C(=O)[N-]S(=O)(=O)O1', 'sweetness'),
('[Na+].CCC1=COS(=O)(=O)[N-]C1=O', 'sweetness'),
('[Na+].CCC1=C(C)OS(=O)(=O)[N-]C1=O', 'sweetness'),
('[Na+].CC1=COS(=O)(=O)[N-]C1=O', 'sweetness'),
('[Na+].CC(C)C1=C(C)C(=O)[N-]S(=O)(=O)O1', 'sweetness'),
('[Na+].CCC1=CC(=O)[N-]S(=O)(=O)O1', 'sweetness'),
('[Na+].CCCC1=C(C)OS(=O)(=O)[N-]C1=O', 'sweetness'),
('[Na+].[O-]S(=O)(=O)NC1CCCCC1', 'sweetness'),
('[Na+].[O-]S(=O)(=O)Nc1cccc(c1)Br', 'sweetness'),
('[Na+].[O-]S(=O)(=O)Nc1cccc(c1)Cl', 'sweetness'),
('[Na+].[O-]S(=O)(=O)Nc1cccc(c1)C#N', 'sweetness'),
('[Na+].[O-]S(=O)(=O)Nc1cccc(c1)F', 'sweetness'),

('[Na+].Cc1cccc(c1)NS([O-])(=O)=O', 'sweetness'),
('[Na+].[O-]S(=O)(=O)NC1=NN=CS1', 'sweetness'),
('[Na+].[O-]S(=O)(=O)NC1=NC=CS1', 'sweetness'),
('[Na+].CCOC(=O)N1CCC(CC1)NS([O-])(=O)=O', 'sweetness'),
('[Na+].CC(C)(C)CNS([O-])(=O)=O', 'sweetness'),
('[Na+].[O-]S(=O)(=O)Nc1cccc2c1CCC2', 'sweetness'),
('[Na+].Cc1ccc(c(c1)Br)NS([O-])(=O)=O', 'sweetness'),
('[Na+].CCOc1ccccc1NS([O-])(=O)=O', 'sweetness'),
('[Na+].CCc1ccc(cc1NS([O-])(=O)=O)C', 'sweetness'),
('[Na+].CCC(CC)CNS([O-])(=O)=O', 'sweetness'), ('[Na+].CCCC(C)CNS([O-])(=O)=O', 'sweetness'), ('[Na+].Cc1ccccc1NS([O-])(=O)=O', 'sweetness'),
('[Na+].CC1CC(CCS1)NS([O-])(=O)=O', 'sweetness'),
('[Na+].CC(C)c1ccccc1NS([O-])(=O)=O', 'sweetness'),
('[Na+].CCCc1ccccc1NS([O-])(=O)=O', 'sweetness'),
('[Na+].[O-]S(=O)(=O)Nc1cc(cc(c1)Cl)Cl', 'sweetness'),
('[Na+].[O-]S(=O)(=O)Nc1cc(cc(c1)F)F', 'sweetness'),
('[Na+].CCOc1cccc(c1)NS([O-])(=O)=O', 'sweetness'),
('[Na+].CCc1cccc(c1)NS([O-])(=O)=O', 'sweetness'),
('[Na+].COc1cc(cc(c1)NS([O-])(=O)=O)C', 'sweetness'),
('[Na+].CC1CCCC(C1)NS([O-])(=O)=O', 'sweetness'),
('[Na+].CC1CCC(C1)NS([O-])(=O)=O', 'sweetness'),
('[Na+].CC(C)c1cccc(c1)NS([O-])(=O)=O', 'sweetness'),
('[Na+].CC1=C(C)N=C(NS([O-])(=O)=O)S1', 'sweetness'),
('[Na+].CCc1cc(ccc1NS([O-])(=O)=O)Br', 'sweetness'),
('[Na+].Cc1cc(ccc1NS([O-])(=O)=O)Br', 'sweetness'),
('[Na+].Cc1cc(ccc1Br)NS([O-])(=O)=O', 'sweetness'),
('[Na+].CCCCOC(=O)c1ccc(cc1)NS([O-])(=O)=O', 'sweetness'),
('[Na+].CCCCC1=CSC(=N1)NS([O-])(=O)=O', 'sweetness'),
('[Na+].COc1cccc2c1N=C(NS([O-])(=O)=O)S2', 'sweetness'),
('[Na+].Cc1cccc2c1N=C(NS([O-])(=O)=O)S2', 'sweetness'),
('[Na+].CC1=CSC(=N1)NS([O-])(=O)=O', 'sweetness'),
('[Na+].[O-]S(=O)(=O)Nc1ccc(cc1)N1CCOCC1', 'sweetness'),
('[Na+].CCCCCC1=CSC(=N1)NS([O-])(=O)=O', 'sweetness'),
('[Na+].CC(C)OC(=O)c1ccc(cc1)NS([O-])(=O)=O', 'sweetness'),
('[Na+].CCCC1=CSC(=N1)NS([O-])(=O)=O', 'sweetness'),
('[Na+].[O-]S(=O)(=O)NC1=NN=C(S1)C1CCC1', 'sweetness'),
('[Na+].CCC1=NN=C(NS([O-])(=O)=O)S1', 'sweetness'),
('[Na+].COC(=O)c1ccc(c(c1)NS([O-])(=O)=O)C', 'sweetness'),
('[Na+].CC1CC(CS1)NS([O-])(=O)=O', 'sweetness'),
('[Na+].CCCC1=NN=C(NS([O-])(=O)=O)S1', 'sweetness'),
('[Na+].CCCCCCCCCCCCC1=NN=C(NS([O-])(=O)=O)S1', 'sweetness'),
('[Na+].[O-]S(=O)(=O)NC1=Nc2ccc(cc2S1)Cl', 'sweetness'),
('[Na+].CCc1cccc(n1)NS([O-])(=O)=O', 'sweetness'),
('[Na+].Cc1ccc2c(c1)SC(=N2)NS([O-])(=O)=O', 'sweetness'),
('[Na+].CCCc1cccc(n1)NS([O-])(=O)=O', 'sweetness'),
('[Na+].CCNc1ccccc1NS([O-])(=O)=O', 'sweetness'), ('[Na+].CCSCCNS([O-])(=O)=O', 'sweetness'), ('[Na+].CSCCNS([O-])(=O)=O', 'sweetness'),

```
('[Na+].CC(O)c1cccc(c1)NS([O-])(=O)=O', 'sweetness'),
('[Na+].[O-]S(=O)(=O)Nc1cccc(c1)OC(F)(F)F', 'sweetness'),
('[Na+].CC(C)COC(=O)c1ccc(cc1)NS([O-])(=O)=O', 'sweetness'),
('[Na+].CC(C)CC1=CSC(=N1)NS([O-])(=O)=O', 'sweetness'),
('[Na+].CN(C)c1ccc(cc1)NS([O-])(=O)=O', 'sweetness'),
('[Na+].CC(C)c1ccc(cc1)NS([O-])(=O)=O', 'sweetness'),
('[Na+].CCSC1=NN=C(NS([O-])(=O)=O)S1', 'sweetness'),
('[Na+].CC(C)C1=NN=C(NS([O-])(=O)=O)S1', 'sweetness'),
('[Na+].[O-]S(=O)(=O)NC1=NN=C(S1)C(F)(F)F', 'sweetness'),
('[Na+].[O-]S(=O)(=O)Nc1ccccc1', 'sweetness'),
('[Na+].O=C1[N-]S(=O)(=O)c2ccccc12', 'sweetness'), ('OCC(O)C(O)C(O)C(=O)CO',
'sweetness'), ('COc1cc2c(cc1O)C1(CC2)SCc2ccccc2S1', 'sweetness'),
('COCC1=CCC(=CC1)C=NO', 'sweetness'), ('OCC1OC(OCC2OC(OCC3OC(OC4(CO)OC(CO)C(O)C4
O)C(O)C(O)C3O)C(O)C(O)C2O)C(O)C(O)C1O', 'sweetness'),
('CC12CCCC(C)(C2CCC23CC(=C)C(O)(CCC12)C3)C(O)=O', 'sweetness'), ('CC12CCCC(C)(C2
CCC23CC(=C)C(CCC12)(C3)OC1OC(CO)C(O)C(O)C1OC1OC(CO)C(O)C(O)C1O)C(O)=O',
'sweetness'), ('CC12CCCC(C)(C2CCC23CC(=C)C(CCC12)(C3)OC1OC(CO)C(O)C(O)C1OC1OC(CO
)C(O)C(O)C1O)C(=O)OC1OC(CO)C(O)C(O)C1O', 'sweetness'), ('CC12CCCC(C)(C2CCC23CC(=
C)C(CCC12)(C3)OC1OC(CO)C(O)C(O)C1OC1OC(CO)C(O)C(O)C1OC1OC(CO)C(OC2OC(CO)C(O)C(O)
C2O)C(O)C1O)C(=O)OC1OC(CO)C(O)C(O)C1O', 'sweetness'), ('CC1OC(OC2C(O)C3(C)CCC4(C
)C(=CCC5C6(C)CCC(OC7OC(C(O)C(O)C7OC7OCC(O)C(O)C7O)C(O)=O)C(C)(CO)C6CCC45C)C3CC2(
C)CO)C(OC(C)=O)C(OC(C)=O)C1OC(C)=O', 'sweetness'), ('CC1OC(OC2C(O)C3(C)CCC4(C)C(
=CCC5C6(C)CCC(OC7OC(C(O)C(O)C7O)C(O)=O)C(C)(CO)C6CCC45C)C3CC2(C)CO)C(OC(C)=O)C(O
C(C)=O)C1OC(C)=O', 'sweetness'), ('CC1OC(OC2C(O)C3(C)CCC4(C)C(=CCC5C6(C)CCC(OC7O
C(C(O)C(O)C7OC7OC(CO)C(O)C(O)C7O)C(O)=O)C(C)(CO)C6CCC45C)C3CC2(C)CO)C(OC(C)=O)C(
OC(C)=O)C1OC(C)=O', 'sweetness'), ('OCC1(OC(CCl)C(O)C1O)OC1OC(CCl)C(Cl)C(O)C1O',
'sweetness'), ('CC1OC(OC2(CCl)OC(CCl)C(O)C2O)C(O)C(O)C1Cl', 'sweetness'),
('COCC1OC(OC2(CCl)OC(CCl)C(O)C2O)C(O)C(O)C1Cl', 'sweetness'),
('OC(=O)CNC(NC1CCCCCCC1)=Nc1ccc(cc1)C#N', 'sweetness'),
('CC(C)(C)C(C(O)=O)S(=O)(=O)CCc1ccccc1', 'sweetness'),
('[Na+].[O-]C(=O)CCNC(=O)Nc1ccc(cc1)N(=O)=O', 'sweetness'),
('OC(=O)CNC(=O)Nc1ccc(cc1)N(=O)=O', 'sweetness'),
('COC(=O)C(Cc1ccccc1)NC(=O)C(CC(O)=O)NC(Nc1ccc(cc1)C#N)=NC#N', 'sweetness'),
('COC(=O)C(Cc1ccccc1)NC(=O)C(CC(O)=O)NC(=S)Nc1ccc(cc1)C(=O)OC', 'sweetness'),
('COC(=O)C(Cc1ccccc1)NC(=O)C(CC(O)=O)NC(=N)Nc1ccc(cc1)C#N', 'sweetness'),
('CC(NC(=O)C(CC(O)=O)NC(Nc1ccc(cc1)C#N)=NC#N)c1ccccc1', 'sweetness'),
('COC(=O)C(Cc1ccccc1)NC(=O)C(CC(O)=O)NC(=O)Nc1ccc(cc1)C#N', 'sweetness'), ('CCC(
C)C(=O)OC(C)C1(O)CCC2(O)C3CCC4CC(CCC4(C)C3CC(OC(C)=O)C12C)OC1CC(O)C(OC2CC(OC)C(O
C3CC(OC)C(OC4OC(CO)C(O)C(O)C4O)C(C)O3)C(C)O2)C(C)O1', 'sweetness'), ('CCC(C)C(=O
)OC(C)C1(O)CCC2(O)C3CCC4CC(CCC4(C)C3CC(OC(C)=O)C12C)OC1CC(O)C(OC2CC(OC)C(OC3OC(C
)C(OC4OC(CO)C(O)C(O)C4O)C(OC)C3O)C(C)O2)C(C)O1', 'sweetness'), ('CCC(C)C(=O)OC(C
)C1(O)CCC2(O)C3CCC4CC(CCC4(C)C3CC(OC(C)=O)C12C)OC1CC(O)C(OC2CC(OC)C(OC3CC(OC)C(O
C4CC(OC)C(OC5CC(OC)C(O)C(C)O5)C(C)O4)C(C)O3)C(C)O2)C(C)O1', 'sweetness'), ('CCC(
C)C(=O)OC(C)C1(O)CCC2(O)C3CCC4CC(CCC4(C)C3CC(OC(C)=O)C12C)OC1CC(O)C(OC2CC(OC)C(O
C3CC(OC)C(OC4CC(OC)C(OC5OC(C)C(O)C(OC)C5O)C(C)O4)C(C)O3)C(C)O2)C(C)O1',
'sweetness'), ('CCC(C)C(=O)OC(C)C1(O)CCC2(O)C3CCC4CC(CCC4(C)C3CC(OC(C)=O)C12C)OC
1CC(O)C(OC2CC(OC)C(OC3CC(OC)C(OC4CC(OC)C(OC5OC(CO)C(O)C(O)C5O)C(C)O4)C(C)O3)C(C)
```

O2)C(C)O1', 'sweetness'), ('CCC(C)C(=O)OC(C)C1(O)CCC2(O)C3CCC4CC(CCC4(C)C3CC(OC(
C)=O)C12C)OC1CC(O)C(OC2CC(O)C(OC3CC(OC)C(OC4CC(OC)C(OC5OC(C)C(O)C(OC)C5O)C(C)O4)
C(C)O3)C(C)O2)C(C)O1', 'sweetness'), ('CCC(C)C(=O)OC(C)C1(O)CCC2(O)C3CCC4CC(CCC4
(C)C3CC(OC(C)=O)C12C)OC1CC(O)C(OC2CC(OC)C(OC3CC(OC)C(OC4CC(OC)C(OC5CC(OC)C(OC6OC
(CO)C(O)C(O)C6O)C(C)O5)C(C)O4)C(C)O3)C(C)O2)C(C)O1', 'sweetness'), ('CCC(C)C(=O)
OC(C)C1(O)CCC2(O)C3CCC4CC(CCC4(C)C3CC(OC(C)=O)C12C)OC1CC(O)C(OC2CC(OC)C(OC3CC(OC
)C(OC4CC(OC)C(OC5OC(C)C(OC6OC(CO)C(O)C(O)C6O)C(OC)C5O)C(C)O4)C(C)O3)C(C)O2)C(C)O
1', 'sweetness'), ('CCC(C)C(=O)OC(C)C1(O)CCC2(O)C3CCC4CC(CCC4(C)C3CC(OC(C)=O)C12
C)OC1CC(O)C(OC2CC(OC)C(OC3CC(OC)C(OC4CC(OC)C(OC5OC(CO)C(OC6OC(CO)C(O)C(O)C6O)C(O
)C5O)C(C)O4)C(C)O3)C(C)O2)C(C)O1', 'sweetness'), ('CCC(C)C(=O)OC(C)C1(O)CCC2(O)C
3CCC4CC(CCC4(C)C3CC(OC(C)=O)C12C)OC1CC(O)C(OC2CC(OC)C(OC3CC(OC)C(OC4OC(C)C(OC5OC
(CO)C(OC6OC(CO)C(O)C(O)C6O)C(O)C5O)C(OC)C4O)C(C)O3)C(C)O2)C(C)O1', 'sweetness'),
('ClC(Cl)C(Cl)Cl', 'sweetness'), ('[Na+].[O-]S(=O)(=O)NC1CCCSC1', 'sweetness'),
('[Na+].[O-]S(=O)(=O)NC1CCCSCCC1', 'sweetness'),
('OC(=O)CCNC(=S)Nc1ccc(cc1)C#N', 'sweetness'), ('O=C1NS(=O)(=O)C2=CSC=C12',
'sweetness'), ('OC(=O)CCNC(=S)Nc1ccc(cc1)N(=O)=O', 'sweetness'),
('COC(=O)C(Cc1ccccc1)NC(=O)C(CC(O)=O)NC(=S)Nc1ccc(cc1)C#N', 'sweetness'),
('COc1ccc(cc1)C=CC', 'sweetness'), ('O=CC=Cc1ccccc1', 'sweetness'),
('OCC1OC(OC2OC(CO)C(O)C(O)C2O)C(O)C(O)C1O', 'sweetness'),
('OCC(O)C(O)C(O)C(=O)COC1OC(CO)C(O)C(O)C1O', 'sweetness'), ('[K+].[K+].[K+].CC1(
C)C(CCC2(C)C1CCC1(C)C2C(=O)C=C2C3CC(C)(CCC3(C)CCC12C)C([O-])=O)OC1OC(C(O)C(O)C1O
C1OC(C(O)C(O)C1O)C([O-])=O)C([O-])=O', 'sweetness'),
('OCC1OC(OC2C(O)C(O)COC2(O)CO)C(O)C(O)C1O', 'sweetness'),
('OC1COC(OC2COC(O)C(O)C2O)C(O)C1O', 'sweetness'), ('COc1ccc(cc1O)CCc1ccccc1',
'sweetness'), ('OCC1OC2OC3C(O)C(O)C(OC3CO)OC3C(O)C(O)C(OC3CO)OC3C(O)C(O)C(OC3CO)
OC3C(O)C(O)C(OC3CO)OC3C(O)C(O)C(OC3CO)OC3C(O)C(O)C(OC3CO)OC1C(O)C2O',
'sweetness'), ('OCC1OC(O)C(O)C1O', 'sweetness'),
('COc1cc(ccc1O)OC1OC(COC(=O)c2cc(c(c(c2)O)O)O)C(O)C(O)C1O', 'bitterness'),
('CC(C)C12CC2C(C)C(=O)C1', 'bitterness'),
('OC1Cc2c(cc(cc2OC1c1ccc(c(c1)O)O)O)O', 'bitterness'),
('Oc1cc(c2c(c1)OC(C(C2)OC(=O)c1cc(c(c(c1)O)O)O)c1ccc(c(c1)O)O)O', 'bitterness'),
('OC1Cc2c(cc(cc2OC1c1cc(c(c(c1)O)O)O)O)O', 'bitterness'),
('CC1=CCCC2(C)OC2C2OC(=O)C(=C)C2CC1', 'bitterness'), ('COc1cc(cc(c1O)OC)C1C(COC2
OC(COC(=O)c3cc(c(c(c3)O)O)O)C(O)C(O)C2O)C(CO)Cc2cc(c(c(c12)OC)O)OC',
'bitterness'), ('OC1C(Oc2cc(cc(c2C1=O)O)O)c1ccc(c(c1)O)O', 'bitterness'),
('COc1cc2c(cc1OC)C1c3ccc(c(c3OCC1(O)C2)OC)OC', 'bitterness'),
('Oc1ccc(cc1)C1COc2cc(ccc2C1)O', 'bitterness'), ('CC(CCC1(O)OC2CC3C4CC=C5CC(CCC5
(C)C4CCC3(C)C2C1C)OC1OC(CO)C(OC2OC(C)C(O)C(O)C2O)C(O)C1O)COC1OC(CO)C(O)C(O)C1O',
'bitterness'), ('CC(CCC1(O)OC2CC3C4CCC5CC(CCC5(C)C4CCC3(C)C2C1C)OC1OC(CO)C(OC2OC
(C)C(O)C(O)C2O)C(O)C1O)COC1OC(CO)C(O)C(O)C1O', 'bitterness'),
('COS(=O)(=O)NC1CCCCC1C', 'bitterness'),
('OC(=O)C(CSCC=C)N1CC2=CC=C(C=O)N2C(CSCC=C)C1=O', 'bitterness'), ('CC1CCC2(OC1)O
C1CC3C4CCC5CC(CCC5(C)C4CCC3(C)C1C2C)OC1OC(CO)C(OC2OCC(O)C(O)C2O)C(O)C1OC1OC(CO)C
(O)C(O)C1O', 'bitterness'), ('O=C1C2CCCN2C(=O)C2CCCN12', 'bitterness'),
('CC1C2CC(OC(C)=O)C(=C)C(O)CCC(=CC2OC1=O)C', 'bitterness'),
('COCC(O)CCCc1ccccc1', 'bitterness'), ('CCCC(CCCOC)CCc1ccccc1', 'bitterness'),
('ON=CC1=CCOCC1', 'bitterness'), ('ON=CC1=CCSCC1', 'bitterness'),

('ON=CC1=CCCOC1', 'bitterness'), ('ON=CC1=CC2CCC1C=C2', 'bitterness'), ('COCC(C)(OC)C1CCC(=CC1)C=NO', 'bitterness'), ('COC(C)C1=CCC=C(C1)C=NO', 'bitterness'), ('COC(C)C1CCC=C(C1)C=NO', 'bitterness'), ('COCC1CCC=C(C1)C=NO', 'bitterness'), ('O=C1CN(CC2N1CCc1ccccc21)C(=O)C1CCCCC1', 'bitterness'), ('CC1=CCC2(O)C1C(O)C(C(O)C=C2C)C(=C)C(O)=O', 'bitterness'), ('CN(C)C(=N)N=C(N)N.OC(=O)CC(O)(CC(O)=O)C(O)=O', 'bitterness'), ('COc1cc2c(c3c1C(=O)CC(O3)c1ccc(cc1)O)CCC(C)(C)O2', 'bitterness'), ('COc1ccc(cc1)C=CC(=O)c1c(c2c(cc1OC)OC(C)(C)CC2)O', 'bitterness'), ('COc1c(c(c2c(c1C(=O)C=Cc1ccc(cc1)O)OC(C)(C)CC2)O)CC=C(C)C', 'bitterness'), ('COc1cc(c2c(c1C(=O)C=Cc1ccc(cc1)O)OC(C)(C)CC2)O', 'bitterness'), ('CN1C(=O)N(C)C2=C(N(C)C(=O)N2C)C1=O', 'bitterness'), ('CC(O)CCO', 'bitterness'), ('OCCCCCO', 'bitterness'), ('OC(=O)c1cccc2cccc(c12)C=O', 'bitterness'), ('OCC1C(=CC2=CC=CO2)C(=O)c2cc(c(c[n+]12)[O-])CC1=CC=CO1', 'bitterness'), ('CC1C2C(O)CC(=C3C(C2OC1=O)C(=CC3=O)CO)C', 'bitterness'), ('CCCCCC=CCC=CCCCCCCCC(O)CC(O)COC(C)=O', 'bitterness'), ('CC(=O)OCC(O)CC(O)CCCCCCCCCCCC=C', 'bitterness'), ('CC(=O)OCC(O)CC(O)CCCCCCCCCCC#C', 'bitterness'), ('CCCCCC=CCC=CCCCCCC=CC(=O)CC(O)COC(C)=O', 'bitterness'), ('CC(=O)OCC(O)CC(=O)CCCCCCCCCCCC=C', 'bitterness'), ('CCCCCCCCCCCCCC(=O)CC(O)COC(C)=O', 'bitterness'), ('CCCCCC=CCCCCCCCC(=O)CC(O)COC(C)=O', 'bitterness'), ('CC(C)C1=CC=C(C)CC1', 'bitterness'), ('COCC(C)(O)C1CCC(=CC1)C', 'bitterness'), ('COCC(O)CC(C)CCc1ccccc1', 'bitterness'), ('COc1cc(c(c(c1)O)CC=C(C)C)O', 'bitterness'), ('COCC(O)CCCCc1ccccc1', 'bitterness'), ('CN1C(=O)C=Cc2ccccc12', 'bitterness'), ('COC(C)C1=CCC=C(C)C1', 'bitterness'), ('COC(C)C1CCC=C(C)C1', 'bitterness'), ('OC(=O)c1cccc2ccccc12', 'bitterness'), ('O=N(=O)c1cccc2ccccc12', 'bitterness'), ('CCC(C)C(=O)c1c(cc(cc1OC1OC(CO)C(O)C(O)C1O)O)O', 'bitterness'), ('CC(C)C(=O)c1c(cc(cc1OC1OC(CO)C(O)C(O)C1O)O)O', 'bitterness'), ('CCCCCCCCC=CCCCCCCCC(=O)OCC(O)CO', 'bitterness'), ('COc1cc(cc(c1O)OC)C=CC(=O)OC1OC(CO)C(O)C(O)C1O', 'bitterness'), ('CCC(O)C=C', 'bitterness'), ('CC(C)C(C(O)=O)S(=O)(=O)c1ccc(cc1Cl)Cl', 'bitterness'), ('COc1ccc(c(c1)OC)C(=O)c1ccccc1C(O)=O', 'bitterness'), ('CC(C)C(C(O)=O)S(=O)(=O)c1ccc(c(c1)Cl)Cl', 'bitterness'), ('OC(=O)c1ccccc1C(=O)c1ccc(cc1)N(=O)=O', 'bitterness'), ('CC(O)(CO)C1CCC=CC1', 'bitterness'), ('Oc1ccc(c(c1)O)C(=O)C=Cc1ccccc1O', 'bitterness'), ('CC(C)=CCc1c(c(c(c2c1OC(C)(C)CC2=O)O)O)O', 'bitterness'), ('CC1(O)CC(=C(N2CCCC2)C1=O)N1CCCC1', 'bitterness'), ('CCCOc1ccc(cc1O)CCC(=O)c1ccccc1O', 'bitterness'), ('COc1cc(c(c(c1C(=O)C=Cc1ccc(cc1)O)O)CC1OC1(C)C)O', 'bitterness'), ('CCCOc1ccc(cc1N(=O)=O)N(=O)=O', 'bitterness'), ('CC1CCCC(C)N1', 'bitterness'), ('CCCCN(CCCC)CCN1C(=O)c2ccccc2S1(=O)=O', 'bitterness'), ('CCN(CC)CCN1C(=O)c2ccccc2S1(=O)=O', 'bitterness'), ('CCN(CC)CCCN1C(=O)c2ccccc2S1(=O)=O', 'bitterness'), ('CC(O)(CO)C1CCC(=CC1)C=NO', 'bitterness'), ('COCC(C)(O)C1CCC(=CC1)C=NO', 'bitterness'), ('COC(C)(CO)C1CCC(=CC1)C=NO', 'bitterness'), ('CC(C)(O)C1CCC(=CC1)C=NO', 'bitterness'), ('NC1=NC=C(S1)N(=O)=O', 'bitterness'), ('CCCCCC(C)O', 'bitterness'), ('CCCCCCCC1=C(O)C(=O)c2ccccc2N1',

'bitterness'), ('COc1c(c(c(c(c1C(=O)C=Cc1ccc(cc1)O)O)CC(O)C(C)(C)OC)O)C',
'bitterness'), ('COC(C)(CO)C1CCC(=CC1)C', 'bitterness'), ('COc1cc(ccc1O)C',
'bitterness'), ('COc1ccc(cc1O)CC(C)c1ccccc1', 'bitterness'), ('CC1=CC2CCC1C=C2',
'bitterness'), ('OCC1OC(OC2=Cc3ccccc3C=C2)C(O)C(O)C1O', 'bitterness'),
('OCC1OC(Oc2ccccc2N(=O)=O)C(O)C(O)C1O', 'bitterness'), ('Nc1ccccc1N(=O)=O',
'bitterness'), ('CCCCCCC(C)=O', 'bitterness'),
('CCCCCCCCC=CCCCCCCCC(=O)OC(CO)CO', 'bitterness'), ('O=C1NC(=S)NC=C1',
'bitterness'), ('OC(=O)CCNC(=O)Cc1ccc(cc1)N(=O)=O', 'bitterness'),
('COc1ccc(cc1OC)C1Cc2cccc(c2C(=O)O1)O', 'bitterness'),
('CC(CC(O)=O)S(=O)(=O)c1ccc(cc1)Cl', 'bitterness'),
('CC(C)OC1=NS(=O)(=O)c2ccccc12', 'bitterness'), ('Cc1ccc(cc1)S(=O)(=O)CCC(O)=O',
'bitterness'), ('Oc1cccc(c1)C=CC(=O)c1ccccc1O', 'bitterness'),
('OC(=O)c1ccc(c(c1)N(=O)=O)N(=O)=O', 'bitterness'), ('OC(=O)c1cc(cc(c1O)I)I',
'bitterness'), ('Oc1ccc2c(c1)C(=O)C(=C(O2)c1ccc(c(c1)O)O)O', 'bitterness'),
('CC1=CCOCC1', 'bitterness'), ('Oc1ccc(cc1)C1=C(O)C(=O)c2ccc(cc2O1)O',
'bitterness'), ('CC(C)CCCC(C)CCO', 'bitterness'),
('OC(=O)CCN(C1CCCCC1)C(=O)Nc1ccc(cc1)C#N', 'bitterness'),
('OC(=O)CCN(Cc1ccccc1)C(=O)Nc1ccc(cc1)C#N', 'bitterness'),
('CC(CC1CCCCC1)N(C)C(=O)C(N)CC(O)=O', 'bitterness'),
('COC(=O)C(CCCCN)NC(=O)C(N)CC(O)=O', 'bitterness'),
('CC(Cc1ccccc1)N(C)C(=O)C(N)CC(O)=O', 'bitterness'),
('CC(CC1=Cc2ccccc2N1)NC(=O)C(N)CC(O)=O', 'bitterness'),
('CC(Cc1ccc2c(c1)OCO2)NC(=O)C(N)CC(O)=O', 'bitterness'),
('CC(Cc1ccc(cc1)NS(C)(=O)=O)NC(=O)C(N)CC(O)=O', 'bitterness'),
('CC(C)CC(C)NC(=O)C(N)CC(O)=O', 'bitterness'),
('CCC(C)C(NC(=O)C(N)CC(O)=O)C(=O)OC', 'bitterness'),
('COC(=O)C(CC(C)C)NC(=O)C(N)CC(O)=O', 'bitterness'), ('Nc1cc(ccc1Cl)C(O)=O',
'bitterness'), ('CC(CNC(=O)C(N)CC(O)=O)c1ccccc1', 'bitterness'), ('CCCCC(O)CC',
'bitterness'), ('COc1ccc(cc1O)COC(=O)c1ccccc1', 'bitterness'),
('CC(C)C(C=O)c1ccccc1', 'bitterness'), ('CC1=CCCOC1', 'bitterness'),
('OC1C(CC2(O)CC1OC2=O)OC(=O)C=Cc1ccc(c(c1)O)O', 'bitterness'),
('OC(=O)CCS(=O)(=O)c1ccccc1', 'bitterness'), ('COCC1CCC=C(C)C1', 'bitterness'),
('Oc1ccc(cc1)C=CC(=O)c1cc(ccc1O)O', 'bitterness'), ('CC(O)CCOCCC(C)O',
'bitterness'), ('CCOc1ccc(cc1)NC(N)=S', 'bitterness'), ('Cc1ccc(cc1)F',
'bitterness'), ('COc1ccc2c(c1)C(=O)C=C(O2)c1ccc(cc1)O', 'bitterness'),
('COc1ccc2c(c1)OC(=CC2=O)c1ccc(cc1)O', 'bitterness'), ('COc1ccc(cc1)O',
'bitterness'), ('Oc1ccc(cc1)C(=O)C=Cc1ccccc1', 'bitterness'),
('Oc1ccc(cc1)C1=CC(=O)c2ccccc2O1', 'bitterness'),
('COC1=CC(=O)C(CC=C(C)C)(CC=C(C)C)C(=O)C1=C(O)C=Cc1ccc(cc1)O', 'bitterness'),
('CC(C)c1ccc(cc1)C=NO', 'bitterness'), ('CC1=CCSCC1', 'bitterness'),
('OC1CC2(O)CC(OC(=O)C=Cc3ccc(c(c3)O)O)C1OC2=O', 'bitterness'),
('CC(C1CCC(C(C1)C(O)=O)C12COC(=O)C2C(C)(C)C(O)C=C1)C(O)=O', 'bitterness'),
('CC(C1CCC(C(C1)C(O)=O)C12CCC(O)C(C)(C)C2C(=O)OC1)C(O)=O', 'bitterness'),
('CC(C1CCC(C(C1)C(O)=O)C12CCC(=O)C(C)(C)C2C(=O)OC1)C(O)=O', 'bitterness'),
('COC1CCC(CCC2CCCCC2)CC1O', 'bitterness'), ('COc1ccc(cc1O)CC(CO)c1ccccc1',
'bitterness'), ('Oc1ccccc1C1=CC(=O)c2c(cccc2O1)O', 'bitterness'),
('Oc1ccc(cc1)C1=CC(=O)c2c(cccc2O1)O', 'bitterness'),

('OC(=O)c1cc(ccc1O)Cc1ccc(c(c1)C(O)=O)O', 'bitterness'),
('Oc1cc(c2c(c1)OC(=CC2=O)c1ccccc1O)O', 'bitterness'),
('COc1ccc(cc1)C1=CC(=O)c2c(cc(cc2O1)OC)OC', 'bitterness'),
('COc1cc(c2c(c1)OC(=CC2=O)c1ccccc1)OC', 'bitterness'),
('Oc1cccc2c1C(=O)C=C(O2)c1ccccc1', 'bitterness'),
('OCC1=CC=C(CN2C(=CC=C2C=O)CO)O1', 'bitterness'),
('COc1ccc2c(c1)C(=O)NS2(=O)=O', 'bitterness'), ('CC1CC(=C(N2CCCC2)C1=O)N1CCCC1',
'bitterness'), ('O=C1NS(=O)(=O)c2ccc(cc12)N(=O)=O', 'bitterness'),
('OC1CC(O)(CC(OC(=O)C=Cc2ccc(c(c2)O)O)C1O)C(O)=O', 'bitterness'),
('COc1c(c(c(c(c1C(=O)C=Cc1ccc(cc1)O)O)CC=C(C)C)O)CC=C(C)C', 'bitterness'),
('CCCC1=CNC(=S)NC1=O', 'bitterness'), ('Oc1ccc(cc1)C1=CC(=O)c2cc(ccc2O1)O',
'bitterness'), ('Oc1ccc(cc1)C1=COc2cc(c(cc2C1=O)O)O', 'bitterness'),
('COc1cc2c(cc1OC)C(=O)C=C(O2)c1ccccc1', 'bitterness'),
('OC1C(O)C(CCl)OC(OC2(CCl)OC(CCl)C(O)C2O)C1Cl', 'bitterness'),
('OCC1OC(CO)(OC2OC(CCl)C(O)C(O)C2O)C(O)C1O', 'bitterness'),
('Ic1ccc2c(c1)S(=O)(=O)NC2=O', 'bitterness'),
('COc1c(cc2c(c1O)C(=O)C=C(O2)c1ccc(c(c1)O)O)O', 'bitterness'),
('COc1cc(c2c(c1)CC(C)OC2=O)O', 'bitterness'), ('Cc1ccc2c(c1)C=CC(=O)O2',
'bitterness'), ('O=C1NS(=O)(=O)c2cc(ccc12)N(=O)=O', 'bitterness'),
('CC(=O)OCC1OC(Oc2cc(c3c(c2)OC=C(C3=O)c2ccc(cc2)O)O)C(O)C(O)C1O', 'bitterness'),
('OCC1OC(CO)(OC2OC(COC(=O)c3ccccc3)C(O)C(O)C2O)C(O)C1O', 'bitterness'),
('OCC1OC(CO)(OC2OC(COCc3ccccc3)C(O)C(O)C2O)C(O)C1O', 'bitterness'),
('OCC1OC(CO)(OC2OC(COP(O)(O)=O)C(O)C(O)C2O)C(O)C1O', 'bitterness'),
('Oc1ccc2c(c1)OC(=CC2=O)c1ccc(c(c1)O)O', 'bitterness'),
('Oc1ccc2c(c1)OC=C(C2=O)c1ccc(c(c1)O)O', 'bitterness'),
('Oc1ccc(cc1)C1=CC(=O)c2ccc(cc2O1)O', 'bitterness'),
('COc1ccc(cc1)C1=COc2cc(ccc2C1=O)OC', 'bitterness'),
('Oc1ccc(cc1)C1=COc2c(c(ccc2C1=O)O)O', 'bitterness'),
('Oc1ccc2c(c1)OC=C(C2=O)c1ccccc1', 'bitterness'), ('COc1cccc2c1S(=O)(=O)NC2=O',
'bitterness'), ('CC1CC2=C(NCCC=C2)C1=O', 'bitterness'),
('O=C1NS(=O)(=O)c2c1cccc2N(=O)=O', 'bitterness'),
('CC1=C2C(C3OC(=O)C(=C)C3CC1)C(=CC2=O)COS([O-])(=O)=O', 'bitterness'),
('CC(C)=CCc1c(cc(c2c1OC(CC2=O)c1ccc(cc1)O)O)O', 'bitterness'),
('CCCCCC(O)C=CC(O)C(O)CCCCCCCC(O)=O', 'bitterness'),
('CC1C2CCC(C)(O)C3C4C5C=C(C)C6(C7OC(=O)C(C)C7CCC(C)(O)C56)C4C(=C3C2OC1=O)C',
'bitterness'), ('CCC(Br)(CC)C(=O)NC(=O)NC(C)=O', 'bitterness'),
('COc1ccc(cc1)C(C)=O', 'bitterness'), ('CC(=O)c1ccccc1', 'bitterness'),
('CC(=O)c1cnccn1', 'bitterness'), ('CC(=O)NC(N)=S', 'bitterness'),
('CCN1CC2(COC)C(O)CC(OC)C34C5CC6(O)C(O)C5C(O)(C(O)C6OC)C(C(OC)C23)C14',
'bitterness'), ('OCC1OC2(CC1O)CN1C(=CC=C1C=O)CO2', 'bitterness'),
('Nc1ncnc2c1N=CN2C1OC(CO)C(O)C1O', 'bitterness'),
('CCC(C)C(=O)C1C(=O)C(=O)C(CC=C(C)C)(CC=C(C)C)C1=O', 'bitterness'),
('CCC(C)C(=O)C1=C(O)C(O)(CC=C(C)C)C(O)(C(=O)CC=C(C)C)C1=O', 'bitterness'),
('CCC(C)C(=O)C1=C(O)C(=C(O)C(O)(CC=C(C)C)C1=O)CC=C(C)C', 'bitterness'),
('CCC(C)C(=O)C1=C(O)C(=C(O)C(CC=C(C)C)(CC=C(C)C)C1=O)CC=C(C)C', 'bitterness'),
('CC=C(C)C(=O)OC1CC2C(CC=C3CC(O)CCC23C)C2(O)CCC(C(C)=O)C12C', 'bitterness'),
('Cl.Cl.OC(=O)COCCN1CCN(CC1)C(c1ccccc1)c1ccc(cc1)Cl', 'bitterness'),

```
('C=CCC1(CC=C)C(=O)NC(=O)NC1=O', 'bitterness'),
('OCC1OC(C(O)C(O)C1O)C1c2cccc(c2C(=O)c2c(cc(cc12)CO)O)O', 'bitterness'),
('CC(C)(O)Cc1ccccc1', 'bitterness'),
('NC(CCC(O)=O)C(=O)NC(CC1=CNc2ccccc12)C(O)=O', 'bitterness'),
('NC(CCC(O)=O)C(=O)NC(Cc1ccc(cc1)O)C(O)=O', 'bitterness'),
('CC(OC(C)=O)c1ccccc1', 'bitterness'), ('CC1=CCC(CC1)C(C)(C)OC(=O)c1ccccc1N',
'bitterness'), ('O=C1CCCc2ccccc12', 'bitterness'),
('CCN(CCCc1ccccc1)CCCc1ccccc1.OC(=O)CC(O)(CC(O)=O)C(O)=O', 'bitterness'),
('OCC1OC(OC2OC=C3C(CCOC3=O)C2C=C)C(OC(=O)c2c(cc(cc2-c2cccc(c2)O)O)O)C(O)C1O',
'bitterness'), ('CN1C(=O)N(C)C2=C(NC=N2)C1=O.CN1C(=O)N(C)C2=C(NC=N2)C1=O.NCCN',
'bitterness'), ('[NH4+].[O-]c1c(cc(cc1N(=O)=O)N(=O)=O)N(=O)=O', 'bitterness'),
('CCC1(CCC(C)C)C(=O)NC(=O)NC1=O', 'bitterness'),
('[Na+].CCC1(CCC(C)C)C(=O)NC(=NC1=O)[O-]', 'bitterness'),
('Cl.Cl.CCN(CC)Cc1cc(ccc1O)Nc1ccnc2cc(ccc12)Cl', 'bitterness'),
('CC(N)Cc1ccccc1.OP(O)(O)=O', 'bitterness'),
('CC(N)Cc1ccccc1.CC(N)Cc1ccccc1.OS(O)(=O)=O', 'bitterness'),
('CCN(CC)CC(C)(C)COC(=O)C(CO)c1ccccc1.OP(O)(O)=O', 'bitterness'),
('Cl.CCC(C)(CN(C)C)OC(=O)c1ccccc1', 'bitterness'),
('CC1(CO)C(O)CCC2(C)C(CC=C3C(O)COC3=O)C(=C)CCC12', 'bitterness'),
('CC12CCC(O)CC2CCC2C1CCC1(C)C2CCC1=O', 'bitterness'),
('COc1ccc(cc1)C1=CC(=S)SS1', 'bitterness'),
('Cl.C1CN=C(CN(Cc2ccccc2)c2ccccc2)N1', 'bitterness'),
('OP(O)(O)=O.C1CN=C(CN(Cc2ccccc2)c2ccccc2)N1', 'bitterness'), ('ON=CC1CCCCC1',
'multitaste'), ('CN1N(C(=O)C=C1C)c1ccccc1', 'bitterness'),
('NC(=S)Nc1cccc2ccccc12', 'bitterness'),
('Oc1ccc(cc1)C1=CC(=O)c2c(cc(cc2O1)O)O', 'bitterness'),
('CC(C)C1(CC=C)C(=O)NC(=O)NC1=O', 'bitterness'),
('[Na+].CC(C)C1(CC=C)C(=O)NC(=NC1=O)[O-]', 'bitterness'),
('CC1C2CCC3(C)OC43CC=C(C)C4C2OC1=O', 'bitterness'),
('OCC1OC(Oc2ccc(cc2)O)C(O)C(O)C1O', 'bitterness'), ('Br.COC(=O)C1=CCCN(C)C1',
'bitterness'), ('CC1=CCC23OC3(C)CCC3C(OC(=O)C3=C)C12', 'bitterness'),
('COc1cccc2c1cc(c1c(cc3c(c21)OCO3)C(O)=O)N(=O)=O', 'bitterness'),
('CC1C2CCC(C)(O)C3=CCC(=C3C2OC1=O)C', 'bitterness'),
('CC1C2C3OC(=O)C(=C)C3CCC2(C)C(O)CC1=O', 'bitterness'),
('CC1C2C(O)CC3(C)C=CC(=O)C(=C3C2OC1=O)C', 'bitterness'),
('CC1=CC2OC(=O)C(=C)C2CCC(=C)C(O)CC1', 'bitterness'), ('CC(=O)Oc1ccccc1C(O)=O',
'bitterness'), ('COC(=O)c1c(cc(c(c1O)C)OC(=O)c1c(cc(c(c1O)C=O)O)C)C',
'bitterness'), ('CN1C2CCC1CC(C2)OC(=O)C(CO)c1ccccc1', 'bitterness'), ('O.CN1C2CC
C1CC(C2)OC(=O)C(CO)c1ccccc1.CN1C2CCC1CC(C2)OC(=O)C(CO)c1ccccc1.OS(O)(=O)=O',
'bitterness'), ('CC1OC(OC2C(O)C(OC3OC(CO)C(O)C(O)C3O)C(CO)OC2OC2CCC3(C)C4CCC5(C)
C(CC6OC7(CCC(C)(COC8OC(CO)C(O)C(O)C8O)O7)C(C)C56)C4CC=C3C2)C(O)C(O)C1O',
'bitterness'), ('CC1OC(OC2C(O)C(OC3OC(CO)C(O)C(OC4OC(CO)C(O)C(O)C4O)C3O)C(CO)OC2
OC2CCC3(C)C4CCC5(C)C(CC6OC7(CCC(C)(COC8OC(CO)C(O)C(O)C8O)O7)C(C)C56)C4CC=C3C2)C(
O)C(O)C1O', 'bitterness'), ('OC(=O)c1ccccc1NC(=O)C=Cc1ccc(c(c1)O)O',
'bitterness'), ('COc1cc(ccc1O)C=CC(=O)Nc1ccccc1C(O)=O', 'bitterness'),
('OC(=O)c1ccccc1NC(=O)C=Cc1ccc(cc1)O', 'bitterness'),
('COc1cc(cc(c1O)OC)C=CC(=O)Nc1ccccc1C(O)=O', 'bitterness'),
```

```
('OC(=O)c1cc(ccc1NC(=O)C=Cc1ccc(c(c1)O)O)O', 'bitterness'),
('COc1cc(ccc1O)C=CC(=O)Nc1ccc(cc1C(O)=O)O', 'bitterness'),
('OC(=O)c1cc(ccc1NC(=O)C=Cc1ccc(cc1)O)O', 'bitterness'),
('COc1cc(cc(c1O)OC)C=CC(=O)Nc1ccc(cc1C(O)=O)O', 'bitterness'),
('OCC(O)CC(O)CCCCCCCCCCC=C', 'bitterness'), ('OCC(O)CC(O)CCCCCCCCCCC#C',
'bitterness'), ('Cl.OC(C1CCNCC1)(c1ccccc1)c1ccccc1', 'bitterness'),
('Cn1cnc(c1Sc1ncnc2nc[nH]c12)N(=O)=O', 'bitterness'),
('CCN(CC)C(=O)CSc1ccc(nn1)Cl', 'bitterness'),
('OCC1OC(OC2OC=C3C(CCNC3=O)C2C=C)C(O)C(O)C1O', 'bitterness'),
('CCC1(CC)C(=O)NC(=O)NC1=O', 'bitterness'),
('[Na+].CCC1(CC)C(=O)NC(=NC1=O)[O-]', 'bitterness'), ('O=Cc1ccccc1',
'bitterness'), ('NC(=O)c1ccccc1', 'bitterness'),
('[Cl-].CC(C)(C)CC(C)(C)c1ccc(cc1)OCCOCC[N+](C)(C)Cc1ccccc1', 'bitterness'),
('OC(=O)C(O)(c1ccccc1)c1ccccc1', 'bitterness'), ('OC(c1ccccc1)C(=O)c1ccccc1',
'bitterness'), ('COc1ccc2cc3[n+](cc2c1OC)CCc1cc2c(cc31)OCO2', 'bitterness'),
('[Cl-].COc1ccc2cc3[n+](cc2c1OC)CCc1cc2c(cc31)OCO2', 'bitterness'),
('COc1c2c(cc3c1C=CC(=O)O3)OC=C2', 'bitterness'),
('CC1=C(C=CC(C)=CC=CC(C)=CC=CC=C(C)C=CC=C(C)C=CC2=C(C)CCCC2(C)C)C(C)(C)CCC1',
'bitterness'), ('COc1ccc2c(c1)C13CCN4CC5=CCOC6CC(=O)N2C3C6C5CC14',
'bitterness'), ('COc1ccc(cc1)C1=COc2cc(cc(c2C1=O)O)O', 'bitterness'),
('CC[N+](C)(C)Cc1ccccc1Br.Cc1ccc(cc1)S([O-])(=O)=O', 'bitterness'),
('COc1cc2c(cc1OC)C13CCN4CC5=CCOC6CC(=O)N2C3C6C5CC14', 'bitterness'), ('CC(=O)OC1
CC2(O)C3CCC4CC(CCC4(C)C3CCC2(C)C1C1=COC(=O)C=C1)OC(=O)CCCCCCC(=O)NC(CCCN=C(N)N)C
(O)=O', 'bitterness'), ('Cl.CC(NC(C)(C)C)C(=O)c1cccc(c1)Cl', 'bitterness'),
('[Na+].CCC(C)C1(CC)C(=O)NC(=NC1=O)[O-]', 'bitterness'),
('CC(C)CC1(CC=C)C(=O)NC(=O)NC1=O', 'bitterness'),
('CCC(C)C1(CC(Br)=C)C(=O)NC(=O)NC1=O', 'bitterness'),
('[Na+].CCC(C)C1(CC(Br)=C)C(=O)NC(=NC1=O)[O-]', 'bitterness'),
('CCCC(CC)NC(=O)C(N)CC(O)=O', 'bitterness'),
('Oc1ccc(c(c1)O)C(=O)C=Cc1ccc(c(c1)O)O', 'bitterness'),
('CCCCC1(CC)C(=O)NC(=O)NC1=O', 'bitterness'), ('Cl.CC(C)CNCCOC(=O)c1cccc(c1)N',
'bitterness'), ('CCCCOC(=O)CCC(C)=O', 'bitterness'), ('OC(=O)C=Cc1ccc(c(c1)O)O',
'bitterness'), ('CCOC(=O)C=Cc1ccc(c(c1)O[Si](C)(C)C)O[Si](C)(C)C',
'bitterness'), ('CN1C=NC2=C1C(=O)N(C)C(=O)N2C', 'bitterness'),
('[OH-].[OH-].[Ca+2]', 'bitterness'), ('[Ca+2].CC(O)C([O-])=O.CC([O-])C(O)=O',
'bitterness'),
('CCN(CC)C(=O)c1ccc[n+](c1)C.CC1(C)C2CCC1(C)C(=O)C2S([O-])(=O)=O',
'bitterness'), ('OC1COC2(CC1O)CN1C(=CC=C1C=O)CO2', 'bitterness'),
('O=C1CCCCCN1', 'bitterness'), ('CCOC(=O)N1C=CN(C)C1=S', 'bitterness'),
('CN(C)CCOC(c1ccc(cc1)Cl)c1ccccn1.OC(=O)C=CC(O)=O', 'bitterness'),
('CCCC(C)(COC(N)=O)COC(=O)NC(C)C', 'bitterness'),
('CC(C)c1cc2c(c(c1O)O)C13CCCC(C)(C)C3CC2OC1=O', 'bitterness'),
('CC1C(CC2(C)C(CCC(O)C2(C)O)C1(CC(O)c1ccoc1)C=O)OC(C)=O', 'bitterness'),
('COCC1=C(N2C(SC1)C(NC(=O)C(=NOC)C1=CSC(=N1)N)C2=O)C(=O)OC(C)OC(=O)OC(C)C',
'bitterness'), ('CCC1CN2CCc3cc(c(cc3C2CC1CC1NCCc2cc(c(cc12)OC)O)OC)OC',
'bitterness'), ('CCOCc1c(c(c(c2c1OC(=O)c1c(cc(c(c1O2)C=O)O)C)C)C(O)=O)O',
'bitterness'), ('O=C(C=Cc1ccccc1)c1ccccc1', 'bitterness'),
```

('CC(CCC(O)=O)C1CCC2C3C(O)CC4CC(O)CCC4(C)C3CCC12C', 'bitterness'),
('OC(O)C(Cl)(Cl)Cl', 'bitterness'), ('OCC(NC(=O)C(Cl)Cl)C(O)c1ccc(cc1)N(=O)=O',
'bitterness'), ('Cl.Cl.Cc1ccccc1CN1CCN(CCOC(c2ccccc2)c2ccccc2Cl)CC1',
'bitterness'), ('OCCN(Cc1ccc(cc1Cl)Cl)C(=O)C(Cl)Cl', 'bitterness'),
('CC(CC(C)(C)O)OC(O)C(Cl)(Cl)Cl', 'bitterness'),
('NC(Nc1ccc(cc1)Cl)=NC(N)=NCCCCCCN=C(N)N=C(N)Nc1ccc(cc1)Cl', 'bitterness'),
('COC(CNC(N)=O)C[Hg]Cl', 'bitterness'),
('Cl.Cl.CCN(CC)CC(O)CNc1c2ccc(cc2nc2ccc(cc12)OC)Cl', 'bitterness'),
('[Cl-].CC[NH+](CC)CCOC(=O)c1ccc(cc1Cl)N', 'bitterness'),
('CCN(CC)CCCC(C)Nc1ccnc2cc(ccc12)Cl', 'bitterness'),
('CCN(CC)CCCC(C)Nc1ccnc2cc(ccc12)Cl.OP(O)(O)=O.OP(O)(O)=O', 'bitterness'),
('CN(C)CCC(c1ccc(cc1)Cl)c1ccccn1', 'bitterness'), ('[Cl-].C[N+](C)(C)CCO',
'bitterness'), ('CC(C1CCC2C3CCC4CC(N)CCC4(C)C3CCC12C)N(C)C', 'bitterness'),
('Oc1c2c(c(cc1N(=O)=O)N(=O)=O)C(=O)c1c(c(c(cc1N(=O)=O)N(=O)=O)O)C2=O',
'bitterness'), ('Oc1cc(c2c(c1)OC(=CC2=O)c1ccccc1)O', 'bitterness'),
('COc1cc(ccc1O)C1=CC(=O)c2c(cc(cc2O1)O)O', 'bitterness'),
('OC(C1CC2CCN1CC2C=C)c1ccnc2ccccc12', 'bitterness'),
('OC(=O)c1cc(nc2ccccc12)-c1ccccc1', 'bitterness'),
('CC(C(O)c1ccccc1)N(C)CC=Cc1ccccc1', 'bitterness'), ('OCC=Cc1ccccc1',
'bitterness'), ('OC(=O)C1=CN(C2CC2)c2cc(c(cc2C1=O)F)N1CCNCC1', 'bitterness'),
('CC(C)CCCCCCC=CC(O)=O', 'bitterness'),
('CCC(C)C(=O)C1=C(O)C(O)C(CC=C(C)C)C1=O', 'bitterness'),
('CCC(C)C(=O)C1=C(O)C(O)(C(CC=C(C)C)C1=O)C(=O)C=CC(C)C', 'bitterness'),
('CC(C)C=CC(=O)C1(O)C(CC=C(C)C)C(=O)C(=C1O)C(=O)C(C)C', 'bitterness'),
('CC(C)CC(=O)C1=C(O)C(O)(C(CC=C(C)C)C1=O)C(=O)C=CC(C)C', 'bitterness'),
('CC(C)C(=O)C1=C(O)C(O)C(CC=C(C)C)C1=O', 'bitterness'),
('O=C1CNC(=O)C(Cc2ccccc2)N1', 'bitterness'), ('O=C1CNC(=O)C2CCCN12',
'bitterness'), ('CC1NC(=O)CNC1=O', 'bitterness'), ('CCC(C)C1NC(=O)C(C)NC1=O',
'bitterness'), ('CC(C)CC1NC(=O)C(C)NC1=O', 'bitterness'),
('CC1NC(=O)C(Cc2ccccc2)NC1=O', 'bitterness'), ('CC1NC(=O)C2CCCN2C1=O',
'bitterness'), ('CC1NC(=O)C(Cc2ccc(cc2)O)NC1=O', 'bitterness'),
('CC(C)C1NC(=O)C(C)NC1=O', 'bitterness'), ('NC(=O)CC1NC(=O)C(Cc2ccccc2)NC1=O',
'bitterness'), ('OC(=O)CC1NC(=O)C(Cc2ccccc2)NC1=O', 'bitterness'),
('CCC(C)C1NC(=O)C(Cc2ccccc2)NC1=O', 'bitterness'), ('CCC(C)C1NC(=O)C2CCCN2C1=O',
'bitterness'), ('CC(C)CC1NC(=O)CNC1=O', 'bitterness'),
('CC(C)CC1NC(=O)C(Cc2ccccc2)NC1=O', 'bitterness'), ('CC(C)CC1NC(=O)C2CCCN2C1=O',
'bitterness'), ('O=C1NC(Cc2ccccc2)C(=O)N2CCCC12', 'bitterness'),
('OCC1NC(=O)C(Cc2ccccc2)NC1=O', 'bitterness'), ('CC(O)C1NC(=O)C2CCCN2C1=O',
'bitterness'), ('Oc1ccc(cc1)CC1NC(=O)C2CCCN2C1=O', 'bitterness'),
('CC(C)CC1NC(=O)C(NC1=O)C(C)C', 'bitterness'),
('CC(C)C1NC(=O)C(Cc2ccccc2)NC1=O', 'bitterness'), ('CC(C)C1NC(=O)C2CCCN2C1=O',
'bitterness'), ('CC(C)C1NC(=O)C(Cc2ccc(cc2)O)NC1=O', 'bitterness'),
('CC(C)C1NC(=O)C(NC1=O)C(C)C', 'bitterness'),
('CC(C)CC(=O)C1=C(O)C(CC=C(C)C)C(O)C1=O', 'bitterness'),
('CCC(C)C(=O)C1=C(O)C(O)(C(CC=C(C)C)C1=O)C(=O)CC=C(C)C', 'bitterness'),
('CC(C)C(=O)C1=C(O)C(O)(C(CC=C(C)C)C1=O)C(=O)CC=C(C)C', 'bitterness'),
('CC(C)CC(=O)C1=C(O)C(O)(C(CC=C(C)C)C1=O)C(=O)CC=C(C)C', 'bitterness'),

```
('CCOC(=O)C=Cc1ccc(cc1)O', 'bitterness'), ('COC(=O)C=Cc1ccc(cc1)O',
'bitterness'), ('CCC1OC(=O)C(C)C(OC2CC(C)(OC)C(O)C(C)O2)C(C)C(OC2OC(C)CC(C2O)N(C
)C)C(C)(CC(C)C(=O)C(C)C(O)C1(C)O)OC', 'bitterness'),
('CCCC1CC(N(C)C1)C(=O)NC(C(C)Cl)C1OC(SC)C(O)C(O)C1O', 'bitterness'),
('Cl.CC(CN(C)C)C(C)(O)Cc1ccc(cc1)Cl', 'bitterness'),
('Cc1c(cccc1Nc1ncccc1C(O)=O)Cl', 'bitterness'),
('CC1=CCCC(=CC2OC(=O)C(=C)C2C(C1)OC(=O)C(=C)C(O)CO)CO', 'bitterness'),
('COC(=O)C1C(CC2CCC1N2C)OC(=O)c1ccccc1', 'bitterness'),
('COc1ccc(cc1OC)CC1N(C)CCc2cc(c(cc12)O)OC', 'bitterness'),
('CC(C)C(=O)C1C(=O)C(=O)C(CC=C(C)C)(CC=C(C)C)C1=O', 'bitterness'),
('CC(C)C(=O)C1=C(O)C(O)(CC=C(C)C)C(O)(C(=O)CC=C(C)C)C1=O', 'bitterness'),
('CC(C)C(=O)C1=C(O)C(=C(O)C(O)(CC=C(C)C)C1=O)CC=C(C)C', 'bitterness'),
('COC1=CC=C2C(=CC1=O)C(CCc1cc(c(c(c21)OC)OC)OC)NC(C)=O', 'bitterness'),
('CC12CC(OC(=O)C2CCC2(C)C1C1OC(=O)C2(O)C=C1)C1=COC=C1', 'bitterness'),
('CC(C)C(=O)C1=C(O)C(=C(O)C(CC=C(C)C)(CC=C(C)C)C1=O)CC=C(C)C', 'bitterness'),
('CCCCN1C(=O)c2ccc(cc2S1(=O)=O)N(=O)=O', 'bitterness'),
('CC(C)N1C(=O)c2ccc(cc2S1(=O)=O)N(=O)=O', 'bitterness'),
('CCCCCN1C(=O)c2ccc(cc2S1(=O)=O)N(=O)=O', 'bitterness'),
('ClC(Cl)(Cl)SN1C(=O)c2ccc(cc2S1(=O)=O)N(=O)=O', 'bitterness'),
('C=CCN1C(=O)c2ccc(cc2S1(=O)=O)N(=O)=O', 'bitterness'),
('CN1C(=O)c2ccc(cc2S1(=O)=O)S(N)(=O)=O', 'bitterness'),
('CCN(CC)CCCN1S(=O)(=O)c2ccccc2S1(=O)=O', 'bitterness'),
('CCCCN(CCCC)CCN1S(=O)(=O)c2ccccc2S1(=O)=O', 'bitterness'),
('CC1(C)CC(CC2(C)C1CCC13CC(CCC21)C(=C)C3=O)OOC1OC(CO)C(O)C(O)C1O',
'bitterness'), ('CC1C2CC(O)C3C4(COC(C)=O)C(CCC(C)(C)C4CO)OC(=O)C3(C2)C1O',
'bitterness'),
('CC1C2CC(OC(C)=O)C3C45COC(OC(C)=O)C5C(C)(C)CCC4OC(=O)C3(C2)C1=O',
'bitterness'), ('CC1C2CC(O)C3C45COC=C5C(C)(C)CCC4OC(=O)C3(C2)C1=O',
'bitterness'),
('COC(=O)C(C)C1CC(OC(C)=O)C2C(C1)C(=O)OC1CCC(C)(C)C(COC(C)=O)C21CO',
'bitterness'), ('CC1C2CCC3C45COC(O)(C(O)C5C(C)(C)CCC4=O)C3(C2O)C1=O',
'bitterness'),
('CC1C2CC(OC(C)=O)C3C45COC(O)(C(O)C5C(C)(C)CCC4OC(C)=O)C3(C2)C1O',
'bitterness'), ('CC1C2CCC3C45COC(=O)C5C(C)(C)C(CC4OC(=O)C3(C2)C1=O)OC(C)=O',
'bitterness'), ('CC1C2CCC(C34CCC(=O)C(C)(C)C4C(=O)OC3)C(C2)(C(O)=O)C1=O',
'bitterness'), ('Br.Br.CC1C2CCC3C4CC=C5CC(CCC5(C)C4CCC23CN1C)N(C)C',
'bitterness'),
('CC1OC(OC2CCC3(C=O)C4CCC5(C)C(CCC5(O)C4CCC3(O)C2)C2=CC(=O)OC2)C(O)C(O)C1O',
'bitterness'), ('CC(=C)C1C2CC3(C)C(O)(C4OC4C43CO4)C1C(=O)O2', 'bitterness'),
('CC1=CCCC(=CC2OC(=O)C(=C)C2CC1)C', 'bitterness'), ('OC(=O)C1=Cc2ccccc2O1',
'bitterness'), ('Oc1ccc2c(c1)OC(=O)C1=C2Oc2cc(ccc12)O', 'bitterness'),
('CN(CC(O)=O)C(N)=N', 'bitterness'), ('CN1CC(=O)N=C1N', 'bitterness'),
('CC12CCC(OO)C(=CCC3C(OC(=O)C3=C)C1O)C2', 'bitterness'),
('OC(COc1ccc2c(c1)C(=O)C=C(O2)C(O)=O)COc1cccc2c1C(=O)C=C(O2)C(O)=O',
'bitterness'), ('CC(=O)OC(C)(C)C=CC(=O)C(C)(O)C1C(O)CC2(C)C3CC=C4C(CC(O)C(=O)C4(
C)C)C3(C)C(=O)CC12C', 'bitterness'),
('CC(C)(O)C=CC(=O)C(C)(O)C1C(O)CC2(C)C3CC=C4C(CC(O)C(=O)C4(C)C)C3(C)C(=O)CC12C',
```

'bitterness'), ('CC(=O)OC(C)(C)C=CC(=O)C(C)(O)C1C(O)CC2(C)C3CC=C4C(C=C(O)C(=O)C4(C)C)C3(C)C(=O)CC12C', 'bitterness'), ('CC(C)(O)C=CC(=O)C(C)(O)C1C(O)CC2(C)C3CC=C4C(C=C(O)C(=O)C4(C)C)C3(C)C(=O)CC12C', 'bitterness'), ('Oc1cc(c2cc(c([o+]c2c1)-c1ccc(c(c1)O)O)O)O', 'bitterness'), ('OCC1OC(Oc2cc3c(cc(cc3[o+]c2-c2ccc(c(c2)O)O)O)O)O)C(O)C(O)C1O', 'bitterness'), ('Cl.CNCC(C)C1CCCCC1', 'bitterness'), ('CCC1(C(=O)NC(=O)NC1=O)C1=CCCCC1', 'bitterness'), ('[Ca+2].CCC1(C(=O)NC(=O)N=C1[O-])C1=CCCCC1.CCC1(C(=O)NC(=O)N=C1[O-])C1=CCCCC1', 'bitterness'), ('COCC(C)(OC)C1CCC(=CC1)C', 'bitterness'), ('CC1CC(C)C(=O)C(C1)C(O)CC1CC(=O)NC(=O)C1', 'bitterness'), ('NC1CCCCC1', 'bitterness'), ('CCC(C)C1NC(=O)C(Cc2ccccc2)NC(=O)C(CC(C)C)NC(=O)C2CCCN2C(=O)C(Cc2ccccc2)NC(=O)C(NC(=O)C(CC(C)C)NC(=O)C(CCS(C)=O)NC1=O)C(C)C', 'bitterness'), ('O=C1CCCCCCC1', 'bitterness'), ('Cl.CNC(C)CC1CCCC1', 'bitterness'), ('C=CCC1(C2CCC=C2)C(=O)NC(=O)NC1=O', 'bitterness'), ('Cl.OC(CCN1CCCCC1)(C1CCCC1)c1ccccc1', 'bitterness'), ('OCC(=C)C(=O)OC1CC(=C)C2CC(O)C(=C)C2C2OC(=O)C(=C)C12', 'bitterness'), ('Oc1ccc(cc1)C1=COc2cc(ccc2C1=O)O', 'bitterness'), ('OCC1OC(Oc2ccc3c(c2)OC=C(C3=O)c2ccc(cc2)O)C(O)C(O)C1O', 'bitterness'), ('OCC1OC(OCC2OC(OC(C#N)c3ccccc3)C(O)C(O)C2O)C(O)C(O)C1O', 'bitterness'), ('Nc1ccc(cc1)S(=O)(=O)c1ccc(cc1)N', 'bitterness'), ('Oc1cc(c2c(c1)OC(=C(O)C2=O)c1ccccc1O)O', 'bitterness'), ('CC1(C)C2CCC1(C)C(=O)C2', 'bitterness'), ('CC(CCO)CCC=C(C)C', 'bitterness'), ('CN1C=CC(=O)C(=C1C)O', 'bitterness'), ('CC(CCC(O)=O)C1CCC2C3C(CC(=O)C12C)C1(C)CCC(=O)CC1CC3=O', 'bitterness'), ('OCC(=C)C(=O)OC1CC(=C)C2CC(=O)C(=C)C2C2OC(=O)C(=C)C12', 'bitterness'), ('CCC(C)C(=O)C1=C(O)C2(CC=C(C)C)CC3C(C)(C)C(CC3(C1=O)C2=O)C(C)=C', 'bitterness'), ('CC(C)C(=O)C1=C(O)C2(CC=C(C)C)CC3C(C)(C)C(CC3(C1=O)C2=O)C(C)=C', 'bitterness'), ('CC(C)CC(=O)C1=C(O)C2(CC=C(C)C)CC3C(C)(C)C(CC3(C1=O)C2=O)C(C)=C', 'bitterness'), ('CC[N+](CC)(CC(=O)Nc1c(cccc1C)C)Cc1ccccc1.[O-]C(=O)c1ccccc1', 'bitterness'), ('[Cl-].CC[N+](CC)(CC(=O)Nc1c(cccc1C)C)Cc1ccccc1', 'bitterness'), ('Cc1cccc(c1NC(=O)C[N+](C)(C)CCl)C', 'bitterness'), ('Cc1cccc(c1NC(=O)C[N+](C)(C)CCBr)C', 'bitterness'), ('Cc1cccc(c1NC(=O)C[N+](C)(C)CCCCI)C', 'bitterness'), ('Cc1cccc(c1NC(=O)C[N+](C)(C)CCCCCCl)C', 'bitterness'), ('Cc1cccc(c1NC(=O)C[N+](C)(C)CCCCCCCl)C', 'bitterness'), ('Cc1cccc(c1NC(=O)C[N+](C)(C)CCCCCCCCCl)C', 'bitterness'), ('Cc1cccc(c1NC(=O)C[N+](C)(C)CCCCCCCCCCCl)C', 'bitterness'), ('Cc1cccc(c1NC(=O)C[N+](C)(C)Cc1ccc(cc1)Cl)C', 'bitterness'), ('Cc1cccc(c1NC(=O)C[N+](C)(C)CCc1ccc(cc1)Cl)C', 'bitterness'), ('Cc1cccc(c1NC(=O)C[N+](C)(C)CCCc1ccc(cc1)Cl)C', 'bitterness'), ('CC[N+](C)(CC(=O)Nc1c(cccc1C)C)Cc1ccc(cc1)Cl', 'bitterness'), ('CCCC[N+](C)(CC(=O)Nc1c(cccc1C)C)Cc1ccc(cc1)Cl', 'bitterness'), ('Cc1cccc(c1NC(=O)C[N+](C)(Cc1ccccc1)Cc1ccc(cc1)Cl)C', 'bitterness'), ('CC[N+](CC)(CC(=O)Nc1c(cccc1C)C)Cc1ccc(cc1)Cl', 'bitterness'), ('CC[N+](CC)(CC(=O)Nc1c(cccc1C)C)Cc1ccccc1.[O-]C1=NS(=O)(=O)c2ccccc12', 'bitterness'), ('CC1CC2C3CCC4=CC(=O)C=CC4(C)C3(F)C(O)CC2(C)C1(O)C(=O)CO',

'bitterness'), ('COc1ccc2c(c1)C13CCCCC3C(C2)N(C)CC1', 'bitterness'),
('CC(CN1CCOCC1)C(C(=O)N1CCCC1)(c1ccccc1)c1ccccc1.OC(C(O)C(O)=O)C(O)=O',
'bitterness'), ('CC1(C)C2CCC(C)(C2)C1=O', 'bitterness'),
('CN1C(=O)CN=C(c2ccccc2)c2cc(ccc12)Cl', 'bitterness'),
('NCC1CCC(N)C(O1)OC1C(N)CC(N)C(OC2OC(CO)C(O)C(N)C2O)C1O.OS(O)(=O)=O',
'bitterness'), ('OC(=O)Cc1ccccc1Nc1c(cccc1Cl)Cl', 'bitterness'),
('OC1=C(CC2=C(O)c3ccccc3OC2=O)C(=O)Oc2ccccc12', 'bitterness'),
('Cl.CCN(CC)CCOC(=O)C1(CCCC1)C1CCCCC1', 'bitterness'),
('CCOC(=O)c1ccccc1C(=O)OCC', 'bitterness'), ('CCC(Br)(CC)C(N)=O', 'bitterness'),
('CC1OC(CC(O)C1O)OC1C(O)CC(OC1C)OC1C(O)CC(OC1C)OC1CCC2(C)C(CCC3C2CCC2(C)C(CCC32O
)C2=CC(=O)OC2)C1', 'bitterness'), ('CC1C2CCC3(C)C(O)CCC(=C3C2OC1=O)C',
'bitterness'), ('CC1C2CC(O)C3C45COC(O)C5C(C)(C)CCC4OC(=O)C3(C2)C1=O',
'bitterness'), ('CC1C2CC(O)C3C4(COC(C)=O)C(CCC(C)(C)C4C=O)OC(=O)C3(C2)C1=O',
'tastelessness'),
('CC1C2CCC3C45COC(OC(C)=C)(C(OC(C)=O)C5C(C)(C)CCC4OC(C)=O)C3(C2OC(C)=O)C1=O',
'bitterness'), ('CC(C)CNCC(C)C', 'bitterness'),
('Cl.COc1ccc(cc1)C1Sc2ccccc2N(CCN(C)C)C(=O)C1OC(C)=O', 'bitterness'),
('CN1CCC23CCCCC3C1Cc1ccc(cc21)C.OP(O)(O)=O', 'bitterness'),
('Cl.CCN(CC)CC(C)(C)COC(=O)c1ccc(cc1)N', 'bitterness'), ('CS(C)=O',
'bitterness'), ('CN(C)C=S', 'bitterness'),
('CC12CC(OC(=O)C2CC(=O)C2C3CC(CC12)OC3=O)C1=COC=C1', 'bitterness'), ('CC1CCC2(OC
1)OC1CC3C4CC=C5CC(CCC5(C)C4CCC3(C)C1C2C)OC1OC(CO)C(OC2OC(C)C(O)C(O)C2O)C(O)C1OC1
OC(C)C(O)C(O)C1O', 'bitterness'), ('CN(C)C(=O)C(=O)N(N(C)C(C)=O)c1ccccc1',
'bitterness'), ('Cl.O=C(Nc1ccccc1)OCC(CN1CCCCC1)OC(=O)Nc1ccccc1', 'bitterness'),
('CN(C)CCOC(c1ccccc1)c1ccccc1', 'bitterness'),
('OC(CCCN1CCCCC1)(c1ccccc1)c1ccccc1', 'bitterness'),
('OCCN(CCO)c1nc(c2nc(nc(c2n1)N1CCCCC1)N(CCO)CCO)N1CCCCC1', 'bitterness'),
('CC1C2CC(O)C3C4(COC(C)=O)C(CCC(C)(C)C4C(O)=O)OC(=O)C3(C2)C1=O', 'bitterness'),
('COC(=O)C1C(C)(C)CCC2OC(=O)C34CC(CC(O)C3C12COC(C)=O)C(C)C4=O', 'bitterness'),
('OC(C=C)C(O)C=C', 'bitterness'), ('C=CS(=O)C=C', 'bitterness'),
('CC(C)CC(N)C(=O)NC(CC(C)C)C(O)=O', 'bitterness'),
('CCC=CCC=CCC=CCC=CCC=CCC=CCCC(O)=O', 'bitterness'),
('[Cl-].CCCCCCCCCCCCNC(=O)C[N+](C)(C)Cc1ccccc1', 'bitterness'),
('[Br-].CCCCCCCCCCCC[N+](C)(C)CCOc1ccccc1', 'bitterness'),
('Cl.COc1cc2c(cc1OC)C(=O)C(CC1CCN(CC1)Cc1ccccc1)C2', 'bitterness'),
('CN(C)CCC=C1c2ccccc2COc2ccccc12', 'bitterness'),
('COc1ccc2c(c1OC)C13CCN(C)C(C2)C3(O)CCC(O)C1', 'bitterness'),
('CN1C(=O)N(C)C2=C(N(CC(O)CO)C=N2)C1=O', 'bitterness'),
('[Br-].CC[N+](C)(C)c1cccc(c1)O', 'bitterness'),
('O=C(OC1CCN2CC3CC(CN4C3CCCC4=O)C2C1)C=Cc1ccccc1', 'bitterness'),
('CCC1CN2CCc3cc(c(cc3C2CC1CC1NCCc2cc(c(cc12)OC)OC)OC)OC', 'bitterness'),
('CCOC(=O)C(CCc1ccccc1)NC(C)C(=O)N1CCCC1C(O)=O', 'bitterness'),
('CC1(C)C(O)CC2OC(=O)C34CC(CCC3C32COC(O)C13)C(=C)C4=O', 'bitterness'),
('CC(=O)OC1CC2OC(=O)C34CC(CCC3C32COC(O)C3C1(C)C)C(=C)C4=O', 'bitterness'),
('CCN1C=C(C(O)=O)C(=O)c2cc(c(nc12)N1CCNCC1)F', 'bitterness'),
('Oc1cc(c2c(c1)OC(C(C2)OC(=O)c1cc(c(c(c1)O)O)O)c1cc(c(c(c1)O)O)O)O',
'bitterness'), ('COc1cc(nc(n1)N1N=C(C)C=C1OC)C', 'bitterness'),

('OC1CC2C(C3OC(=O)C(=C)C3CCC2=C)C1=C', 'bitterness'),
('Cl.Cl.CCOC(CN1CCN(CC1)CC(C)C(=O)c1ccccc1)c1ccccc1', 'bitterness'),
('Oc1cc(c2c(c1)OC(CC2=O)c1ccc(c(c1)O)O)O', 'bitterness'),
('Oc1cc(c(c(c1)O)C(=O)C=Cc1ccc(c(c1)O)O)O', 'bitterness'), ('CCC1OC(=O)C(C)C(OC2
CC(C)(OC)C(O)C(C)O2)C(C)C(OC2OC(C)CC(C2O)N(C)C)C(C)(O)CC(C)C(=O)C(C)C(O)C1(C)O',
'bitterness'), ('OCC1OC(Oc2cc3c(cc2O)OC(=O)C=C3)C(O)C(O)C1O', 'bitterness'),
('CCOC(=O)C(Cc1ccc(c(c1)O)OC)c1ccccc1', 'bitterness'), ('CCOC(=O)c1ccccc1',
'bitterness'), ('CCOC(=O)CC(=O)c1ccccc1', 'bitterness'),
('CCOC(=O)C(C1=C(O)c2ccccc2OC1=O)C1=C(O)c2ccccc2OC1=O', 'bitterness'),
('CCOC=O', 'bitterness'), ('CCC1=C(O)C(=O)C=CO1', 'bitterness'),
('CCOC(=O)C=Cc1ccc(c(c1)O)O', 'bitterness'),
('Cl.CCN(CC)CCOC(c1ccccc1)c1ccccc1', 'bitterness'), ('S=C1NCCN1', 'bitterness'),
('CCOc1ccc2nccc(c2c1)C(O)C1CC2CCN1CC2CC', 'bitterness'), ('CCc1cnccn1',
'bitterness'), ('Cl.CCNCC(O)c1cccc(c1)O', 'bitterness'),
('COc1cc(c2c(c1)OC(=CC2=O)C)O', 'bitterness'), ('COc1ccc(cc1OC)CC=C',
'bitterness'), ('CCCCCCCC=CC(O)C#CC#CC(O)C=C', 'bitterness'),
('CCCCCCCC=CC(O)C#CC#CC(OC(C)=O)C=C', 'bitterness'),
('CCCCCCCC=CCC#CC#CC(O)C=C', 'bitterness'),
('NC(N)=Nc1nc(cs1)CSCCC(N)=NS(N)(=O)=O', 'bitterness'),
('Cl.CC(Cc1ccccc1)N(C)CCNC1=NC2=C(N1C)C(=O)N(C)C(=O)N2C', 'bitterness'),
('CC1(C)C2CCC(C)(C2)C1O', 'bitterness'),
('COc1ccc(cc1)CC=C.COc1ccc(cc1)C=CC.CC1(C)C2CCC(C)(C2)C1=O', 'bitterness'),
('Cl.CC(Cc1ccccc1)NCCC#N', 'bitterness'),
('CCC(=O)N(C1CCN(CC1)CCc1ccccc1)c1ccccc1.OC(=O)CC(O)(CC(O)=O)C(O)=O',
'bitterness'), ('[Fe+2].[O-]S([O-])(=O)=O.c1cnc2c(c1)ccc1cccnc21.c1cnc2c(c1)ccc1
cccnc21.c1cnc2c(c1)ccc1cccnc21', 'bitterness'), ('CCOC(=O)C=Cc1ccc(c(c1)OC)O',
'bitterness'), ('Oc1ccc2c(c1)OC(=C(O)C2=O)c1ccc(c(c1)O)O', 'bitterness'),
('O=C1CC(Oc2ccccc12)c1ccccc1', 'bitterness'), ('O=C1C=C(Oc2ccccc12)c1ccccc1',
'bitterness'), ('OC(=O)c1ccccc1Nc1cccc(c1)C(F)(F)F', 'bitterness'),
('COc1ccc(cc1)C1=COc2cc(ccc2C1=O)O', 'bitterness'),
('COc1cc2c(c(c1O)OC1OC(CO)C(O)C(O)C1O)OC(=O)C=C2', 'bitterness'),
('OCC1=CC=CO1', 'bitterness'), ('CC(CCC1(O)OC2CC3C4CCC5CC(CCC5(C)C4CCC3(C)C2C1C)
OC1(C)OC(C)(CO)C(C)(COCC2(C)OC(C)(CO)C(C)(O)C(C)(O)C2(C)OC2(C)OC(C)(CO)C(C)(O)C(
C)(O)C2(C)O)C(C)(O)C1(C)O)COCC1(C)OC(C)(CO)C(C)(O)C(C)(O)C1(C)O', 'bitterness'),
('OC1C(Oc2cc(ccc2C1=O)O)c1ccc(c(c1)O)O', 'bitterness'), ('CC(C)=CCN=C(N)N',
'bitterness'), ('CCOC(=O)c1cc(c(c(c1)O)O)O', 'bitterness'),
('CC12CCC(O)CC2CCC2C1C(O)CC1(C)C(CCC21O)C1=COC(=O)C=C1', 'bitterness'),
('c1cc(ccn1)-c1ccncc1', 'bitterness'),
('NC(CCC(=O)NC(Cc1ccc(cc1)O)C(O)=O)C(O)=O', 'bitterness'),
('CC(CC(=O)CC(C)C(O)=O)C1CC(O)C2(C)C3=C(C(=O)CC12C)C1(C)CCC(=O)C(C)(C)C1CC3O',
'bitterness'), ('CC12CCCC3(CN4CCOC14)C2CCC12CC(CCC31)C(=C)C2O', 'bitterness'),
('COc1c(c(cc2c1N(C=C(C(O)=O)C2=O)C1CC1)F)N1CCNC(C)C1', 'bitterness'),
('CC(CCC1C(C)(O)CC(OC2OCC(O)C(O)C2O)C2C(C)(CO)CCCC12C)=CCO', 'bitterness'),
('COc1cc(c(cc1OC)C=O)OC', 'bitterness'), ('Oc1ccc(cc1)C1=COc2cc(cc(c2C1=O)O)O',
'bitterness'), ('COc1cc(c2c(c1)OC(=CC2=O)c1ccc(cc1)O)O', 'bitterness'),
('C=Cc1cncc2c1CCOC2=O', 'bitterness'),
('OCC1OC(OCC2OC(O)C(O)C(O)C2O)C(O)C(O)C1O', 'non-sweetness'),

('OCC1OC(OC2OC=C3C(=O)OCC=C3C2C=C)C(O)C(O)C1O', 'bitterness'),
('CC(=O)OCC=C(C)CCC=C(C)C', 'bitterness'), ('CC(C)=CCCC(C)=CCOC=O',
'bitterness'), ('CCC(=O)OCC=C(C)CCC=C(C)C', 'bitterness'),
('CC1=CC(O)C2C(CC(=CCC1)C)OC(=O)C2(C)O', 'bitterness'),
('CC1CCCC2(C)CCC3C(C)C(=O)OC3C12', 'bitterness'),
('OCC1OC(Oc2cccc3cc4c(c(c23)O)C(=O)OC4)C(O)C(O)C1O', 'bitterness'),
('CC1C(=O)OC2CC34C5CC(C(C)(C)C)C64C(O)C(=O)OC6OC3(C(=O)O5)C12O', 'bitterness'),
('CC1C(=O)OC2C(O)C34C5CC(C(C)(C)C)C64C(O)C(=O)OC6OC3(C(=O)O5)C12O',
'bitterness'),
('CC1C(=O)OC2C(O)C34C5OC(=O)C4(OC4OC(=O)C(O)C34C(C5O)C(C)(C)C)C12O',
'bitterness'),
('CCC(C)(O)C(=O)OC1C2C(C)C(O)C3(O)OCC42C(CC2C(=CC(O)C(O)C2(C)C34)C)OC1=O',
'bitterness'),
('CCC1=C(C)CN(C(=O)NCCc2ccc(cc2)S(=O)(=O)NC(=O)NC2CCC(C)CC2)C1=O',
'bitterness'), ('CC(C)CC(NC(=O)C(N)CCC(N)=O)C(=O)NC(Cc1ccccc1)C(=O)NC(CC(N)=O)C(
=O)N1CCCC1C(=O)NC(CO)C(=O)NC(C(C)O)C(=O)NC(CC(N)=O)C(=O)N1CCCC1C(=O)NC(CC1=CNc2c
cccc12)C(=O)NC(CC1=CN=CN1)C(=O)NC(CO)C(=O)N1CCCC1C(O)=O', 'bitterness'),
('OCC1OC(SC(CC2=CNc3ccccc23)=NOS(O)(=O)=O)C(O)C(O)C1O', 'bitterness'),
('CC(=O)OCC(OC(C)=O)C(OC(C)=O)C(OC(C)=O)C(OC(C)=O)C=O', 'bitterness'),
('COc1cc(ccc1OC1OC(CO)C(O)C(O)C1O)C=O', 'bitterness'),
('CCCCCCCCCC(=O)OCC(O)CO', 'bitterness'), ('CCCCCC=CCC=CCCCCCCCC(=O)OCC(O)CO',
'bitterness'), ('CCCCCCCCCCCCCCCCCC(=O)OCC(O)CO', 'bitterness'),
('CCCCCCCCCCCCCCCC(=O)OCC(O)CO', 'bitterness'),
('CCC(=O)OCC(COC(=O)CC)OC(=O)CC', 'bitterness'),
('COc1cc2c(cc1O)OC=C(C2=O)c1ccc(cc1)O', 'bitterness'),
('COc1cc2c(cc1OC1OC(CO)C(O)C(O)C1O)OC=C(C2=O)c1ccc(cc1)O', 'bitterness'),
('C=CC1CNC(=S)O1', 'bitterness'),
('Oc1ccc(cc1O)C1=C(O)C(=O)c2c(cc(c(c2O1)O)O)O', 'bitterness'),
('CC1C2C(CC1=O)C(=C)CC(O)C1C2OC(=O)C1=C', 'bitterness'), ('COc1ccccc1OCC(O)CO',
'bitterness'), ('OC1(CCN(CCCC(=O)c2ccc(cc2)F)CC1)c1ccc(cc1)Cl', 'bitterness'),
('Cc1nccc2c1Nc1ccccc21', 'bitterness'),
('CC1CC2OC(=O)C(=C)C2C(O)C2(C)C1C=CC2=O', 'bitterness'),
('OCC1OC(Oc2ccccc2C=O)C(O)C(O)C1O', 'bitterness'),
('CCC1(C(=O)NC(=O)NC1=O)C1=CCCCCC1', 'bitterness'),
('Oc1ccc(cc1)C1=C(O)C(=O)c2c(cc(c(c2O1)O)O)O', 'bitterness'),
('CC1C2CC(OC(C)=O)C(=CCCC(=CC2OC1=O)C)C', 'bitterness'),
('CC1C2CC(OC(C)=O)C(=C)C(CCC(=CC2OC1=O)C)OC(C)=O', 'bitterness'),
('[Na+].CCCCCCC1(CC)C(=O)NC(=NC1=O)[O-]', 'bitterness'),
('[Na+].CN1C(=O)[N-]C(=O)C(C)(C1=O)C1=CCCCC1', 'bitterness'), ('COC1CC(OC(C)C1OC
1CC(OC)C(OC2OC(C)C(O)C(OC)C2O)C(C)O1)OC1CCC2(C)C3CC(OC(=O)C(C)=CC)C4(C)C(CCC4(O)
C3CC=C2C1)C(C)=O', 'bitterness'),
('[Br-].CC[n+]1c2cc(ccc2c2ccc(cc2c1-c1ccccc1)N)N', 'bitterness'),
('CC(C)C1CC(=CC(=O)C1)C', 'bitterness'),
('COc1cc(ccc1O)C1CC(=O)c2c(cc(cc2O1)O)O', 'bitterness'),
('OCC1C(=Cc2ccco2)C(=O)c2c(c(c(c[n+]12)[O-])Cc1ccco1)CO', 'bitterness'),
('CCCC(=O)CC(=O)NC1CCOC1=O', 'bitterness'), ('CCCCCC(=O)CC(=O)NC1CCOC1=O',
'bitterness'), ('CCCC(=O)NC1CCOC1=O', 'bitterness'), ('CCCCCC(=O)NC1CCOC1=O',

'bitterness'), ('CC(C)=CCC1(CC=C(C)C)C(=O)C(=C(O)C1=O)O', 'bitterness'),
('CC(C)CC(=O)C1=C(O)C(=O)C(CC=C(C)C)(CC=C(C)C)C1=O', 'bitterness'),
('CC(C)CC(=O)C1=C(O)C(O)(CC=C(C)C)C(O)(C(=O)CC=C(C)C)C1=O', 'bitterness'),
('CC(C)CC(=O)C1=C(O)C(=C(O)C(O)(CC=C(C)C)C1=O)CC=C(C)C', 'bitterness'),
('CC12CCC(=O)C=C2CCC2C3CCC(O)(C(=O)CO)C3(C)CC(O)C12', 'bitterness'), ('OO',
'bitterness'), ('CCC(C)C(=O)C1=C(O)C(O)(C(CC=C(C)C)C1=O)C(=O)C=CC(C)(C)OO',
'bitterness'), ('CC(C)C(=O)C1=C(O)C(O)(C(CC=C(C)C)C1=O)C(=O)C=CC(C)(C)OO',
'bitterness'), ('CC(C)CC(=O)C1=C(O)C(O)(C(CC=C(C)C)C1=O)C(=O)C=CC(C)(C)OO',
'bitterness'),
('CCC(C)C(=O)C1=C(O)C23CC(C(C)(C)OO)C(C)(C)C3CC(CC=C(C)C)(C1=O)C2=O',
'bitterness'),
('CC(C)C(=O)C1=C(O)C23CC(C(C)(C)OO)C(C)(C)C3CC(CC=C(C)C)(C1=O)C2=O',
'bitterness'),
('CC(C)CC(=O)C1=C(O)C23CC(C(C)(C)OO)C(C)(C)C3CC(CC=C(C)C)(C1=O)C2=O',
'bitterness'), ('CCN(CCO)CCCC(C)Nc1ccnc2cc(ccc12)Cl.OS(O)(=O)=O', 'bitterness'),
('CCC(C)C(=O)C1=C(O)C(O)(C(CC=C(C)C)C1=O)C(=O)C=CC(C)(C)O', 'bitterness'),
('CC(C)C(=O)C1=C(O)C(O)(C(CC=C(C)C)C1=O)C(=O)C=CC(C)(C)O', 'bitterness'),
('CC(C)CC(=O)C1=C(O)C(O)(C(CC=C(C)C)C1=O)C(=O)C=CC(C)(C)O', 'bitterness'),
('OC1CCN2CC3CC(CN4C3CCCC4=O)C2C1', 'bitterness'),
('CCC(C)C(=O)C1=C(O)C2(CC=C(C)C)CC3C(C)(C)C(CC3(C1=O)C2=O)C(C)(C)O',
'bitterness'),
('CC(C)C(=O)C1=C(O)C2(CC=C(C)C)CC3C(C)(C)C(CC3(C1=O)C2=O)C(C)(C)O',
'bitterness'),
('CC(C)CC(=O)C1=C(O)C2(CC=C(C)C)CC3C(C)(C)C(CC3(C1=O)C2=O)C(C)(C)O',
'bitterness'), ('Cl.Cl.OCCOCCN1CCN(CC1)C(c1ccccc1)c1ccc(cc1)Cl', 'bitterness'),
('O=C1N=CNC2=C1NC=N2', 'bitterness'), ('N1C=CN=C1', 'bitterness'),
('Cl.CN(C)CCC[N+]1([O-])c2ccccc2CCc2ccccc12', 'bitterness'),
('Cc1ccc(cc1)S(O)(=O)=O.CS(=O)(=O)OCCCNCCCOS(C)(=O)=O', 'bitterness'),
('Cl.CCc1ccc(cc1)C(=O)C(C)CN1CCCC1', 'bitterness'),
('OCC1OC(C(O)C1O)N1C=NC2=C1NC=NC2=O', 'bitterness'), ('CC(I)C1OCC(CO)O1',
'bitterness'), ('[Na+].CN(C)C=Nc1c(cc(c(c1I)CCC([O-])=O)I)I', 'bitterness'),
('CC(C)COC(C)=O', 'bitterness'), ('CCC(=O)OCC(C)C', 'bitterness'),
('CC(C)COC(=O)c1ccccc1O', 'bitterness'), ('CCOS(=O)(=O)NCC(C)C', 'bitterness'),
('CC(=O)OCC12C(CCC(C)(C)C1C=O)OC(=O)C13CC(CC(O)C21)C(=C)C3=O', 'tastelessness'),
('COc1ccc(cc1OC)C=CC', 'bitterness'), ('O=C1C(=COc2ccccc12)c1ccccc1',
'bitterness'), ('Oc1ccc(cc1)C=CC(=O)c1ccc(cc1O)O', 'bitterness'),
('OCC1CCCC2CCCC[N+]12[O-]', 'bitterness'),
('CNC(C)CCC=C(C)C.OC(C(O)C(O)C(O)=O)C(O)C(O)=O', 'bitterness'),
('COc1c2c(c(c3c1OC(=O)C=C3)OC)C=CO2', 'bitterness'), ('CC(C)O', 'bitterness'),
('COc1cc(ccc1O)C1=C(O)C(=O)c2c(cc(cc2O1)O)O', 'bitterness'),
('COc1cc(c(c2c1C(=O)CC(O2)c1ccc(cc1)O)CC=C(C)C)O', 'bitterness'),
('COc1cc(c(c2c1C(=O)CC(O2)c1ccc(cc1)O)CCC(C)(C)O)O', 'bitterness'),
('COc1cc(c(c2c1C(=O)CC(O2)c1ccc(cc1)O)CCC(C)(C)OC)O', 'bitterness'),
('CCOC(C)(C)CCc1c(cc(c2c1OC(CC2=O)c1ccc(cc1)O)OC)O', 'bitterness'),
('CC(C)=CCC1=C(O)C(CC=C(C)C)(CC=C(C)C)C(=O)C2=C1OC(CC2=O)c1ccc(cc1)O',
'bitterness'), ('Cl.CC(COc1ccccc1)NC(C)C(O)c1ccc(cc1)O', 'bitterness'),
('Oc1ccc(cc1)C1=C(O)C(=O)c2c(cc(cc2O1)O)O', 'bitterness'),

('OC1C(O)C(COC(=O)CC(O)=O)OC(OC2=C(Oc3cc(cc(c3C2=O)O)O)c2ccc(cc2)O)C1O',
'bitterness'), ('OCC1OC(OC2=C(Oc3cc(cc(c3C2=O)O)O)c2ccc(cc2)O)C(O)C(O)C1O',
'bitterness'), ('COc1c2c(c(c3c1OC(=CC3=O)C)OC)C=CO2', 'bitterness'),
('Cl.CC(CCc1ccccc1)NCC(O)c1ccc(c(c1)C(N)=O)O', 'bitterness'),
('CC1=C2C(C3OC(=O)C(=C)C3C(O)C1)C(=CC2=O)CO', 'bitterness'),
('CC1=C2C(C3OC(=O)C(=C)C3C(O)C1)C(=CC2=O)COC(=O)C(O)=O', 'bitterness'),
('CC1=C2C(C3OC(=O)C(=C)C3C(O)C1)C(=CC2=O)COC(=O)Cc1ccc(cc1)O', 'bitterness'),
('CC1=C2C(C3OC(=O)C(=C)C3C(C1)OC(=O)Cc1ccc(cc1)O)C(=CC2=O)COC(=O)C(O)=O',
'bitterness'), ('CC1=C2C(C3OC(=O)C(=C)C3C(C1)OC(=O)Cc1ccc(cc1)O)C(=CC2=O)CO',
'bitterness'), ('O=C(CS(=O)CC1=CC=CO1)NCC=CCOc1cc(ccn1)CN1CCCCC1',
'bitterness'), ('CC(C)CC(NC(=O)C(C)N)C(O)=O', 'bitterness'),
('CC(C)C(NC(=O)C(C)N)C(O)=O', 'bitterness'),
('CCN1CC2(CCC(OC)C34C2CC(C13)C1(O)CC(OC)C2CC4C1(O)C2OC)OC(=O)c1ccccc1NC(C)=O',
'bitterness'), ('NC(CCCNC(N)=N)C(O)=O', 'miscellaneous'),
('CC(C)CC(N)C(=O)NC(CC(C)C)C(=O)NC(CC(C)C)C(O)=O', 'bitterness'),
('CC(C)CC(N)C(=O)NC(Cc1ccccc1)C(O)=O', 'bitterness'),
('CC(C)CC(N)C(=O)NC(CC1=CNc2ccccc12)C(O)=O', 'bitterness'),
('CC(C)CC(NC(=O)CN)C(O)=O', 'bitterness'), ('NCC(=O)NC(Cc1ccccc1)C(O)=O',
'bitterness'), ('CCN(CC)CC(=O)Nc1c(cccc1C)C', 'bitterness'),
('Cl.CCN(CC)CC(=O)Nc1c(cccc1C)C', 'bitterness'),
('CC1(C)OC2CC(=O)OCC32C2CCC4(C)C(OC(=O)C5OC45C2(C)C(=O)CC13)C1=COC=C1',
'bitterness'), ('CC(C)(OC1OC(CO)C(O)C(O)C1O)C#N', 'bitterness'),
('Cl.CCCC1CC(N(C)C1)C(=O)NC(C(C)O)C1OC(SC)C(O)C(O)C1O', 'bitterness'),
('Oc1ccc(cc1)C1CC(=O)c2ccc(cc2O1)O', 'bitterness'), ('[Li+].[Br-]',
'bitterness'), ('CC(C)CC(N)C(=O)NC(C)C(O)=O', 'bitterness'),
('CC(C)CC(N)C(=O)NCC(O)=O', 'bitterness'), ('CC(C)C1CCC(C)CC1=O', 'bitterness'),
('CCCCC(N)C(O)=O', 'multitaste'),
('CCN1C=C(C(O)=O)C(=O)c2cc(c(c(c12)F)N1CCNC(C)C1)F', 'bitterness'),
('NC(Cc1ccccc1)C(=O)NCC(O)=O', 'bitterness'),
('NC(Cc1ccccc1)C(=O)NCC(=O)NCC(=O)NC(Cc1ccccc1)C(O)=O', 'bitterness'),
('NC(Cc1ccccc1)C(=O)NCC(=O)NC(Cc1ccccc1)C(=O)NCC(O)=O', 'bitterness'),
('OCC1CCCN2CCCCC12', 'bitterness'),
('CC(C)CC(=O)C1=C(O)C(=C(O)C(CC=C(C)C)(CC=C(C)C)C1=O)CC=C(C)C', 'bitterness'),
('Oc1cc(c2c(c1)OC(=CC2=O)c1ccc(c(c1)O)O)O', 'bitterness'),
('CC(C)C(N)C(=O)NC(C)C(O)=O', 'bitterness'),
('CCN1CC2(CO)CCC(OC)C34C5CC6C(CC(O)(C5C6OC)C(O)(C(OC)C23)C14)OC', 'bitterness'),
('CC1CC2CC(=O)C3CCCN4CCCC2C34C1', 'bitterness'), ('[Mg+2].[Br-].[Br-]',
'bitterness'), ('[Mg+2].[O-][Cl](=O)=O.[O-][Cl](=O)=O', 'bitterness'),
('[Mg+2].[Cl-].[Cl-]', 'bitterness'), ('[Mg+2].CC(O)C([O-])=O.CC(O)C([O-])=O',
'bitterness'), ('[Mg+2].[O-]S([O-])(=O)=O', 'bitterness'),
('CCOC(=O)CC(SP(=S)(OC)OC)C(=O)OCC', 'bitterness'),
('COc1cc(cc(c1O)OC)-c1[o+]c2cc(cc(c2cc1OC1OC(CO)C(O)C(O)C1O)O)O', 'bitterness'),
('CC1CC2OC(=O)C3(C)CCCC(C)(C23)C1(O)CCC1=COC=C1', 'bitterness'),
('O=C1CCCC2C3CCCN4CCCC(CN12)C34', 'bitterness'),
('[Na+].[Na+].[O-]S(=O)(=O)Nc1cccc(c1)NS([O-])(=O)=O', 'bitterness'),
('COc1ccc2c(c1OC)C(=O)OC2', 'bitterness'), ('Cc1cccc(c1C)Nc1ccccc1C(O)=O',
'bitterness'), ('OCC1OC(Oc2ccccc2C=CC(O)=O)C(O)C(O)C1O', 'bitterness'),

('CC(C)CC(=O)OC1CC(C)CCC1C(C)C', 'bitterness'),
('Cl.CCOC(=O)C1(CCN(C)CC1)c1ccccc1', 'bitterness'), ('Cc1ccccc1OCC(O)CO',
'bitterness'), ('CCCC(C)(COC(N)=O)COC(N)=O', 'bitterness'),
('COc1ccc2c(c1CC1OC1(C)C)OC(=O)C=C2', 'bitterness'),
('O.[Na+].COC(C[Hg])CNC(=O)c1ccccc1OCC([O-])=O', 'bitterness'),
('Cc1cccc(c1)NC(N)=O', 'bitterness'),
('Cl.CN(C)C1C2C(O)C3C(=C)c4cccc(c4C(=C3C(=O)C2(O)C(=C(C(N)=O)C1=O)O)O)O',
'bitterness'), ('CCC(=O)C(CC(C)N(C)C)(c1ccccc1)c1ccccc1', 'bitterness'),
('Cl.CCC(=O)C(CC(C)N(C)C)(c1ccccc1)c1ccccc1', 'bitterness'),
('Cl.CNC(C)Cc1ccccc1', 'bitterness'),
('[Br-].CC[N+](C)(CC)CCOC(=O)C1c2ccccc2Oc2ccccc12', 'bitterness'),
('CN1C=CNC1=S', 'bitterness'), ('COc1c2c(cc3c1OC(=O)C=C3)C=CO2', 'bitterness'),
('Cl.CCC(NC)c1ccccc1OC', 'bitterness'), ('COC(=O)c1ccccc1CCc1ccc(c(c1)O)OC',
'bitterness'), ('COC(C)=O', 'bitterness'), ('COC(=O)c1ccccc1N', 'bitterness'),
('[Cl-].Cc1cc(ccc1C(C)(C)CC(C)(C)C)OCCOCC[N+](C)(C)Cc1ccccc1', 'bitterness'),
('CCC(CO)NC(=O)C1CN(C)C2CC3=CNc4cccc(c34)C2=C1.OC(=O)C=CC(O)=O', 'bitterness'),
('CC1=CC(=O)NC(=S)N1', 'bitterness'), ('COC1CC(O)C(O)C(C)O1', 'bitterness'),
('COC1OC(C)C(O)CC1O', 'bitterness'), ('COC1OC(C)CC(O)C1O', 'bitterness'),
('CCC1(CC)C(=O)NCC(C)C1=O', 'bitterness'),
('Cl.COc1ccc2c(c1)C1=C3N2CCN(C)C3=NCC1', 'bitterness'),
('Cl.Cl.COCC(=O)OC1(CCN(C)CCCC2=Nc3ccccc3N2)CCc2cc(ccc2C1C(C)C)F',
'bitterness'), ('Clc1ccc(c(c1)Cl)COC(Cn1ccnc1)c1ccc(cc1Cl)Cl', 'bitterness'),
('O=N(=O)c1cccc(c1)N(=O)=O', 'bitterness'),
('CC(CC(O)C=C(C)C)C1CCC2(C)C3C(O)C=C4C(CCC(O)C4(C)C)C3(CCC12C)C=O',
'bitterness'), ('CC1(C)CCC2(CCC3(C)C(=CCC4C5(C)CCC(OC6OC(C(O)C(OC7OCC(O)C(O)C7O)
C6O)C(O)=O)C(C)(C)C5CCC34C)C2C1)C(O)=O', 'bitterness'), ('CC1(C)CCC2(CCC3(C)C(=C
CC4C5(C)CCC(OC6OC(C(O)C(OC7OCC(O)C(O)C7O)C6O)C(O)=O)C(C)(C)C5CCC34C)C2C1)C(=O)OC
1OC(CO)C(O)C(O)C1O', 'bitterness'),
('Oc1ccc(c(c1)O)C1=C(O)C(=O)c2c(cc(cc2O1)O)O', 'bitterness'),
('Cl.CN1CCC23C4Oc5c(ccc(c25)CC1C3C=CC4O)O', 'bitterness'), ('[Ca+2].CC(CSC(=O)C(
C)NC(=O)C1CCCCC1)C(=O)N1CCCC1C([O-])=O.CC(CSC(=O)C(C)NC(=O)C1CCCCC1)C(=O)N1CCCC1
C([O-])=O', 'bitterness'), ('Oc1cc(c2c(c1)OC(=C(O)C2=O)c1cc(c(c(c1)O)O)O)O',
'bitterness'), ('[Na+].[Na+].[O-]S(=O)(=O)N(c1cccc2ccccc12)S([O-])(=O)=O',
'bitterness'), ('[Na+].[Na+].CCOC(CN(S([O-])(=O)=O)S([O-])(=O)=O)OCC',
'bitterness'), ('CC(=C)C=NO', 'bitterness'),
('[Na+].[Na+].[O-]S(=O)(=O)N(CCN1CCOCC1)S([O-])(=O)=O', 'bitterness'),
('[Na+].[Na+].[O-]S(=O)(=O)N(CCN1CCCCC1)S([O-])(=O)=O', 'bitterness'),
('COCCC=C(C)C=NO', 'bitterness'), ('COCCCCC=C(C)C=NO', 'bitterness'),
('CCN(CC)C1=NS(=O)(=O)c2cc(ccc12)N(=O)=O', 'bitterness'), ('ON=CC1=CC(CCC1)C=C',
'bitterness'), ('CC1CCC(=CC1)C=NO', 'bitterness'),
('[Na+].[Na+].CCN(CC)CCN(S([O-])(=O)=O)S([O-])(=O)=O', 'bitterness'),
('[Na+].[Na+].CN(C)CCN(S([O-])(=O)=O)S([O-])(=O)=O', 'bitterness'),
('[Na+].[Na+].CN(C)CCCN(S([O-])(=O)=O)S([O-])(=O)=O', 'bitterness'), ('[Na+].[Na
+].[Na+].[Na+].[O-]S(=O)(=O)N(CCCCCCN(S([O-])(=O)=O)S([O-])(=O)=O)S([O-])(=O)=O'
, 'bitterness'), ('CC(C)C1=CCC(=CC1)C=NO', 'bitterness'), ('COCC1CCC=C1C=NO',
'bitterness'),
('Oc1ccc(cc1O)CCC(=O)NCCCCN(CCCNC(=O)CCc1ccc(c(c1)O)O)C(=O)CCc1ccc(c(c1)O)O',

'bitterness'), ('Oc1ccc(cc1O)CCC(=O)NCCCCNCCCNC(=O)CCc1ccc(c(c1)O)O',
'bitterness'), ('Cl.C1CN=C(Cc2cccc3ccccc23)N1', 'bitterness'),
('CC(C)C1(CC(Br)=C)C(=O)NC(=O)N(C)C1=O', 'bitterness'),
('CC1OC(OC2C(O)C(O)C(CO)OC2Oc2cc(c3c(c2)OC(CC3=O)c2ccc(cc2)O)O)C(O)C(O)C1O',
'bitterness'), ('CCC(C)(C)C1(CC=C)C(=O)NC(=O)NC1=O', 'bitterness'),
('CC1OC(OC2C(O)C(O)C(CO)OC2Oc2cc(c3c(c2)OC(CC3=O)c2ccc(c(c2)O)O)O)C(O)C(O)C1O',
'bitterness'),
('COc1ccc(cc1O)C1CC(=O)c2c(cc(cc2O1)OC1OC(CO)C(O)C(O)C1OC1OC(C)C(O)C(O)C1O)O',
'bitterness'), ('COC1=CC(C)C2CC3OC(O)CC4C(=C(OC)C(=O)C(C34C)C2(C)C1=O)C',
'bitterness'), ('[Br-].CN(C)C(=O)Oc1cccc(c1)[N+](C)(C)C', 'bitterness'),
('CC(C)=CCCC(C)=CCO', 'bitterness'), ('CC(C)CC(=O)OCC=C(C)CCC=C(C)C',
'bitterness'), ('CCNC(N)=S', 'bitterness'), ('O=C(CCNNC(=O)c1ccncc1)NCc1ccccc1',
'bitterness'), ('CN1CCCC1c1cccnc1', 'bitterness'),
('OC(=O)c1cccnc1Nc1cccc(c1)C(F)(F)F', 'bitterness'), ('CCN(CC)C(=O)c1cccnc1',
'bitterness'), ('CC(C)CNC(C)=O', 'bitterness'), ('NC(=O)NN=CC1=CC=C(O1)N(=O)=O',
'bitterness'), ('CNC(N)=S', 'bitterness'),
('COc1ccc(cc1OC)C1=CC(=O)c2c(c(c(c(c2O1)OC)OC)OC)OC', 'bitterness'),
('CC=C(C)C(=O)OC1CC(=CCC(O)C(=CC2OC(=O)C(=C)C12)C)C', 'bitterness'),
('CC1(C)CCC2OC(=O)C34CC(CC(O)C3C32COC(O)C13)C(=C)C4=O', 'bitterness'),
('CC1C2CC(=O)C3C45COC(OC(C)=O)C5C(C)(C)CCC4OC(=O)C3(C2)C1=O', 'bitterness'),
('CC(=O)OC1CC(=O)OC(C)(C)C2CC(=O)C3(C)C(CCC4(C)C(OC(=O)C5OC345)C3=COC=C3)C12C',
'bitterness'), ('CCCCCCCCCOC(C)=O', 'bitterness'), ('CCCCCCCCCO', 'bitterness'),
('CCC(C)C(=O)C1=C(O)C2(CC=C(C)C)CC3C(C)(C)CCC3(C1=O)C2=O', 'bitterness'),
('CC(C)C(=O)C1=C(O)C2(CC=C(C)C)CC3C(C)(C)CCC3(C1=O)C2=O', 'bitterness'),
('CC(C)CC(=O)C1=C(O)C2(CC=C(C)C)CC3C(C)(C)CCC3(C1=O)C2=O', 'bitterness'),
('COc1ccc2c(c1OC)C(=O)OC2C1N(C)CCc2cc3c(c(c12)OC)OCO3', 'bitterness'),
('[Na+].[Na+].[O-]S(=O)(=O)Nc1ccc(cc1)N(c1ccccc1)S([O-])(=O)=O', 'bitterness'),
('CC1(C)OC(=O)C=CC2(C)C3CCC4(C)C(OC(=O)C5OC45C3(C)C(=O)CC12)C1=COC=C1',
'bitterness'), ('CC(CCC1(O)OC2CC3C4CCC5CC(CCC5(C)C4CCC3(C)C2C1C)OC1OC(CO)C(O)C(O
)C1OC1OC(CO)C(O)C(O)C1O)COC1OC(CO)C(O)C(O)C1O', 'bitterness'), ('CC(CCC1(O)OC2CC
3C4CCC5CC(CCC5(C)C4CCC3(C)C2C1C)OC1OC(CO)C(OC2OCC(O)C(O)C2O)C(O)C1OC1OC(CO)C(O)C
(O)C1O)COC1OC(CO)C(O)C(O)C1O', 'bitterness'),
('CC1COc2c(c(cc3c2N1C=C(C(O)=O)C3=O)F)N1CCN(C)CC1', 'bitterness'),
('COC(=O)C1=COC(OC2OC(CO)C(O)C(O)C2O)C(=CC)C1CC(=O)OCCc1ccc(c(c1)O)O',
'bitterness'), ('COc1ccc2c(c1)NC(=N2)S(=O)Cc1ncc(c(c1C)OC)C', 'bitterness'),
('COc1ccccc1C=O', 'bitterness'),
('CC1(C)CCC(O)C23COC(O)(C(O)C12)C12C(O)C(CCC31)C(=C)C2=O', 'bitterness'),
('CN(C)CCOC(c1ccccc1)c1ccccc1C', 'bitterness'), ('CC1OC(OC2CC(O)C3(CO)C4C(O)CC5(
C)C(CCC5(O)C4CCC3(O)C2)C2=CC(=O)OC2)C(O)C(O)C1O', 'bitterness'),
('CCN(CC)CCOCCOC(=O)C(CC)(CC)c1ccccc1', 'bitterness'),
('COc1cc2c3cc1Oc1c(c(cc4c1C(Cc1ccc(cc1)Oc1cc(ccc1O)CC3N(C)CC2)N(C)CC4)OC)OC',
'bitterness'), ('COc1ccc(cc1)NC(C)=O', 'bitterness'), ('[Br-].[Br-].CC(=O)OC1CC2
CCC3C(CCC4(C)C3CC(C4OC(C)=O)[N+]3(C)CCCCC3)C2(C)CC1[N+]1(C)CCCCC1',
'bitterness'), ('COc1ccnc(c1OC)CS(=O)C1=Nc2cc(ccc2N1)OC(F)F', 'bitterness'),
('CC(C)(CO)C(O)C(=O)NCCC(O)=O', 'bitterness'),
('[Ca+2].CC(C)(CO)C(O)C(=O)NCCC([O-])=O.CC(C)(CO)C(O)C(=O)NCCC([O-])=O',
'bitterness'), ('COc1ccc(cc1OC)Cc1nccc2cc(c(cc12)OC)OC', 'bitterness'),

('CC(=O)Nc1ccc(cc1)O', 'bitterness'), ('CC1CCC2C(OC(=O)C2=C)C2(C)C(=O)C=CC12O',
'bitterness'), ('COc1ccc(cc1)OC', 'bitterness'),
('Oc1ccc(cc1)-c1[o+]c2cc(cc(c2cc1O)O)O', 'bitterness'),
('CC1=CC=CN2C(=O)C(=CN=C12)C1=NNN=N1', 'bitterness'),
('[Br-].[Br-].C[N+](C)(C)CCCCC[N+](C)(C)C', 'bitterness'),
('NC(=N)c1ccc(cc1)OCCCCCOc1ccc(cc1)C(N)=N.OCCS(O)(=O)=O.OCCS(O)(=O)=O',
'bitterness'), ('[Na+].CCCC(C)C1(CC)C(=O)NC(=NC1=O)[O-]', 'bitterness'),
('CN1C=NC2=C1C(=O)N(CCCCC(C)=O)C(=O)N2C', 'bitterness'), ('C1CCN2N=NN=C2CC1',
'bitterness'),
('COC1CC(OC(C)C1O)OC1CCC2(C)C3CCC4(C)C(CCC4(O)C3CCC2(O)C1)C1=CC(=O)OC1',
'bitterness'), ('CC1=CC2OC(=O)C(=C)C2CCC(=C)C(CC1)OO', 'bitterness'),
('CCCCCC=CCC=CCCCCCCCC(=O)CC(O)COC(C)=O', 'bitterness'), ('Oc1cc(cc(c1O)O)C(=O)O
CC1OC(OC(=O)c2cc(c(c(c2)O)O)O)C(OC(=O)c2cc(c(c(c2)O)O)O)C(OC(=O)c2cc(c(c(c2)O)O)
O)C1OC(=O)c1cc(c(c(c1)O)O)O', 'bitterness'),
('CC(C)CC(NC(=O)C(N)Cc1ccccc1)C(O)=O', 'bitterness'), ('CCOc1ccc(cc1)NC(C)=O',
'bitterness'), ('C=CCC1(C(=O)NC(=O)NC1=O)c1ccccc1', 'bitterness'),
('Cl.Nc1ccc(c(n1)N)N=Nc1ccccc1', 'bitterness'), ('OCCc1ccccc1', 'bitterness'),
('S=C=NCCc1ccccc1', 'bitterness'), ('CCC1(C(=O)NC(=O)NC1=O)c1ccccc1',
'bitterness'), ('NC(=O)OCCCc1ccccc1', 'bitterness'),
('Cc1ccc(cc1)N(CC1=NCCN1)c1cccc(c1)O', 'bitterness'),
('OCC1OC(Oc2ccccc2)C(O)C(O)C1O', 'bitterness'), ('O=CCc1ccccc1', 'bitterness'),
('COC(Cc1ccccc1)OC', 'bitterness'), ('NC(N)=NC(N)=Nc1ccccc1', 'bitterness'),
('CCCCC1C(=O)N(N(C1=O)c1ccccc1)c1ccccc1', 'bitterness'),
('Cl.CNCC(O)c1cccc(c1)O', 'bitterness'), ('O=C=NCCc1ccccc1', 'bitterness'),
('CC1(C(=O)NC(=O)NC1=O)c1ccccc1', 'bitterness'), ('Cl.CC(N)C(O)c1ccccc1',
'bitterness'), ('NC(=S)Nc1ccccc1', 'bitterness'),
('[Na+].O=C1NC(C(=O)[N-]1)(c1ccccc1)c1ccccc1', 'bitterness'),
('NC(Cc1ccccc1)C(=O)NC(Cc1ccccc1)C(=O)NC(Cc1ccccc1)C(O)=O', 'bitterness'),
('NC(Cc1ccccc1)C(=O)NC(CC1=CNc2ccccc12)C(O)=O', 'bitterness'),
('Oc1ccc(cc1)CCC(=O)c1c(cc(cc1O)O)O', 'bitterness'),
('CC(C)CC(=O)c1c(c(c(c(c1O)C1OC(CO)C(O)C(O)C1O)O)C1OC(CO)C(O)C(O)C1O)O',
'bitterness'), ('CN1CCC23C4Oc5c(ccc(c25)CC1C3C=CC4O)OCCN1CCOCC1', 'bitterness'),
('OC(=O)c1ccccc1C(=O)Nc1ccc(cc1)S(=O)(=O)NC1=NC=CS1', 'bitterness'),
('Cc1ccccn1', 'bitterness'), ('Cl.C1CCN(CC1)CCN(Cc1ccccn1)c1ccccc1',
'bitterness'), ('Oc1c(cc(cc1N(=O)=O)N(=O)=O)N(=O)=O', 'bitterness'),
('CC1=C(C=O)C(C)(C)CC(C1)OC1OC(CO)C(O)C(O)C1O', 'bitterness'),
('CCC1OC(=O)C(C)C(=O)C(C)C(OC2OC(C)CC(C2O)N(C)C)C(C)CC(C)C(=O)C=CC1(C)O',
'bitterness'), ('CC(C)(O)C1C2OC(=O)C1C1(O)CC3OC43C(=O)OC2C14C', 'bitterness'), (
'CC(=C)C1C2OC(=O)C1C1(O)CC3OC43C(=O)OC2C14C.CC(C)(O)C1C2OC(=O)C1C1(O)CC3OC43C(=O
)OC2C14C', 'bitterness'), ('CC(=C)C1C2OC(=O)C1C1(O)CC3OC43C(=O)OC2C14C',
'bitterness'), ('CC(C)=CCCC(C)=CCCC(C)=CCN1CCN(CC1)Cc1ccc2c(c1)OCO2',
'bitterness'), ('CCCCCC(O)C(O)C=CC(O)CCCCCCCC(O)=O', 'bitterness'),
('Oc1cc(c2c(c1)OC(CC2=O)c1ccccc1)O', 'bitterness'),
('CCN1C=C(C(O)=O)C(=O)c2cnc(nc12)N1CCNCC1', 'bitterness'), ('C1CNCCN1',
'bitterness'), ('C1CCNCC1', 'bitterness'), ('CCC1(CC)C(=O)CCNC1=O',
'bitterness'), ('CC(=O)OCc1ccc2c(c1)OCO2', 'bitterness'),
('OC(=O)c1ccc2c(c1)OCO2', 'bitterness'), ('Cl.OC(C1CCCCN1)(c1ccccc1)c1ccccc1',

'bitterness'), ('CN1CCN(CC1)CC(=O)N1c2ccccc2C(=O)Nc2cccnc12', 'bitterness'),
('CCOc1ccc(cc1)NC(=O)C(C)O', 'bitterness'),
('CC(C)=CCCC(C)=CCCC(CO)=CCCC(C)=CCO', 'bitterness'), ('CC(C)C1=CCC(=CC1)C',
'bitterness'), ('COc1ccc(cc1)C=O', 'bitterness'),
('CCCCCCCCCCCC(=O)OCCOCC(OCCO)C1OCC(OCCO)C1OCCO', 'bitterness'),
('CCCCCCCCCCCCCCCCCC(=O)OCCOCC(OCCOC)C1OCC(OCCOC)C1OCCOC', 'bitterness'),
('CCCCCCCCC=CCCCCCCCC(=O)OCCOCC(OCCO)C1OCC(OCCO)C1OCCO', 'bitterness'),
('COc1ccc(cc1)C1CC(=O)c2c(cc(cc2O1)OC1OC(CO)C(O)C(O)C1OC1OC(C)C(O)C(O)C1O)O',
'bitterness'), ('[K+].OCOc1ccccc1S([O-])(=O)=O', 'bitterness'),
('[K+].[K+].[O-]S([O-])(=O)=O', 'bitterness'),
('CC12CC(O)C3C(CCC4=CC(=O)C=CC34C)C2CCC1(O)C(=O)CO', 'bitterness'),
('CC12CC(=O)C3C(CCC4=CC(=O)C=CC34C)C2CCC1(O)C(=O)CO', 'bitterness'),
('NC(N)=NCCCC(NC(=O)C1CCCN1)C(O)=O', 'bitterness'),
('CCN(CC)CCNC(=O)c1ccc(cc1)N', 'bitterness'), ('CCN(CC)CCOC(=O)c1ccc(cc1)N',
'bitterness'), ('OC1C(Oc2cc(cc(c2C1c1c(cc(c2c1OC(C(C2)OC(=O)c1cc(c(c(c1)O)O)O)c1
ccc(c(c1)O)O)O)O)O)O)c1ccc(c(c1)O)O', 'bitterness'),
('OC1Cc2c(cc(c(c2OC1c1ccc(c(c1)O)O)C1C(O)C(Oc2cc(cc(c12)O)O)c1ccc(c(c1)O)O)O)O',
'bitterness'), ('OC1Cc2c(cc(c(c2OC1c1ccc(c(c1)O)O)C1C(O)C(Oc2c(c(cc(c12)O)O)C1C(
O)C(Oc2cc(cc(c12)O)O)c1ccc(c(c1)O)O)c1ccc(c(c1)O)O)O)O', 'bitterness'),
('CC(=O)C1CCC2C3CCC4=CC(=O)CCC4(C)C3CCC12C', 'bitterness'),
('OCC1OC(SC(CC(O)C=C)=NOS(O)(=O)=O)C(O)C(O)C1O', 'bitterness'),
('CC(CN1c2ccccc2Sc2ccccc12)N(C)C', 'bitterness'),
('Cl.CCCNCC(O)COc1ccccc1C(=O)CCc1ccccc1', 'bitterness'),
('CC(C)C1(CC(Br)=C)C(=O)NC(=O)NC1=O', 'bitterness'),
('NC(=N)c1ccc(cc1)OCCCOc1ccc(cc1)C(N)=N.OCCS(O)(=O)=O.OCCS(O)(=O)=O',
'bitterness'), ('CC(C)NCC(O)COc1cccc2ccccc12', 'bitterness'),
('CCC(=O)OC(Cc1ccccc1)(C(C)CN(C)C)c1ccccc1.OS(=O)(=O)c1ccc2ccccc2c1',
'bitterness'), ('Cl.CCC(=O)OC(Cc1ccccc1)(C(C)CN(C)C)c1ccccc1', 'bitterness'),
('CCCOC(=O)C=CC1=CC=CO1', 'bitterness'), ('CCCOC(=O)c1cc(c(c(c1)O)O)O',
'bitterness'), ('CCCOC(=O)CC', 'bitterness'), ('CCCC1=CC(=O)NC(=S)N1',
'bitterness'), ('CC(C)C1=C(C)N(C)N(c2ccccc2)C1=O', 'bitterness'),
('CCOC(=O)c1ccc(c(c1)O)O', 'bitterness'), ('CC(CCC1(O)OC2CC3C4CC=C5CC(CCC5(C)C4C
CC3(C)C2C1C)OC1OC(CO)C(OC2OC(C)C(O)C(O)C2O)C(O)C1OC1OC(C)C(O)C(O)C1O)COC1OC(CO)C
(O)C(O)C1O', 'bitterness'), ('COc1cc(c2c(c1)OC=C(C2=O)c1ccc(cc1)O)O',
'bitterness'), ('OCC1OC(Oc2cc(c3c(c2)OC(CC3=O)c2ccc(cc2)O)O)C(O)C(O)C1O',
'bitterness'), ('CNC(C)C(O)c1ccccc1.OS(O)(=O)=O', 'bitterness'),
('CCC1CN2CCc3cc(c(cc3C2CC1CC1=C2C=C(OC)C(=O)C=C2CCN1)OC)OC', 'bitterness'),
('N1C=Nc2ncncc12', 'bitterness'), ('c1cnccn1', 'bitterness'), ('N1C=CC=N1',
'bitterness'), ('c1ccnnc1', 'bitterness'), ('c1ccncc1', 'bitterness'),
('[Br-].CN(C)C(=O)Oc1ccc[n+](c1)C', 'bitterness'), ('c1cncnc1', 'bitterness'),
('Oc1ccccc1O', 'bitterness'), ('COC1=CC=C2Nc3cc(ccc3C(=C2N1)N=C1C=C(CN2CCCC2)C(=
O)C(=C1)CN1CCCC1)Cl.OP(O)(O)=O.OP(O)(O)=O.OP(O)(O)=O.OP(O)(O)=O', 'bitterness'),
('N1C=CC=C1', 'bitterness'), ('C1CCNC1', 'bitterness'),
('CC1(O)CC(=C(O)C1=O)N1CCCC1', 'bitterness'), ('CC(C)CC(NC(=O)C(N)CCC(N)=O)C(=O)
NC(Cc1ccccc1)C(=O)NCC(=O)N1CCCC1C(=O)NC(CC(N)=O)C(=O)NC(C(C)C)C(=O)NC(CC(N)=O)C(
=O)N1CCCC1C(=O)NC(CC1=CNc2ccccc12)C(=O)NC(CC1=CN=CN1)C(=O)NC(CC(N)=O)C(=O)N1CCCC
1C(O)=O', 'bitterness'),

('COC1=CC(C)C2CC3OC(=O)CC4C(=C(OC)C(=O)C(C34C)C2(C)C1=O)C', 'bitterness'), ('CCC12CCCN(CCC3=C(CC1)Nc1ccccc31)C2', 'bitterness'), ('Oc1ccc(cc1O)C1=C(O)C(=O)c2c(c(c(cc2O1)O)O)O', 'bitterness'), ('Oc1cc(c2c(c1)OC(=C(O)C2=O)c1ccc(c(c1)O)O)O', 'bitterness'), ('OCC1CC(OC2=C(Oc3cc(cc(c3C2=O)O)O)c2ccc(c(c2)O)O)C(O)C(O)C1O', 'bitterness'), ('Cl.Cl.CCN(CC)CCCC(C)Nc1c2ccc(cc2nc2ccc(cc12)OC)Cl', 'bitterness'), ('c1ccc2c(c1)C=NC=N2', 'bitterness'), ('OC1CC(O)(CC(O)C1O)C(O)=O', 'bitterness'), ('COc1ccc2nccc(c2c1)C(O)C1CC2CCN1CC2C=C.COc1ccc2nccc(c2c1)C(O)C1C C2CCN1CC2C=C.OS(O)(=O)=O', 'bitterness'), ('COc1ccc2nccc(c2c1)C(O)C1CC2CCN1CC2C=C', 'bitterness'), ('Cl.Cl.COc1ccc2nccc(c2c1)C(O)C1CC2CCN1CC2C=C', 'bitterness'), ('Cl.COc1ccc2nccc(c2c1)C(O)C1CC2CCN1CC2C=C', 'bitterness'), ('COc1ccc2nccc(c2c1)C(O)C1CC2CCN1CC2C=C.OS(O)(=O)=O', 'bitterness'), ('OCC1C(=Cc2ccco2)C(=O)c2cc(c(c[n+]12)[O-])Cc1ccco1', 'bitterness'), ('CC1CCC2(CCC3(C(O)=O)C(=CCC4C5(C)CCC(O)C(C)(C)C5CCC34C)C2C1C)C(O)=O', 'bitterness'), ('CC1(C)C=CC(=O)C23COC(O)(C(O)C12)C12CC(CCC31)C(=C)C2=O', 'bitterness'), ('CC(=O)OC1C(=C)C2CCC3C45COC(O)(C(O)C5C(C)(C)C=CC4=O)C13C2', 'bitterness'), ('CC(=O)OCC1OC(OCC2OC(OC3(COC(C)=O)OC(COC(C)=O)C(OC(C)=O)C3OC(C)= O)C(OC(C)=O)C(OC(C)=O)C2OC(C)=O)C(OC(C)=O)C(OC(C)=O)C1OC(C)=O', 'bitterness'), ('Cl.CNC(NCCSCC1=CC=C(CN(C)C)O1)=CN(=O)=O', 'bitterness'), ('Oc1ccc(cc1)C=Cc1cc(cc(c1)O)O', 'bitterness'), ('OC1CCCN2CC3CC(CN4CCCCC34)C12', 'bitterness'), ('CC(CCCC(C)=C)CCOC(C)=O', 'bitterness'), ('CC(CCc1ccc(cc1)O)OC1OC(CO)C(O)C(O)C1O', 'bitterness'), ('OCC1OC(Oc2ccccc2CO)C(O)C(O)C1O', 'bitterness'), ('NC(=O)c1ccccc1O', 'bitterness'), ('OC(=O)c1ccccc1OC(=O)c1ccccc1O', 'bitterness'), ('CC1=CCC(O)C2(C)CCC3C(OC(=O)C3=C)C12', 'bitterness'), ('CC(=O)OCC1OC(OC2OC=C3C( CCOC3=O)C2C=C)C(OC(=O)c2ccccc2)C(OC(C)=O)C1OC(=O)c1cccc(c1O)OC1OC(CO)C(O)C(O)C1O ', 'bitterness'), ('CC1OC(OC2CCC3(C)C4CCC5(C)C(CCC5(O)C4CCC3=C2)C2=COC(=O)C=C2)C (O)C(OC2OC(CO)C(O)C(O)C2O)C1O', 'bitterness'), ('CC(C)C(=O)C1=C(O)C2(O)C(CC3C(C)(C)OC(C=C(C)C)C23O)C1=O', 'bitterness'), ('CC(C)CC(=O)C1=C(O)C2(O)C(CC3C(C)(C)OC(C=C(C)C)C23O)C1=O', 'bitterness'), ('Oc1ccc(cc1)C1=CC(=O)c2c(c(c(cc2O1)O)O)O', 'bitterness'), ('[Na+].CCCC(C)C1(CC=C)C(=O)NC(=O)[N-]C1=O', 'bitterness'), ('CC=C1CC(C)C(C)(O)C(=O)OCC2=CCN3CCC(OC1=O)C23', 'bitterness'), ('CC(=O)OC1CC2CC3(C1C14COC3(O)C(O)C4C(C)(C)CCC1OC(C)=O)C(=O)C2=C', 'bitterness'), ('COc1cc(ccc1O)C1Oc2cc(ccc2OC1CO)C1Oc2cc(cc(c2C(=O)C1O)O)O', 'bitterness'), ('COc1cc(cc(c1O)OC)C=CC(O)=O', 'bitterness'), ('COc1cc(cc(c1O)OC)C=CC(=O)OCC[N+](C)(C)C', 'bitterness'), ('COc1ccc(cc1OC)C1=CC(=O)c2c(cc(c(c2OC)OC)OC)O1', 'bitterness'), ('OCC1OC(SC(CC=C)=NOS([O-])(=O)=O)C(O)C(O)C1O', 'bitterness'), ('[K+].OCC1OC(SC(CC=C)=NOS([O-])(=O)=O)C(O)C(O)C1O', 'bitterness'), ('CC1C2CC(OC(C)=O)C(=CCC(OC(C)=O)C(=CC2OC1=O)C)C', 'bitterness'), ('COc1ccc2c(c3c(nc2c1OC)OC=C3)OC', 'bitterness'), ('[Na+].[O-]C(=O)c1ccccc1', 'bitterness'), ('[Na+].[Br-]', 'bitterness'), ('[Na+].[O-]S(=O)(=O)Nc1ccc2c(c1)OCCO2', 'bitterness'), ('[Na+].[O-]S(=O)(=O)Nc1ccc(cc1Br)Br', 'bitterness'), ('[Na+].CCc1cccc(c1NS([O-])(=O)=O)CC', 'bitterness'),

```
('[Na+].[O-]S(=O)(=O)Nc1ccccc1Cc1ccccc1', 'bitterness'),
('[Na+].CC(C)c1ccc(c(c1)Br)NS([O-])(=O)=O', 'bitterness'),
('[Na+].CCCCOC(=O)c1ccccc1NS([O-])(=O)=O', 'bitterness'),
('[Na+].[O-]S(=O)(=O)Nc1ccc(cc1Cl)F', 'bitterness'),
('[Na+].[O-]S(=O)(=O)Nc1ccc(cc1Cl)N(=O)=O', 'bitterness'),
('[Na+].Cc1cc(ccc1NS([O-])(=O)=O)N(=O)=O', 'bitterness'),
('[Na+].CCOC(=O)c1cccc(c1)NS([O-])(=O)=O', 'bitterness'),
('[Na+].CCc1ccc(nc1NS([O-])(=O)=O)C', 'bitterness'),
('[Na+].Cc1ccc(cc1F)NS([O-])(=O)=O', 'bitterness'),
('[Na+].Cc1ccc(cc1I)NS([O-])(=O)=O', 'bitterness'),
('[Na+].[O-]S(=O)(=O)Nc1cccc(c1)Oc1ccccc1', 'bitterness'),
('[Na+].[O-]S(=O)(=O)Nc1cccc(c1)OCc1ccccc1', 'bitterness'),
('[Na+].CC(C)(C)c1cccc(c1)NS([O-])(=O)=O', 'bitterness'),
('[Na+].[O-]S(=O)(=O)Nc1ccc(cc1Cl)Br', 'bitterness'),
('[Na+].[O-]S(=O)(=O)Nc1ccc(cc1F)Br', 'bitterness'),
('[Na+].[O-]S(=O)(=O)NC1=Nc2c(cccc2S1)Cl', 'bitterness'),
('[Na+].[O-]S(=O)(=O)Nc1ccc(cc1N(=O)=O)Cl', 'bitterness'),
('[Na+].Cc1cc(ccc1NS([O-])(=O)=O)F', 'bitterness'),
('[Na+].[O-]S(=O)(=O)Nc1ccc(cc1N(=O)=O)F', 'bitterness'),
('[Na+].Cc1cc(ccc1F)NS([O-])(=O)=O', 'bitterness'),
('[Na+].COc1ccc(c(c1)C)NS([O-])(=O)=O', 'bitterness'),
('[Na+].COC(=O)c1ccc(cc1)NS([O-])(=O)=O', 'bitterness'),
('[Na+].CC(C)(C)C1=CSC(=N1)NS([O-])(=O)=O', 'bitterness'),
('[Na+].Cc1nc(ccc1Br)NS([O-])(=O)=O', 'bitterness'),
('[Na+].[O-]S(=O)(=O)Nc1cc(ccc1N(=O)=O)Cl', 'bitterness'),
('[Na+].[O-]S(=O)(=O)NC1=Nc2ccc(cc2S1)N(=O)=O', 'bitterness'),
('[Na+].CCC(C)c1ccccc1NS([O-])(=O)=O', 'bitterness'),
('[Na+].CC(C)c1cccc(c1NS([O-])(=O)=O)C(C)C', 'bitterness'),
('[Na+].CN(C)c1cccc(c1)NS([O-])(=O)=O', 'bitterness'),
('[Na+].COC(=O)c1cc(cc(c1)C(=O)OC)NS([O-])(=O)=O', 'bitterness'),
('[Na+].CC(O)c1ccc(cc1)NS([O-])(=O)=O', 'bitterness'),
('[Na+].[O-]S(=O)(=O)Nc1ccc(cc1)S(=O)(=O)c1ccc(cc1)N(=O)=O', 'bitterness'),
('[Na+].CSc1ccc(cc1)NS([O-])(=O)=O', 'bitterness'),
('[Na+].[O-]S(=O)(=O)Nc1ccc(cc1)S(=O)(=O)N1CCCCC1', 'bitterness'),
('[Na+].CC(C)(C)OC(=O)c1ccc(cc1)NS([O-])(=O)=O', 'bitterness'),
('[Na+].[O-]S(=O)(=O)Nc1ccccc1-c1ccccc1', 'bitterness'),
('[Na+].[O-]S(=O)(=O)c1cccc(c1)N(=O)=O', 'bitterness'),
('[Na+].Oc1ccccc1S([O-])(=O)=O', 'bitterness'),
('[Na+].[Na+].[O-]S([O-])(=O)=O', 'bitterness'), ('[Na+].[S-]C#N',
'bitterness'), ('CC1CCC2C(C)C3C(CC4C5CC=C6CC(O)CCC6(C)C5CCC34C)N2C1',
'bitterness'), ('OS(O)(=O)=O.C1CCN2CC3CC(CN4CCCCC34)C2C1', 'bitterness'),
('C1CCN2CC3CC(CN4CCCCC34)C2C1', 'bitterness'),
('CC(=O)OC1C2OC2(C)CCC=C(C)CC2OC(=O)C(=C)C12', 'bitterness'),
('CNC1C(O)C(O)C(CO)OC1OC1C(OC(C)C1(O)C=O)OC1C(O)C(O)C(N=C(N)N)C(O)C1N=C(N)N',
'bitterness'), ('O=C1CC2OCC=C3CN4CCC56C4CC3C2C6N1c1ccccc51', 'bitterness'),
('[O-][N+]12CCC34C2CC2C5C(CC(=O)N(C35)c3ccccc43)OCC=C2C1', 'bitterness'),
('[Cl-].[Cl-].C[N+](C)(C)CCOC(=O)CCC(=O)OCC[N+](C)(C)C', 'bitterness'), ('CC(=O)
```

OCC1OC(OC2(COC(C)=O)OC(COC(C)=O)C(OC(C)=O)C2OC(C)=O)C(OC(C)=O)C(OC(C)=O)C1OC(C)=
O', 'bitterness'),
('CC1(C)C2CCC34CC(CCC3C2(C)CCC1=O)C(O)(COC1OC(CO)C(O)C(O)C1O)C4', 'bitterness'),
('[Na+].Cc1ccnc(n1)[N-]S(=O)(=O)c1ccc(cc1)N', 'bitterness'),
('COc1cnc(nc1)NS(=O)(=O)c1ccc(cc1)N', 'bitterness'),
('CC1=CC(=NO1)NS(=O)(=O)c1ccc(cc1)N', 'bitterness'),
('COc1ccc(nn1)NS(=O)(=O)c1ccc(cc1)N', 'bitterness'),
('CC1=NOC(=C1C)NS(=O)(=O)c1ccc(cc1)N', 'bitterness'), ('[Na+].[Na+].Oc1ccc(cc1S(
[O-])(=O)=O)C1(OC(=O)c2c(c(c(c(c12)Br)Br)Br)Br)c1ccc(c(c1)S([O-])(=O)=O)O',
'bitterness'), ('CCC(C)(S(=O)(=O)CC)S(=O)(=O)CC', 'bitterness'), ('O=S=O',
'bitterness'), ('Oc1ccc2c(c1)OC(=Cc1ccc(c(c1)O)O)C2=O', 'bitterness'),
('COC(=O)C(NC(=O)C(CC(O)=O)NC(=S)Nc1ccc(cc1)C#N)C(=O)OC1C(C)(C)C2CCC1(C)C2',
'bitterness'), ('CCCCC1(COC(=O)CCC(O)=O)C(=O)N(N(C1=O)c1ccccc1)c1ccccc1',
'bitterness'), ('OCC1OC(OC2OC=C3C(=O)OCCC3(O)C2C=C)C(O)C(O)C1O', 'bitterness'),
('CCOC(=O)c1cc(c(c(c1)OC)O)OC', 'bitterness'), ('Cl.NC1=C2CCCCC2=Nc2ccccc12',
'bitterness'), ('CCC(C)C1(CC=C)C(=O)NC(=O)NC1=O', 'bitterness'),
('COc1ccc(cc1)C1=CC(=O)c2c(c(c(c(c2O1)OC)OC)OC)OC', 'bitterness'),
('CC1=CC(O)C2C(OC(=O)C2=C)C=C(C)C(O)CC1', 'bitterness'),
('CC1=CC(O)C2C(CC(=C)C(O)CC1)OC(=O)C2=C', 'bitterness'),
('CC(=O)OC1CCC(=CC(OC(C)=O)C2C(OC(=O)C2=C)C=C1C)C', 'bitterness'),
('NCCS(O)(=O)=O', 'bitterness'),
('CC(CCC(=O)NCCS(O)(=O)=O)C1CCC2C3C(O)CC4CC(O)CCC4(C)C3CC(O)C12C',
'bitterness'), ('CCC(C)C(=O)OC(C)C1(O)CCC2(O)C3CCC4CC(CCC4(C)C3CC(OC(C)=O)C12C)O
C1CC(OC)C(OC2CC(OC)C(OC3OC(C)C(OC4OC(CO)C(O)C(O)C4O)C(OC)C3O)C(C)O2)C(C)O1',
'bitterness'), ('CC(C)(O)C1CCC(C)(O)CC1', 'bitterness'),
('CC1=CCC(CC1)C(C)(C)OC=O', 'bitterness'), ('CC(C)CC(=O)OC(C)(C)C1CCC(=CC1)C',
'bitterness'), ('CCC(=O)OC(C)(C)C1CCC(=CC1)C', 'bitterness'),
('Cl.CCCCNc1ccc(cc1)C(=O)OCCN(C)C', 'bitterness'),
('CCC(C)C1=C2C(=O)C3CC(C(C)(C)O)C4(O)CC(C(C)(C)O1)C2(O)C34O', 'bitterness'),
('CC(C)C1=C2C(=O)C3CC(C(C)(C)O)C4(O)CC(C(C)(C)O1)C2(O)C34O', 'bitterness'),
('CC(C)CC1=C2C(=O)C3CC(C(C)(C)O)C4(O)CC(C(C)(C)O1)C2(O)C34O', 'bitterness'),
('[OH-].CC[N+](CC)(CC)CC', 'bitterness'),
('CC1C2CC(O)C3C45COC(O)C5C(C)(C)CCC4OC(=O)C3(C2)C1O', 'bitterness'),
('CC1CC(O)C2(COC(C)=O)C(CCC(=O)C32CO3)C21CC(OC2O)C1=COC=C1', 'bitterness'),
('CC1CC(OC2OC(CO)C(O)C(O)C2O)C2=C(CCCC2C21CC(OC2=O)C1=COC=C1)COC(C)=O',
'bitterness'), ('CC1CC(O)C2(COC(C)=O)C(CC(O)CC32CO3)C21CC(OC2=O)C1=COC=C1',
'bitterness'), ('CN1C=NC2=C1C(=O)NC(=O)N2C', 'bitterness'),
('CN1C(=O)N(C)C2=C(NC=N2)C1=O', 'bitterness'),
('[Na+].CN1C(=O)N(C)C2=C(NC=N2)C1=O.CC([O-])=O', 'bitterness'),
('CC(=O)Nc1ccc(cc1)C=NNC(N)=S', 'bitterness'),
('Cc1ncc(c(n1)N)C[N+]1=CSC(=C1C)CCO', 'bitterness'),
('Cl.[Cl-].Cc1ncc(c(n1)N)C[N+]1=CSC(=C1C)CCO', 'bitterness'),
('CC(=S)Nc1ccccc1', 'bitterness'), ('S=C(Nc1ccccc1)Nc1ccccc1', 'bitterness'),
('NC(=S)NCC=C', 'bitterness'), ('Cl.OCCN1CCN(CC1)C(=O)CN1C(=O)Sc2ccc(cc12)Cl',
'bitterness'),
('COc1ccc2cc1-c1cc(ccc1O)CC1N(C)CCc3cc4c(cc13)Oc1c(c(cc3c1C(C2)N(C)CC3)OC)O4',
'bitterness'), ('Cl.CCOC(=O)C1=C(N)SC2=C1CCN(Cc1ccccc1)C2', 'bitterness'),

('Cl.C1CN=C(Cc2ccccc2)N1', 'bitterness'), ('CC1CCC2(NC1)OC1CC3C4CCC5CC(CCC5(C)C4
CCC3(C)C1C2C)OC1OC(CO)C(OC2OC(CO)C(O)C(OC3OCC(O)C(O)C3O)C2OC2OC(CO)C(O)C(O)C2O)C
(O)C1O', 'bitterness'), ('CC1(C)OC2COC3(COS(N)(=O)=O)OC(C)(C)OC3C2O1',
'bitterness'), ('Cl.COc1cccc(c1)C1(O)CCCCC1CN(C)C', 'bitterness'),
('COc1cc(ccc1O)C=CC(=O)NCCc1ccc(cc1)O', 'bitterness'),
('CCC(C)C(=O)Oc1ccc(cc1C=CC)OC', 'bitterness'), ('CCN(CC)C1=CC(=NC2=NC=NN12)C',
'bitterness'), ('CC(=O)OCC(COC(C)=O)OC(C)=O', 'bitterness'),
('CCCC(=O)OCC(COC(=O)CCC)OC(=O)CCC', 'bitterness'),
('Oc1cc(c2c(c1)OC(=CC2=O)c1cc(c(c(c1)O)O)O)O', 'bitterness'),
('CCC(C)C(=O)C1=C(O)C2(O)C(CC3C(C)(C)C(CC23O)C(C)=C)C1=O', 'bitterness'),
('CCC(C)C(=O)C1=C(O)C2(O)C(CC3C(C)(C)C(CC23O)C(C)(C)O)C1=O', 'bitterness'),
('CCC(C)C(=O)C1=C(O)C2(O)C(CC3C(C)(C)OC(O)CC23O)C1=O', 'bitterness'),
('CCC(C)C(=O)C1=C(O)C2(CC=C(C)C)CC3C(C)(C)C(CC3(C1=O)C2=O)C(C)C', 'bitterness'),
('CC(C)C(=O)C1=C(O)C2(O)C(CC3C(C)(C)C(CC23O)C(C)=C)C1=O', 'bitterness'),
('CC(C)C(=O)C1=C(O)C2(O)C(CC3C(C)(C)C(CC23O)C(C)(C)O)C1=O', 'bitterness'),
('CC(C)C(=O)C1=C(O)C2(O)C(CC3C(C)(C)OC(O)CC23O)C1=O', 'bitterness'),
('CC(C)C1CC23C(CC(CC=C(C)C)(C(=C(C(=O)C(C)C)C2=O)O)C3=O)C1(C)C', 'bitterness'),
('CC(C)CC(=O)C1=C(O)C2(O)C(CC3C(C)(C)C(CC23O)C(C)=C)C1=O', 'bitterness'),
('CC(C)CC(=O)C1=C(O)C2(O)C(CC3C(C)(C)C(CC23O)C(C)(C)O)C1=O', 'bitterness'),
('CC(C)CC(=O)C1=C(O)C2(O)C(CC3C(C)(C)OC(O)CC23O)C1=O', 'bitterness'),
('CC(C)CC(=O)C1=C(O)C2(CC=C(C)C)CC3C(C)(C)C(CC3(C1=O)C2=O)C(C)C', 'bitterness'),
('[I-].CC[N+](CC)(CC)CCC(O)(C1CCCCC1)c1ccccc1', 'bitterness'),
('CCOC(=O)CC(O)(CC(=O)OCC)C(=O)OCC', 'bitterness'), ('CN1C(=O)OC(C)(C)C1=O',
'bitterness'), ('CC1(C)C2CCC1(CS([O-])(=O)=O)C(=O)C2.O=C1N(Cc2ccccc2)C2C[S+]3CCC
C3C2N1Cc1ccccc1', 'bitterness'), ('COc1cc(cc(c1OC)OC)Cc1cnc(nc1N)N',
'bitterness'), ('CC(C)CC(NC(=O)C(N)CC1=CNc2ccccc12)C(O)=O', 'bitterness'),
('NC(CC1=CNc2ccccc12)C(=O)NC(Cc1ccccc1)C(O)=O', 'bitterness'),
('NC(CC1=CNc2ccccc12)C(=O)N1CCCC1C(O)=O', 'bitterness'),
('NC(CC1=CNc2ccccc12)C(=O)NC(CC1=CNc2ccccc12)C(O)=O', 'bitterness'),
('NC(CC1=CNc2ccccc12)C(=O)NC(CC1=CNc2ccccc12)C(=O)NC(CC1=CNc2ccccc12)C(O)=O',
'bitterness'), ('Oc1ccc2c(c1)OC(=O)C=C2', 'bitterness'),
('CC(=O)CCCC1(C)CC2OC(=O)C(=C)C2CC1=O', 'bitterness'), ('NC(N)=O',
'bitterness'), ('CCCC(CCC)C(N)=O', 'bitterness'), ('CNC(CC(C)C)C(=O)NC1C(O)c2ccc
(c(c2)Cl)Oc2cc3cc(c2OC2OC(CO)C(O)C(O)C2OC2CC(C)(N)C(O)C(C)O2)Oc2ccc(cc2Cl)C(O)C2
NC(=O)C(NC(=O)C3NC(=O)C(CC(N)=O)NC1=O)c1ccc(c(c1)-c1c(cc(cc1C(NC2=O)C(O)=O)O)O)O
', 'bitterness'), ('CCOC(=O)c1ccc(c(c1)OC)O', 'bitterness'),
('CC12CCCC3(C1CCC14CC(CCC31)C(=C)C4O)C1OCCN1C2', 'bitterness'),
('COC(=O)C1=COC(OC2OC(CO)C(O)C(O)C2O)C2C(C)CC(=O)C12', 'bitterness'),
('[Na+].CCC=C(C)C1(CC)C(=O)NC(=O)[N-]C1=O', 'bitterness'),
('Cl.COc1ccc2nccc(c2c1)C(=O)CCC1CCNCC1C=C', 'bitterness'), ('CC1(C)CCC2(CCC3(C)C
(=CCC4C5(C)CCC(OC6OC(CO)C(O)C(O)C6O)C(C)(C)C5CCC34C)C2C1)C(O)=O', 'bitterness'),
('CC12CCC(O)OC2C2OC(=O)C(=C)C2CCC(=O)C1', 'bitterness'),
('[Na+].CC(=O)CC(c1ccccc1)C1=C([O-])c2ccccc2OC1=O', 'bitterness'),
('O=C1NC(=O)C2=C(N1)N=CN2', 'bitterness'),
('COc1cc(c(c(c1C(=O)C=Cc1ccc(cc1)O)O)CC=C(C)C)O', 'bitterness'),
('COc1cc2c(c(c1C(=O)C=Cc1ccc(cc1)O)O)CC(O)C(C)(C)O2', 'bitterness'),
('COc1cc2c(c(c1C(=O)C=Cc1ccc(cc1)O)O)C=CC(C)(C)O2', 'bitterness'),

```
('COc1cc(c(c(c1C(=O)C=Cc1ccc(cc1)O)O)CC(O)C(C)=C)O', 'bitterness'),
('COc1cc(c(c(c1C(=O)C=Cc1ccc(cc1)O)O)CC(O)C(C)(C)O)O', 'bitterness'),
('COc1cc(c(c(c1C(=O)C=Cc1ccc(cc1)O)O)CCC(C)(C)O)O', 'bitterness'),
('COc1cc(c2c(c1C(=O)C=Cc1ccc(cc1)O)OC(C2)C(C)(C)O)O', 'bitterness'),
('COc1cc(c2c(c1C(=O)C=Cc1ccc(cc1)O)OC(C)(C)C(O)C2)O', 'bitterness'),
('COc1cc(c(c(c1C(=O)C=Cc1ccc(cc1)O)O)CCC(C)(C)OC)O', 'bitterness'),
('COc1cc(c(c(c1C(=O)C=Cc1ccc(cc1)O)O)CCC(C)=C)O', 'bitterness'),
('COc1cc(c2c(c1C(=O)C=Cc1ccc(cc1)O)OC=C2)O', 'bitterness'),
('CCOC(C)(C)CCc1c(cc(c(c1O)C(=O)C=Cc1ccc(cc1)O)OC)O', 'bitterness'),
('O=C1c2ccccc2Oc2ccccc12', 'bitterness'),
('COC(=O)C1C(O)CCC2CN3CCc4c([nH]c5ccccc45)C3CC12', 'bitterness'),
('CC(=O)OC1CC2C(C3OC(=O)C(=C)C3CCC2=C)C1=C', 'bitterness'),
('Cc1cc2c(cc1C)N(CC(O)C(O)C(O)CO)C1=NC(=O)NC(=O)C1=N2', 'bitterness'), ('CC1CCC2
C(C)C3C(CC4C5CC=C6CC(CCC6(C)C5CCC34C)OC3OC(CO)C(OC4OC(C)C(O)C(O)C4O)C(O)C3OC3OC(
C)C(O)C(O)C3O)N2C1', 'bitterness'), ('CCC=CCC=CCC=CCCCCCCCC(O)=O',
'bitterness'), ('CC1C2CCC3(C)C=CC(=O)C(=C3C2OC1=O)C', 'bitterness'), ('CC1CCC2C(
C)C3C(CC4C5CC=C6CC(CCC6(C)C5CCC34C)OC3OC(CO)C(O)C(OC4OC(CO)C(O)C(O)C4O)C3OC3OC(C
)C(O)C(O)C3O)N2C1', 'bitterness'),
('OCC1OC(OC(=O)c2cc(c(c(c2)O)O)O)C(O)C(O)C1O', 'bitterness'),
('CC(C([O-])=O)[n+]1cc(ccc1CO)O', 'umaminess'), ('CCC=CCCC=CC(=O)NC1CC1',
'umaminess'), ('CC(C)=CCCC(C)=CCNC(=O)C1CC1', 'umaminess'),
('CC(NCC(O)=O)C(O)=O', 'umaminess'), ('OCC1OC(OC(CC(O)=O)C(O)=O)C(O)C(O)C1O',
'umaminess'), ('CCCCNC(=O)C(C)NC1=NC2=C(N=CN2C2OC(COP(O)(O)=O)C(O)C2O)C(=O)N1',
'umaminess'), ('OC1C(O)C(OC1COP(O)(O)=O)N1C=NC2=C1N=C(NC2=O)SCC=C',
'umaminess'), ('OC1C(O)C(OC1COP(O)(O)=O)N1C=NC2=C1N=C(NC2=O)SCC1=CC=CO1',
'umaminess'), ('OC1C(O)C(OC1COP(O)(O)=O)N1C=NC2=C1NC(=S)NC2=O', 'umaminess'),
('NC1=NC2=C(N=CN2C2OC(COP(O)(O)=O)C(O)C2O)C(=S)N1', 'umaminess'),
('Nc1ncnc2c1N=CN2C1OC(COP(O)(O)=O)C(O)C1O', 'umaminess'),
('NC(CC(O)=O)C(=O)NC(CCC(O)=O)C(=O)NC(CO)C(O)=O', 'umaminess'),
('[Ca+2].OC1C(O)C(OC1COP([O-])([O-])=O)N1C=NC2=C1N=C(NC2=O)OCC=C', 'umaminess'),
('[Ca+2].CC(C)=CCSC1=NC2=C(N=CN2C2OC(COP([O-])([O-])=O)C(O)C2O)C(=O)N1',
'umaminess'), ('[Ca+2].NC(CCC(O)=O)C([O-])=O.NC(CCC(O)=O)C([O-])=O',
'umaminess'), ('[Ca+2].NC1=NC2=C(N=CN2C2OC(COP([O-])([O-])=O)C(O)C2O)C(=O)N1',
'umaminess'), ('[Ca+2].OC1C(O)C(OC1COP([O-])([O-])=O)N1C=NC2=C1N=CNC2=O',
'umaminess'), ('[Ca+2].[Ca+2].NC1=NC2=C(N=CN2C2OC(COP([O-])([O-])=O)C(O)C2O)C(=O
)N1.OC1C(O)C(OC1COP([O-])([O-])=O)N1C=NC2=C1N=CNC2=O', 'umaminess'),
('CC(C)C1CCC(C)CC1NC(=O)C1CC1', 'umaminess'),
('[K+].[K+].NC1=NC2=C(N=CN2C2OC(COP([O-])([O-])=O)C(O)C2O)C(=O)N1',
'umaminess'), ('[K+].[K+].OC1C(O)C(OC1COP([O-])([O-])=O)N1C=NC2=C1N=CNC2=O',
'umaminess'),
('[Na+].[Na+].CC(=C)CSC1=NC2=C(N=CN2C2OC(COP([O-])([O-])=O)C(O)C2O)C(=O)N1',
'umaminess'),
('[Na+].[Na+].CC=CCSC1=NC2=C(N=CN2C2OC(COP([O-])([O-])=O)C(O)C2O)C(=O)N1',
'umaminess'),
('[Na+].[Na+].CC1(C)OC2C(COP([O-])([O-])=O)OC(C2O1)N1C=NC2=C1N=C(N)NC2=O',
'umaminess'),
('[Na+].[Na+].CC1(C)OC2C(COP([O-])([O-])=O)OC(C2O1)N1C=NC2=C1N=CNC2=O',
```

```
'umaminess'),
('[Na+].[Na+].OC1C(O)C(OC1COP([O-])([O-])=O)N1C=NC2=C1N=C(Cl)NC2=O',
'umaminess'),
('[Na+].[Na+].CCOC1=NC2=C(N=CN2C2OC(COP([O-])([O-])=O)C(O)C2O)C(=O)N1',
'umaminess'),
('[Na+].[Na+].CCOC(=O)CCSC1=NC2=C(N=CN2C2OC(COP([O-])([O-])=O)C(O)C2O)C(=O)N1',
'umaminess'),
('[Na+].[Na+].CCOCCSC1=NC2=C(N=CN2C2OC(COP([O-])([O-])=O)C(O)C2O)C(=O)N1',
'umaminess'),
('[Na+].[Na+].CCC1=NC2=C(N=CN2C2OC(COP([O-])([O-])=O)C(O)C2O)C(=O)N1',
'umaminess'),
('[Na+].[Na+].CCSC1=NC2=C(N=CN2C2OC(COP([O-])([O-])=O)C(O)C2O)C(=O)N1',
'umaminess'),
('[Na+].[Na+].OC1C(O)C(OC1COP([O-])([O-])=O)N1C=NC2=C1N=C(NC2=O)SCC1=CC=CO1',
'umaminess'),
('[Na+].[Na+].CC(C)OC1=NC2=C(N=CN2C2OC(COP([O-])([O-])=O)C(O)C2O)C(=O)N1',
'umaminess'),
('[Na+].[Na+].COC1=NC2=C(N=CN2C2OC(COP([O-])([O-])=O)C(O)C2O)C(=O)N1',
'umaminess'),
('[Na+].[Na+].CC1=NC2=C(N=CN2C2OC(COP([O-])([O-])=O)C(O)C2O)C(=O)N1',
'umaminess'),
('[Na+].[Na+].Cc1nc(c2c(n1)N(C=N2)C1OC(COP([O-])([O-])=O)C(O)C1O)S',
'umaminess'),
('[Na+].[Na+].CSC1=NC2=C(N=CN2C2OC(COP([O-])([O-])=O)C(O)C2O)C(=O)N1',
'umaminess'),
('[Na+].[Na+].CSc1nc(c2c(n1)N(C=N2)C1OC(COP([O-])([O-])=O)C(O)C1O)S',
'umaminess'),
('[Na+].[Na+].CCCOC1=NC2=C(N=CN2C2OC(COP([O-])([O-])=O)C(O)C2O)C(=O)N1',
'umaminess'),
('[Na+].[Na+].OC1C(O)C(OC1COP([O-])([O-])=O)N1C=NC2=C1N=C(NC2=O)c1ccccc1',
'umaminess'),
('[Na+].[Na+].OC1C(O)C(OC1COP([O-])([O-])=O)N1C=NC2=C1N=C(NC2=O)SCC1CCCO1',
'umaminess'), ('[Na+].[Na+].Nc1ncnc2c1N=CN2C1OC(COP([O-])([O-])=O)C(O)C1O',
'umaminess'), ('[Na+].[Na+].OC1C(O)C(OC1COP([O-])([O-])=O)N1C=Nc2c(ncnc12)Cl',
'umaminess'), ('[Na+].[Na+].OC1C(O)C(OC1COP([O-])([O-])=O)N1C=Nc2c(ncnc12)S',
'umaminess'),
('[Na+].[Na+].NC1=NC2=C(N=CN2C2CC(O)C(COP([O-])([O-])=O)O2)C(=O)N1',
'umaminess'), ('[Na+].[Na+].OC1C(O)C(OC1COP([O-])([O-])=O)N1C=NC2=C1N=CNC2=O',
'umaminess'),
('[Na+].[Na+].CN1C=NC2=C(N=CN2C2OC(COP([O-])([O-])=O)C(O)C2O)C1=O',
'umaminess'),
('[Na+].[Na+].CN(C)C1=NC2=C(N=CN2C2OC(COP([O-])([O-])=O)C(O)C2O)C(=O)N1',
'umaminess'),
('[Na+].[Na+].CNC1=NC2=C(N=CN2C2OC(COP([O-])([O-])=O)C(O)C2O)C(=O)N1',
'umaminess'),
('[Na+].[Na+].CSC1=NC2=C(N=CN2C2OC(COP([O-])([O-])=O)C(O)C2O)C(=O)N1C',
'umaminess'),
```

('[Na+].[Na+].CN1C(=NC2=C(N=CN2C2OC(COP([O-])([O-])=O)C(O)C2O)C1=O)N',
'umaminess'), ('[Na+].[Na+].[Na+].[Na+].NC1=NC2=C(N=CN2C2OC(COP([O-])([O-])=O)C(
O)C2O)C(=O)N1.OC1C(O)C(OC1COP([O-])([O-])=O)N1C=NC2=C1N=CNC2=O', 'umaminess'),
('CCOC(=O)CCCNC(=O)OC1CC(C)CCC1C(C)C', 'umaminess'),
('NC(CCC(O)=O)C(=O)NC(CC(O)=O)C(=O)NC(CCC(O)=O)C(O)=O', 'umaminess'),
('NC1=NC2=C(N=CN2C2OC(COP(O)(O)=O)C(O)C2O)C(=O)N1', 'umaminess'),
('[Na+].[Na+].NC1=NC2=C(N=CN2C2OC(COP([O-])([O-])=O)C(O)C2O)C(=O)N1',
'umaminess'), ('OC1C(O)C(OC1COP(O)(O)=O)N1C=NC2=C1N=CNC2=O', 'umaminess'),
('NC(C(O)=O)C1=CC(=O)NO1', 'umaminess'), ('CCNC(=O)CCC(N)C(O)=O', 'umaminess'),
('NC(C1CC(=O)NO1)C(O)=O', 'umaminess'),
('[Mg+2].NC(CCC(O)=O)C([O-])=O.NC(CCC(O)=O)C([O-])=O', 'umaminess'),
('COC(=O)c1ccc(cc1)C(=O)NC1CC(C)CCC1C(C)C', 'umaminess'),
('[NH4+].NC(CCC(O)=O)C([O-])=O', 'umaminess'), ('[K+].NC(CCC(O)=O)C([O-])=O',
'umaminess'), ('[Na+].NC(CCS([O-])(=O)=O)C(O)=O', 'umaminess'),
('[Na+].NC(C(O)CC(O)=O)C([O-])=O', 'umaminess'), ('[Na+].NC(CC(O)=O)C([O-])=O',
'umaminess'), ('[Na+].NC(CCC(O)=O)C([O-])=O', 'umaminess'),
('[Na+].NC(C([O-])=O)C1=CC(=O)NO1', 'umaminess'),
('[Na+].NC(C1CC(=NO1)O)C([O-])=O', 'umaminess'),
('[Na+].NC(CCCC([O-])=O)C(O)=O', 'umaminess'),
('OC1COC(O)(CNC(CCC(O)=O)C(O)=O)C(O)C1O', 'umaminess'),
('OC1COC(O)(CN2C(CCC2=O)C(O)=O)C(O)C1O', 'umaminess'),
('COc1cc(ccc1O)CNC(=O)CCCCO', 'umaminess'), ('CC(O)C(O)C(O)C(=O)NCCc1ccc(cc1)O',
'umaminess'), ('OCC(O)C(O)C(O)C(O)C(=O)NCCc1ccc(cc1)O', 'umaminess'),
('OC(=O)CCC(=O)NCCc1ccc(cc1)O', 'umaminess'), ('CCCC(CCC)NC(=O)c1ccc2c(c1)OCO2',
'umaminess'),
('OCC(O)C(O)CC(NC1=NC2=C(N=CN2C2OC(COP(O)(O)=O)C(O)C2O)C(=O)N1)C(O)=O',
'umaminess'), ('OCCC(NC1=NC2=C(N=CN2C2OC(COP(O)(O)=O)C(O)C2O)C(=O)N1)C(O)=O',
'umaminess'), ('CC(NC1=NC2=C(N=CN2C2OC(COP(O)(O)=O)C(O)C2O)C(=O)N1)C(O)=O',
'umaminess'), ('CSCCNC1=NC2=C(N=CN2C2OC(COP(O)(O)=O)C(O)C2O)C(=O)N1',
'umaminess'), ('CCCSCNC1=NC2=C(N=CN2C2OC(COP(O)(O)=O)C(O)C2O)C(=O)N1',
'umaminess'), ('COc1ccc(c(c1)OC)CNC(=O)C(=O)NCCc1ccccn1', 'umaminess'),
('CCCCNC1=NC2=C(N=CN2C2OC(COP(O)(O)=O)C(O)C2O)C(=O)N1', 'umaminess'),
('CC(O)C(=O)NC1=NC2=C(N=CN2C2OC(COP(O)(O)=O)C(O)C2O)C(=O)N1', 'umaminess'),
('CC(C)C1CCC(C)C2C1C2C(=O)NC1CCCC1', 'umaminess'),
('OCC(O)C(O)C(O)C(O)C(=O)NCCOP(O)(O)=O', 'umaminess'),
('CC(O)CCC(=O)NCCc1ccccc1', 'umaminess'), ('OC(=O)C1CCCN1C(=O)C1CCC(=O)N1',
'umaminess'), ('OC(=O)CCC(NC(=O)C1CCCN1C(=O)C1CCC(=O)N1)C(O)=O', 'umaminess'),
('OCC(NC(=O)C1CCCN1C(=O)C1CCC(=O)N1)C(O)=O', 'umaminess'),
('OC1CC(O)(CC(OC(=O)c2cc(c(c(c2)O)O)O)C1O)C(O)=O', 'umaminess'),
('CC(O)C(N)C(=O)NC(CCC(O)=O)C(O)=O', 'umaminess'),
('[Na+].[Na+].OC1C(O)C(OC1COP([O-])([O-])=O)N1C=NC2=C1NC(=O)NC2=O',
'umaminess'), ('OC1C(O)C(OC1COP(O)(O)=O)N1C=NC2=C1NC(=O)NC2=O', 'umaminess'),
('CSCCC(NC(=O)CCC(N)C(O)=O)C(O)=O', 'umaminess'),
('NC(CCC(=O)NC(CS)C(=O)NCCC(O)=O)C(O)=O', 'umaminess'),
('[Na+].[O-]S(=O)(=O)Nc1ccc(c(c1)Cl)N1CCOCC1', 'sourness'),
('COc1ccc(cc1O)CCC(=O)c1ccccc1', 'sourness'), ('CC(O)=O', 'sourness'),
('OC(=O)CCCCC(O)=O', 'sourness'), ('OCC(O)C1OC(=O)C(=C1O)O', 'sourness'),

('OC(=O)c1ccccc1', 'sourness'), ('OC(O)=O', 'sourness'),
('OC(=O)CC(O)(CC(O)=O)C(O)=O', 'sourness'), ('OC=O', 'sourness'),
('OC(=O)C=CC(O)=O', 'sourness'), ('CC(O)C(O)=O', 'sourness'),
('OC(CC(O)=O)C(O)=O', 'sourness'),
('[Na+].[Na+].CCCCCC(C)N(S([O-])(=O)=O)S([O-])(=O)=O', 'sourness'),
('[Na+].[Na+].CCCCCCCC(C)N(S([O-])(=O)=O)S([O-])(=O)=O', 'sourness'),
('[Na+].[Na+].COC(CN(S([O-])(=O)=O)S([O-])(=O)=O)OC', 'sourness'),
('[Na+].[Na+].COCCN(S([O-])(=O)=O)S([O-])(=O)=O', 'sourness'),
('[Na+].[Na+].CCOCCCN(S([O-])(=O)=O)S([O-])(=O)=O', 'sourness'),
('[Na+].[Na+].[Na+].CN(CCN(S([O-])(=O)=O)S([O-])(=O)=O)S([O-])(=O)=O',
'sourness'), ('[Na+].[Na+].CC(C)CN(S([O-])(=O)=O)S([O-])(=O)=O', 'sourness'),
('[Na+].[Na+].CC(C)N(S([O-])(=O)=O)S([O-])(=O)=O', 'sourness'),
('[Na+].[Na+].CCCN(S([O-])(=O)=O)S([O-])(=O)=O', 'sourness'), ('OC(=O)C(O)=O',
'sourness'), ('OP(O)(O)=O', 'sourness'), ('CCC(O)=O', 'sourness'),
('[Na+].[O-]S(=O)(=O)Nc1cccc(c1)S([O-])(=O)=O', 'sourness'),
('[Na+].[O-]S(=O)(=O)NC1=Nc2ccccc2S1', 'sourness'),
('[Na+].CC(=O)c1ccccc1NS([O-])(=O)=O', 'sourness'),
('[Na+].Cc1ccc(c(c1)Cl)NS([O-])(=O)=O', 'sourness'),
('[Na+].[O-]S(=O)(=O)Nc1ccc(cc1F)I', 'sourness'),
('[Na+].COc1ccc(cc1NS([O-])(=O)=O)C', 'sourness'),
('[Na+].CCCC1=C(C)N=C(NS([O-])(=O)=O)S1', 'sourness'),
('[Na+].[O-]S(=O)(=O)NC1=NN=C(Cc2ccccc2)S1', 'sourness'),
('[Na+].Cc1ccc(c(c1)NS([O-])(=O)=O)N(=O)=O', 'sourness'),
('[Na+].CC(C)(C)C1=CC(=NO1)NS([O-])(=O)=O', 'sourness'),
('[Na+].CCC(C)c1ccc(cc1)NS([O-])(=O)=O', 'sourness'), ('OC(=O)CCC(O)=O',
'sourness'), ('OC(C(O)C(O)=O)C(O)=O', 'sourness'), ('NCCCC(O)=O', 'sourness'),
('[NH4+].[Cl-]', 'saltiness'),
('[Na+].[Na+].CCCCCN(S([O-])(=O)=O)S([O-])(=O)=O', 'saltiness'), ('[Li+].[Cl-]',
'saltiness'), ('NCCCC(N)C(=O)NCCS(O)(=O)=O', 'saltiness'),
('NCCCC(N)C(=O)NCCCC(O)=O', 'saltiness'), ('Cl.NCCCCC(N)C(=O)NCCS(O)(=O)=O',
'saltiness'), ('[Na+].[Na+].CCC(C)CCCCN(S([O-])(=O)=O)S([O-])(=O)=O',
'saltiness'), ('Cl.NCCCC(N)C(=O)NCCS(O)(=O)=O', 'saltiness'),
('Cl.NCCCC(N)C(=O)NCCC(O)=O', 'saltiness'), ('Cl.NCCCC(N)C(=O)NCCCC(O)=O',
'saltiness'), ('[Cl-].[K+]', 'saltiness'), ('[Na+].[Cl-]', 'saltiness'),
('COc1ccc(cc1O)C1CC(=O)c2c(cc(cc2O1)O)O', 'multitaste'),
('CCCCCC(O)C=CC=CCCCCCCCC(O)=O', 'multitaste'), ('NC1(CCCCC1)C(O)=O',
'multitaste'), ('NC1(CCCCCCC1)C(O)=O', 'multitaste'), ('NC1(CCCC1)C(O)=O',
'multitaste'), ('ON=CC1=CCCCCC1', 'multitaste'), ('OCCC1=CNc2ccccc12',
'multitaste'), ('CC(C)C(C(O)=O)S(=O)(=O)c1ccc(cc1)Br', 'multitaste'),
('CCCCCCCCCCC=CC=O', 'multitaste'), ('CC(CC(O)=O)S(=O)(=O)c1ccc(cc1)C',
'multitaste'), ('CC(=O)OCCCc1ccccc1', 'multitaste'), ('CC(C)C(=O)OCCCc1ccccc1',
'multitaste'), ('Clc1ccc2c(c1)C(=O)NS2(=O)=O', 'multitaste'),
('Brc1ccc2c(c1)S(=O)(=O)NC2=O', 'multitaste'), ('Clc1ccc2c(c1)S(=O)(=O)NC2=O',
'multitaste'), ('CC(C)=CCCC(C)=O', 'multitaste'), ('Clc1cccc2c1S(=O)(=O)NC2=O',
'multitaste'), ('CCCCCC=CC=CC(O)CCCCCCCC(O)=O', 'multitaste'), ('C=CCN=C=S',
'multitaste'), ('OC(=O)CC(NC(=O)Nc1ccc(cc1)C#N)C1CCCCCCC1', 'multitaste'),
('OC(=O)CC(NC(=O)Nc1ccc(cc1)C#N)C1CCCCC1', 'multitaste'),

('OC(=O)CC1(CCCCC1)NC(=O)Nc1ccc(cc1)C#N', 'multitaste'), ('[Cl-].[Cl-].[Ba+2]', 'multitaste'), ('[Na+].[O-]S(=O)(=O)Nc1ccc2c(c1)SC(=N2)S', 'multitaste'), ('CC(=O)OCc1ccccc1', 'multitaste'), ('CC(C)C(C(O)=O)S(=O)(=O)c1ccc(cc1)Cl', 'multitaste'), ('CCCCNS(=O)(=O)OCC', 'multitaste'), ('CCCCNS(=O)(=O)OCCC', 'multitaste'), ('[Cl-].[Cl-].[Ca+2]', 'multitaste'), ('[Ca+2].Oc1ccc(cc1)S([O-])(=O)=O.Oc1ccc(cc1)S([O-])(=O)=O', 'multitaste'), ('[Na+].CC1=NN=C(SCC2=C(N3C(SC2)C(NC(=O)CN2C=NN=N2)C3=O)C([O-])=O)S1', 'multitaste'), ('O=COCC=Cc1ccccc1', 'multitaste'), ('CC(C)=CCCC(C)=CC=O', 'multitaste'), ('CCC(=O)OCCC(C)CCC=C(C)C', 'multitaste'), ('O=C1Oc2ccccc2C=C1', 'multitaste'), ('CCCCOS(=O)(=O)NC1CCCCC1', 'multitaste'), ('O=S(=O)(NC1CCCCC1)OC1CCCCC1', 'multitaste'), ('CCOS(=O)(=O)NC1CCCCC1', 'multitaste'), ('CC(C)COS(=O)(=O)NC1CCCCC1', 'multitaste'), ('CC(C)OS(=O)(=O)NC1CCCCC1', 'multitaste'), ('COS(=O)(=O)NC1CCCCC1', 'multitaste'), ('CCCOS(=O)(=O)NC1CCCCC1', 'multitaste'), ('CCC(C)OS(=O)(=O)NC1CCCCC1', 'multitaste'), ('NC(CS)C(O)=O', 'multitaste'), ('NC(CCC(O)=O)C(O)=O', 'multitaste'), ('CCOC(=O)Cc1ccccc1', 'multitaste'), ('OC(=O)c1cc(c(c(c1)O)O)O', 'multitaste'), ('CC(CCC1=C(CO)CC(OC2OCC(O)C(O)C2O)C2C(C)(CO)CCCC12C)=CCO', 'multitaste'), ('CC(CCC1C(C)(O)CC(OC2OCC(O)C(O)C2O)C2C(C)(C)CC(O)CC12C)=CCO', 'multitaste'), ('OCC1OC(Oc2cc(c3c(c2)OC=C(C3=O)c2ccc(cc2)O)O)C(O)C(O)C1O', 'multitaste'), ('N#CCCCC#N', 'multitaste'), ('NCCCC(NC(=O)CN)C(O)=O', 'multitaste'), ('CCCCCCOC(C)=O', 'multitaste'), ('C#N', 'multitaste'), ('CC(C)CCOC(C)=O', 'multitaste'), ('CCC(=O)OCCC(C)C', 'multitaste'), ('CC(C)CCOC(=O)c1ccccc1O', 'multitaste'), ('CCCOS(=O)(=O)NCC(C)C', 'multitaste'), ('CCC(=O)OC(C)C', 'multitaste'), ('COC(=O)CCC(N)C(=O)OC', 'multitaste'), ('CC(C)=CCCC(C)(OC=O)C=C', 'multitaste'), ('CCCCCC=CCC=CCCCCCCCC(O)=O', 'miscellaneous'), ('CC(C)C1CCC(C)CC1O', 'multitaste'), ('NCCCC(N)C(=O)NCC(O)=O', 'multitaste'), ('NCCCC(N)C(=O)NCCC(O)=O', 'multitaste'), ('N.N.[Mg+2].OS([O-])(=O)=O.OS([O-])(=O)=O', 'multitaste'), ('COCC1OC(CO)C(O)C(O)C1O', 'multitaste'), ('[Na+].[Na+].CC(C)CCCC(C)N(S([O-])(=O)=O)S([O-])(=O)=O', 'multitaste'), ('CCCCCCCCC=CCCCCCCCC(O)=O', 'miscellaneous'), ('O=COCCc1ccccc1', 'multitaste'), ('CC(C)C(=O)OCCc1ccccc1', 'multitaste'), ('CC(C)CC(=O)OCCc1ccccc1', 'multitaste'), ('O=Cc1ccc2c(c1)OCO2', 'multitaste'), ('O=C1NS(=O)(=O)c2cccc(c12)N(=O)=O', 'multitaste'), ('[K+].OC(=O)C([O-])=O', 'multitaste'), ('CCCOC(C)=O', 'multitaste'), ('CCCOC=O', 'multitaste'), ('CCCOC(=O)CC(C)C', 'multitaste'), ('CCCNS(=O)(=O)OCCC', 'multitaste'), ('OC(=O)c1ccc(c(c1)O)O', 'multitaste'), ('CCC1=C(CC(NC(=O)C2CCCN2C(=O)C(CC(N)=O)NC(=O)C(NC(=O)C(CC(N)=O)NC(=O)C2CCCN2C(=O)CNC(=O)C(Cc2ccccc2)NC(=O)C(CC(C)C)NC(=O)C(N)CCC(N)=O)C(C)C)C(O)=O)c2ccccc2N1', 'multitaste'), ('CC(C)CC(NC(=O)C(N)CCC(N)=O)C(=O)NC(Cc1ccccc1)C(=O)NC(CC(N)=O)C(=O)N1CCCC1C(=O)NC(CO)C(=O)NC(C(C)O)C(=O)NC(CC(N)=O)C(=O)N1CCCC1C(=O)NC(CC1=CNc2ccccc12)C(O)=O', 'multitaste'), ('CCOC(=O)C(CC1=CNc2ccccc12)NC(=O)C1CCCN1C(=O)C(CC(N)=O)NC(=O)C(NC(=O)C(CO)NC(=O)C1CCCN1C(=O)C(CC(N)=O)NC(=O)C(Cc1ccccc1)NC(=O)C(CC(C)C)NC(=O)C(N)CCC(N)=O)C(C)O', 'multitaste'), ('CC(CCCC(C)=C)CCOC=O', 'multitaste'), ('CC(C)CC(=O)OCCC(C)CCCC(C)=C', 'multitaste'), ('OC(=O)c1ccccc1O', 'multitaste'),

('CC(=O)OCC(C)=CCCC1(C)C2CC3C(C2)C13.CC(=O)OCC(C)=CCCC1(C)C2CCC(C2)C1=C', 'multitaste'), ('[Na+].Cc1c(cccc1NS([O-])(=O)=O)Cl', 'multitaste'), ('[Na+].NC(CC([O-])=O)C(O)=O', 'multitaste'), ('[Na+].COc1ccc(c(c1)OC)NS([O-])(=O)=O', 'multitaste'), ('[Na+].CCc1cccc(c1NS([O-])(=O)=O)C', 'multitaste'), ('[Na+].COc1cccc(c1NS([O-])(=O)=O)C', 'multitaste'), ('[Na+].Cc1cnc(c(c1)Br)NS([O-])(=O)=O', 'multitaste'), ('[Na+].COC(=O)c1cccc(c1)NS([O-])(=O)=O', 'multitaste'), ('[Na+].[O-]S(=O)(=O)Nc1ccc(cc1F)Cl', 'multitaste'), ('[Na+].Cc1cc(ccc1NS([O-])(=O)=O)Cl', 'multitaste'), ('[Na+].[O-]S(=O)(=O)Nc1ccc(c(c1)N(=O)=O)Cl', 'multitaste'), ('[Na+].COc1ccc(c(c1)N(=O)=O)NS([O-])(=O)=O', 'multitaste'), ('[Na+].Cc1ccc(cc1N(=O)=O)NS([O-])(=O)=O', 'multitaste'), ('[Na+].CC(C)(C)OC(=O)c1cccc(c1)NS([O-])(=O)=O', 'multitaste'), ('[Na+].CC(C)C1=CSC(=N1)NS([O-])(=O)=O', 'multitaste'), ('[Br-].[Br-].[Sr+2]', 'multitaste'), ('COc1ccc(cc1)C=NO', 'multitaste'), ('CC=C(C)CC=NO', 'multitaste'), ('CCC=C(C)CC=NO', 'multitaste'), ('NCCCC(NC(=O)CCN)C(O)=O', 'multitaste'), ('CCCCCC=CCC=CCC=CCCCCC(O)=O', 'multitaste'), ('COc1ccc(cc1O)C(=O)Sc1ccccc1', 'tastelessness'), ('NC(=O)Nc1ccc(cc1)N(=O)=O', 'tastelessness'), ('OC(=O)CC(NC(=O)Nc1ccc(cc1)CN(=O)=O)c1ccccc1', 'tastelessness'), ('NC(=O)Nc1ccc(cc1)CCc1ccccc1', 'tastelessness'), ('COc1cc(ccc1O)CCC(=O)c1c(cccc1O)O', 'tastelessness'), ('Oc1ccc(cc1)CCC(=O)c1c(cccc1O)O', 'tastelessness'), ('O=C(NCCC#N)Nc1ccc(cc1)N(=O)=O', 'tastelessness'), ('COc1ccc(cc1O)C(=O)Cc1ccccc1', 'tastelessness'), ('CC1OC(OC2C(O)C(O)C(CO)OC2Oc2cc(c(c(c2)O)C(=O)CCc2ccccc2O)O)C(O)C(O)C1O', 'tastelessness'), ('CC1OC(OC2C(O)C(O)C(CO)OC2Oc2cc(c(c(c2)O)C(=O)CCc2cc(c(c(c2)O
)O)O)O)C(O)C(O)C1O', 'tastelessness'), ('CCOc1cc(ccc1O)CCC(=O)c1c(cc(cc1O)OC1OC(CO)C(O)C(O)C1OC1OC(C)C(O)C(O)C1O)O', 'tastelessness'), ('COc1cc(ccc1O)CCC(=O)c1c(cc(cc1O)OC1OC(CO)C(O)C(O)C1OC1OC(C)C(O)C(O)C1O)O', 'tastelessness'), ('CC1OC(OC2C(O)C(O)C(CO)OC2Oc2cc(c(c(c2)O)C(=O)CCc2ccccc2)O)C(O)C(O)C1O', 'tastelessness'), ('CCCNC(=O)Nc1ccc(cc1)N(=O)=O', 'tastelessness'), ('COc1ccc(cc1O)C1=COc2ccccc2O1', 'tastelessness'), ('[Na+].[Na+].[O-]S(=O)(=O)NC1CCC(CC1)NS([O-])(=O)=O', 'tastelessness'), ('COc1cc2c(cc1O)CC1(O)COc3ccccc3C21', 'tastelessness'), ('CC1C2CC(=O)C3C4(COC(C)=O)C(CCC(C)(C)C4C=O)OC(=O)C3(C2)C1=O', 'tastelessness'), ('CSc1ccc(cc1)C(=O)c1ccccc1C(O)=O', 'tastelessness'), ('BrCCN1C(=O)c2ccccc2S1(=O)=O', 'tastelessness'), ('COc1ccc(cc1OC)C(=O)c1ccccc1C(O)=O', 'tastelessness'), ('COc1ccc(cc1O)C1NC(=O)c2ccccc2S1', 'tastelessness'), ('COc1ccc(cc1O)C1OCc2ccccc2O1', 'tastelessness'), ('CSc1ccc(cc1O)C1CSc2cc(ccc2O1)O', 'tastelessness'), ('CSc1ccc(cc1O)C1CSc2ccccc2O1', 'tastelessness'), ('OC(=O)c1ccc(cc1)C(=O)c1ccccc1C(O)=O', 'tastelessness'), ('COc1ccc(c(c1)O)C(=O)c1ccccc1C(O)=O', 'tastelessness'),

('COc1cc(ccc1O)C(=O)c1ccccc1C(O)=O', 'tastelessness'),
('COc1ccc(cc1)Cc1ccccc1C(O)=O', 'tastelessness'),
('COc1ccc(cc1)Oc1ccccc1C(O)=O', 'tastelessness'), ('COc1ccc(cc1)C1OCc2ccccc2S1', 'tastelessness'), ('COc1ccc(cc1)Sc1ccccc1C(O)=O', 'tastelessness'),
('OC(=O)c1ccccc1C(=O)c1ccc(cc1)Oc1ccccc1', 'tastelessness'),
('CCCOc1ccc(cc1)C(=O)c1ccccc1C(O)=O', 'tastelessness'),
('CCCCCOc1ccc(cc1O)CCc1ccccc1', 'tastelessness'),
('CCOc1ccc(cc1O)CCC(=O)c1c(cccc1O)O', 'tastelessness'),
('COc1ccc(cc1O)CCC(=O)c1c(cccc1O)O', 'tastelessness'),
('CCCOc1ccc(cc1O)CCC(=O)c1c(cccc1O)O', 'tastelessness'),
('COc1ccc(cc1O)CCc1ccccc1C(O)=O', 'tastelessness'),
('COc1ccc(cc1OCC(O)=O)CCc1ccccc1', 'tastelessness'),
('OC(=O)c1ccccc1C(=O)c1ccccc1', 'tastelessness'),
('CCN1C(=O)c2cc(ccc2S1(=O)=O)N(=O)=O', 'tastelessness'),
('CCN1C(=O)c2ccc(cc2S1(=O)=O)N(=O)=O', 'tastelessness'),
('CCN1C(=O)c2cccc(c2S1(=O)=O)N(=O)=O', 'tastelessness'),
('COc1ccc(cc1)C(O)c1ccccc1CO', 'tastelessness'),
('COc1ccc(cc1O)C1Oc2ccccc2OC1C', 'tastelessness'), ('COc1ccc(cc1O)CCCCc1ccccc1', 'tastelessness'), ('COc1ccc(cc1O)CCc1cccc2ccccc12', 'tastelessness'),
('COc1ccc(cc1)CCc1ccc(c(c1)O)OC', 'tastelessness'),
('COc1ccc(cc1O)CCc1ccc(cc1)C', 'tastelessness'),
('COc1ccc(cc1O)CCc1ccccc1C(C)C', 'tastelessness'),
('COc1ccc(cc1O)C1(C)COc2ccccc2O1', 'tastelessness'),
('CN1C(=O)c2ccc(cc2S1(=O)=O)N(=O)=O', 'tastelessness'),
('CCCOc1ccc(cc1N(=O)=O)N', 'tastelessness'), ('COc1cc2c(cc1O)C(COc1ccccc1)OCC2', 'tastelessness'), ('COc1ccc(c(c1O)CC=Cc1ccc(cc1)N(=O)=O)C1OCc2ccccc2S1', 'tastelessness'), ('Oc1ccc(cc1O)C1Cc2ccccc2C(=O)O1', 'tastelessness'),
('COc1ccc(cc1OC)C1Cc2cccc(c2C(=O)O1)OC', 'tastelessness'),
('COc1ccc(cc1O)C1OC(=O)c2ccccc12', 'tastelessness'),
('COc1ccc(cc1O)C1OCc2ccccc2CO1', 'tastelessness'),
('COc1ccc(cc1O)C1NC(=O)c2ccccc2O1', 'tastelessness'),
('COc1ccc(cc1O)CC(C(O)=O)c1ccccc1', 'tastelessness'),
('COc1ccc(cc1O)C1CSc2ccccc2C1', 'tastelessness'),
('COc1ccc(cc1O)C1NC(=O)c2ccccc2N1', 'tastelessness'),
('COc1ccc(cc1O)C1=Cc2ccccc2C(=O)O1', 'tastelessness'),
('COc1ccc(cc1O)C1Cc2ccccc2C(=O)N1', 'tastelessness'),
('Oc1cccc(c1)C1Cc2ccccc2C(=O)O1', 'tastelessness'),
('COc1cc(ccc1O)C1Cc2ccccc2C(=O)O1', 'tastelessness'),
('COc1ccc(cc1)C1OC(=O)c2ccccc12', 'tastelessness'),
('COc1ccc(cc1)C1Cc2ccccc2C(=O)O1', 'tastelessness'),
('COc1ccc2c(c1O)OCC1(O)Cc3cc(c(cc3C21)OC)OC', 'tastelessness'),
('COc1ccc(cc1O)CC1OC(=O)c2ccccc12', 'tastelessness'),
('OC(=O)CCNNC(=O)c1ccc(cc1)N(=O)=O', 'tastelessness'),
('OC(=O)CCNC(=O)Nc1ccccc1', 'tastelessness'),
('COc1ccc(cc1O)CCc1cccc(c1)C(O)=O', 'tastelessness'),
('COc1ccccc1C(CC(O)=O)NC(=O)Nc1ccc(cc1)C#N', 'tastelessness'),
('OC(=O)C=CNC(=O)Nc1ccc(cc1)N(=O)=O', 'tastelessness'),

('OC(=O)CCOC(=O)Nc1ccc(cc1)N(=O)=O', 'tastelessness'),
('CCCC(C)NC(=O)C(N)CC(O)=O', 'tastelessness'),
('[O-]S(=O)(=O)C1=CC2=C(N=C(N2Nc2ccccc2)c2ccccc2)c2ccccc12', 'tastelessness'),
('CCCCCCCC(C)NC(=O)C(N)CC(O)=O', 'tastelessness'),
('CCCCC(NC(=O)C(N)CC(O)=O)C(=O)OC(C)C', 'tastelessness'),
('COc1ccc(cc1O)C(=O)Nc1ccccc1', 'tastelessness'),
('COc1ccc(cc1O)C1CC(=O)c2ccccc2O1', 'tastelessness'),
('COc1ccc(cc1O)C=Cc1ccccc1', 'tastelessness'), ('COc1ccc(cc1O)CCc1cccc(c1)O', 'tastelessness'), ('Cc1c(cccc1N(=O)=O)N', 'tastelessness'),
('Cc1ccc(cc1N(=O)=O)N', 'tastelessness'), ('NC(=O)NCCC(O)=O', 'tastelessness'),
('COc1cc(ccc1O)C1OCCc2ccccc2O1', 'tastelessness'),
('COc1ccc(cc1)C(=O)C(C)=CC(O)=O', 'tastelessness'),
('COc1cc(ccc1O)C1CC(=O)c2c(cccc2O1)O', 'tastelessness'),
('COc1cc2c(cc1OC)C1c3ccc(c(c3OCC1(O)C2)OC)O', 'tastelessness'),
('OC(=O)CCCNC(=O)c1ccc(cc1)N(=O)=O', 'tastelessness'),
('OC(=O)CCCNC(=O)Nc1ccc(cc1)N(=O)=O', 'tastelessness'),
('COc1ccc(cc1O)CCc1ccc(cc1)C(O)=O', 'tastelessness'),
('OC(=O)CCC(=O)NNc1ccc(cc1)N(=O)=O', 'tastelessness'),
('OC(=O)CCNC(=O)Nc1ccc(cc1)C(O)=O', 'tastelessness'),
('COc1ccc(cc1)C(=O)c1ccccc1', 'tastelessness'),
('CC(C)C1C(=O)OCC1(CC(O)=O)C1CCC(CC1C(O)=O)C(C)C(O)=O', 'tastelessness'),
('COc1cc(c(cc1O)C1OCc2ccccc2S1)N(=O)=O', 'tastelessness'),
('CCNC(=O)Oc1cc(ccc1OC)C1OCc2ccccc2S1', 'tastelessness'),
('CCCOc1ccc(cc1O)CCc1ccccc1', 'tastelessness'),
('COc1ccc(cc1O)C1SCc2cc(c(cc2CS1)C)C', 'tastelessness'),
('Oc1ccc(cc1)C1CC(=O)c2c(cccc2O1)O', 'tastelessness'),
('COc1ccc(cc1O)CCc1ccc(cc1)Br', 'tastelessness'),
('COc1ccc(cc1O)CCc1ccc(cc1)Cl', 'tastelessness'),
('COc1ccc(cc1O)CCc1ccc(c(c1)OC)O', 'tastelessness'),
('COc1ccc(cc1O)C1OCc2ccccc2C1=NO', 'tastelessness'),
('COc1ccc(cc1O)CCc1cccc(c1)CO', 'tastelessness'),
('COc1ccc(cc1O)CCc1ccc(cc1)CO', 'tastelessness'), ('Nc1ccc2c(c1)C(=O)NS2(=O)=O', 'tastelessness'), ('COc1ccc(cc1O)Cc1ccccc1', 'tastelessness'),
('COc1ccc(cc1O)C1=CC(=O)c2c(cccc2O1)O', 'tastelessness'),
('OCC1SC(O)(CO)C(O)C1O', 'tastelessness'),
('OCC1(OC(CCl)C(O)C1O)OC1OC(CCl)C(O)C(O)C1O', 'tastelessness'),
('COC1OC(CCl)C(O)C(O)C1O', 'tastelessness'), ('CCOc1ccc2c(c1)S(=O)(=O)NC2=O', 'tastelessness'), ('COc1ccc2c(c1)S(=O)(=O)NC2=O', 'tastelessness'),
('CC1C2CCC3C45COC(=O)C5C(C)(C)C(=O)CC4OC(=O)C3(C2)C1=O', 'tastelessness'),
('CC1C2CCC(C(C2)C1=O)C12CCC(=O)C(C)(C)C2C(=O)OC1', 'tastelessness'),
('COc1ccc(cc1O)C1SCc2ccccc2CS1', 'tastelessness'),
('COc1ccc(cc1O)C1=Cc2cccc(c2C(=O)O1)O', 'tastelessness'),
('COc1cc2c(cc1O)C1COc3ccccc3C1O2', 'tastelessness'),
('CC(=O)OCC12C(CCC(C)(C)C1C=O)OC(=O)C13CC(CC(OC(C)=O)C21)C(=C)C3=O', 'tastelessness'), ('CC(N)C(=O)NC1C(C)(C)SC1(C)C', 'tastelessness'),
('NC(CC(O)=O)C(=O)NC(Cc1ccccc1)C(=O)OC1CCCCC1', 'tastelessness'),
('CC(C)CCCNC(=O)C(N)CC(O)=O', 'tastelessness'),

('CC1C2CC(OC(C)=O)C3C45COC=C5C(C)(C)CCC4OC(=O)C3(C2)C1=O', 'tastelessness'),
('COC(=O)C(C)C1CC(O)C2C(C1)C(=O)OC1CCC(C)(C)C(CO)C21CO', 'tastelessness'),
('CC1C2CC(=O)C3C45COC(=O)C5C(C)(C)CCC4OC(=O)C3(C2)C1=O', 'tastelessness'),
('CC1C2CC(O)C3C45COCC5C(C)(C)CCC4OC(=O)C3(C2)C1=O', 'tastelessness'),
('OC1CC(O)C(O)CC1O', 'tastelessness'), ('NC(CSSCC(N)C(O)=O)C(O)=O',
'tastelessness'),
('CC1C2CC(OC(C)=O)C3C4(COC(C)=O)C(CCC(C)(C)C4C=O)OC(=O)C3(C2)C1=O',
'tastelessness'), ('CC(C)(Cl)C1CCC(=CC1)C=NO', 'tastelessness'),
('CC(C)(C)C(N)C(O)=O', 'tastelessness'),
('CC1C2CCC3C45COC(=O)C5C(C)(C)C(O)CC4OC(=O)C3(C2)C1=O', 'tastelessness'),
('CC1OC(OCC2OC(Oc3cc(c4c(c3)OC(CC4=O)c3ccc(c(c3)O)O)O)C(O)C(O)C2O)C(O)C(O)C1O',
'tastelessness'), ('CC(CCC1=C(CO)CC(OC2OCC(O)C(O)C2O)C2C(C)(CO)CC(O)CC12C)=CCO',
'tastelessness'), ('COc1ccccc1O', 'tastelessness'),
('NC(CC(O)=O)C(=O)NCCc1ccccc1', 'tastelessness'),
('COc1ccc(cc1O)C1CC(=O)c2c(cc(cc2O1)OC1OC(COC2OC(C)C(O)C(O)C2O)C(O)C(O)C1O)O',
'tastelessness'), ('Oc1ccc(cc1)C1Cc2cccc(c2C(=O)O1)O', 'tastelessness'),
('COc1ccc(cc1)C1CC(=O)c2c(cc(cc2O1)OC1OC(COC2OC(C)C(O)C(O)C2O)C(O)C(O)C1O)O',
'tastelessness'), ('NC(CC(O)=O)C(O)=O', 'miscellaneous'),
('NC(CC(O)=O)C(=O)NC(Cc1ccccc1)C(O)=O', 'tastelessness'),
('COC(=O)c1ccccc1C(=O)c1ccc(cc1)OC', 'tastelessness'),
('COC(=O)CCNC(=O)Nc1ccc(cc1)N(=O)=O', 'tastelessness'),
('COC(=O)c1cccc(c1)CCc1ccc(c(c1)O)OC', 'tastelessness'),
('COC(=O)c1ccc(cc1)CCc1ccc(c(c1)O)OC', 'tastelessness'),
('CC(=O)OC1OCC23C(CCC(C)(C)C12)OC(=O)C12CC(CC(O)C31)C(=C)C2=O',
'tastelessness'), ('COc1ccc(cc1O)CNCC(C)C', 'tastelessness'),
('CCCNCc1ccc(c(c1)O)OC', 'tastelessness'), ('COc1ccc(cc1O)CNCC(C)(C)C',
'tastelessness'), ('[Na+].[Na+].CC(C)OCCCN(S([O-])(=O)=O)S([O-])(=O)=O',
'tastelessness'), ('[Na+].[Na+].[O-]S(=O)(=O)N(CCCN1CCOCC1)S([O-])(=O)=O',
'tastelessness'), ('COc1ccc(c(c1O)NC(=O)c1ccc(cc1)N(=O)=O)C1OCc2ccccc2S1',
'tastelessness'), ('COCCC1CCC(=CC1)C=NO', 'tastelessness'),
('CC1OC(OCC2OC(Oc3cc(c4c(c3)OC(CC4=O)c3ccc(cc3)O)O)C(O)C(O)C2O)C(O)C(O)C1O',
'tastelessness'), ('CCN1C(=O)c2ccccc2S1(=O)=O', 'tastelessness'),
('CN1C(=O)c2ccccc2S1(=O)=O', 'tastelessness'),
('OC(=O)c1ccccc1C(=O)c1ccc(cc1)-c1ccccc1', 'tastelessness'),
('CCOc1ccccc1NC(N)=O', 'tastelessness'), ('Cc1ccccc1NC(N)=O', 'tastelessness'),
('COc1ccc(cc1O)CCc1ccc2ccccc2c1', 'tastelessness'),
('COc1ccc(cc1O)CCc1ccc(c(c1)O)OC', 'tastelessness'),
('COc1ccc(cc1O)CCc1ccc(c(c1)OC)OC', 'tastelessness'), ('NC(NCC(O)=O)=Nc1ccccc1',
'tastelessness'), ('[Na+].[O-]C(=O)c1cccc(c1)NS([O-])(=O)=O', 'tastelessness'),
('[Na+].[O-]C(=O)c1ccc(cc1)NS([O-])(=O)=O', 'tastelessness'),
('[Na+].[O-]S(=O)(=O)Nc1ccccc1Oc1ccccc1', 'tastelessness'),
('[Na+].CCOC(=O)c1ccc(cc1)NS([O-])(=O)=O', 'tastelessness'),
('[Na+].CCCOC(=O)c1ccc(cc1)NS([O-])(=O)=O', 'tastelessness'),
('[Na+].CSc1cccc(c1)NS([O-])(=O)=O', 'tastelessness'),
('[Na+].OCCc1ccc(cc1)NS([O-])(=O)=O', 'tastelessness'),
('[Na+].CCN(CCO)c1ccc(cc1)NS([O-])(=O)=O', 'tastelessness'),
('COc1ccc2c(c1O)CCC12SCc2ccccc2S1', 'tastelessness'),

```
('COc1cc2c(cc1O)CCC12SCc2ccccc2S1', 'tastelessness'),
('[O-]S(=O)(=O)OC(=O)CCNC(=O)Nc1ccc(cc1)N(=O)=O', 'tastelessness'),
('OCC1=CC=C(O1)C=NO', 'tastelessness'), ('CCC(C)C(=O)OC(C)C1(O)CCC2(O)C3CCC4CC(C
CC4(C)C3CC(OC(C)=O)C12C)OC1CC(OC)C(OC2CC(OC)C(OC3OC(C)C(O)C(OC)C3O)C(C)O2)C(C)O1
', 'tastelessness'), ('CCC(C)C(=O)OC(C)C1(O)CCC2(O)C3CCC4CC(CCC4(C)C3CC(OC(C)=O)
C12C)OC1CC(OC)C(OC2CC(OC)C(OC3OC(C)C(OC4OC(CO)C(OC5OC(CO)C(O)C(O)C5O)C(O)C4O)C(O
C)C3O)C(C)O2)C(C)O1', 'tastelessness'), ('CCC(C)C(=O)OC(C)C1(O)CCC2(O)C3CCC4CC(C
CC4(C)C3CC(OC(C)=O)C12C)OC1CC(OC)C(OC2CC(OC)C(OC3CC(OC)C(O)C(C)O3)C(C)O2)C(C)O1'
, 'tastelessness'), ('CCC(C)C(=O)OC(C)C1(O)CCC2(O)C3CCC4CC(CCC4(C)C3CC(OC(C)=O)C
12C)OC1CC(OC)C(OC2CC(OC)C(OC3CC(OC)C(OC4OC(CO)C(OC5OC(CO)C(O)C(O)C5O)C(O)C4O)C(C
)O3)C(C)O2)C(C)O1', 'tastelessness'), ('CCC(C)C(=O)OC(C)C1(O)CCC2(O)C3CCC4CC(CCC
4(C)C3CC(OC(C)=O)C12C)OC1CC(O)C(OC2CC(OC)C(OC3CC(OC)C(O)C(C)O3)C(C)O2)C(C)O1',
'tastelessness'), ('CCC(C)C(=O)OC(C)C1(O)CCC2(O)C3CCC4CC(CCC4(C)C3CC(OC(C)=O)C12
C)OC1CC(O)C(OC2CC(OC)C(OC3CC(OC)C(OC4CC(OC)C(O)C(C)O4)C(C)O3)C(C)O2)C(C)O1',
'tastelessness'), ('Oc1ccc(cc1O)C1Cc2cccc(c2C(=O)O1)O', 'tastelessness'),
('CC(=O)OC1C2C(C)(C)CCC(O)C32COC1(O)C12CC(CCC31)C(=C)C2OC(C)=O',
'tastelessness'), ('COc1ccc(cc1N)CCc1ccccc1', 'tastelessness'),
('COc1cc(ccc1O)CCc1ccccc1', 'tastelessness'),
('CC(NC(=O)CC(N)C(O)=O)C(=O)NC1C(C)(C)SC1(C)C', 'tastelessness'),
('[Na+].[O-]S(=O)(=O)Nc1ccc(cc1)Cl', 'non-sweetness'),
('[Na+].[O-]S(=O)(=O)Nc1ccc(cc1)F', 'non-sweetness'),
('[Na+].[O-]S(=O)(=O)Nc1ccc(cc1)N(=O)=O', 'non-sweetness'),
('OC(=O)CS(=O)(=O)c1ccccc1', 'non-sweetness'),
('COc1ccc(cc1)S(=O)(=O)C1(CCCC1)C(O)=O', 'non-sweetness'),
('COc1ccc(cc1)S(=O)(=O)C1(CCCCC1)C(O)=O', 'non-sweetness'),
('Cc1ccc(cc1)S(=O)(=O)C1(CCCC1)C(O)=O', 'non-sweetness'),
('OC(=O)C1(CCCC1)S(=O)(=O)c1ccccc1', 'non-sweetness'),
('OC(=O)C1(CC1)S(=O)(=O)c1ccccc1', 'non-sweetness'), ('OC1COCC1O', 'non-
sweetness'), ('CC(=O)OCC1OC(COC(C)=O)(OC2OC(CO)C(O)C(O)C2O)C(O)C1O', 'non-
sweetness'), ('CC(=O)OC(C)(CCC=C(C)C)C1CCC(=CC1=O)C', 'non-sweetness'),
('CC1CC(C)C(CC1C)(C(O)=O)S(=O)(=O)c1ccccc1', 'non-sweetness'),
('CC1CCC(C(C)C1)(C(O)=O)S(=O)(=O)c1ccccc1', 'non-sweetness'),
('CC1CCCCC1(C(O)=O)S(=O)(=O)c1ccccc1', 'non-sweetness'),
('OC(=O)C1(CCC=CC1)S(=O)(=O)c1ccccc1', 'non-sweetness'),
('CC1CCC(CC1)(C(O)=O)S(=O)(=O)c1ccccc1', 'non-sweetness'),
('OC(=O)C1(CCC1)S(=O)(=O)c1ccccc1', 'non-sweetness'),
('OC(=O)C1(CCCCC1)S(=O)(=O)c1ccccc1', 'non-sweetness'),
('CC(C)C(C(O)=O)S(=O)(=O)c1ccc(cc1)C(C)C', 'non-sweetness'),
('CC(C)C(C(O)=O)S(=O)(=O)c1ccc(cc1)C(C)(C)C', 'non-sweetness'),
('CCC(C)(C(O)=O)S(=O)(=O)c1ccccc1', 'non-sweetness'),
('OC(=O)C(C(=O)c1ccccc1)S(=O)(=O)c1ccccc1', 'non-sweetness'),
('CCCCC(C(O)=O)S(=O)(=O)c1ccccc1', 'non-sweetness'),
('OCC1OC(OC2(CO)OC(CO)C(O)C2O)C(Cl)C(O)C1O', 'non-sweetness'),
('CCC(CC)(C(O)=O)S(=O)(=O)c1ccccc1', 'non-sweetness'),
('CC(C)(C(O)=O)S(=O)(=O)c1ccccc1', 'non-sweetness'),
('[Na+].CC1SCCCC1NS([O-])(=O)=O', 'non-sweetness'),
('OC(=O)C(Cc1ccccc1)S(=O)(=O)c1ccccc1', 'non-sweetness'),
```

```
('CCCC(C(O)=O)S(=O)(=O)c1ccccc1', 'non-sweetness'),
('CC(C(O)=O)S(=O)(=O)c1ccccc1', 'non-sweetness'),
('[Na+].[O-]S(=O)(=O)Nc1ccccc1N1CCCCC1', 'non-sweetness'),
('CCCC(CCC)(C(O)=O)S(=O)(=O)c1ccccc1', 'non-sweetness'),
('[Na+].[O-]S(=O)(=O)Nc1ncccn1', 'non-sweetness'),
('[Na+].[O-]S(=O)(=O)NC1=NCCS1', 'non-sweetness'),
('CC(CC(O)=O)S(=O)(=O)c1ccc(cc1)Br', 'non-sweetness'),
('CCC(C)C(COC(C)=O)NC(=O)C(N)CC(O)=O', 'non-sweetness'),
('CC(=O)OCC(Cc1ccccc1)NC(=O)C(N)CC(O)=O', 'non-sweetness'),
('CC(C)CC(COC(C)=O)NC(=O)C(N)CC(O)=O', 'non-sweetness'),
('CC1CCCCC1OC(=O)CCNC(=O)C(N)CC(O)=O', 'non-sweetness'),
('COC(=O)C(NC(=O)C(N)CC(O)=O)C(=O)OC1CC(C)CCC1C(C)C', 'non-sweetness'),
('COC(=O)C(CNC(=O)C(N)CC(O)=O)Cc1ccccc1', 'non-sweetness'),
('CCCCOC(=O)CCNC(=O)C(N)CC(O)=O', 'non-sweetness'),
('CCOC(=O)CC(Cc1ccccc1)NC(=O)C(N)CC(O)=O', 'non-sweetness'),
('CC(COC(=O)C1CCCC(C)(C)C1)NC(=O)C(N)CC(O)=O', 'non-sweetness'),
('CC(COC(=O)c1ccccc1)NC(=O)C(N)CC(O)=O', 'non-sweetness'),
('CCC(COC(=O)C1CCCCC1)NC(=O)C(N)CC(O)=O', 'non-sweetness'),
('CC(COC(=O)C1CCCCC1)NC(=O)C(N)CC(O)=O', 'non-sweetness'),
('CCC(CC(=O)OC1CCCCC1)NC(=O)C(N)CC(O)=O', 'non-sweetness'),
('CCCCC(=O)OCC(CC)NC(=O)C(N)CC(O)=O', 'non-sweetness'),
('CCCC(COC(=O)CC)NC(=O)C(N)CC(O)=O', 'non-sweetness'),
('CCC(CC(=O)OC(C)C)NC(=O)C(N)CC(O)=O', 'non-sweetness'),
('COC(=O)C(COC(=O)c1ccccc1)NC(=O)C(N)CC(O)=O', 'non-sweetness'),
('COC(=O)C(CNC(=O)C(N)CC(O)=O)C(=O)OC1CCCCC1', 'non-sweetness'),
('COC(=O)C(CNC(=O)C(N)CC(O)=O)C(=O)OC1CCCC1', 'non-sweetness'),
('COC(=O)C(CNC(=O)C(N)CC(O)=O)C(=O)OC', 'non-sweetness'),
('COC(=O)C(CC(=O)OC1CCCCC1)NC(=O)C(N)CC(O)=O', 'non-sweetness'),
('CC(C)CC(NC(=O)C(N)CC(O)=O)C(=O)OC(C)C', 'non-sweetness'),
('[Na+].[O-]S(=O)(=O)Nc1ncc(cc1Cl)C(F)(F)F', 'non-sweetness'),
('[Na+].Cc1cccnc1NS([O-])(=O)=O', 'non-sweetness'),
('CC(=O)OC1C(O)C(CO)OC1(CO)OC1OC(CO)C(O)C(O)C1O', 'non-sweetness'),
('CC(CC(O)=O)S(=O)(=O)c1ccccc1', 'non-sweetness'),
('[Na+].[O-]S(=O)(=O)Nc1cc(ccn1)C(F)(F)F', 'non-sweetness'),
('[Na+].[O-]S(=O)(=O)Nc1nc(cc(n1)Cl)Cl', 'non-sweetness'),
('OCC1OC(OC2(CO)OC(CCl)C(Cl)C2O)C(O)C(O)C1O', 'non-sweetness'),
('[Na+].Cc1cc(nc(n1)NS([O-])(=O)=O)C', 'non-sweetness'),
('[Na+].CCc1ccnc(c1)NS([O-])(=O)=O', 'non-sweetness'),
('[Na+].COc1cc(nc(n1)NS([O-])(=O)=O)C', 'non-sweetness'),
('[Na+].Cc1ccnc(c1)NS([O-])(=O)=O', 'non-sweetness'),
('[Na+].Cc1ccnc(n1)NS([O-])(=O)=O', 'non-sweetness'),
('[Na+].CC1CCSCC1NS([O-])(=O)=O', 'non-sweetness'),
('[Na+].[O-]S(=O)(=O)NN1CCC(CC1)c1ccccc1', 'non-sweetness'),
('[Na+].Cc1nnc(nc1C)NS([O-])(=O)=O', 'non-sweetness'),
('[Na+].[O-]S(=O)(=O)Nc1ccc(cn1)Br', 'non-sweetness'),
('[Na+].[O-]S(=O)(=O)Nc1ccc(cn1)Cl', 'non-sweetness'),
('OCC1OC(OC2(CCl)OC(CO)C(O)C2O)C(Cl)C(O)C1O', 'non-sweetness'),
```

```
('[Na+].CCOc1ccc2c(c1)N=C(NS([O-])(=O)=O)S2', 'non-sweetness'),
('[Na+].[O-]S(=O)(=O)Nc1cccc2cnccc12', 'non-sweetness'),
('[Na+].Cc1ccc(nc1)NS([O-])(=O)=O', 'non-sweetness'),
('[Na+].CC1=CN=C(NS([O-])(=O)=O)S1', 'non-sweetness'),
('[Na+].CC1CSCC(C1)NS([O-])(=O)=O', 'non-sweetness'),
('[Na+].[O-]S(=O)(=O)Nc1ccc(cn1)N(=O)=O', 'non-sweetness'),
('CC(C)=CCCC(C)(O)C1CCC(=CC1O)C', 'non-sweetness'),
('OCC1OC(CCl)(OC2OC(CCl)C(O)C(O)C2O)C(O)C1O', 'non-sweetness'),
('[Na+].[O-]S(=O)(=O)Nc1cc(cc(n1)Cl)C(F)(F)F', 'non-sweetness'),
('[Na+].CC1CCC(CS1)NS([O-])(=O)=O', 'non-sweetness'),
('CC(=O)OCC1OC(OC2(CO)OC(CO)C(O)C2O)C(O)C(O)C1O', 'non-sweetness'),
('CC(C)OCC1OC(OC2(CCl)OC(CCl)C(O)C2O)C(O)C(O)C1Cl', 'non-sweetness'), ('COC(=O)C
1OC(OC2C(O)C(O)C(CO)OC2OC2CCC34CC54CCC4(C)C(CCC4(C)C5CCC3C2(C)C(=O)OC)C(C)C2CC=C
(C)C(=O)O2)C(O)C(O)C1O', 'non-sweetness'),
('[Na+].[Na+].[O-]S(=O)(=O)Nc1ccccc1[S-]', 'non-sweetness'),
('[Na+].[Na+].[O-]S(=O)(=O)Nc1cccc(c1)[S-]', 'non-sweetness'),
('[Na+].[Na+].[O-]S(=O)(=O)Nc1ccc(cc1)[S-]', 'non-sweetness'), ('CCO', 'non-
sweetness'), ('COC1(CO)OC(CO)C(O)C1O', 'non-sweetness'),
('[Na+].[O-]S(=O)(=O)Nc1cccc(c1)N(=O)=O', 'non-sweetness'),
('[Na+].CCC(C)NS([O-])(=O)=O', 'non-sweetness'), ('CC=C(C)C(C)=NO', 'non-
sweetness'), ('CC1CC=C(C=NO)C(C)O1', 'non-sweetness'), ('CC(=NO)C1=CCCCC1',
'non-sweetness'), ('COCC(OC)C1CCC(=CC1)C=NO', 'non-sweetness'),
('COCC1=CCC=C(C1)C=NO', 'non-sweetness'), ('CCCC(CC)(C(O)=O)S(=O)(=O)c1ccccc1',
'non-sweetness'), ('COC1C(O)C(C)OC(OC2OC(CCC2C)C(C)C2CCC3C4=CC(=O)C5CC(CCC5(C)C4
CCC23C)OC2OC(CO)C(O)C(O)C2O)C1O', 'non-sweetness'), ('COC(=O)CC1C(O)C(Oc2c1c(cc(
c2C1C(O)C(Oc2c1c(cc1c2C2C(O)C(Oc3cc(cc(c23)O)O)(O1)c1ccc(cc1)O)O)c1ccc(cc1)O)O)O
)c1ccc(cc1)O', 'non-sweetness'), ('[Na+].CCC(CO)NS([O-])(=O)=O', 'non-
sweetness'), ('[Na+].CC(C)(OCC([O-])=O)C1CCC(=CC1)C=NO', 'non-sweetness'),
('[Na+].CC(C)(CO)NS([O-])(=O)=O', 'non-sweetness'),
('[Na+].[O-]c1cccc(c1)NS([O-])(=O)=O', 'non-sweetness'),
('[Na+].[O-]S(=O)(=O)N[n+]1ccc(cc1)-c1ccccc1', 'non-sweetness'),
('[Na+].CCCCCCNS([O-])(=O)=O', 'non-sweetness'), ('[Na+].CC(C)NS([O-])(=O)=O',
'non-sweetness'), ('[Na+].CNS([O-])(=O)=O', 'non-sweetness'),
('[Na+].[O-]S(=O)(=O)Nc1ccc2c(c1)OCO2', 'non-sweetness'),
('[Na+].COCC(C)NS([O-])(=O)=O', 'non-sweetness'),
('[Na+].COC(CNS([O-])(=O)=O)OC', 'non-sweetness'),
('[Na+].[O-]S(=O)(=O)Nc1cccc(c1Cl)Cl', 'non-sweetness'),
('[Na+].[O-]S(=O)(=O)Nc1cccc(c1F)F', 'non-sweetness'),
('[Na+].[O-]S(=O)(=O)Nc1ccc2c(c1)CCC2', 'non-sweetness'),
('[Na+].CC(C)(C)CC(C)(C)NS([O-])(=O)=O', 'non-sweetness'),
('[Na+].[O-]S(=O)(=O)Nc1ccc(cc1Cl)Cl', 'non-sweetness'),
('[Na+].[O-]S(=O)(=O)Nc1ccc(cc1F)F', 'non-sweetness'),
('[Na+].Cc1ccc(c(c1)C)NS([O-])(=O)=O', 'non-sweetness'),
('[Na+].[O-]S(=O)(=O)Nc1ccc(cc1N(=O)=O)N(=O)=O', 'non-sweetness'),
('[Na+].[O-]S(=O)(=O)Nc1cc(ccc1Cl)Cl', 'non-sweetness'),
('[Na+].[O-]S(=O)(=O)Nc1c(cccc1F)F', 'non-sweetness'),
('[Na+].Cc1cccc(c1NS([O-])(=O)=O)C', 'non-sweetness'),
```

('[Na+].Nc1ccccc1NS([O-])(=O)=O', 'non-sweetness'),
('[Na+].[O-]S(=O)(=O)Nc1ccccc1Br', 'non-sweetness'),
('[Na+].[O-]S(=O)(=O)Nc1ccccc1Cl', 'non-sweetness'),
('[Na+].[O-]S(=O)(=O)Nc1cccnc1Cl', 'non-sweetness'),
('[Na+].[O-]S(=O)(=O)Nc1ccccc1C#N', 'non-sweetness'),
('[Na+].[O-]S(=O)(=O)NCCC1CCCCC1', 'non-sweetness'),
('[Na+].CCOC(C)CNS([O-])(=O)=O', 'non-sweetness'),
('[Na+].CCc1ccccc1NS([O-])(=O)=O', 'non-sweetness'),
('[Na+].Cc1ccc(c(c1)NS([O-])(=O)=O)F', 'non-sweetness'),
('[Na+].[O-]S(=O)(=O)Nc1ccccc1F', 'non-sweetness'), ('[Na+].OCCNS([O-])(=O)=O', 'non-sweetness'), ('[Na+].[O-]S(=O)(=O)Nc1ccccc1I', 'non-sweetness'),
('[Na+].COc1cc(ccc1NS([O-])(=O)=O)N(=O)=O', 'non-sweetness'),
('[Na+].COc1ccc(cc1NS([O-])(=O)=O)N(=O)=O', 'non-sweetness'),
('[Na+].COCCNS([O-])(=O)=O', 'non-sweetness'), ('[Na+].CCC(C)(C)NS([O-])(=O)=O', 'non-sweetness'), ('[Na+].[O-]S(=O)(=O)Nc1ccccc1N(=O)=O', 'non-sweetness'),
('[Na+].[O-]S(=O)(=O)NCCN1CCCCC1', 'non-sweetness'),
('[Na+].[O-]S(=O)(=O)Nc1ccc2c(c1)N=C(S)S2', 'non-sweetness'),
('[Na+].CC(NS([O-])(=O)=O)C(C)(C)C', 'non-sweetness'),
('[Na+].[O-]S(=O)(=O)Nc1ccc(c(c1)Cl)Cl', 'non-sweetness'),
('[Na+].[O-]S(=O)(=O)Nc1ccc(c(c1)F)F', 'non-sweetness'),
('[Na+].Cc1ccc(cc1C)NS([O-])(=O)=O', 'non-sweetness'),
('[Na+].CC(=O)c1cccc(c1)NS([O-])(=O)=O', 'non-sweetness'),
('[Na+].Nc1cccc(c1)NS([O-])(=O)=O', 'non-sweetness'),
('[Na+].[O-]S(=O)(=O)Nc1ccc(c(c1)Cl)F', 'non-sweetness'),
('[Na+].Cc1ccc(cc1Cl)NS([O-])(=O)=O', 'non-sweetness'),
('[Na+].CCOCCCNS([O-])(=O)=O', 'non-sweetness'),
('[Na+].Cc1c(cccc1NS([O-])(=O)=O)F', 'non-sweetness'),
('[Na+].Oc1cccc(c1)NS([O-])(=O)=O', 'non-sweetness'),
('[Na+].OCCCNS([O-])(=O)=O', 'non-sweetness'),
('[Na+].[O-]S(=O)(=O)Nc1cccc(c1)I', 'non-sweetness'),
('[Na+].COCCCNS([O-])(=O)=O', 'non-sweetness'),
('[Na+].CCC(C)C(C)NS([O-])(=O)=O', 'non-sweetness'),
('[Na+].CCCc1cccc(c1)NS([O-])(=O)=O', 'non-sweetness'),
('[Na+].CC(=O)c1ccc(cc1)NS([O-])(=O)=O', 'non-sweetness'),
('[Na+].CCCCc1ccc(cc1)NS([O-])(=O)=O', 'non-sweetness'),
('[Na+].[O-]S(=O)(=O)NC1=C(N=CN1)C#N', 'non-sweetness'),
('[Na+].CCOc1ccc(c(c1)N(=O)=O)NS([O-])(=O)=O', 'non-sweetness'),
('[Na+].CCC1=C(C)SC(=N1)NS([O-])(=O)=O', 'non-sweetness'),
('[Na+].[O-]S(=O)(=O)Nc1ccc(c(c1)N(=O)=O)F', 'non-sweetness'),
('[Na+].CC(C)CC(C)NS([O-])(=O)=O', 'non-sweetness'),
('[Na+].CC(C)CCCNS([O-])(=O)=O', 'non-sweetness'),
('[Na+].CC1CCN(CC1)NS([O-])(=O)=O', 'non-sweetness'),
('[Na+].[O-]S(=O)(=O)NC1=NC(=CS1)c1ccccc1', 'non-sweetness'),
('[Na+].[O-]S(=O)(=O)Nc1ccc(cc1)-c1ccccc1', 'non-sweetness'),
('[Na+].CCCc1ccc(cc1)NS([O-])(=O)=O', 'non-sweetness'),
('[Na+].CC(C)(C)c1ccc(cc1)NS([O-])(=O)=O', 'non-sweetness'),
('[Na+].Cc1cc(cnc1NS([O-])(=O)=O)Br', 'non-sweetness'),

```
('[Na+].CCCCC1=NN=C(NS([O-])(=O)=O)S1', 'non-sweetness'),
('[Na+].CCCCC1=C(C)N=C(NS([O-])(=O)=O)S1', 'non-sweetness'),
('[Na+].Cc1ccc(cc1NS([O-])(=O)=O)Cl', 'non-sweetness'),
('[Na+].[O-]S(=O)(=O)Nc1cnc(cn1)Cl', 'non-sweetness'),
('[Na+].CCC1=C(C)N=C(NS([O-])(=O)=O)S1', 'non-sweetness'),
('[Na+].Cc1ccc(cc1NS([O-])(=O)=O)F', 'non-sweetness'),
('[Na+].CC(C)CCC(C)NS([O-])(=O)=O', 'non-sweetness'),
('[Na+].CCCCCC1=NN=C(NS([O-])(=O)=O)S1', 'non-sweetness'),
('[Na+].CCOc1ccc2c(c1)SC(=N2)NS([O-])(=O)=O', 'non-sweetness'),
('[Na+].[O-]S(=O)(=O)NC1=Nc2ccc(cc2S1)F', 'non-sweetness'),
('[Na+].COc1ccc2c(c1)SC(=N2)NS([O-])(=O)=O', 'non-sweetness'),
('[Na+].CCCCCC(C)NS([O-])(=O)=O', 'non-sweetness'),
('[Na+].CCCCC(C)NS([O-])(=O)=O', 'non-sweetness'),
('[Na+].[O-]S(=O)(=O)NN1CCCC1', 'non-sweetness'),
('[Na+].CCC(C)c1cccc(c1NS([O-])(=O)=O)CC', 'non-sweetness'),
('[Na+].CN(C)CCNS([O-])(=O)=O', 'non-sweetness'),
('[Na+].[O-]S(=O)(=O)Nc1ccccc1N1CCOCC1', 'non-sweetness'),
('[Na+].CC(C)OCCNS([O-])(=O)=O', 'non-sweetness'),
('[Na+].[O-]S(=O)(=O)Nc1ccccc1C(F)(F)F', 'non-sweetness'),
('[Na+].[O-]S(=O)(=O)Nc1cc(ccc1Br)C(F)(F)F', 'non-sweetness'),
('[Na+].[O-]S(=O)(=O)Nc1ccc(cc1Cl)C(F)(F)F', 'non-sweetness'),
('[Na+].[O-]S(=O)(=O)Nc1cc(ccc1Cl)C(F)(F)F', 'non-sweetness'),
('[Na+].CCc1cccc(c1NS([O-])(=O)=O)C(C)C', 'non-sweetness'),
('[Na+].[O-]S(=O)(=O)Nc1cc(ccc1F)C(F)(F)F', 'non-sweetness'),
('[Na+].CC(C)c1cccc(c1NS([O-])(=O)=O)C', 'non-sweetness'),
('[Na+].CN(C)CCCNS([O-])(=O)=O', 'non-sweetness'),
('[Na+].[O-]S(=O)(=O)Nc1cccc(c1)C(F)(F)F', 'non-sweetness'),
('[Na+].[O-]S(=O)(=O)Nc1cc(cc(c1)C(F)(F)F)C(F)(F)F', 'non-sweetness'),
('[Na+].[O-]S(=O)(=O)Nc1cc(cc(c1)C(F)(F)F)Cl', 'non-sweetness'),
('[Na+].CCOC(=O)CC1=CSC(=N1)NS([O-])(=O)=O', 'non-sweetness'),
('[Na+].[O-]S(=O)(=O)NC1=NC(=CS1)c1ccccc1N(=O)=O', 'non-sweetness'),
('[Na+].[O-]S(=O)(=O)NC1=NC(=CS1)c1cccc(c1)N(=O)=O', 'non-sweetness'),
('[Na+].[O-]S(=O)(=O)NC1=NC(=CS1)c1ccc(cc1)Cl', 'non-sweetness'),
('[Na+].CCC(CC)c1ccc(cc1)NS([O-])(=O)=O', 'non-sweetness'),
('[Na+].CC(Cc1ccccc1)C1=NN=C(NS([O-])(=O)=O)S1', 'non-sweetness'),
('[Na+].CC(C)CC1=NN=C(NS([O-])(=O)=O)S1', 'non-sweetness'),
('[Na+].[O-]S(=O)(=O)NC1=NN=C(CCc2ccccc2)S1', 'non-sweetness'),
('[Na+].[O-]S(=O)(=O)NC1=NN=C(S1)c1cccc(c1)Cl', 'non-sweetness'),
('[Na+].CC(C)CCC1=NN=C(NS([O-])(=O)=O)S1', 'non-sweetness'),
('[Na+].[O-]S(=O)(=O)NC1=NN=C(S1)c1ccc(cc1)Cl', 'non-sweetness'),
('[Na+].CCC(C)C1=NN=C(NS([O-])(=O)=O)S1', 'non-sweetness'),
('[Na+].CSC1=NN=C(NS([O-])(=O)=O)S1', 'non-sweetness'),
('[Na+].CCCC(C)C1=NN=C(NS([O-])(=O)=O)S1', 'non-sweetness'),
('[Na+].CCCCCNS([O-])(=O)=O', 'non-sweetness'), ('[Na+].CCNS([O-])(=O)=O', 'non-
sweetness'), ('[Na+].CCCCCCCNS([O-])(=O)=O', 'non-sweetness'),
('[Na+].CCCCCCCCNS([O-])(=O)=O', 'non-sweetness'),
('[Na+].CCC(CC)NS([O-])(=O)=O', 'non-sweetness'),
```

```
('[Na+].CC(C)(C)NS([O-])(=O)=O', 'non-sweetness'),
('[Na+].Cc1ccc(cc1)NS([O-])(=O)=O', 'non-sweetness'),
('CC(=O)OC1C(Cl)C(CO)OC(OC2(CCl)OC(CCl)C3OC23)C1OC(C)=O', 'non-sweetness'),
('[Na+].[O-]S(=O)(=O)NC1CSC1', 'non-sweetness'),
('[Na+].[O-]S(=O)(=O)NC1CCCCSC1', 'non-sweetness'),
('[Na+].[O-]S(=O)(=O)NC1CCCSCC1', 'non-sweetness'),
('Oc1ccc2c(c1)OCC1(O)Cc3cc(c(cc3C21)O)O', 'miscellaneous'),
('OC1CC(O)C(O)C(O)C1O', 'miscellaneous'),
('COc1cc(ccc1O)C(=O)OCC(=CCOC1OC(CO)C(O)C(O)C1O)C#N', 'miscellaneous'),
('OCC1OC(OCC=C(COC(=O)c2ccc(cc2)O)C#N)C(O)C(O)C1O', 'miscellaneous'),
('CCCCCCCC(=O)CCc1ccc(c(c1)OC)O', 'miscellaneous'),
('CCCCCCCC(O)CC(=O)CCc1ccc(c(c1)OC)O', 'miscellaneous'),
('OCC1OC(C(O)C(O)C1O)N1C=C(CC(O)=O)c2ccccc12', 'miscellaneous'),
('[Na+].[O-]S(=O)(=O)NC1CSCCSC1', 'miscellaneous'),
('COc1ccc2c(c1O)CCC1C2CCC2(C)CCCC12', 'miscellaneous'),
('Nc1ccc2c(c1)S(=O)(=O)NC2=O', 'miscellaneous'),
('CCCCCC=CC=CC(CCCCCCCC(O)=O)OO', 'miscellaneous'),
('CCCCCC(OO)C=CC=CCCCCCCCC(O)=O', 'miscellaneous'),
('CC=CC=CC=CCCC=CC(=O)NCC(C)(C)O', 'miscellaneous'),
('CCCCCC=CCC=CCC=CCC=CCCCC(O)=O', 'miscellaneous'),
('COc1ccc(cc1)S(=O)(=O)C(C(C)C)C(O)=O', 'miscellaneous'),
('COc1cc(ccc1O)CNC(=O)CCCCC=CC(C)C', 'miscellaneous'),
('CCCCCC=CC=CC(=O)NCC(C)C', 'miscellaneous'),
('OC(=O)C(Cc1ccc(c(c1)O)O)NC(=O)C=Cc1ccc(c(c1)O)O', 'miscellaneous'),
('CC(C)C1CCC(C)CC1OCC(O)CO', 'miscellaneous'),
('CC(C)C1CCC(C)C23CCC(C)(O)C3C12', 'miscellaneous'),
('[Na+].[O-]S(=O)(=O)NC1CCC1', 'miscellaneous'),
('COc1cc(ccc1O)CNC(=O)CCCCCCC(C)C', 'miscellaneous'),
('CC(C)C1CCC(C)CC1C(=O)Nc1ccc(cc1)CC#N', 'miscellaneous'),
('CC(C)C1CCC(C)CC21OCC(CO)O2', 'miscellaneous'), ('CC(C)C1CCC(C)CC1OC(=O)OCCO',
'miscellaneous'), ('CCCCCC(O)CC(=O)CCc1ccc(c(c1)OC)O', 'miscellaneous'),
('OC1OC(COC(=O)c2cc(c(c(c2)O)O)O)C(O)C1(O)COC(=O)c1cc(c(c(c1)O)O)O',
'miscellaneous'), ('Oc1ccccc1N1CC=C(NC1=O)c1cccc(c1)N(=O)=O', 'miscellaneous'),
('CC1(C)OC2CC(=O)OCC32C2CCC(C)(C(O)C4=COC=C4)C4(OC4(C)C(O)=O)C2(C)C(=O)CC13',
'miscellaneous'), ('CC(C)C1CCC(C)CC1OC(=O)CCCC(O)=O', 'miscellaneous'),
('CC(C)C1CCC(C)CC1OC(=O)C(C)O', 'miscellaneous'),
('CC(C)C1CCC(C)CC1OC(=O)CCC(O)=O', 'miscellaneous'),
('COC(=O)CC1=CN(C2OC(CO)C(O)C(O)C2O)c2ccccc12', 'miscellaneous'),
('CC1(C)CCCC2(C)C3CC=C(C=O)C(=CC3CCC12)C=O', 'miscellaneous'),
('Oc1ccc(cc1)C1CC(=O)c2c(cc(cc2O1)O)O', 'miscellaneous'),
('COc1ccc(cc1OC)CCC(=O)c1c(cc(cc1O)OC1OC(CO)C(O)C(O)C1OC1OC(C)C(O)C(O)C1O)O',
'miscellaneous'), ('CCCCCCCCC(=O)NCc1ccc(c(c1)OC)O', 'miscellaneous'),
('COc1cc(ccc1O)CNC(=O)CCCCCC(C)C', 'miscellaneous'),
('O=C(C=CCCc1ccc2c(c1)OCO2)N1CCCCC1', 'miscellaneous'),
('O=C(C=CC=Cc1ccc2c(c1)OCO2)N1CCCCC1', 'miscellaneous'),
('O=C(C=CC=Cc1ccc2c(c1)OCO2)N1CCCC1', 'miscellaneous'),
('CC1(C)CCCC2(C)C1CC=C(C=O)C2C=O', 'miscellaneous'), ('OC1Cc2c(cc3c(c2OC1c1ccc(c
```

```
(c1)O)O)C1C(O)C(Oc2cc(cc(c12)O)O)(O3)c1ccc(c(c1)O)O)O', 'miscellaneous'), ('CC1O
C(OCC2OC(OC3=C(Oc4cc(cc(c4C3=O)O)O)c3ccc(c(c3)O)O)C(O)C(O)C2O)C(O)C(O)C1O',
'miscellaneous'), ('CCCCCC=CC(=O)CCc1ccc(c(c1)OC)O', 'miscellaneous'),
('[Na+].Cc1cccc(c1C)NS([O-])(=O)=O', 'miscellaneous'),
('[Na+].CCOc1ccc(cc1)NS([O-])(=O)=O', 'miscellaneous'),
('[Na+].COc1ccc(cc1)NS([O-])(=O)=O', 'miscellaneous'),
('[Na+].[O-]c1ccc(cc1NS([O-])(=O)=O)N(=O)=O', 'miscellaneous'),
('[Na+].[O-]S(=O)(=O)Nc1cc(ccc1F)F', 'miscellaneous'),
('[Na+].Cc1ccc(c(c1)NS([O-])(=O)=O)C', 'miscellaneous'),
('[Na+].[O-]S(=O)(=O)Nc1cc(ccc1Cl)N(=O)=O', 'miscellaneous'),
('[Na+].CCOC(=O)c1ccccc1NS([O-])(=O)=O', 'miscellaneous'),
('[Na+].[O-]S(=O)(=O)Nc1cc(ccc1F)N(=O)=O', 'miscellaneous'),
('[Na+].COc1ccccc1NS([O-])(=O)=O', 'miscellaneous'),
('[Na+].Cc1ccc(cc1NS([O-])(=O)=O)N(=O)=O', 'miscellaneous'),
('[Na+].COc1ccc(cc1OC)NS([O-])(=O)=O', 'miscellaneous'),
('[Na+].Cc1cc(cc(c1)NS([O-])(=O)=O)C', 'miscellaneous'),
('[Na+].COc1cccc(c1)NS([O-])(=O)=O', 'miscellaneous'),
('[Na+].[O-]S(=O)(=O)Nc1ccc(cc1)C#N', 'miscellaneous'),
('[Na+].CCc1ccc(cc1)NS([O-])(=O)=O', 'miscellaneous'),
('[Na+].[O-]S(=O)(=O)Nc1ccc(cc1)I', 'miscellaneous'),
('[Na+].[O-]S(=O)(=O)Nc1ccc(cc1)OCc1ccccc1', 'miscellaneous'),
('[Na+].Cc1cc2c(cc1C)N=C(NS([O-])(=O)=O)S2', 'miscellaneous'),
('[Na+].COc1ccc(cc1NS([O-])(=O)=O)Cl', 'miscellaneous'),
('[Na+].CC(C)(C)C1=NN=C(NS([O-])(=O)=O)S1', 'miscellaneous'),
('[Na+].CCCC(C)NS([O-])(=O)=O', 'miscellaneous'),
('[Na+].OCc1ccccc1NS([O-])(=O)=O', 'miscellaneous'),
('[Na+].OCc1cccc(c1)NS([O-])(=O)=O', 'miscellaneous'),
('[Na+].[O-]S(=O)(=O)NC1=NC(=CS1)c1ccc(cc1)Br', 'miscellaneous'),
('[Na+].Cc1ccc(cc1)C1=CSC(=N1)NS([O-])(=O)=O', 'miscellaneous'),
('[Na+].[O-]S(=O)(=O)NC1=NN=C(S1)c1ccccc1Br', 'miscellaneous'),
('[Na+].[O-]S(=O)(=O)NC1=NN=C(S1)c1cccc(c1)Br', 'miscellaneous'),
('CC=CC=CCCC=CC(=O)NCC(C)C', 'miscellaneous'), ('Oc1cc(cc(c1O)O)C(=O)Oc1cc(cc(c1
O)O)C(=O)OCC1OC(OC(=O)c2cc(c(c(c2)OC(=O)c2cc(c(c(c2)O)O)O)O)O)C(OC(=O)c2cc(c(c(c
2)OC(=O)c2cc(c(c(c2)O)O)O)O)O)C(OC(=O)c2cc(c(c(c2)OC(=O)c2cc(c(c(c2)O)O)O)O)O)C1
OC(=O)c1cc(c(c(c1)OC(=O)c1cc(c(c(c1)O)O)O)O)O)O', 'miscellaneous'),
('CCNC(=O)C(C)(C(C)C)C(C)C', 'miscellaneous'), ('CCNC(=O)C1CC(C)CCC1C(C)C',
'miscellaneous')])
```

[ ]:
```python
len(new_data)
```

[ ]: 2785

[ ]:
```python
temp_data=pd.DataFrame(data=new_data.items(),columns=["canonical SMILES","Class␣
↪taste"])
temp_data.head()
```

[ ]:

```
                              canonical SMILES Class taste
0         Oc1cc2c(cc1O)C1c3ccc(c(c3OCC1(O)C2)O)O   sweetness
1              CC(C)=CCCC(C)(O)C1CC(O)C(=CC1=O)C   sweetness
2  CC(=O)OC1C(Oc2cc(cc(c2C1=O)O)O)c1ccc(c(c1)O)O   sweetness
3                         OC1C(O)C(O)C(O)C(O)C1O   sweetness
4                        CON=C(C)C1CCC(=CC1)C=NO   sweetness
```

[ ]:

```
#Now let's see how many instances we have belonging to each class of odor
from collections import defaultdict
def scatter_taste_dist(data):
  taste_count = defaultdict(int)
  for sentence in data:
      for taste in sentence:
          taste_count[taste] += 1
  plt.figure(figsize=(20, 10))
  plt.xticks(rotation="vertical")
  plt.xlabel("Taste")
  plt.ylabel("No of each Taste label")
  plt.scatter([x for x in taste_count.keys()],[x for x in taste_count.
 ↪values()], s = [x for x in taste_count.values()])

def scatter_class_taste_dist(data):
  class_taste_count = defaultdict(int)
  for taste in data:
        class_taste_count[taste] += 1
  plt.figure(figsize=(20, 10))
  plt.xticks(rotation="vertical")
  plt.xlabel("Class Taste")
  plt.ylabel("No of each Taste label")
  plt.scatter([x for x in class_taste_count.keys()],[x for x in␣
 ↪class_taste_count.values()], s = [x for x in class_taste_count.values()])
```

[ ]:

```
#Label distribution in fermenich dataset
scatter_taste_dist(data["Taste"])
```

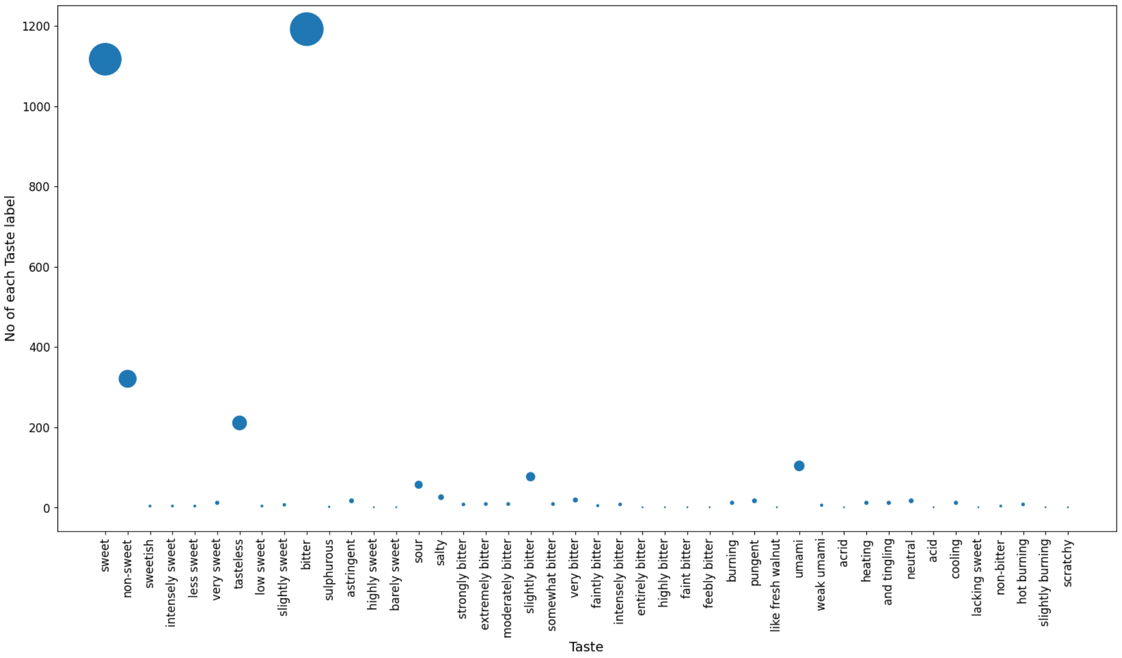


[ ]:
```python
#Label distribution in fermenich dataset
scatter_class_taste_dist(data["Class taste"])
```

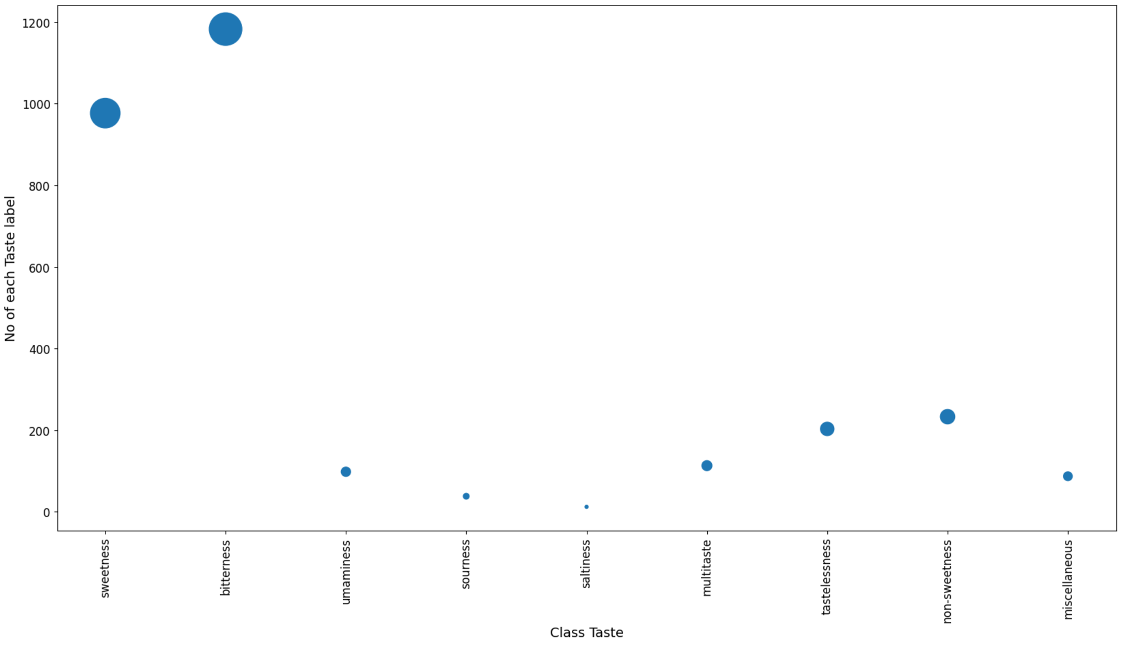

```
[ ]: #Now let's see how many instances we have belonging to each class of odor
from collections import defaultdict
def label_taste_dist(data):
  taste_count = defaultdict(int)
  for sentence in data:
      for taste in sentence:
          taste_count[taste] += 1
  plt.figure(figsize=(20, 10))
  plt.xticks(rotation="vertical")
  plt.xlabel("Taste")
  plt.ylabel("Percantage of each Taste label")
  plt.bar([x for x in taste_count.keys()],[(x/len(data))*100 for x in taste_count.values()])
```

```
[ ]: #Label distribution in fermenich dataset
label_taste_dist(data["Taste"])
```

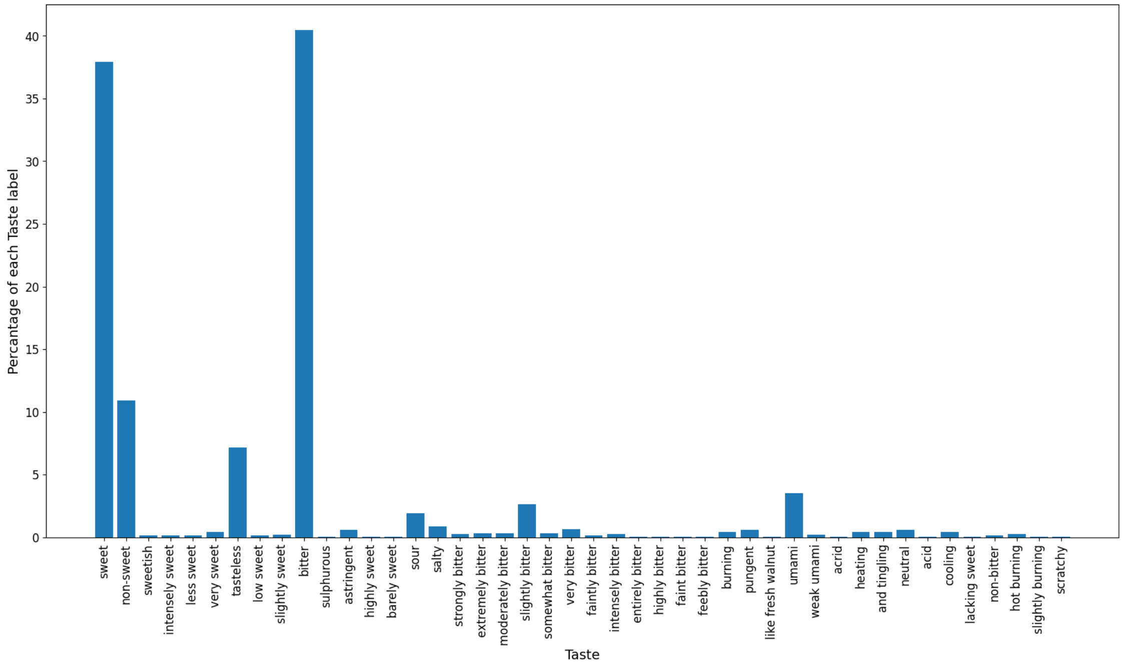


```
[ ]: taste_count = defaultdict(int)
for sentence in data["Taste"]:
    for taste in sentence:
        taste_count[taste] += 1
table=pd.DataFrame(sorted(taste_count.items(),key=lambda x: x[1],reverse=True),columns=["Taste","No of associated samples"])
table["Percentage_of_samples"]=(table["No of associated samples"]/data.shape[0])*100
```

```python
table.head(100)
#We thus get a list of top 10 most frequent odor descriptors. We observe that␣
↪fruity is most common
#with 2050 molecules having the associated Odor.
```

[ ]:

| | Taste | No of associated samples | Percentage_of_samples |
|---|---|---|---|
| 0 | bitter | 1192 | 40.489130 |
| 1 | sweet | 1117 | 37.941576 |
| 2 | non-sweet | 321 | 10.903533 |
| 3 | tasteless | 211 | 7.167120 |
| 4 | umami | 104 | 3.532609 |
| 5 | slightly bitter | 77 | 2.615489 |
| 6 | sour | 57 | 1.936141 |
| 7 | salty | 26 | 0.883152 |
| 8 | very bitter | 19 | 0.645380 |
| 9 | astringent | 17 | 0.577446 |
| 10 | pungent | 17 | 0.577446 |
| 11 | neutral | 17 | 0.577446 |
| 12 | very sweet | 12 | 0.407609 |
| 13 | burning | 12 | 0.407609 |
| 14 | heating | 12 | 0.407609 |
| 15 | and tingling | 12 | 0.407609 |
| 16 | cooling | 12 | 0.407609 |
| 17 | extremely bitter | 9 | 0.305707 |
| 18 | moderately bitter | 9 | 0.305707 |
| 19 | somewhat bitter | 9 | 0.305707 |
| 20 | strongly bitter | 8 | 0.271739 |
| 21 | intensely bitter | 8 | 0.271739 |
| 22 | hot burning | 8 | 0.271739 |
| 23 | slightly sweet | 7 | 0.237772 |
| 24 | weak umami | 6 | 0.203804 |
| 25 | faintly bitter | 5 | 0.169837 |
| 26 | sweetish | 4 | 0.135870 |
| 27 | intensely sweet | 4 | 0.135870 |
| 28 | less sweet | 4 | 0.135870 |
| 29 | low sweet | 4 | 0.135870 |
| 30 | non-bitter | 4 | 0.135870 |
| 31 | sulphurous | 2 | 0.067935 |
| 32 | highly sweet | 1 | 0.033967 |
| 33 | barely sweet | 1 | 0.033967 |
| 34 | entirely bitter | 1 | 0.033967 |
| 35 | highly bitter | 1 | 0.033967 |
| 36 | faint bitter | 1 | 0.033967 |
| 37 | feebly bitter | 1 | 0.033967 |
| 38 | like fresh walnut | 1 | 0.033967 |
| 39 | acrid | 1 | 0.033967 |
| 40 | acid | 1 | 0.033967 |

```
41      lacking sweet                          1              0.033967
42   slightly burning                          1              0.033967
43           scratchy                          1              0.033967
```

[ ]:
```
#Now let's see how many instances we have belonging to each class of odor
from collections import defaultdict
def label_classtaste_dist(data):
  classtaste_count = defaultdict(int)
  for classtaste in data:
      classtaste_count[classtaste] += 1
  plt.figure(figsize=(20, 10))
  plt.xticks(rotation="vertical")
  plt.xlabel("Class Taste")
  plt.ylabel("Percantage of each Class Taste label")
  plt.bar([x for x in classtaste_count.keys()],[(x/len(data))*100 for x in␣
↪classtaste_count.values()])
```

[ ]:
```
#Label distribution in fermenich dataset
label_classtaste_dist(data["Class taste"])
```

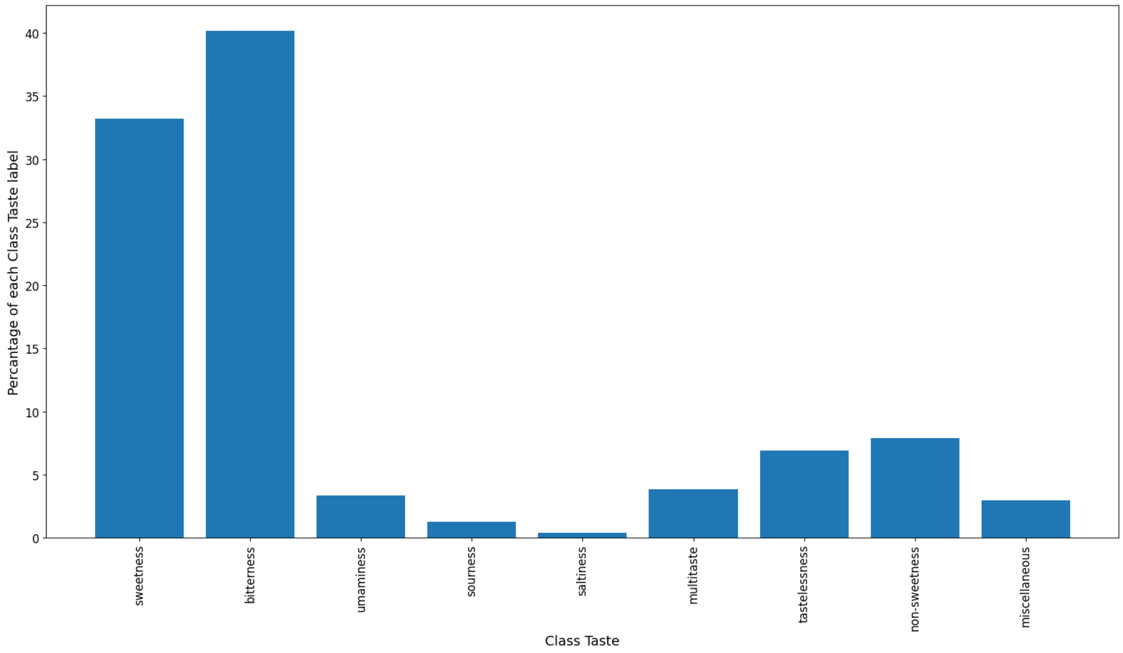


[ ]:
```
classtaste_count = defaultdict(int)
for classtaste in data["Class taste"]:
      classtaste_count[classtaste] += 1
table=pd.DataFrame(sorted(classtaste_count.items(),key=lambda x:␣
↪x[1],reverse=True),columns=["Class Taste","No of associated samples"])
```

```
table["Percentage_of_samples"]=(table["No of associated samples"]/data.
  ↪shape[0])*100
table.head(100)
#We thus get a list of top 10 most frequent odor descriptors. We observe that␣
  ↪fruity is most common
#with 2050 molecules having the associated Odor.
```

[ ]:

| | Class Taste | No of associated samples | Percentage_of_samples |
|---|---|---|---|
| 0 | bitterness | 1183 | 40.183424 |
| 1 | sweetness | 977 | 33.186141 |
| 2 | non-sweetness | 233 | 7.914402 |
| 3 | tastelessness | 203 | 6.895380 |
| 4 | multitaste | 113 | 3.838315 |
| 5 | umaminess | 98 | 3.328804 |
| 6 | miscellaneous | 87 | 2.955163 |
| 7 | sourness | 38 | 1.290761 |
| 8 | saltiness | 12 | 0.407609 |

[ ]:
```
sorted_classtaste=sorted(classtaste_count.items(),key=lambda x:␣
  ↪x[1],reverse=True)
print(sorted_classtaste)
```

```
[('bitterness', 1183), ('sweetness', 977), ('non-sweetness', 233),
('tastelessness', 203), ('multitaste', 113), ('umaminess', 98),
('miscellaneous', 87), ('sourness', 38), ('saltiness', 12)]
```

[ ]:
```
#Utility function to get the characteristics of our multi label dataset
def char_MLD(data):
  inst_sum=0
  for item in data["Class taste"]:
    inst_sum+=len(item)
  IRLbl=[]
  most_freq=sorted_classtaste[0][1]
  for item in sorted_classtaste:
    IRLbl.append(most_freq/item[1])
  print("Cardinality= {}".format(inst_sum/data.shape[0]))
  print("Density= {}".format((inst_sum/data.shape[0])*(1/(len(vocabt)))))
  print("MeanIR= {}".format(np.array(IRLbl).mean()))
```

[ ]:
```
char_MLD(data)
```

```
Cardinality= 10.137907608695652
Density= 0.2304069911067194
MeanIR= 19.885417501712798
```

[ ]:
```
df=data.sample(frac=1,random_state=42).reset_index(drop=True)
```

```
#Since we've concatenated two datsets on top of each other, to ensure there are
↪is no inherent
#Ordering in data which can bias our model, we reshuffle the data
```

[ ]:
```
df
```

[ ]:
```
         ID                                               Name PubChem CID  \
0      0566                                 Compound (R)-(+)-9           *
1      1221                                          Artemisin       65030
2      1678                                           L-Leucine        6106
3      1578                                     Hexethal sodium    23690440
4     2832*  Sodium N-[3-chloro-5-(trifluoromethyl)phenyl]s…           *
…       …                                                  …          …
2939   1639                                     Isoxanthohumol M   122363515
2940  1096*  3-Amino-3-{[1-(2H-1,3-benzodioxol-5-yl)propan-…           *
2941   1131                              5,7,2'-Trihydroxyflavone     5322064
2942   1295                                       Chlormerodrin       25210
2943  0861*            Sodium N-(2-methylthian-4-yl)sulfamate           *

     CAS number                                  canonical SMILES  \
0             *             COc1ccc(cc1O)C1OC2CC3CCC2(CS1)C3(C)C
1      481-05-0           CC1C2C(O)CC3(C)C=CC(=O)C(=C3C2OC1=O)C
2       61-90-5                                   CC(C)CC(N)C(O)=O
3      144-00-3            [Na+].CCCCCCC1(CC)C(=O)NC(=NC1=O)[O-]
4             *          [Na+].[O-]S(=O)(=O)Nc1cc(cc(c1)C(F)(F)F)Cl
…           …                                                  …
2939          *  COc1cc(c(c2c1C(=O)CC(O2)c1ccc(cc1)O)CCC(C)(C)OC)O
2940          *             CC(Cc1ccc2c(c1)OCO2)NC(=O)C(N)CC(O)=O
2941 73046-40-9               Oc1cc(c2c(c1)OC(=CC2=O)c1ccccc1O)O
2942    62-37-3                               COC(CNC(N)=O)C[Hg]Cl
2943          *                     [Na+].CC1CC(CCS1)NS([O-])(=O)=O

                                     Taste   Class taste  \
0                                  [sweet]     sweetness
1                                 [bitter]    bitterness
2     [bitter, slightly bitter, non-sweet]    bitterness
3                                 [bitter]    bitterness
4                              [non-sweet]  non-sweetness
…                                        …            …
2939                              [bitter]    bitterness
2940                              [bitter]    bitterness
2941                              [bitter]    bitterness
2942                              [bitter]    bitterness
2943                               [sweet]     sweetness

                              Reference_(cod)/[pp]
0                              Bassoli2000_((+)-9)
```

```
1                                   Dagan-Wiener2019_(463)
2      Belitz2009_[35]; Dagan-Wiener2019_(751); Glase…
3                                   Dagan-Wiener2019_(587)
4                                      Spillane2009b_(51A)
…                                                    …
2939                               Dagan-Wiener2019_(1029)
2940                                      Iwamura1981_(35)
2941                                Dagan-Wiener2019_(878)
2942                                Dagan-Wiener2019_(638)
2943                                   Spillane2009b_(17H)

[2944 rows x 8 columns]
```

[ ]:
```
df.shape[0]
```

[ ]: 2944

[ ]:
```
# df.to_excel('/content/drive/MyDrive/taste_predict/Dataset/integrated_dataset.
↪xlsx')
```

[ ]:
```
len(get_vocab_taste(df["Taste"]))
```

[ ]: 44

[ ]:
```
#Let's explore our label space by building a co-occurence matrix
from itertools import combinations
n_taste = len(vocabt)
sorted_taste=sorted(taste_count, key=lambda x: taste_count[x])[::-1]
taste_matrix = np.zeros((n_taste, n_taste))
taste_index = {k: i for i, k in enumerate(sorted_taste)}
for tastes in df["Taste"]:
    for o1, o2 in combinations(tastes, 2):
        if o1 == o2: continue
        taste_matrix[taste_index[o1], taste_index[o2]] += 1
        taste_matrix[taste_index[o2], taste_index[o1]] += 1
```

[ ]:
```
# get the index of highest count
indices = np.dstack(np.unravel_index(np.argsort(taste_matrix.ravel()),␣
↪(n_taste, n_taste)))[0][::-1]
print('      Top 20 common odor association')
temp_dict=dict()
for idx in indices[:40:2]:
    temp_dict[sorted_taste[idx[0]] +"-"+␣
↪sorted_taste[idx[1]]]=taste_matrix[idx[0], idx[1]]
pd.DataFrame(temp_dict.items(),columns=["Label Pairs","Co-occurence Count"])
```

```
      Top 20 common odor association
```

[ ]:

|  | Label Pairs | Co-occurence Count |
|---|---|---|
| 0 | bitter-sweet | 113.0 |
| 1 | sweet-non-sweet | 56.0 |
| 2 | non-sweet-bitter | 39.0 |
| 3 | tasteless-sweet | 19.0 |
| 4 | bitter-burning | 13.0 |
| 5 | heating-and tingling | 12.0 |
| 6 | and tingling-heating | 12.0 |
| 7 | and tingling-pungent | 12.0 |
| 8 | bitter-sour | 11.0 |
| 9 | non-sweet-tasteless | 10.0 |
| 10 | very sweet-sweet | 10.0 |
| 11 | non-sweet-sour | 9.0 |
| 12 | sweet-sour | 9.0 |
| 13 | salty-bitter | 8.0 |
| 14 | salty-umami | 7.0 |
| 15 | pungent-bitter | 7.0 |
| 16 | sour-umami | 6.0 |
| 17 | umami-sour | 6.0 |
| 18 | bitter-slightly bitter | 5.0 |
| 19 | slightly sweet-sweet | 4.0 |

[ ]:

```
import seaborn as sns
plt.figure(figsize=(14,10))
# for i in range(100):
sns.heatmap(taste_matrix, vmax=5)
plt.xticks(range(len(sorted_taste)), sorted_taste,rotation='vertical')
plt.yticks(range(len(sorted_taste)), sorted_taste,rotation='horizontal')
plt.show()
```

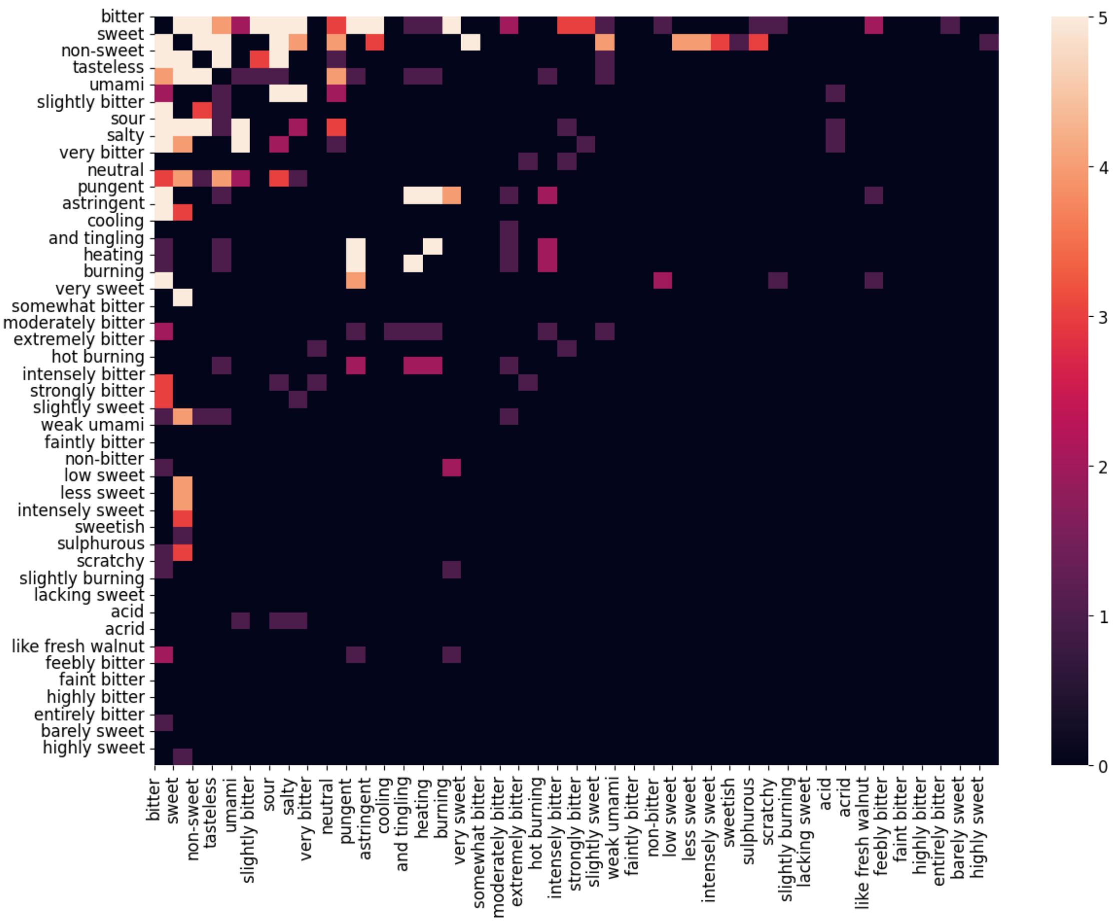


```
[ ]: import seaborn as sns
     plt.figure(figsize=(7,5))
     # for i in range(100):
     sns.heatmap(taste_matrix[:20,:20], vmax=5)
     plt.xticks(range(len(sorted_taste[:20])), sorted_taste[:20],rotation='vertical')
     plt.yticks(range(len(sorted_taste[:20])), sorted_taste[:
       ↪20],rotation='horizontal')
     plt.show()
```

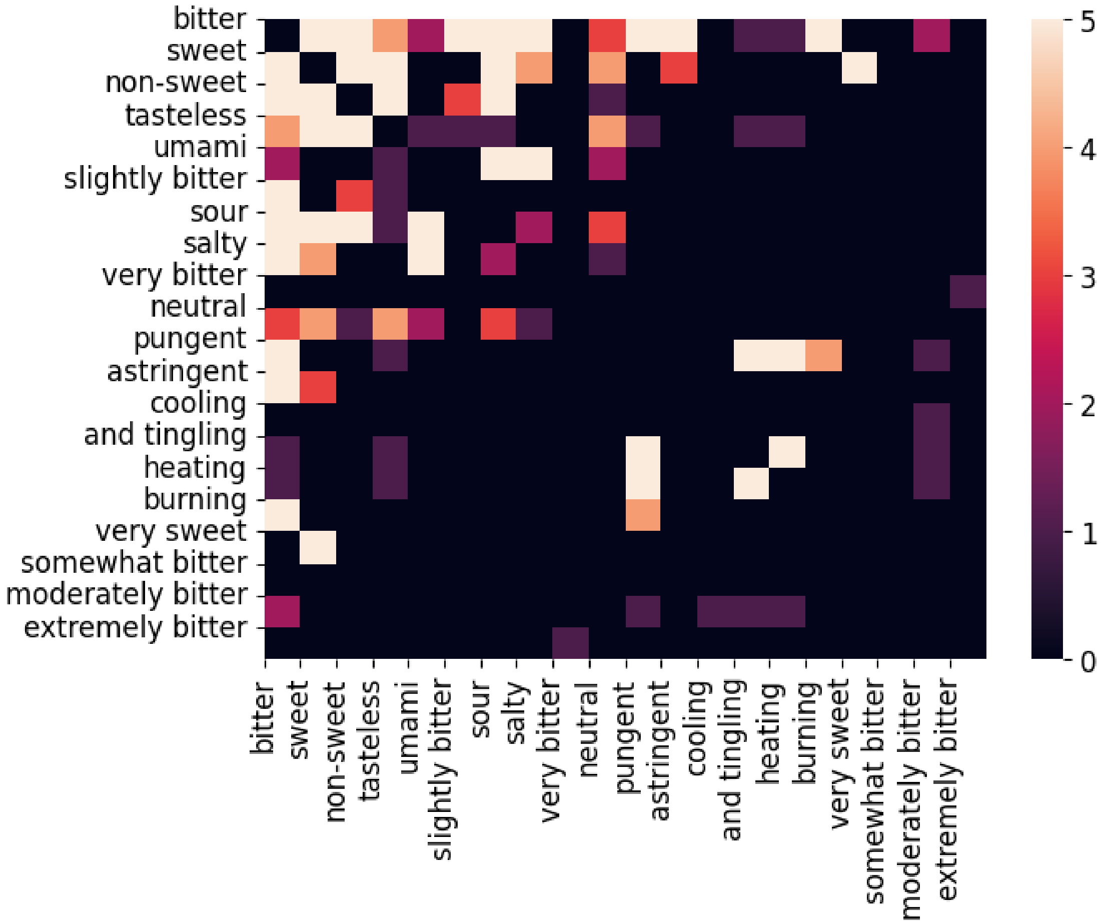


```
[ ]: plt.hist(df["Class taste"],bins=20)
     plt.xticks(rotation='vertical')
     plt.xlabel("Class Taste")
     plt.ylabel("No of molecules")
     plt.show()
```

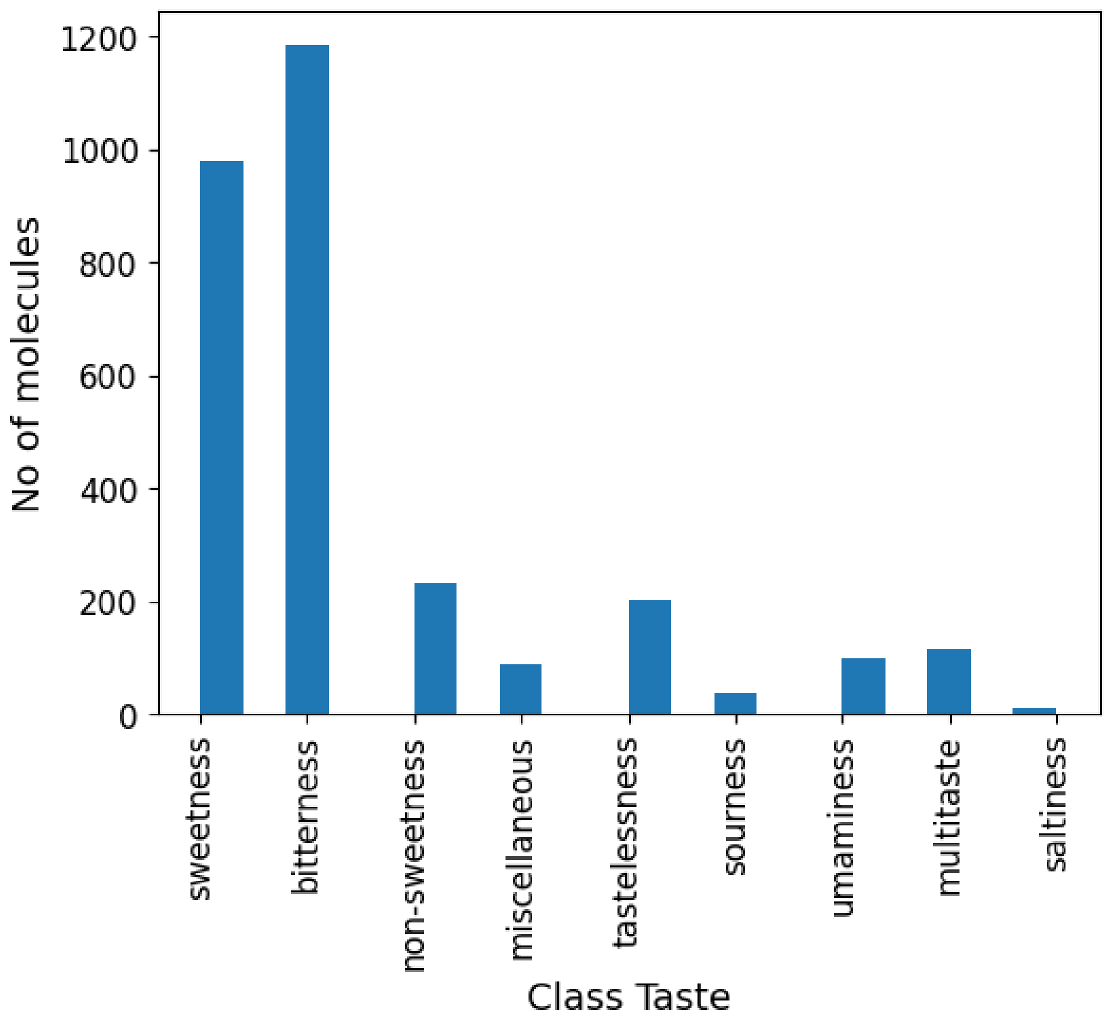


[ ]:
```
print(df.dtypes)  # Print the data types of all columns
```

```
ID                         object
Name                       object
PubChem CID                object
CAS number                 object
canonical SMILES           object
Taste                      object
Class taste                object
Reference_(cod)/[pp]       object
dtype: object
```

[ ]:
```
import pandas as pd
import seaborn as sns
import matplotlib.pyplot as plt
```

```
# Example feature engineering: length of SMILES string, count of commas in␣
↪Taste, etc.
df['SMILES length'] = df['canonical SMILES'].apply(lambda x: len(x) if pd.
↪notna(x) else 0)
# Modify the lambda function to handle lists and strings
df['Taste count'] = df['Taste'].apply(lambda x: len(x) if isinstance(x, list)␣
↪else (x.count(',') + 1 if isinstance(x, str) else 0))
# df['Class taste count'] = df['Class taste'].apply(lambda x: x.count(',') + 1␣
↪if pd.notna(x) else 0)

# Now proceed with the rest of the code
# Select the newly created numeric columns and add 'Class taste'
df_numeric_with_features = pd.concat([df[['SMILES length', 'Taste count']],␣
↪df['Class taste']], axis=1)


# Drop rows with NaN values in numeric columns
df_numeric_with_features = df_numeric_with_features.dropna()

# Create a pairplot with the engineered features
sns.pairplot(df_numeric_with_features, hue='Class taste')
plt.show()
```

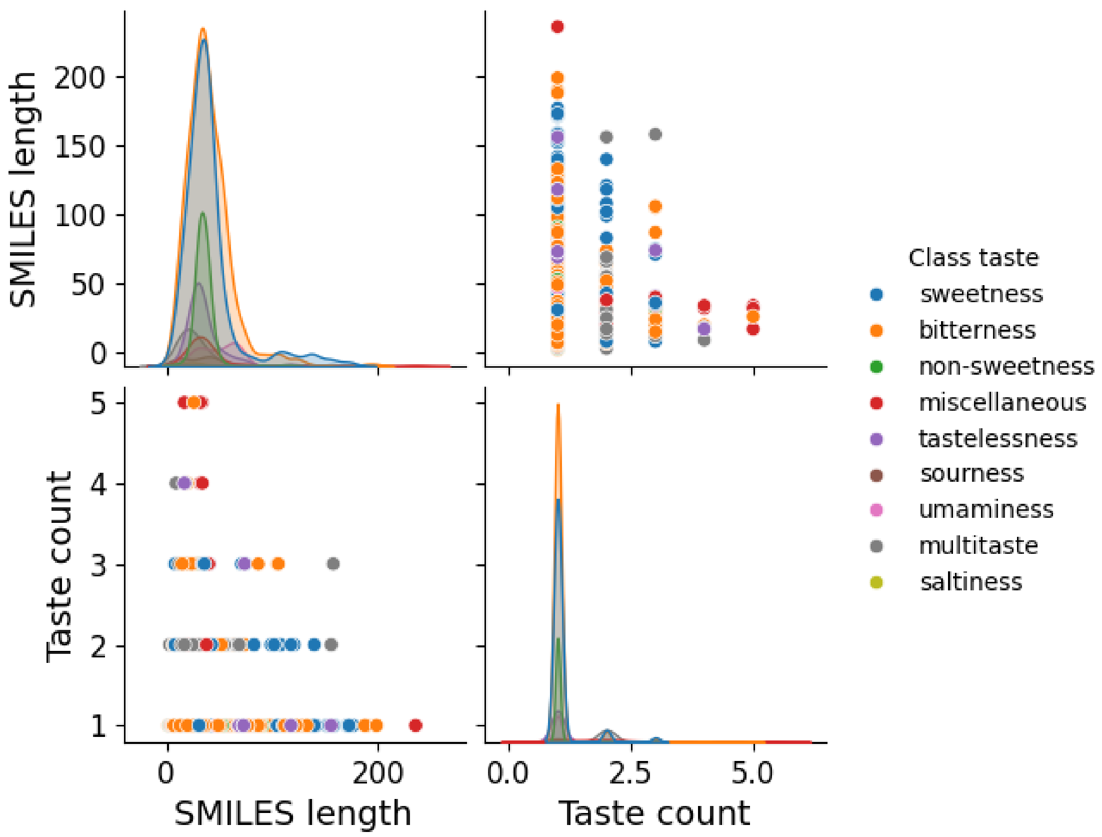


```
[ ]: df
```

```
[ ]:         ID                                           Name  PubChem CID  \
     0     0566                          Compound (R)-(+)-9            *
     1     1221                                   Artemisin        65030
     2     1678                                   L-Leucine         6106
     3     1578                             Hexethal sodium     23690440
     4     2832*  Sodium N-[3-chloro-5-(trifluoromethyl)phenyl]s…         *
     …     …                                              …          …
     2939  1639                               Isoxanthohumol M   122363515
     2940  1096*  3-Amino-3-{[1-(2H-1,3-benzodioxol-5-yl)propan-…         *
     2941  1131                          5,7,2'-Trihydroxyflavone      5322064
     2942  1295                                   Chlormerodrin        25210
     2943  0861*           Sodium N-(2-methylthian-4-yl)sulfamate            *

          CAS number                            canonical SMILES  \
     0            *          COc1ccc(cc1O)C1OC2CC3CCC2(CS1)C3(C)C
     1      481-05-0        CC1C2C(O)CC3(C)C=CC(=O)C(=C3C2OC1=O)C
     2       61-90-5                             CC(C)CC(N)C(O)=O
     3      144-00-3       [Na+].CCCCCCC1(CC)C(=O)NC(=NC1=O)[O-]
```

```
4              *         [Na+].[O-]S(=O)(=O)Nc1cc(cc(c1)C(F)(F)F)Cl
…          …                                                  …
2939           *  COc1cc(c(c2c1C(=O)CC(O2)c1ccc(cc1)O)CCC(C)(C)OC)O
2940           *            CC(Cc1ccc2c(c1)OCO2)NC(=O)C(N)CC(O)=O
2941  73046-40-9              Oc1cc(c2c(c1)OC(=CC2=O)c1ccccc1O)O
2942     62-37-3                           COC(CNC(N)=O)C[Hg]Cl
2943           *                   [Na+].CC1CC(CCS1)NS([O-])(=O)=O

                                     Taste    Class taste  \
0                                  [sweet]      sweetness
1                                 [bitter]     bitterness
2     [bitter, slightly bitter, non-sweet]     bitterness
3                                 [bitter]     bitterness
4                              [non-sweet]  non-sweetness
…                                      …            …
2939                              [bitter]     bitterness
2940                              [bitter]     bitterness
2941                              [bitter]     bitterness
2942                              [bitter]     bitterness
2943                               [sweet]      sweetness

                                   Reference_(cod)/[pp]  SMILES length  \
0                                    Bassoli2000_((+)-9)             36
1                                  Dagan-Wiener2019_(463)            37
2     Belitz2009_[35]; Dagan-Wiener2019_(751); Glase…              16
3                                  Dagan-Wiener2019_(587)            37
4                                    Spillane2009b_(51A)             42
…                                                   …            …
2939                              Dagan-Wiener2019_(1029)            49
2940                                    Iwamura1981_(35)             37
2941                               Dagan-Wiener2019_(878)            34
2942                               Dagan-Wiener2019_(638)            20
2943                                 Spillane2009b_(17H)             31

      Taste count
0               1
1               1
2               3
3               1
4               1
…             …
2939            1
2940            1
2941            1
2942            1
2943            1
```

[2944 rows x 10 columns]

[ ]:
```
sns.pairplot(df, hue='Taste count')
# plt.show()
```

[ ]: <seaborn.axisgrid.PairGrid at 0x787682d7e0e0>

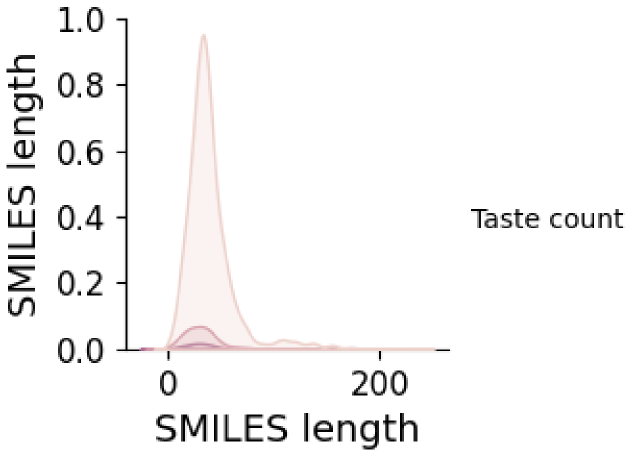


[ ]:
```
pip install mordred
```

```
Collecting mordred
  Downloading mordred-1.2.0.tar.gz (128 kB)
                               128.8/128.8

kB 3.1 MB/s eta 0:00:00
  Preparing metadata (setup.py) … done
Requirement already satisfied: six==1.* in /usr/local/lib/python3.10/dist-
packages (from mordred) (1.16.0)
Requirement already satisfied: numpy==1.* in /usr/local/lib/python3.10/dist-
packages (from mordred) (1.26.4)
Collecting networkx==2.* (from mordred)
  Downloading networkx-2.8.8-py3-none-any.whl.metadata (5.1 kB)
Downloading networkx-2.8.8-py3-none-any.whl (2.0 MB)
                            2.0/2.0 MB
30.7 MB/s eta 0:00:00
Building wheels for collected packages: mordred
  Building wheel for mordred (setup.py) … done
  Created wheel for mordred: filename=mordred-1.2.0-py3-none-any.whl size=176718
sha256=4e6aded03675fd4505d00d9a65d9e82e1e3811bb00dc8866599d168123776335
  Stored in directory: /root/.cache/pip/wheels/a7/4f/b8/d4c6591f6ac944aaced7865b
349477695f662388ad958743c7
Successfully built mordred
```

```
Installing collected packages: networkx, mordred
  Attempting uninstall: networkx
    Found existing installation: networkx 3.4.2
    Uninstalling networkx-3.4.2:
      Successfully uninstalled networkx-3.4.2
ERROR: pip's dependency resolver does not currently take into account all
the packages that are installed. This behaviour is the source of the following
dependency conflicts.
nx-cugraph-cu12 24.10.0 requires networkx>=3.0, but you have networkx 2.8.8
which is incompatible.
Successfully installed mordred-1.2.0 networkx-2.8.8
```

[ ]:
```
import mordred
from mordred import Calculator,descriptors
from rdkit import Chem, DataStructs
from rdkit.Chem import rdMolDescriptors, Draw
```

[ ]:
```
def mord(data):
  #Under the Rdkit package all the operations
  #are done on the molecule object which can be obtained by using the␣
↪Molfromsmiles method
  mol_data_set=[Chem.MolFromSmiles(x) for x in data["canonical SMILES"]]
  l=[]
  for x in mol_data_set:
    calc=Calculator(descriptors)
    new_feat=calc(x)
    new_dict=new_feat.drop_missing().asdict()
    l.append(new_dict)
  return pd.DataFrame(l)
```

[ ]:
```
def path_based_fing(df):
    # Convert SMILES strings into molecules
    mols = [Chem.rdmolfiles.MolFromSmiles(SMILES_string) for SMILES_string in␣
↪df["canonical SMILES"]]
    fps = [Chem.RDKFingerprint(m,fpSize=1024) for m in mols ]

    # Convert training fingerprints into binary
    np_fps = []
    for fp in fps:
      arr = np.zeros((1,), dtype= int)
      DataStructs.ConvertToNumpyArray(fp, arr)
      np_fps.append(arr)
    return pd.DataFrame(np_fps)
```

[ ]:
```
def morgan_fing(df):
    # Convert SMILES strings into molecules
    mols = [Chem.rdmolfiles.MolFromSmiles(SMILES_string) for SMILES_string in␣
  ↪df["canonical SMILES"]]
    fps = [rdMolDescriptors.GetMorganFingerprintAsBitVect(m, radius=2, bitInfo=␣
  ↪{}, nBits=1024, useChirality=True) for m in mols]

    np_fps = []
    for fp in fps:
      arr = np.zeros((1,), dtype= int)
      DataStructs.ConvertToNumpyArray(fp, arr)
      np_fps.append(arr)
    return pd.DataFrame(np_fps)
```

[ ]:
```
table.tail(10).sort_values(by="No of associated samples")
```

[ ]:
```
     Class Taste  No of associated samples  Percentage_of_samples
8      saltiness                        12               0.407609
7       sourness                        38               1.290761
6  miscellaneous                        87               2.955163
5      umaminess                        98               3.328804
4     multitaste                       113               3.838315
3  tastelessness                       203               6.895380
2  non-sweetness                       233               7.914402
1      sweetness                       977              33.186141
0     bitterness                      1183              40.183424
```

[ ]:
```
#Before we make the split, we need to binarize our target labels because the␣
  ↪implementation of
#the stratifiedsplit require the targed labels to be in a binarized format
from sklearn.preprocessing import MultiLabelBinarizer
enc=MultiLabelBinarizer()
y=enc.fit_transform(df["Taste"])
```

[ ]:
```
Y=pd.DataFrame(y,columns=enc.classes_)
```

[ ]:
```
df.head(100)
```

[ ]:
```
       ID                                                Name PubChem CID  \
0    0566                                  Compound (R)-(+)-9           *
1    1221                                           Artemisin       65030
2    1678                                           L-Leucine        6106
3    1578                                      Hexethal sodium    23690440
4   2832*  Sodium N-[3-chloro-5-(trifluoromethyl)phenyl]s…           *
..    …                                                    …         …
95   1898                                      Procyanidin C2    11182062
96   1836                                  Phenobarbital sodium    23674889
```

```
97   0522                                      Abrusoside D    44575936
98  0645^     Hexahydrothiepin-3-ylsulfamic acid sodium salt    23721056
99  1979^              Sodium N-(2,6-diethylphenyl)sulfamate    23674129

     CAS number                                   canonical SMILES  \
0             *              COc1ccc(cc1O)C1OC2CC3CCC2(CS1)C3(C)C
1      481-05-0             CC1C2C(O)CC3(C)C=CC(=O)C(=C3C2OC1=O)C
2       61-90-5                                  CC(C)CC(N)C(O)=O
3      144-00-3             [Na+].CCCCCCC1(CC)C(=O)NC(=NC1=O)[O-]
4             *        [Na+].[O-]S(=O)(=O)Nc1cc(cc(c1)C(F)(F)F)Cl
..          …                                                 …
95            *  OC1Cc2c(cc(c(c2OC1c1ccc(c(c1)O)O)C1C(O)C(Oc2c(…
96      57-30-7                    CCC1(C(=O)NC(=O)NC1=O)c1ccccc1
97  125003-00-1  CC(C1CC=C(C)C(=O)O1)C1CCC2(C)C3CCC4C(C)(C(CCC5…
98            *                     [Na+].[O-]S(=O)(=O)NC1CCCCSC1
99            *                 [Na+].CCc1cccc(c1NS([O-])(=O)=O)CC

                                  Taste    Class taste  \
0                               [sweet]      sweetness
1                              [bitter]     bitterness
2   [bitter, slightly bitter, non-sweet]     bitterness
3                              [bitter]     bitterness
4                           [non-sweet]  non-sweetness
..                                  …            …
95                             [bitter]     bitterness
96                   [slightly bitter]     bitterness
97                              [sweet]      sweetness
98                              [sweet]      sweetness
99                             [bitter]     bitterness

                               Reference_(cod)/[pp]  SMILES length  \
0                                Bassoli2000_((+)-9)             36
1                             Dagan-Wiener2019_(463)             37
2   Belitz2009_[35]; Dagan-Wiener2019_(751); Glase…           16
3                             Dagan-Wiener2019_(587)             37
4                                Spillane2009b_(51A)             42
..                                              …            …
95                           Dagan-Wiener2019_(1249)            117
96                            Dagan-Wiener2019_(385)             30
97  Bouysset2020_(172); Kinghorn1998_(13); Kinghor…          108
98             Spillane2000_(70); Spillane2009b_(11H)             29
99                                 Spillane2006_(72)             34

    Taste count
0             1
1             1
2             3
```

```
3              1
4              1
..           ...
95             1
96             1
97             1
98             1
99             1

[100 rows x 10 columns]
```

[ ]:
```
Y.head()
```

[ ]:
```
   acid  acrid  and tingling  astringent  barely sweet  bitter  burning  \
0     0      0             0           0             0       0        0
1     0      0             0           0             0       1        0
2     0      0             0           0             0       1        0
3     0      0             0           0             0       1        0
4     0      0             0           0             0       0        0

   cooling  entirely bitter  extremely bitter  ...  sour  strongly bitter  \
0        0                0                 0  ...     0                0
1        0                0                 0  ...     0                0
2        0                0                 0  ...     0                0
3        0                0                 0  ...     0                0
4        0                0                 0  ...     0                0

   sulphurous  sweet  sweetish  tasteless  umami  very bitter  very sweet  \
0           0      1         0          0      0            0           0
1           0      0         0          0      0            0           0
2           0      0         0          0      0            0           0
3           0      0         0          0      0            0           0
4           0      0         0          0      0            0           0

   weak umami
0           0
1           0
2           0
3           0
4           0

[5 rows x 44 columns]
```

[ ]:
```
tastes=enc.classes_
print(len(tastes), tastes)
```

```
44 ['acid' 'acrid' 'and tingling' 'astringent' 'barely sweet' 'bitter'
```

```
 'burning' 'cooling' 'entirely bitter' 'extremely bitter' 'faint bitter'
 'faintly bitter' 'feebly bitter' 'heating' 'highly bitter' 'highly sweet'
 'hot burning' 'intensely bitter' 'intensely sweet' 'lacking sweet'
 'less sweet' 'like fresh walnut' 'low sweet' 'moderately bitter'
 'neutral' 'non-bitter' 'non-sweet' 'pungent' 'salty' 'scratchy'
 'slightly bitter' 'slightly burning' 'slightly sweet' 'somewhat bitter'
 'sour' 'strongly bitter' 'sulphurous' 'sweet' 'sweetish' 'tasteless'
 'umami' 'very bitter' 'very sweet' 'weak umami']
```

[ ]:
```
pip install iterative-stratification
```

```
Collecting iterative-stratification
  Downloading iterative_stratification-0.1.9-py3-none-any.whl.metadata (1.3 kB)
Requirement already satisfied: numpy in /usr/local/lib/python3.10/dist-packages
(from iterative-stratification) (1.26.4)
Requirement already satisfied: scipy in /usr/local/lib/python3.10/dist-packages
(from iterative-stratification) (1.13.1)
Requirement already satisfied: scikit-learn in /usr/local/lib/python3.10/dist-
packages (from iterative-stratification) (1.5.2)
Requirement already satisfied: joblib>=1.2.0 in /usr/local/lib/python3.10/dist-
packages (from scikit-learn->iterative-stratification) (1.4.2)
Requirement already satisfied: threadpoolctl>=3.1.0 in
/usr/local/lib/python3.10/dist-packages (from scikit-learn->iterative-
stratification) (3.5.0)
Downloading iterative_stratification-0.1.9-py3-none-any.whl (8.5 kB)
Installing collected packages: iterative-stratification
Successfully installed iterative-stratification-0.1.9
```

[ ]:
```
import skmultilearn
from skmultilearn.model_selection import iterative_train_test_split
from sklearn.model_selection import train_test_split
from iterstrat.ml_stratifiers import MultilabelStratifiedKFold
#We do a hold out testing for our model validation, under stratified sampling.
```

[ ]:
```
#The main function we'll call for splitting the data in accordance with
#the algorithm in the paper cited above
def iterative_split(x,y,test_size):
  x=np.array(x)
  y=np.array(y)
  np.random.seed(42)
  return iterative_train_test_split(x,y,test_size=test_size)
```

[ ]:
```
#Let's visualize our split and compare it with a random split.
 #We test our split function using morgan featurization.
x_train,y_train,x_test,y_test=iterative_split(morgan_fing(df),Y,test_size=0.2)
```

```
[ ]: #This is a random split
     ␣
     ↪X_train,X_test,Y_train,Y_test=train_test_split(morgan_fing(df),Y,random_state=42,test_size=
     ↪2)
```

```
[ ]: print("y_train{}  ".format(y_train.shape) +"\n"+"y_test{}   ".format(y_test.
     ↪shape)+"\n"+"x_train{}  ".format(x_train.shape)+"\n"+"x_test{}  ".
     ↪format(x_test.shape))
```

```
y_train(585, 44)
y_test(2359, 44)
x_train(585, 1024)
x_test(2359, 1024)
```

```
[ ]: def label_distribution(y_train,y_test):
       voc= defaultdict(int)
       for arr in y_train:
         for i in range(44):
           if(arr[i]==1):
             voc[i]+=1
       plt.figure(figsize=(30,10))
       plt.bar(voc.keys(),[(i/len(y_train))*100 for i in voc.values()])
       voc= defaultdict(int)
       for arr in y_test:
         for i in range(44):
           if(arr[i]==1):
             voc[i]+=1
       plt.xlabel("Taste")
       plt.ylabel("Percentage of samples associated")
       plt.bar(voc.keys(),[(i/len(y_test))*100 for i in voc.values()])
       plt.legend(["train-set","test-set"])
       plt.show()
```

```
[ ]: label_distribution(np.array(Y_train),np.array(Y_test))#label distribution while␣
     ↪doing a random split
```

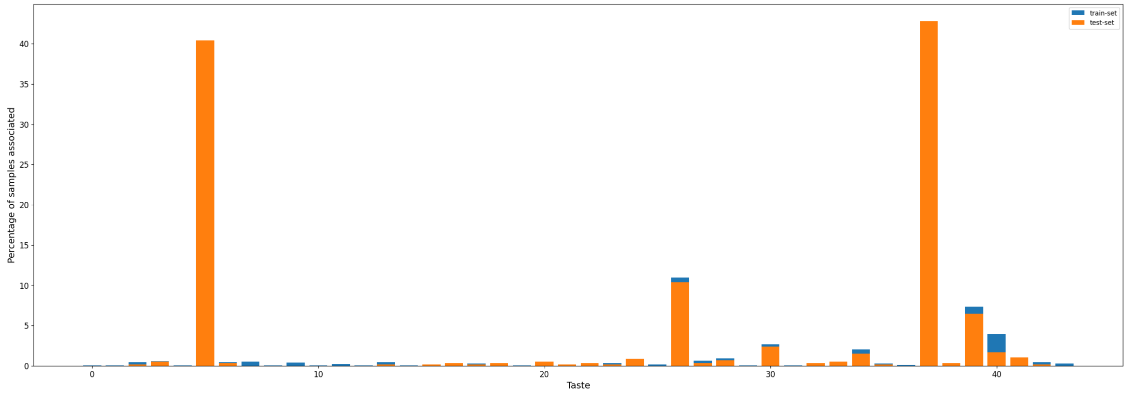

[ ]:
```
label_distribution(y_train,y_test)# label distribution while doing a stratified␣
↪split
```

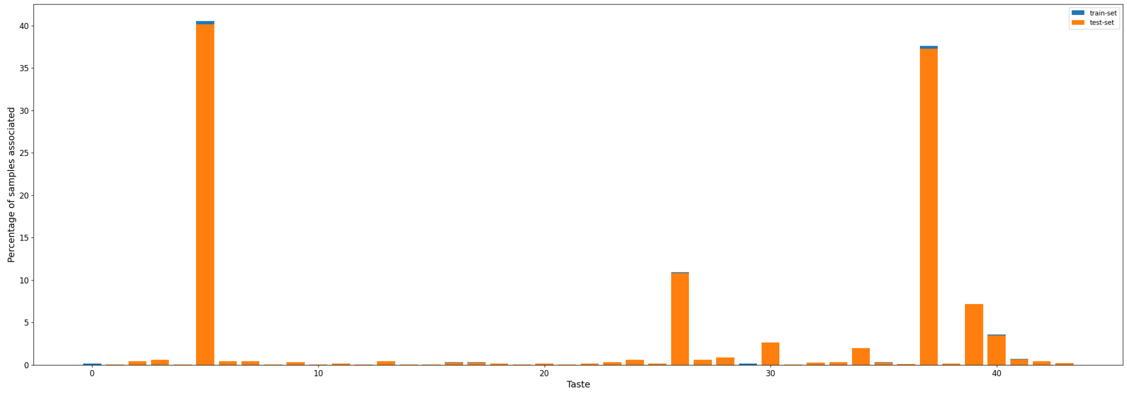


[ ]:
```
!pip install python-louvain
```

```
Requirement already satisfied: python-louvain in /usr/local/lib/python3.10/dist-
packages (0.16)
Requirement already satisfied: networkx in /usr/local/lib/python3.10/dist-
packages (from python-louvain) (2.8.8)
Requirement already satisfied: numpy in /usr/local/lib/python3.10/dist-packages
(from python-louvain) (1.26.4)
```

[ ]:
```
import community.community_louvain as community_louvain
print(dir(community_louvain))  # Should list the available attributes including␣
↪'best_partition'
```

```
['Status', '__MIN', '__PASS_MAX', '__author__', '__builtins__', '__cached__',
'__doc__', '__file__', '__insert', '__loader__', '__modularity', '__name__',
'__neighcom', '__one_level', '__package__', '__randomize', '__remove',
'__renumber', '__spec__', 'array', 'best_partition', 'check_random_state',
'generate_dendrogram', 'induced_graph', 'load_binary', 'modularity', 'np',
'numbers', 'nx', 'partition_at_level', 'print_function', 'warnings']
```

[ ]:
```
from skmultilearn.cluster import LabelCooccurrenceGraphBuilder,␣
↪NetworkXLabelGraphClusterer
import community.community_louvain as community_louvain
import networkx as nx
import numpy as np

class CustomNetworkXLabelGraphClusterer(NetworkXLabelGraphClusterer):
    def fit_predict(self, X, y):
```

```
        # Create the graph using the graph builder and get the co-occurrence␣
↪dictionary
        cooccurrence_dict = self.graph_builder.transform(y)

        print("Co-occurrence Dictionary:")
        print(cooccurrence_dict)

        # Check if cooccurrence_dict is a dictionary
        if not isinstance(cooccurrence_dict, dict):
            raise ValueError("Invalid co-occurrence matrix format. Expected a␣
↪dictionary of edge weights.")

        # Create a NetworkX graph
        self.graph_ = nx.Graph()

        # Add edges and weights to the graph
        for (node1, node2), weight in cooccurrence_dict.items():
            self.graph_.add_edge(node1, node2, weight=weight)

        if self.method == 'louvain':
            # Now pass the NetworkX graph to community_louvain.best_partition
            partition_dict = community_louvain.best_partition(self.graph_)
            print("Partition Dict:")  # Debugging line
            print(partition_dict)  # Check if this is as expected

            # Calculate number of labels based on maximum node index
            num_labels = max(partition_dict.keys()) + 1
            print("Number of Labels:", num_labels)

            # Construct memberships based on the partition dictionary
            memberships = [partition_dict.get(i, -1) for i in range(num_labels)]
            print("Memberships Vector:", memberships)  # Debugging line

        else:
            raise NotImplementedError("Only Louvain method is currently␣
↪supported.")

        return memberships

# Adjusted to convert list of cluster memberships into a dictionary format
def to_membership_vector(memberships):
    # Create a dictionary where each node is assigned to its cluster
    return {i: memberships[i] for i in range(len(memberships))}

# Create the graph builder
graph_builder = LabelCooccurrenceGraphBuilder(weighted=True,␣
↪include_self_edges=False)
```

```
# Initialize the custom clusterer with the graph builder
clusterer = CustomNetworkXLabelGraphClusterer(graph_builder=graph_builder,␣
↪method='louvain')

# Fit and predict clusters
try:
    partition = clusterer.fit_predict(x_train, y_train)

    # Use the adjusted function to handle memberships format
    membership_vector = to_membership_vector(partition)
    print("Membership Vector Dictionary:", membership_vector)
except AttributeError as e:
    print(f"Attribute Error: {e}")
except ValueError as e:
    print(f"Value Error: {e}")
except TypeError as e:
    print(f"Type Error: {e}")
except Exception as e:
    print(f"Unexpected Error: {e}")
```

```
Co-occurrence Dictionary:
{(5, 26): 8.0, (5, 30): 1.0, (26, 30): 1.0, (20, 37): 1.0, (34, 37): 2.0, (5,
37): 21.0, (37, 42): 2.0, (26, 37): 9.0, (26, 34): 2.0, (0, 28): 1.0, (0, 34):
1.0, (28, 34): 1.0, (0, 40): 1.0, (28, 40): 1.0, (34, 40): 1.0, (26, 39): 3.0,
(37, 39): 4.0, (5, 17): 1.0, (5, 6): 2.0, (5, 27): 1.0, (6, 27): 1.0, (5, 34):
2.0, (5, 29): 1.0, (6, 29): 1.0, (24, 39): 1.0, (2, 13): 2.0, (2, 27): 2.0, (13,
27): 2.0, (32, 37): 1.0, (5, 28): 1.0, (2, 16): 1.0, (13, 16): 1.0, (16, 27):
1.0, (2, 39): 1.0, (13, 39): 1.0, (16, 39): 1.0, (27, 39): 1.0, (22, 37): 1.0,
(5, 39): 1.0, (5, 24): 1.0, (24, 37): 1.0, (5, 35): 1.0, (3, 5): 1.0, (18, 37):
1.0, (24, 34): 1.0}
Partition Dict:
{5: 1, 26: 1, 30: 1, 20: 1, 37: 1, 34: 3, 42: 1, 0: 3, 28: 3, 40: 3, 39: 0, 17:
1, 6: 2, 27: 0, 29: 2, 24: 3, 2: 0, 13: 0, 32: 1, 16: 0, 22: 1, 35: 1, 3: 1, 18:
1}
Number of Labels: 43
Memberships Vector: [3, -1, 0, 1, -1, 1, 2, -1, -1, -1, -1, -1, -1, 0, -1, -1,
0, 1, 1, -1, 1, -1, 1, -1, 3, -1, 1, 0, 3, 2, 1, -1, 1, -1, 3, 1, -1, 1, -1, 0,
3, -1, 1]
Membership Vector Dictionary: {0: 3, 1: -1, 2: 0, 3: 1, 4: -1, 5: 1, 6: 2, 7:
-1, 8: -1, 9: -1, 10: -1, 11: -1, 12: -1, 13: 0, 14: -1, 15: -1, 16: 0, 17: 1,
18: 1, 19: -1, 20: 1, 21: -1, 22: 1, 23: -1, 24: 3, 25: -1, 26: 1, 27: 0, 28: 3,
29: 2, 30: 1, 31: -1, 32: 1, 33: -1, 34: 3, 35: 1, 36: -1, 37: 1, 38: -1, 39: 0,
40: 3, 41: -1, 42: 1}
```

[ ]:
```
# import numpy as np
# import community.community_louvain as community_louvain
```

```
# from skmultilearn.cluster import LabelCooccurrenceGraphBuilder,␣
↪NetworkXLabelGraphClusterer

# # Create the graph builder
# graph_builder = LabelCooccurrenceGraphBuilder(weighted=True,␣
↪include_self_edges=False)

# # Generate edges based on label co-occurrence
# label_names = [i for i in range(44)]
# edge_map = graph_builder.transform(np.array(y_train))
# print("{} labels, {} edges".format(len(label_names), len(edge_map)))

# # Define a helper function for visualization purposes
# def to_membership_vector(partition):
#     return {
#         member: partition_id
#         for partition_id, members in enumerate(partition)
#         for member in members
#     }

# # Initialize the clusterer with the graph builder
# clusterer = NetworkXLabelGraphClusterer(graph_builder=graph_builder,␣
↪method='louvain')

# # Fit and predict clusters
# try:
#     partition = clusterer.fit_predict(x_train, y_train)
#     membership_vector = to_membership_vector(partition)
#     print(membership_vector)
# except AttributeError as e:
#     print(f"Error: {e}")
#     print("It seems there is an issue with the community library. Please␣
↪check the installation.")
```

[ ]:
```
# from skmultilearn.cluster import LabelCooccurrenceGraphBuilder
# graph_builder = LabelCooccurrenceGraphBuilder(weighted=True,
#                                                include_self_edges=False)

# label_names=[i for i in range(44)]
# edge_map = graph_builder.transform(np.array(y_train))
# print("{} labels, {} edges".format(len(label_names), len(edge_map)))
```

[ ]:
```
# from skmultilearn.cluster import NetworkXLabelGraphClusterer

# # we define a helper function for visualization purposes
# def to_membership_vector(partition):
#     return {
```

```
#         member :  partition_id
#         for partition_id, members in enumerate(partition)
#         for member in members
#     }
# clusterer = NetworkXLabelGraphClusterer(graph_builder, method='louvain')
```

[ ]:
```
# partition = clusterer.fit_predict(x_train,y_train)
```

[ ]:
```
membership_vector = to_membership_vector(partition)
print('There are', len(partition),'clusters')
```

```
There are 43 clusters
```

[ ]:
```
import numpy as np
import networkx as nx
import matplotlib.pyplot as plt
from skmultilearn.cluster import LabelCooccurrenceGraphBuilder,␣
 ↪NetworkXLabelGraphClusterer
import community.community_louvain as community_louvain

class CustomNetworkXLabelGraphClusterer(NetworkXLabelGraphClusterer):
    def fit_predict(self, X, y):
        # Create the graph from the labels
        co_occurrence_graph = self.graph_builder.transform(y)

        # Convert co-occurrence dictionary to a NetworkX graph
        self.graph_ = nx.Graph()  # Initialize an empty graph
        for (u, v), weight in co_occurrence_graph.items():
            self.graph_.add_edge(u, v, weight=weight)  # Add edges with weights

        # Calculate weights based on the graph edges
        self.weights_ = {'weight': np.array([self.graph_[u][v]['weight'] for u,␣
 ↪v in self.graph_.edges()])}

        if self.method == 'louvain':
            partition_dict = community_louvain.best_partition(self.graph_)
            memberships = [partition_dict.get(i, -1) for i in range(y.shape[1])]

        else:
            raise NotImplementedError("Only Louvain method is currently␣
 ↪supported.")

        return memberships

# Create the graph builder
graph_builder = LabelCooccurrenceGraphBuilder(weighted=True,␣
 ↪include_self_edges=False)
```

```python
# Initialize the custom clusterer with the graph builder
clusterer = CustomNetworkXLabelGraphClusterer(graph_builder=graph_builder, method='louvain')

# Fit and predict clusters
partition = clusterer.fit_predict(x_train, y_train)
membership_vector = to_membership_vector(partition)

# Create the graph again (to ensure it's available for visualization)
co_occurrence_graph = clusterer.graph_builder.transform(y_train)
clusterer.graph_ = nx.Graph()  # Initialize an empty graph
for (u, v), weight in co_occurrence_graph.items():
    clusterer.graph_.add_edge(u, v, weight=weight)  # Add edges with weights

# Create a mapping of node indices to colors based on membership_vector
node_color = [membership_vector[i] for i in range(len(clusterer.graph_.nodes()))]

# Create a mapping from node to label
names_dict = {node: tastes[node] for node in clusterer.graph_.nodes() if node < len(tastes)}

# print(names_dict)

plt.figure(1, figsize=(15, 9.5))
nx.draw(
    clusterer.graph_,
    pos=nx.spring_layout(clusterer.graph_, k=4),
    labels=names_dict,
    with_labels=True,
    width=[0.1 * clusterer.graph_[u][v]['weight'] for u, v in clusterer.graph_.edges()],  # Use the computed weights
    node_color=node_color,  # Use the node colors that correspond to the graph nodes
    cmap=plt.cm.viridis,
    node_size=1000,
    font_size=12,
    font_color='black',
    alpha=0.7
)
# plt.title("Label Co-occurrence Graph with Clusters")
plt.show()
```

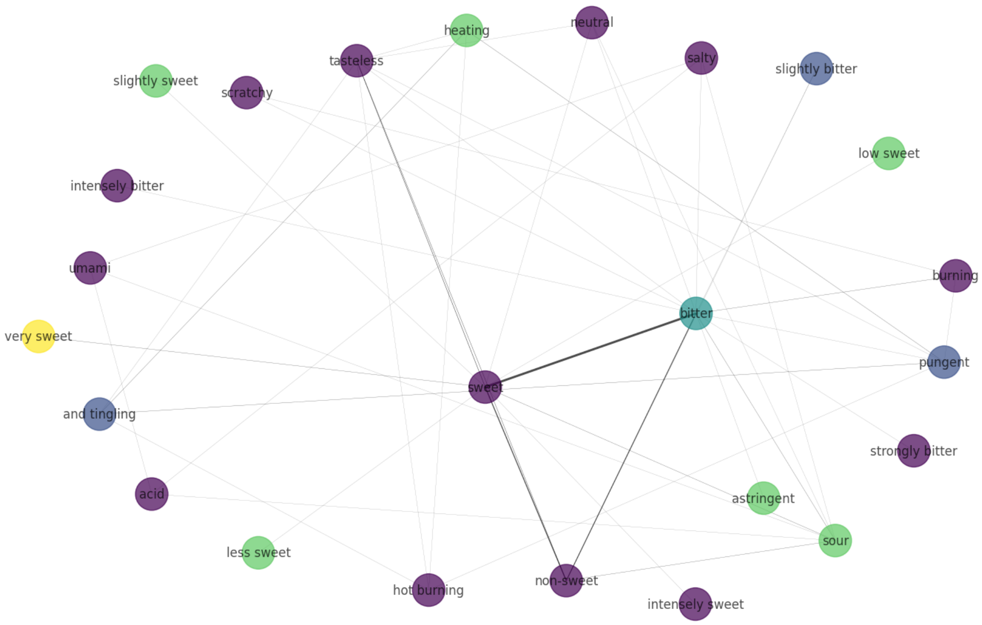


```
[ ]: # import networkx as nx
     # names_dict = dict(enumerate(x for x in classes))
     # import matplotlib.pyplot as plt
     # %matplotlib inline
     # plt.figure(1,figsize=(15,9.5))
     # nx.draw(
     #     clusterer.graph_,
     #     pos=nx.spring_layout(clusterer.graph_,k=4),
     #     labels=names_dict,
     #     with_labels = True,
     #     width = [10*x/y_train.shape[0] for x in clusterer.weights_['weight']],
     #     node_color = [membership_vector[i] for i in range(y_train.shape[1])],
     #     cmap=plt.cm.viridis,
     #     node_size=1000,
     #     font_size=12,
     #     font_color='black',
     #     alpha=0.7
     # )
     # plt.show()
```

```
[ ]: x_path=path_based_fing(df)
     x_path.head()
```

[ ]:
```
   0     1     2     3     4     5     6     7     8     9     …  1014  \
0     1     1     1     0     0     0     1     0     1     1  …     1
1     1     0     1     0     1     1     1     0     1     0  …     1
2     0     0     0     0     0     0     0     0     0     0  …     0
3     0     0     1     0     0     0     1     0     0     0  …     0
4     1     0     1     1     0     0     1     0     0     0  …     0

   1015  1016  1017  1018  1019  1020  1021  1022  1023
0     1     1     1     1     0     0     0     0     0
1     1     0     1     0     1     1     1     1     1
2     0     0     0     0     0     0     0     0     1
3     0     0     1     1     0     1     1     0     1
4     1     0     1     0     0     0     0     0     0

[5 rows x 1024 columns]
```

[ ]:
```
x_morg=morgan_fing(df)
x_morg.head()
```

[ ]:
```
   0     1     2     3     4     5     6     7     8     9     …  1014  \
0     0     0     0     0     0     0     0     0     0     0  …     0
1     0     0     0     0     0     0     0     0     0     0  …     0
2     0     1     0     0     0     0     0     0     0     0  …     0
3     0     0     0     0     0     0     0     0     0     0  …     0
4     0     0     0     0     0     0     0     0     0     0  …     0

   1015  1016  1017  1018  1019  1020  1021  1022  1023
0     0     0     0     0     1     0     0     0     0
1     0     0     0     0     1     0     0     0     0
2     0     0     0     0     0     0     0     0     0
3     1     0     0     0     0     0     0     0     0
4     0     0     0     0     0     0     0     0     0

[5 rows x 1024 columns]
```

[ ]:
```
x_mord = mord(df)
x_mord.head()
```

[ ]:
```
   nAcid  nBase    SpAbs_A   SpMax_A  SpDiam_A     SpAD_A   SpMAD_A   LogEE_A  \
0      0      0  28.786209  2.668412  5.136990  28.786209  1.308464  4.071927
1      0      0  23.553633  2.596372  5.136815  23.553633  1.239665  3.910657
2      1      1   9.335326  2.116883  4.233766   9.335326  1.037258  3.028326
3      1      3        NaN       NaN       NaN        NaN       NaN       NaN
4      1      1        NaN       NaN       NaN        NaN       NaN       NaN

     VE1_A     VE2_A  …  Zagreb2  mZagreb1  mZagreb2   MDEO-11  MDEN-22  \
0  3.634632  0.165211  …    165.0  7.291667  4.618056       NaN      NaN
```

```
1  3.949251  0.207855  ...    138.0  7.951389  3.944444  0.479438       NaN
2  2.774515  0.308279  ...     36.0  5.583333  2.111111  0.500000       NaN
3       NaN       NaN  ...     93.0       NaN  4.083333  0.750000       0.5
4       NaN       NaN  ...     89.0       NaN  3.208333  1.500000       NaN

   MDEN-12  MDEN-23  MDEN-13  MDEN-33  MDEN-11
0      NaN      NaN      NaN      NaN      NaN
1      NaN      NaN      NaN      NaN      NaN
2      NaN      NaN      NaN      NaN      NaN
3      NaN      NaN      NaN      NaN      NaN
4      NaN      NaN      NaN      NaN      NaN

[5 rows x 1262 columns]
```

[ ]:
```
x_mord=x_mord.astype(float)
```

[ ]:
```
x_mord.shape
```

[ ]:
```
(2944, 1262)
```

[ ]:
```
l=dict(x_mord.isna().sum(axis=0))#Feature wise
m=dict(x_mord.isna().sum(axis=1))#sample wise
```

[ ]:
```
x=sorted(l.items(),key=lambda x:x[1],reverse=True)
y=sorted(m.items(),key=lambda x:x[1],reverse=True)
print(x)
print(y)
```

```
[('MDEN-11', 2881), ('MDEN-13', 2870), ('MDEN-33', 2836), ('MDEN-23', 2684),
('MDEC-44', 2635), ('MDEN-12', 2451), ('MDEN-22', 2412), ('MDEC-14', 2371),
('MDEC-24', 2231), ('MDEC-34', 2224), ('MDEO-22', 2152), ('MDEC-11', 1714),
('MDEO-12', 1406), ('MDEC-12', 994), ('MDEC-13', 989), ('AATS8s', 837),
('AATSC8s', 837), ('MATS8s', 837), ('GATS8s', 837), ('AATS7s', 727), ('AATSC7s',
727), ('MATS7s', 727), ('GATS7s', 727), ('AATS6s', 641), ('AATSC6s', 641),
('MATS6s', 641), ('GATS6s', 641), ('BCUTc-1h', 624), ('BCUTc-1l', 624),
('SpAbs_A', 623), ('SpMax_A', 623), ('SpDiam_A', 623), ('SpAD_A', 623),
('SpMAD_A', 623), ('LogEE_A', 623), ('VE1_A', 623), ('VE2_A', 623), ('VE3_A',
623), ('VR1_A', 623), ('VR2_A', 623), ('VR3_A', 623), ('BCUTdv-1h', 623),
('BCUTdv-1l', 623), ('BCUTd-1h', 623), ('BCUTd-1l', 623), ('BCUTs-1h', 623),
('BCUTs-1l', 623), ('BCUTZ-1h', 623), ('BCUTZ-1l', 623), ('BCUTm-1h', 623),
('BCUTm-1l', 623), ('BCUTv-1h', 623), ('BCUTv-1l', 623), ('BCUTse-1h', 623),
('BCUTse-1l', 623), ('BCUTpe-1h', 623), ('BCUTpe-1l', 623), ('BCUTare-1h', 623),
('BCUTare-1l', 623), ('BCUTp-1h', 623), ('BCUTp-1l', 623), ('BCUTi-1h', 623),
('BCUTi-1l', 623), ('SpAbs_DzZ', 623), ('SpMax_DzZ', 623), ('SpDiam_DzZ', 623),
('SpAD_DzZ', 623), ('SpMAD_DzZ', 623), ('LogEE_DzZ', 623), ('SM1_DzZ', 623),
('VE1_DzZ', 623), ('VE2_DzZ', 623), ('VE3_DzZ', 623), ('VR1_DzZ', 623),
('VR2_DzZ', 623), ('VR3_DzZ', 623), ('SpAbs_Dzm', 623), ('SpMax_Dzm', 623),
```

('SpDiam_Dzm', 623), ('SpAD_Dzm', 623), ('SpMAD_Dzm', 623), ('LogEE_Dzm', 623), ('SM1_Dzm', 623), ('VE1_Dzm', 623), ('VE2_Dzm', 623), ('VE3_Dzm', 623), ('VR1_Dzm', 623), ('VR2_Dzm', 623), ('VR3_Dzm', 623), ('SpAbs_Dzv', 623), ('SpMax_Dzv', 623), ('SpDiam_Dzv', 623), ('SpAD_Dzv', 623), ('SpMAD_Dzv', 623), ('LogEE_Dzv', 623), ('SM1_Dzv', 623), ('VE1_Dzv', 623), ('VE2_Dzv', 623), ('VE3_Dzv', 623), ('VR1_Dzv', 623), ('VR2_Dzv', 623), ('VR3_Dzv', 623), ('SpAbs_Dzse', 623), ('SpMax_Dzse', 623), ('SpDiam_Dzse', 623), ('SpAD_Dzse', 623), ('SpMAD_Dzse', 623), ('LogEE_Dzse', 623), ('SM1_Dzse', 623), ('VE1_Dzse', 623), ('VE2_Dzse', 623), ('VE3_Dzse', 623), ('VR1_Dzse', 623), ('VR2_Dzse', 623), ('VR3_Dzse', 623), ('SpAbs_Dzpe', 623), ('SpMax_Dzpe', 623), ('SpDiam_Dzpe', 623), ('SpAD_Dzpe', 623), ('SpMAD_Dzpe', 623), ('LogEE_Dzpe', 623), ('SM1_Dzpe', 623), ('VE1_Dzpe', 623), ('VE2_Dzpe', 623), ('VE3_Dzpe', 623), ('VR1_Dzpe', 623), ('VR2_Dzpe', 623), ('VR3_Dzpe', 623), ('SpAbs_Dzare', 623), ('SpMax_Dzare', 623), ('SpDiam_Dzare', 623), ('SpAD_Dzare', 623), ('SpMAD_Dzare', 623), ('LogEE_Dzare', 623), ('SM1_Dzare', 623), ('VE1_Dzare', 623), ('VE2_Dzare', 623), ('VE3_Dzare', 623), ('VR1_Dzare', 623), ('VR2_Dzare', 623), ('VR3_Dzare', 623), ('SpAbs_Dzp', 623), ('SpMax_Dzp', 623), ('SpDiam_Dzp', 623), ('SpAD_Dzp', 623), ('SpMAD_Dzp', 623), ('LogEE_Dzp', 623), ('SM1_Dzp', 623), ('VE1_Dzp', 623), ('VE2_Dzp', 623), ('VE3_Dzp', 623), ('VR1_Dzp', 623), ('VR2_Dzp', 623), ('VR3_Dzp', 623), ('SpAbs_Dzi', 623), ('SpMax_Dzi', 623), ('SpDiam_Dzi', 623), ('SpAD_Dzi', 623), ('SpMAD_Dzi', 623), ('LogEE_Dzi', 623), ('SM1_Dzi', 623), ('VE1_Dzi', 623), ('VE2_Dzi', 623), ('VE3_Dzi', 623), ('VR1_Dzi', 623), ('VR2_Dzi', 623), ('VR3_Dzi', 623), ('SpAbs_D', 623), ('SpMax_D', 623), ('SpDiam_D', 623), ('SpAD_D', 623), ('SpMAD_D', 623), ('LogEE_D', 623), ('VE1_D', 623), ('VE2_D', 623), ('VE3_D', 623), ('VR1_D', 623), ('VR2_D', 623), ('VR3_D', 623), ('ETA_alpha', 623), ('AETA_alpha', 623), ('ETA_shape_p', 623), ('ETA_shape_y', 623), ('ETA_shape_x', 623), ('ETA_beta', 623), ('AETA_beta', 623), ('ETA_beta_s', 623), ('AETA_beta_s', 623), ('ETA_beta_ns', 623), ('AETA_beta_ns', 623), ('ETA_beta_ns_d', 623), ('AETA_beta_ns_d', 623), ('ETA_eta', 623), ('AETA_eta', 623), ('ETA_eta_L', 623), ('AETA_eta_L', 623), ('ETA_eta_R', 623), ('AETA_eta_R', 623), ('ETA_eta_RL', 623), ('AETA_eta_RL', 623), ('ETA_eta_F', 623), ('AETA_eta_F', 623), ('ETA_eta_FL', 623), ('AETA_eta_FL', 623), ('ETA_eta_B', 623), ('AETA_eta_B', 623), ('ETA_eta_BR', 623), ('AETA_eta_BR', 623), ('ETA_dAlpha_A', 623), ('ETA_dAlpha_B', 623), ('ETA_epsilon_1', 623), ('ETA_epsilon_2', 623), ('ETA_epsilon_3', 623), ('ETA_epsilon_4', 623), ('ETA_epsilon_5', 623), ('ETA_dEpsilon_A', 623), ('ETA_dEpsilon_B', 623), ('ETA_dEpsilon_C', 623), ('ETA_dEpsilon_D', 623), ('ETA_dBeta', 623), ('AETA_dBeta', 623), ('ETA_psi_1', 623), ('ETA_dPsi_A', 623), ('ETA_dPsi_B', 623), ('MID', 623), ('AMID', 623), ('MID_h', 623), ('AMID_h', 623), ('MID_C', 623), ('AMID_C', 623), ('MID_N', 623), ('AMID_N', 623), ('MID_O', 623), ('AMID_O', 623), ('MID_X', 623), ('AMID_X', 623), ('AATS5s', 608), ('AATSC5s', 608), ('MATS5s', 608), ('GATS5s', 608), ('AATS4s', 593), ('AATSC4s', 593), ('MATS4s', 593), ('GATS4s', 593), ('AATS3s', 590), ('AATSC3s', 590), ('MATS3s', 590), ('GATS3s', 590), ('ATS0s', 587), ('ATS1s', 587), ('ATS2s', 587), ('ATS3s', 587), ('ATS4s', 587), ('ATS5s', 587), ('ATS6s', 587), ('ATS7s', 587), ('ATS8s', 587), ('AATS0s', 587), ('AATS1s', 587), ('AATS2s', 587), ('ATSC0s', 587), ('ATSC1s', 587), ('ATSC2s', 587), ('ATSC3s', 587), ('ATSC4s', 587), ('ATSC5s', 587), ('ATSC6s', 587),

('ATSC7s', 587), ('ATSC8s', 587), ('AATSC0s', 587), ('AATSC1s', 587), ('AATSC2s', 587), ('MATS1s', 587), ('MATS2s', 587), ('GATS1s', 587), ('GATS2s', 587), ('Xp-0d', 587), ('AXp-0d', 587), ('mZagreb1', 587), ('MDEO-11', 541), ('Vabc', 510), ('Xp-0dv', 502), ('AXp-0dv', 502), ('AATSC8c', 481), ('MATS8c', 481), ('GATS8c', 481), ('AATS8dv', 479), ('AATS8d', 479), ('AATS8Z', 479), ('AATS8m', 479), ('AATS8v', 479), ('AATS8se', 479), ('AATS8pe', 479), ('AATS8are', 479), ('AATS8p', 479), ('AATS8i', 479), ('AATSC8dv', 479), ('AATSC8d', 479), ('AATSC8Z', 479), ('AATSC8m', 479), ('AATSC8v', 479), ('AATSC8se', 479), ('AATSC8pe', 479), ('AATSC8are', 479), ('AATSC8p', 479), ('AATSC8i', 479), ('MATS8dv', 479), ('MATS8d', 479), ('MATS8Z', 479), ('MATS8m', 479), ('MATS8v', 479), ('MATS8se', 479), ('MATS8pe', 479), ('MATS8are', 479), ('MATS8p', 479), ('MATS8i', 479), ('GATS8dv', 479), ('GATS8d', 479), ('GATS8Z', 479), ('GATS8m', 479), ('GATS8v', 479), ('GATS8se', 479), ('GATS8pe', 479), ('GATS8are', 479), ('GATS8p', 479), ('GATS8i', 479), ('MDEC-33', 249), ('AXp-7d', 233), ('AXp-7dv', 233), ('MDEC-22', 223), ('AATSC7c', 210), ('MATS7c', 210), ('GATS7c', 210), ('AATS7dv', 208), ('AATS7d', 208), ('AATS7Z', 208), ('AATS7m', 208), ('AATS7v', 208), ('AATS7se', 208), ('AATS7pe', 208), ('AATS7are', 208), ('AATS7p', 208), ('AATS7i', 208), ('AATSC7dv', 208), ('AATSC7d', 208), ('AATSC7Z', 208), ('AATSC7m', 208), ('AATSC7v', 208), ('AATSC7se', 208), ('AATSC7pe', 208), ('AATSC7are', 208), ('AATSC7p', 208), ('AATSC7i', 208), ('MATS7dv', 208), ('MATS7d', 208), ('MATS7Z', 208), ('MATS7m', 208), ('MATS7v', 208), ('MATS7se', 208), ('MATS7pe', 208), ('MATS7are', 208), ('MATS7p', 208), ('MATS7i', 208), ('GATS7dv', 208), ('GATS7d', 208), ('GATS7Z', 208), ('GATS7m', 208), ('GATS7v', 208), ('GATS7se', 208), ('GATS7pe', 208), ('GATS7are', 208), ('GATS7p', 208), ('GATS7i', 208), ('MDEC-23', 155), ('AXp-6d', 141), ('AXp-6dv', 141), ('AXp-5d', 88), ('AXp-5dv', 88), ('AATSC6c', 86), ('MATS6c', 86), ('GATS6c', 86), ('AATS6dv', 84), ('AATS6d', 84), ('AATS6Z', 84), ('AATS6m', 84), ('AATS6v', 84), ('AATS6se', 84), ('AATS6pe', 84), ('AATS6are', 84), ('AATS6p', 84), ('AATS6i', 84), ('AATSC6dv', 84), ('AATSC6d', 84), ('AATSC6Z', 84), ('AATSC6m', 84), ('AATSC6v', 84), ('AATSC6se', 84), ('AATSC6pe', 84), ('AATSC6are', 84), ('AATSC6p', 84), ('AATSC6i', 84), ('MATS6dv', 84), ('MATS6d', 84), ('MATS6Z', 84), ('MATS6m', 84), ('MATS6v', 84), ('MATS6se', 84), ('MATS6pe', 84), ('MATS6are', 84), ('MATS6p', 84), ('MATS6i', 84), ('GATS6dv', 84), ('GATS6d', 84), ('GATS6Z', 84), ('GATS6m', 84), ('GATS6v', 84), ('GATS6se', 84), ('GATS6pe', 84), ('GATS6are', 84), ('GATS6p', 84), ('GATS6i', 84), ('AXp-4d', 54), ('AXp-4dv', 54), ('AATSC5c', 45), ('MATS5c', 45), ('GATS5c', 45), ('AATS5dv', 43), ('AATS5d', 43), ('AATS5Z', 43), ('AATS5m', 43), ('AATS5v', 43), ('AATS5se', 43), ('AATS5pe', 43), ('AATS5are', 43), ('AATS5p', 43), ('AATS5i', 43), ('AATSC5dv', 43), ('AATSC5d', 43), ('AATSC5Z', 43), ('AATSC5m', 43), ('AATSC5v', 43), ('AATSC5se', 43), ('AATSC5pe', 43), ('AATSC5are', 43), ('AATSC5p', 43), ('AATSC5i', 43), ('MATS5dv', 43), ('MATS5d', 43), ('MATS5Z', 43), ('MATS5m', 43), ('MATS5v', 43), ('MATS5se', 43), ('MATS5pe', 43), ('MATS5are', 43), ('MATS5p', 43), ('MATS5i', 43), ('GATS5dv', 43), ('GATS5d', 43), ('GATS5Z', 43), ('GATS5m', 43), ('GATS5v', 43), ('GATS5se', 43), ('GATS5pe', 43), ('GATS5are', 43), ('GATS5p', 43), ('GATS5i', 43), ('AXp-3d', 32), ('AXp-3dv', 32), ('Kier3', 32), ('AATSC4c', 27), ('MATS4c', 27), ('GATS4c', 27), ('AATS4dv', 25), ('AATS4d', 25), ('AATS4Z', 25), ('AATS4m', 25), ('AATS4v', 25), ('AATS4se', 25), ('AATS4pe', 25), ('AATS4are', 25), ('AATS4p',

25), ('AATS4i', 25), ('AATSC4dv', 25), ('AATSC4d', 25), ('AATSC4Z', 25), ('AATSC4m', 25), ('AATSC4v', 25), ('AATSC4se', 25), ('AATSC4pe', 25), ('AATSC4are', 25), ('AATSC4p', 25), ('AATSC4i', 25), ('MATS4dv', 25), ('MATS4d', 25), ('MATS4Z', 25), ('MATS4m', 25), ('MATS4v', 25), ('MATS4se', 25), ('MATS4pe', 25), ('MATS4are', 25), ('MATS4p', 25), ('MATS4i', 25), ('GATS4dv', 25), ('GATS4d', 25), ('GATS4Z', 25), ('GATS4m', 25), ('GATS4v', 25), ('GATS4se', 25), ('GATS4pe', 25), ('GATS4are', 25), ('GATS4p', 25), ('GATS4i', 25), ('AATSC3c', 23), ('MATS3c', 23), ('GATS3c', 23), ('HybRatio', 23), ('MATS3d', 22), ('GATS3d', 22), ('AATS3dv', 21), ('AATS3d', 21), ('AATS3Z', 21), ('AATS3m', 21), ('AATS3v', 21), ('AATS3se', 21), ('AATS3pe', 21), ('AATS3are', 21), ('AATS3p', 21), ('AATS3i', 21), ('AATSC3dv', 21), ('AATSC3d', 21), ('AATSC3Z', 21), ('AATSC3m', 21), ('AATSC3v', 21), ('AATSC3se', 21), ('AATSC3pe', 21), ('AATSC3are', 21), ('AATSC3p', 21), ('AATSC3i', 21), ('MATS3dv', 21), ('MATS3Z', 21), ('MATS3m', 21), ('MATS3v', 21), ('MATS3se', 21), ('MATS3pe', 21), ('MATS3are', 21), ('MATS3p', 21), ('MATS3i', 21), ('GATS3dv', 21), ('GATS3Z', 21), ('GATS3m', 21), ('GATS3v', 21), ('GATS3se', 21), ('GATS3pe', 21), ('GATS3are', 21), ('GATS3p', 21), ('GATS3i', 21), ('AXp-2d', 15), ('AXp-2dv', 15), ('Kier2', 15), ('AATSC2c', 14), ('MATS2c', 14), ('MATS2d', 14), ('GATS2c', 14), ('GATS2d', 14), ('AATSC1c', 13), ('MATS1c', 13), ('MATS1d', 13), ('GATS1c', 13), ('GATS1d', 13), ('AXp-1d', 13), ('AXp-1dv', 13), ('Kier1', 13), ('RotRatio', 13), ('VAdjMat', 13), ('AATS2dv', 12), ('AATS2d', 12), ('AATS2Z', 12), ('AATS2m', 12), ('AATS2v', 12), ('AATS2se', 12), ('AATS2pe', 12), ('AATS2are', 12), ('AATS2p', 12), ('AATS2i', 12), ('AATSC2dv', 12), ('AATSC2d', 12), ('AATSC2Z', 12), ('AATSC2m', 12), ('AATSC2v', 12), ('AATSC2se', 12), ('AATSC2pe', 12), ('AATSC2are', 12), ('AATSC2p', 12), ('AATSC2i', 12), ('MATS2dv', 12), ('MATS2Z', 12), ('MATS2m', 12), ('MATS2v', 12), ('MATS2se', 12), ('MATS2pe', 12), ('MATS2are', 12), ('MATS2p', 12), ('MATS2i', 12), ('GATS2dv', 12), ('GATS2Z', 12), ('GATS2m', 12), ('GATS2v', 12), ('GATS2se', 12), ('GATS2pe', 12), ('GATS2are', 12), ('GATS2p', 12), ('GATS2i', 12), ('AATS1dv', 11), ('AATS1d', 11), ('AATS1Z', 11), ('AATS1m', 11), ('AATS1v', 11), ('AATS1se', 11), ('AATS1pe', 11), ('AATS1are', 11), ('AATS1p', 11), ('AATS1i', 11), ('AATSC1dv', 11), ('AATSC1d', 11), ('AATSC1Z', 11), ('AATSC1m', 11), ('AATSC1v', 11), ('AATSC1se', 11), ('AATSC1pe', 11), ('AATSC1are', 11), ('AATSC1p', 11), ('AATSC1i', 11), ('MATS1dv', 11), ('MATS1Z', 11), ('MATS1m', 11), ('MATS1v', 11), ('MATS1se', 11), ('MATS1pe', 11), ('MATS1are', 11), ('MATS1p', 11), ('MATS1i', 11), ('GATS1dv', 11), ('GATS1Z', 11), ('GATS1m', 11), ('GATS1v', 11), ('GATS1se', 11), ('GATS1pe', 11), ('GATS1are', 11), ('GATS1p', 11), ('GATS1i', 11), ('BIC0', 11), ('BIC1', 11), ('BIC2', 11), ('BIC3', 11), ('BIC4', 11), ('BIC5', 11), ('ATSC0c', 2), ('ATSC1c', 2), ('ATSC2c', 2), ('ATSC3c', 2), ('ATSC4c', 2), ('ATSC5c', 2), ('ATSC6c', 2), ('ATSC7c', 2), ('ATSC8c', 2), ('AATSC0c', 2), ('RNCG', 2), ('RPCG', 2), ('nAcid', 0), ('nBase', 0), ('nAromAtom', 0), ('nAromBond', 0), ('nAtom', 0), ('nHeavyAtom', 0), ('nSpiro', 0), ('nBridgehead', 0), ('nHetero', 0), ('nH', 0), ('nB', 0), ('nC', 0), ('nN', 0), ('nO', 0), ('nS', 0), ('nP', 0), ('nF', 0), ('nCl', 0), ('nBr', 0), ('nI', 0), ('nX', 0), ('ATS0dv', 0), ('ATS1dv', 0), ('ATS2dv', 0), ('ATS3dv', 0), ('ATS4dv', 0), ('ATS5dv', 0), ('ATS6dv', 0), ('ATS7dv', 0), ('ATS8dv', 0), ('ATS0d', 0), ('ATS1d', 0), ('ATS2d', 0), ('ATS3d', 0), ('ATS4d', 0), ('ATS5d', 0), ('ATS6d', 0), ('ATS7d', 0), ('ATS8d', 0), ('ATS0Z', 0),

('ATS1Z', 0), ('ATS2Z', 0), ('ATS3Z', 0), ('ATS4Z', 0), ('ATS5Z', 0), ('ATS6Z', 0), ('ATS7Z', 0), ('ATS8Z', 0), ('ATS0m', 0), ('ATS1m', 0), ('ATS2m', 0), ('ATS3m', 0), ('ATS4m', 0), ('ATS5m', 0), ('ATS6m', 0), ('ATS7m', 0), ('ATS8m', 0), ('ATS0v', 0), ('ATS1v', 0), ('ATS2v', 0), ('ATS3v', 0), ('ATS4v', 0), ('ATS5v', 0), ('ATS6v', 0), ('ATS7v', 0), ('ATS8v', 0), ('ATS0se', 0), ('ATS1se', 0), ('ATS2se', 0), ('ATS3se', 0), ('ATS4se', 0), ('ATS5se', 0), ('ATS6se', 0), ('ATS7se', 0), ('ATS8se', 0), ('ATS0pe', 0), ('ATS1pe', 0), ('ATS2pe', 0), ('ATS3pe', 0), ('ATS4pe', 0), ('ATS5pe', 0), ('ATS6pe', 0), ('ATS7pe', 0), ('ATS8pe', 0), ('ATS0are', 0), ('ATS1are', 0), ('ATS2are', 0), ('ATS3are', 0), ('ATS4are', 0), ('ATS5are', 0), ('ATS6are', 0), ('ATS7are', 0), ('ATS8are', 0), ('ATS0p', 0), ('ATS1p', 0), ('ATS2p', 0), ('ATS3p', 0), ('ATS4p', 0), ('ATS5p', 0), ('ATS6p', 0), ('ATS7p', 0), ('ATS8p', 0), ('ATS0i', 0), ('ATS1i', 0), ('ATS2i', 0), ('ATS3i', 0), ('ATS4i', 0), ('ATS5i', 0), ('ATS6i', 0), ('ATS7i', 0), ('ATS8i', 0), ('AATS0dv', 0), ('AATS0d', 0), ('AATS0Z', 0), ('AATS0m', 0), ('AATS0v', 0), ('AATS0se', 0), ('AATS0pe', 0), ('AATS0are', 0), ('AATS0p', 0), ('AATS0i', 0), ('ATSC0dv', 0), ('ATSC1dv', 0), ('ATSC2dv', 0), ('ATSC3dv', 0), ('ATSC4dv', 0), ('ATSC5dv', 0), ('ATSC6dv', 0), ('ATSC7dv', 0), ('ATSC8dv', 0), ('ATSC0d', 0), ('ATSC1d', 0), ('ATSC2d', 0), ('ATSC3d', 0), ('ATSC4d', 0), ('ATSC5d', 0), ('ATSC6d', 0), ('ATSC7d', 0), ('ATSC8d', 0), ('ATSC0Z', 0), ('ATSC1Z', 0), ('ATSC2Z', 0), ('ATSC3Z', 0), ('ATSC4Z', 0), ('ATSC5Z', 0), ('ATSC6Z', 0), ('ATSC7Z', 0), ('ATSC8Z', 0), ('ATSC0m', 0), ('ATSC1m', 0), ('ATSC2m', 0), ('ATSC3m', 0), ('ATSC4m', 0), ('ATSC5m', 0), ('ATSC6m', 0), ('ATSC7m', 0), ('ATSC8m', 0), ('ATSC0v', 0), ('ATSC1v', 0), ('ATSC2v', 0), ('ATSC3v', 0), ('ATSC4v', 0), ('ATSC5v', 0), ('ATSC6v', 0), ('ATSC7v', 0), ('ATSC8v', 0), ('ATSC0se', 0), ('ATSC1se', 0), ('ATSC2se', 0), ('ATSC3se', 0), ('ATSC4se', 0), ('ATSC5se', 0), ('ATSC6se', 0), ('ATSC7se', 0), ('ATSC8se', 0), ('ATSC0pe', 0), ('ATSC1pe', 0), ('ATSC2pe', 0), ('ATSC3pe', 0), ('ATSC4pe', 0), ('ATSC5pe', 0), ('ATSC6pe', 0), ('ATSC7pe', 0), ('ATSC8pe', 0), ('ATSC0are', 0), ('ATSC1are', 0), ('ATSC2are', 0), ('ATSC3are', 0), ('ATSC4are', 0), ('ATSC5are', 0), ('ATSC6are', 0), ('ATSC7are', 0), ('ATSC8are', 0), ('ATSC0p', 0), ('ATSC1p', 0), ('ATSC2p', 0), ('ATSC3p', 0), ('ATSC4p', 0), ('ATSC5p', 0), ('ATSC6p', 0), ('ATSC7p', 0), ('ATSC8p', 0), ('ATSC0i', 0), ('ATSC1i', 0), ('ATSC2i', 0), ('ATSC3i', 0), ('ATSC4i', 0), ('ATSC5i', 0), ('ATSC6i', 0), ('ATSC7i', 0), ('ATSC8i', 0), ('AATSC0dv', 0), ('AATSC0d', 0), ('AATSC0Z', 0), ('AATSC0m', 0), ('AATSC0v', 0), ('AATSC0se', 0), ('AATSC0pe', 0), ('AATSC0are', 0), ('AATSC0p', 0), ('AATSC0i', 0), ('BalabanJ', 0), ('BertzCT', 0), ('nBonds', 0), ('nBondsO', 0), ('nBondsS', 0), ('nBondsD', 0), ('nBondsT', 0), ('nBondsA', 0), ('nBondsM', 0), ('nBondsKS', 0), ('nBondsKD', 0), ('C1SP1', 0), ('C2SP1', 0), ('C1SP2', 0), ('C2SP2', 0), ('C3SP2', 0), ('C1SP3', 0), ('C2SP3', 0), ('C3SP3', 0), ('C4SP3', 0), ('FCSP3', 0), ('Xch-3d', 0), ('Xch-4d', 0), ('Xch-5d', 0), ('Xch-6d', 0), ('Xch-7d', 0), ('Xch-3dv', 0), ('Xch-4dv', 0), ('Xch-5dv', 0), ('Xch-6dv', 0), ('Xch-7dv', 0), ('Xc-3d', 0), ('Xc-4d', 0), ('Xc-5d', 0), ('Xc-6d', 0), ('Xc-3dv', 0), ('Xc-4dv', 0), ('Xc-5dv', 0), ('Xc-6dv', 0), ('Xpc-4d', 0), ('Xpc-5d', 0), ('Xpc-6d', 0), ('Xpc-4dv', 0), ('Xpc-5dv', 0), ('Xpc-6dv', 0), ('Xp-1d', 0), ('Xp-2d', 0), ('Xp-3d', 0), ('Xp-4d', 0), ('Xp-5d', 0), ('Xp-6d', 0), ('Xp-7d', 0), ('Xp-1dv', 0), ('Xp-2dv', 0), ('Xp-3dv', 0), ('Xp-4dv', 0), ('Xp-5dv', 0), ('Xp-6dv', 0), ('Xp-7dv', 0), ('SZ', 0), ('Sm', 0), ('Sv', 0), ('Sse', 0),

('Spe', 0), ('Sare', 0), ('Sp', 0), ('Si', 0), ('MZ', 0), ('Mm', 0), ('Mv', 0), ('Mse', 0), ('Mpe', 0), ('Mare', 0), ('Mp', 0), ('Mi', 0), ('ECIndex', 0), ('fragCpx', 0), ('fMF', 0), ('nHBAcc', 0), ('nHBDon', 0), ('IC0', 0), ('IC1', 0), ('IC2', 0), ('IC3', 0), ('IC4', 0), ('IC5', 0), ('TIC0', 0), ('TIC1', 0), ('TIC2', 0), ('TIC3', 0), ('TIC4', 0), ('TIC5', 0), ('SIC0', 0), ('SIC1', 0), ('SIC2', 0), ('SIC3', 0), ('SIC4', 0), ('SIC5', 0), ('CIC0', 0), ('CIC1', 0), ('CIC2', 0), ('CIC3', 0), ('CIC4', 0), ('CIC5', 0), ('MIC0', 0), ('MIC1', 0), ('MIC2', 0), ('MIC3', 0), ('MIC4', 0), ('MIC5', 0), ('ZMIC0', 0), ('ZMIC1', 0), ('ZMIC2', 0), ('ZMIC3', 0), ('ZMIC4', 0), ('ZMIC5', 0), ('Lipinski', 0), ('GhoseFilter', 0), ('FilterItLogS', 0), ('VMcGowan', 0), ('LabuteASA', 0), ('PEOE_VSA1', 0), ('PEOE_VSA2', 0), ('PEOE_VSA3', 0), ('PEOE_VSA4', 0), ('PEOE_VSA5', 0), ('PEOE_VSA6', 0), ('PEOE_VSA7', 0), ('PEOE_VSA8', 0), ('PEOE_VSA9', 0), ('PEOE_VSA10', 0), ('PEOE_VSA11', 0), ('PEOE_VSA12', 0), ('PEOE_VSA13', 0), ('SMR_VSA1', 0), ('SMR_VSA2', 0), ('SMR_VSA3', 0), ('SMR_VSA4', 0), ('SMR_VSA5', 0), ('SMR_VSA6', 0), ('SMR_VSA7', 0), ('SMR_VSA8', 0), ('SMR_VSA9', 0), ('SlogP_VSA1', 0), ('SlogP_VSA2', 0), ('SlogP_VSA3', 0), ('SlogP_VSA4', 0), ('SlogP_VSA5', 0), ('SlogP_VSA6', 0), ('SlogP_VSA7', 0), ('SlogP_VSA8', 0), ('SlogP_VSA9', 0), ('SlogP_VSA10', 0), ('SlogP_VSA11', 0), ('MPC2', 0), ('MPC3', 0), ('MPC4', 0), ('MPC5', 0), ('MPC6', 0), ('MPC7', 0), ('MPC8', 0), ('MPC9', 0), ('MPC10', 0), ('TMPC10', 0), ('piPC1', 0), ('piPC2', 0), ('piPC3', 0), ('piPC4', 0), ('piPC5', 0), ('piPC6', 0), ('piPC7', 0), ('piPC8', 0), ('piPC9', 0), ('piPC10', 0), ('TpiPC10', 0), ('apol', 0), ('bpol', 0), ('nRing', 0), ('n3Ring', 0), ('n4Ring', 0), ('n5Ring', 0), ('n6Ring', 0), ('n7Ring', 0), ('n8Ring', 0), ('n9Ring', 0), ('n10Ring', 0), ('n11Ring', 0), ('n12Ring', 0), ('nG12Ring', 0), ('nHRing', 0), ('n3HRing', 0), ('n4HRing', 0), ('n5HRing', 0), ('n6HRing', 0), ('n7HRing', 0), ('n8HRing', 0), ('n9HRing', 0), ('n10HRing', 0), ('n11HRing', 0), ('n12HRing', 0), ('nG12HRing', 0), ('naRing', 0), ('n3aRing', 0), ('n4aRing', 0), ('n5aRing', 0), ('n6aRing', 0), ('n7aRing', 0), ('n8aRing', 0), ('n9aRing', 0), ('n10aRing', 0), ('n11aRing', 0), ('n12aRing', 0), ('nG12aRing', 0), ('naHRing', 0), ('n3aHRing', 0), ('n4aHRing', 0), ('n5aHRing', 0), ('n6aHRing', 0), ('n7aHRing', 0), ('n8aHRing', 0), ('n9aHRing', 0), ('n10aHRing', 0), ('n11aHRing', 0), ('n12aHRing', 0), ('nG12aHRing', 0), ('nARing', 0), ('n3ARing', 0), ('n4ARing', 0), ('n5ARing', 0), ('n6ARing', 0), ('n7ARing', 0), ('n8ARing', 0), ('n9ARing', 0), ('n10ARing', 0), ('n11ARing', 0), ('n12ARing', 0), ('nG12ARing', 0), ('nAHRing', 0), ('n3AHRing', 0), ('n4AHRing', 0), ('n5AHRing', 0), ('n6AHRing', 0), ('n7AHRing', 0), ('n8AHRing', 0), ('n9AHRing', 0), ('n10AHRing', 0), ('n11AHRing', 0), ('n12AHRing', 0), ('nG12AHRing', 0), ('nFRing', 0), ('n4FRing', 0), ('n5FRing', 0), ('n6FRing', 0), ('n7FRing', 0), ('n8FRing', 0), ('n9FRing', 0), ('n10FRing', 0), ('n11FRing', 0), ('n12FRing', 0), ('nG12FRing', 0), ('nFHRing', 0), ('n4FHRing', 0), ('n5FHRing', 0), ('n6FHRing', 0), ('n7FHRing', 0), ('n8FHRing', 0), ('n9FHRing', 0), ('n10FHRing', 0), ('n11FHRing', 0), ('n12FHRing', 0), ('nG12FHRing', 0), ('nFaRing', 0), ('n4FaRing', 0), ('n5FaRing', 0), ('n6FaRing', 0), ('n7FaRing', 0), ('n8FaRing', 0), ('n9FaRing', 0), ('n10FaRing', 0), ('n11FaRing', 0), ('n12FaRing', 0), ('nG12FaRing', 0), ('nFaHRing', 0), ('n4FaHRing', 0), ('n5FaHRing', 0), ('n6FaHRing', 0), ('n7FaHRing', 0), ('n8FaHRing', 0), ('n9FaHRing', 0), ('n10FaHRing', 0), ('n11FaHRing', 0), ('n12FaHRing', 0), ('nG12FaHRing', 0), ('nFARing', 0),

('n4FARing', 0), ('n5FARing', 0), ('n6FARing', 0), ('n7FARing', 0), ('n8FARing', 0), ('n9FARing', 0), ('n10FARing', 0), ('n11FARing', 0), ('n12FARing', 0), ('nG12FARing', 0), ('nFAHRing', 0), ('n4FAHRing', 0), ('n5FAHRing', 0), ('n6FAHRing', 0), ('n7FAHRing', 0), ('n8FAHRing', 0), ('n9FAHRing', 0), ('n10FAHRing', 0), ('n11FAHRing', 0), ('n12FAHRing', 0), ('nG12FAHRing', 0), ('nRot', 0), ('SLogP', 0), ('SMR', 0), ('TopoPSA(NO)', 0), ('TopoPSA', 0), ('GGI1', 0), ('GGI2', 0), ('GGI3', 0), ('GGI4', 0), ('GGI5', 0), ('GGI6', 0), ('GGI7', 0), ('GGI8', 0), ('GGI9', 0), ('GGI10', 0), ('JGI1', 0), ('JGI2', 0), ('JGI3', 0), ('JGI4', 0), ('JGI5', 0), ('JGI6', 0), ('JGI7', 0), ('JGI8', 0), ('JGI9', 0), ('JGI10', 0), ('JGT10', 0), ('Diameter', 0), ('Radius', 0), ('TopoShapeIndex', 0), ('PetitjeanIndex', 0), ('MWC01', 0), ('MWC02', 0), ('MWC03', 0), ('MWC04', 0), ('MWC05', 0), ('MWC06', 0), ('MWC07', 0), ('MWC08', 0), ('MWC09', 0), ('MWC10', 0), ('TMWC10', 0), ('SRW02', 0), ('SRW03', 0), ('SRW04', 0), ('SRW05', 0), ('SRW06', 0), ('SRW07', 0), ('SRW08', 0), ('SRW09', 0), ('SRW10', 0), ('TSRW10', 0), ('MW', 0), ('AMW', 0), ('WPath', 0), ('WPol', 0), ('Zagreb1', 0), ('Zagreb2', 0), ('mZagreb2', 0)]
[(553, 656), (580, 656), (1014, 656), (1092, 656), (1168, 656), (1776, 656), (1818, 656), (2085, 656), (2137, 656), (2400, 656), (2835, 656), (961, 607), (1277, 563), (784, 556), (1219, 555), (1803, 555), (2239, 555), (2657, 555), (43, 512), (1833, 465), (678, 464), (605, 462), (608, 419), (1117, 419), (1110, 418), (1509, 418), (2638, 418), (2244, 414), (281, 413), (2176, 411), (1860, 374), (329, 373), (818, 373), (1066, 373), (2832, 373), (1775, 372), (2567, 372), (2653, 372), (872, 371), (1520, 371), (2023, 371), (2132, 371), (2295, 371), (2530, 371), (2538, 371), (2819, 371), (2907, 371), (1977, 370), (347, 369), (461, 369), (742, 369), (1422, 369), (2310, 369), (810, 368), (831, 368), (1211, 368), (1759, 368), (1962, 368), (2420, 368), (2532, 368), (385, 367), (480, 367), (1087, 367), (1836, 367), (991, 366), (1524, 366), (2682, 366), (918, 365), (849, 329), (1959, 328), (2406, 328), (2516, 328), (336, 327), (1141, 327), (1439, 327), (1609, 327), (2427, 327), (98, 326), (253, 326), (649, 326), (836, 326), (1076, 326), (1281, 326), (1360, 326), (1465, 326), (1971, 326), (2064, 326), (2115, 326), (2350, 326), (2367, 326), (2576, 326), (27, 325), (104, 325), (157, 325), (292, 325), (410, 325), (528, 325), (587, 325), (630, 325), (638, 325), (651, 325), (740, 325), (756, 325), (770, 325), (792, 325), (1013, 325), (1037, 325), (1172, 325), (1190, 325), (1224, 325), (1252, 325), (1297, 325), (1328, 325), (1382, 325), (1403, 325), (1428, 325), (1526, 325), (1561, 325), (1578, 325), (1581, 325), (1635, 325), (1659, 325), (1719, 325), (1855, 325), (1874, 325), (1890, 325), (1951, 325), (1974, 325), (1989, 325), (2046, 325), (2073, 325), (2096, 325), (2105, 325), (2428, 325), (2456, 325), (2471, 325), (2473, 325), (2488, 325), (2528, 325), (2564, 325), (2565, 325), (2601, 325), (2665, 325), (2788, 325), (2804, 325), (2911, 325), (224, 324), (233, 324), (320, 324), (370, 324), (389, 324), (493, 324), (660, 324), (1056, 324), (1131, 324), (1268, 324), (1290, 324), (1322, 324), (1339, 324), (1487, 324), (1653, 324), (1853, 324), (1863, 324), (1954, 324), (1960, 324), (2030, 324), (2108, 324), (2296, 324), (2375, 324), (2518, 324), (2931, 324), (2932, 324), (4, 323), (67, 323), (87, 323), (291, 323), (488, 323), (542, 323), (575, 323), (871, 323), (1151, 323), (1495, 323), (1565, 323), (1604, 323), (1611, 323), (1624, 323), (1840, 323), (1847, 323), (1927, 323), (1928, 323), (2000, 323), (2291, 323), (2322, 323), (2362, 323), (2437, 323), (2617, 323),

(2627, 323), (2756, 323), (2812, 323), (2843, 323), (2890, 323), (2943, 323), (332, 322), (450, 322), (549, 322), (670, 322), (724, 322), (752, 322), (995, 322), (1123, 322), (1161, 322), (1223, 322), (1453, 322), (1573, 322), (1617, 322), (1806, 322), (2269, 322), (2472, 322), (2555, 322), (2602, 322), (2644, 322), (2749, 322), (2784, 322), (2814, 322), (1011, 321), (1620, 321), (2726, 321), (665, 320), (1471, 320), (957, 318), (1093, 314), (2316, 312), (2802, 312), (2729, 284), (242, 283), (571, 283), (736, 283), (936, 283), (1100, 283), (1189, 283), (1247, 283), (1315, 283), (1461, 283), (1493, 283), (2195, 283), (2438, 283), (2480, 283), (187, 282), (197, 282), (294, 282), (489, 282), (548, 282), (693, 282), (998, 282), (1097, 282), (1104, 282), (1256, 282), (1353, 282), (1656, 282), (1707, 282), (1912, 282), (1917, 282), (184, 281), (243, 281), (305, 281), (495, 281), (595, 281), (893, 281), (910, 281), (944, 281), (969, 281), (988, 281), (1077, 281), (1094, 281), (1160, 281), (1208, 281), (1264, 281), (1321, 281), (1359, 281), (1549, 281), (1588, 281), (1642, 281), (1721, 281), (1858, 281), (1955, 281), (1968, 281), (2032, 281), (2049, 281), (2135, 281), (2138, 281), (2205, 281), (2215, 281), (2248, 281), (2417, 281), (2493, 281), (2539, 281), (2631, 281), (2739, 281), (2754, 281), (2833, 281), (20, 280), (41, 280), (103, 280), (110, 280), (117, 280), (118, 280), (151, 280), (196, 280), (200, 280), (205, 280), (501, 280), (577, 280), (591, 280), (669, 280), (681, 280), (707, 280), (746, 280), (760, 280), (773, 280), (800, 280), (830, 280), (837, 280), (858, 280), (869, 280), (965, 280), (970, 280), (1034, 280), (1098, 280), (1143, 280), (1299, 280), (1498, 280), (1544, 280), (1568, 280), (1622, 280), (1724, 280), (1767, 280), (1915, 280), (2155, 280), (2422, 280), (2687, 280), (2740, 280), (2803, 280), (2872, 280), (14, 279), (93, 279), (99, 279), (125, 279), (128, 279), (131, 279), (244, 279), (245, 279), (303, 279), (390, 279), (515, 279), (530, 279), (532, 279), (535, 279), (602, 279), (618, 279), (667, 279), (726, 279), (820, 279), (863, 279), (882, 279), (924, 279), (958, 279), (1024, 279), (1124, 279), (1182, 279), (1199, 279), (1232, 279), (1265, 279), (1267, 279), (1282, 279), (1286, 279), (1325, 279), (1352, 279), (1414, 279), (1424, 279), (1445, 279), (1489, 279), (1506, 279), (1589, 279), (1597, 279), (1751, 279), (1844, 279), (1867, 279), (1953, 279), (2014, 279), (2031, 279), (2050, 279), (2102, 279), (2156, 279), (2183, 279), (2256, 279), (2283, 279), (2365, 279), (2394, 279), (2407, 279), (2546, 279), (2579, 279), (2595, 279), (2603, 279), (2605, 279), (2624, 279), (2628, 279), (2786, 279), (2810, 279), (2858, 279), (2868, 279), (23, 278), (33, 278), (90, 278), (210, 278), (213, 278), (328, 278), (330, 278), (397, 278), (619, 278), (637, 278), (642, 278), (691, 278), (737, 278), (759, 278), (774, 278), (833, 278), (939, 278), (1027, 278), (1049, 278), (1062, 278), (1065, 278), (1179, 278), (1269, 278), (1375, 278), (1511, 278), (1582, 278), (1661, 278), (1671, 278), (1791, 278), (1903, 278), (2070, 278), (2088, 278), (2169, 278), (2171, 278), (2213, 278), (2223, 278), (2230, 278), (2251, 278), (2288, 278), (2304, 278), (2379, 278), (2478, 278), (2616, 278), (2621, 278), (2654, 278), (2660, 278), (2663, 278), (2676, 278), (2710, 278), (2720, 278), (2796, 278), (2856, 278), (2860, 278), (209, 277), (221, 277), (323, 277), (351, 277), (478, 277), (483, 277), (739, 277), (757, 277), (1181, 277), (1332, 277), (1409, 277), (1497, 277), (1513, 277), (1551, 277), (1591, 277), (1678, 277), (1919, 277), (1985, 277), (2079, 277), (2203, 277), (2312, 277), (2482, 277), (2684, 277), (2885, 277), (2927, 277), (60, 276), (62, 276), (102, 276), (140, 276), (167,

276), (235, 276), (252, 276), (475, 276), (560, 276), (676, 276), (712, 276), (794, 276), (905, 276), (1028, 276), (1296, 276), (1602, 276), (1613, 276), (1765, 276), (2002, 276), (2018, 276), (2075, 276), (2158, 276), (2164, 276), (2166, 276), (2263, 276), (2276, 276), (2300, 276), (2560, 276), (2696, 276), (2830, 276), (2924, 276), (3, 275), (340, 275), (441, 275), (565, 275), (576, 275), (579, 275), (735, 275), (767, 275), (883, 275), (1025, 275), (1185, 275), (1222, 275), (1340, 275), (1342, 275), (1418, 275), (1419, 275), (1449, 275), (1615, 275), (1645, 275), (1774, 275), (1838, 275), (1920, 275), (1937, 275), (1979, 275), (2025, 275), (2083, 275), (2152, 275), (2185, 275), (2202, 275), (2221, 275), (2401, 275), (2435, 275), (2515, 275), (2799, 275), (2904, 275), (414, 274), (640, 274), (779, 274), (832, 274), (1040, 274), (1063, 274), (1782, 274), (2082, 274), (2089, 274), (2479, 274), (2790, 274), (144, 273), (363, 273), (481, 273), (558, 273), (672, 273), (881, 273), (940, 273), (1248, 273), (1650, 273), (1672, 273), (1787, 273), (1893, 273), (2044, 273), (2571, 273), (2669, 273), (255, 272), (283, 272), (337, 272), (386, 272), (945, 272), (1335, 272), (1395, 272), (1736, 272), (100, 270), (1351, 270), (2534, 270), (2142, 264), (1502, 262), (1980, 261), (780, 224), (1554, 223), (1188, 222), (1811, 222), (574, 221), (922, 221), (1366, 221), (1772, 221), (2338, 221), (2432, 221), (2436, 221), (791, 220), (1005, 220), (1388, 220), (514, 219), (616, 219), (653, 219), (714, 219), (2074, 219), (2469, 219), (2655, 219), (105, 218), (1408, 218), (2133, 218), (2453, 218), (116, 217), (257, 217), (429, 217), (993, 217), (1068, 217), (1393, 217), (2173, 217), (2731, 217), (2855, 217), (29, 216), (492, 216), (1242, 216), (1389, 216), (1616, 216), (1810, 216), (2694, 216), (2080, 215), (2828, 215), (156, 213), (2106, 213), (172, 212), (626, 212), (1109, 211), (1479, 211), (1355, 208), (1101, 167), (1012, 166), (2047, 166), (2157, 166), (2594, 166), (21, 165), (721, 165), (1894, 165), (2562, 165), (2672, 165), (298, 164), (459, 164), (627, 164), (1510, 164), (189, 163), (1139, 163), (1249, 163), (2116, 163), (175, 162), (529, 162), (1601, 162), (2661, 162), (1529, 159), (11, 158), (383, 158), (1773, 158), (2003, 158), (2182, 158), (875, 157), (1074, 157), (1828, 156), (2364, 155), (808, 150), (2275, 117), (201, 116), (290, 116), (510, 116), (1319, 116), (2184, 116), (2271, 116), (2273, 116), (2556, 116), (2575, 116), (2758, 116), (566, 115), (1346, 115), (2439, 115), (2674, 115), (2824, 115), (263, 114), (362, 114), (589, 114), (1823, 114), (1978, 114), (2487, 114), (275, 113), (1369, 113), (1958, 113), (2274, 113), (1118, 112), (1391, 112), (134, 111), (186, 111), (240, 111), (365, 111), (1128, 111), (1546, 111), (1804, 111), (1817, 111), (1994, 111), (2589, 111), (2771, 111), (44, 110), (960, 110), (1567, 110), (1633, 110), (2718, 110), (2732, 110), (82, 109), (288, 109), (300, 109), (307, 109), (350, 109), (829, 109), (900, 109), (978, 109), (1221, 109), (1255, 109), (1397, 109), (1491, 109), (1627, 109), (1679, 109), (2302, 109), (2542, 109), (2580, 109), (2826, 109), (2922, 109), (176, 108), (354, 108), (360, 108), (436, 108), (517, 108), (692, 108), (709, 108), (1029, 108), (1142, 108), (1621, 108), (1666, 108), (1737, 108), (1832, 108), (2017, 108), (2392, 108), (1761, 107), (2048, 107), (146, 106), (764, 106), (1610, 106), (1535, 103), (2470, 103), (2730, 68), (597, 67), (994, 67), (2713, 67), (2792, 67), (203, 66), (953, 66), (1475, 66), (1603, 66), (2742, 66), (1559, 65), (2149, 65), (2483, 65), (2614, 65), (2, 64), (195, 64), (511, 64), (598, 64), (674, 64), (682, 64), (855, 64), (867, 64), (1197, 64), (1317, 64), (1323, 64), (1337, 64), (1371, 64), (1446, 64), (1935, 64),

(2321, 64), (2497, 64), (2550, 64), (2574, 64), (2583, 64), (2930, 64), (143, 63), (1088, 63), (1090, 63), (1279, 63), (1837, 63), (1895, 63), (2122, 63), (2585, 63), (2914, 63), (45, 62), (250, 62), (282, 62), (804, 62), (888, 62), (890, 62), (952, 62), (977, 62), (1081, 62), (1149, 62), (1234, 62), (1263, 62), (1392, 62), (1632, 62), (1777, 62), (1848, 62), (1921, 62), (2331, 62), (2358, 62), (2666, 62), (2880, 62), (2910, 62), (376, 61), (393, 61), (457, 61), (1120, 61), (1379, 61), (1608, 61), (1619, 61), (1735, 61), (1801, 61), (1862, 61), (2162, 61), (2398, 61), (2629, 61), (2701, 61), (2761, 61), (509, 60), (1399, 60), (1731, 60), (2314, 60), (2347, 60), (2360, 60), (2395, 60), (485, 59), (711, 59), (1220, 59), (1230, 59), (1372, 59), (1788, 59), (1808, 59), (2468, 59), (2481, 59), (2517, 59), (2639, 59), (2716, 59), (2841, 59), (886, 58), (948, 58), (671, 57), (1114, 57), (2600, 57), (126, 56), (909, 56), (2622, 55), (2942, 51), (2862, 19), (138, 17), (226, 17), (606, 17), (906, 17), (1059, 17), (1262, 17), (1398, 17), (1455, 17), (1512, 17), (1717, 17), (1740, 17), (1827, 17), (1850, 17), (1961, 17), (2097, 17), (2558, 17), (2757, 17), (2817, 17), (2923, 17), (80, 16), (232, 16), (312, 16), (673, 16), (696, 16), (738, 16), (796, 16), (816, 16), (1164, 16), (1191, 16), (1442, 16), (1652, 16), (1854, 16), (2193, 16), (2292, 16), (2376, 16), (2413, 16), (2640, 16), (2851, 16), (79, 15), (139, 15), (148, 15), (153, 15), (193, 15), (218, 15), (219, 15), (287, 15), (319, 15), (367, 15), (382, 15), (424, 15), (519, 15), (572, 15), (604, 15), (643, 15), (644, 15), (703, 15), (717, 15), (730, 15), (733, 15), (753, 15), (769, 15), (771, 15), (877, 15), (976, 15), (999, 15), (1078, 15), (1111, 15), (1176, 15), (1194, 15), (1201, 15), (1209, 15), (1213, 15), (1231, 15), (1306, 15), (1404, 15), (1411, 15), (1458, 15), (1518, 15), (1539, 15), (1540, 15), (1555, 15), (1556, 15), (1583, 15), (1596, 15), (1663, 15), (1720, 15), (1723, 15), (1755, 15), (1781, 15), (1834, 15), (1846, 15), (1924, 15), (1940, 15), (1970, 15), (1995, 15), (2012, 15), (2112, 15), (2113, 15), (2118, 15), (2129, 15), (2150, 15), (2194, 15), (2201, 15), (2218, 15), (2229, 15), (2245, 15), (2261, 15), (2333, 15), (2410, 15), (2464, 15), (2499, 15), (2501, 15), (2512, 15), (2536, 15), (2587, 15), (2632, 15), (2641, 15), (2664, 15), (2670, 15), (2748, 15), (2751, 15), (2752, 15), (2755, 15), (2767, 15), (2770, 15), (2853, 15), (2876, 15), (2895, 15), (15, 14), (47, 14), (63, 14), (86, 14), (124, 14), (129, 14), (133, 14), (160, 14), (166, 14), (185, 14), (188, 14), (217, 14), (228, 14), (230, 14), (238, 14), (241, 14), (271, 14), (276, 14), (293, 14), (302, 14), (326, 14), (345, 14), (366, 14), (398, 14), (401, 14), (411, 14), (443, 14), (448, 14), (449, 14), (467, 14), (473, 14), (474, 14), (479, 14), (504, 14), (578, 14), (586, 14), (666, 14), (702, 14), (732, 14), (747, 14), (751, 14), (754, 14), (777, 14), (789, 14), (827, 14), (840, 14), (842, 14), (850, 14), (851, 14), (854, 14), (861, 14), (862, 14), (866, 14), (892, 14), (894, 14), (946, 14), (962, 14), (979, 14), (985, 14), (1018, 14), (1019, 14), (1041, 14), (1045, 14), (1057, 14), (1061, 14), (1091, 14), (1096, 14), (1112, 14), (1115, 14), (1135, 14), (1166, 14), (1167, 14), (1178, 14), (1202, 14), (1233, 14), (1244, 14), (1253, 14), (1305, 14), (1349, 14), (1368, 14), (1436, 14), (1448, 14), (1459, 14), (1478, 14), (1492, 14), (1515, 14), (1516, 14), (1522, 14), (1530, 14), (1542, 14), (1543, 14), (1550, 14), (1618, 14), (1631, 14), (1643, 14), (1689, 14), (1705, 14), (1713, 14), (1728, 14), (1729, 14), (1763, 14), (1800, 14), (1807, 14), (1849, 14), (1882, 14), (1891, 14), (1930, 14), (1938, 14), (1948, 14), (1950, 14), (1952, 14), (1957, 14),

(1967, 14), (2016, 14), (2059, 14), (2069, 14), (2110, 14), (2128, 14), (2151, 14), (2170, 14), (2174, 14), (2217, 14), (2220, 14), (2228, 14), (2249, 14), (2252, 14), (2264, 14), (2268, 14), (2320, 14), (2330, 14), (2383, 14), (2389, 14), (2391, 14), (2408, 14), (2421, 14), (2460, 14), (2477, 14), (2491, 14), (2533, 14), (2552, 14), (2581, 14), (2590, 14), (2598, 14), (2610, 14), (2612, 14), (2637, 14), (2648, 14), (2651, 14), (2680, 14), (2733, 14), (2765, 14), (2798, 14), (2847, 14), (2854, 14), (2867, 14), (2894, 14), (2913, 14), (2916, 14), (2919, 14), (2921, 14), (2928, 14), (2941, 14), (7, 13), (16, 13), (19, 13), (24, 13), (26, 13), (31, 13), (37, 13), (42, 13), (48, 13), (53, 13), (54, 13), (70, 13), (71, 13), (73, 13), (81, 13), (89, 13), (92, 13), (95, 13), (101, 13), (120, 13), (121, 13), (123, 13), (135, 13), (137, 13), (150, 13), (158, 13), (161, 13), (162, 13), (163, 13), (164, 13), (168, 13), (180, 13), (183, 13), (192, 13), (202, 13), (211, 13), (214, 13), (225, 13), (248, 13), (249, 13), (262, 13), (274, 13), (278, 13), (296, 13), (304, 13), (306, 13), (314, 13), (316, 13), (322, 13), (325, 13), (338, 13), (348, 13), (352, 13), (355, 13), (368, 13), (378, 13), (379, 13), (381, 13), (394, 13), (399, 13), (405, 13), (407, 13), (418, 13), (426, 13), (427, 13), (462, 13), (468, 13), (477, 13), (486, 13), (496, 13), (497, 13), (498, 13), (512, 13), (518, 13), (527, 13), (531, 13), (533, 13), (540, 13), (564, 13), (582, 13), (588, 13), (593, 13), (594, 13), (596, 13), (613, 13), (614, 13), (615, 13), (617, 13), (623, 13), (629, 13), (634, 13), (636, 13), (647, 13), (655, 13), (656, 13), (658, 13), (659, 13), (661, 13), (675, 13), (683, 13), (687, 13), (715, 13), (728, 13), (741, 13), (749, 13), (750, 13), (772, 13), (797, 13), (812, 13), (817, 13), (823, 13), (843, 13), (844, 13), (857, 13), (870, 13), (874, 13), (878, 13), (880, 13), (885, 13), (891, 13), (913, 13), (927, 13), (935, 13), (954, 13), (963, 13), (974, 13), (983, 13), (984, 13), (997, 13), (1001, 13), (1003, 13), (1030, 13), (1036, 13), (1042, 13), (1048, 13), (1069, 13), (1085, 13), (1103, 13), (1108, 13), (1122, 13), (1145, 13), (1158, 13), (1163, 13), (1173, 13), (1175, 13), (1250, 13), (1272, 13), (1275, 13), (1285, 13), (1288, 13), (1289, 13), (1291, 13), (1303, 13), (1350, 13), (1373, 13), (1376, 13), (1380, 13), (1383, 13), (1385, 13), (1390, 13), (1400, 13), (1415, 13), (1426, 13), (1429, 13), (1438, 13), (1447, 13), (1451, 13), (1454, 13), (1484, 13), (1500, 13), (1501, 13), (1519, 13), (1521, 13), (1525, 13), (1528, 13), (1545, 13), (1562, 13), (1566, 13), (1575, 13), (1576, 13), (1595, 13), (1606, 13), (1626, 13), (1648, 13), (1649, 13), (1658, 13), (1662, 13), (1674, 13), (1685, 13), (1686, 13), (1690, 13), (1691, 13), (1696, 13), (1697, 13), (1702, 13), (1726, 13), (1746, 13), (1757, 13), (1766, 13), (1768, 13), (1780, 13), (1805, 13), (1813, 13), (1820, 13), (1829, 13), (1839, 13), (1851, 13), (1868, 13), (1885, 13), (1886, 13), (1887, 13), (1892, 13), (1910, 13), (1925, 13), (1945, 13), (1946, 13), (1966, 13), (1984, 13), (1999, 13), (2001, 13), (2010, 13), (2033, 13), (2034, 13), (2042, 13), (2054, 13), (2058, 13), (2078, 13), (2092, 13), (2104, 13), (2119, 13), (2125, 13), (2139, 13), (2140, 13), (2143, 13), (2147, 13), (2148, 13), (2163, 13), (2175, 13), (2191, 13), (2199, 13), (2204, 13), (2207, 13), (2211, 13), (2214, 13), (2225, 13), (2227, 13), (2236, 13), (2240, 13), (2247, 13), (2260, 13), (2272, 13), (2279, 13), (2281, 13), (2282, 13), (2306, 13), (2308, 13), (2309, 13), (2315, 13), (2327, 13), (2341, 13), (2349, 13), (2356, 13), (2359, 13), (2368, 13), (2371, 13), (2372, 13), (2373, 13), (2377, 13), (2385, 13), (2404, 13), (2409, 13), (2414, 13), (2433, 13), (2440,

13), (2441, 13), (2449, 13), (2450, 13), (2458, 13), (2467, 13), (2476, 13), (2496, 13), (2502, 13), (2507, 13), (2519, 13), (2520, 13), (2527, 13), (2537, 13), (2554, 13), (2559, 13), (2568, 13), (2570, 13), (2604, 13), (2619, 13), (2643, 13), (2658, 13), (2668, 13), (2698, 13), (2702, 13), (2706, 13), (2717, 13), (2723, 13), (2745, 13), (2753, 13), (2762, 13), (2763, 13), (2772, 13), (2781, 13), (2783, 13), (2787, 13), (2800, 13), (2816, 13), (2818, 13), (2820, 13), (2839, 13), (2840, 13), (2842, 13), (2861, 13), (2864, 13), (2870, 13), (2902, 13), (2903, 13), (2906, 13), (2917, 13), (2918, 13), (2936, 13), (6, 12), (18, 12), (22, 12), (36, 12), (46, 12), (52, 12), (55, 12), (58, 12), (61, 12), (75, 12), (77, 12), (84, 12), (107, 12), (109, 12), (112, 12), (113, 12), (170, 12), (173, 12), (182, 12), (215, 12), (216, 12), (231, 12), (237, 12), (247, 12), (251, 12), (264, 12), (266, 12), (285, 12), (313, 12), (334, 12), (335, 12), (377, 12), (387, 12), (388, 12), (402, 12), (406, 12), (419, 12), (434, 12), (435, 12), (438, 12), (445, 12), (447, 12), (451, 12), (452, 12), (484, 12), (507, 12), (508, 12), (521, 12), (536, 12), (537, 12), (547, 12), (555, 12), (563, 12), (567, 12), (569, 12), (570, 12), (584, 12), (601, 12), (612, 12), (641, 12), (645, 12), (664, 12), (680, 12), (686, 12), (695, 12), (698, 12), (729, 12), (744, 12), (758, 12), (768, 12), (778, 12), (787, 12), (801, 12), (803, 12), (814, 12), (821, 12), (822, 12), (835, 12), (864, 12), (865, 12), (884, 12), (887, 12), (899, 12), (902, 12), (915, 12), (928, 12), (929, 12), (934, 12), (949, 12), (959, 12), (972, 12), (986, 12), (987, 12), (1006, 12), (1017, 12), (1032, 12), (1035, 12), (1058, 12), (1060, 12), (1071, 12), (1073, 12), (1086, 12), (1105, 12), (1121, 12), (1126, 12), (1146, 12), (1148, 12), (1153, 12), (1162, 12), (1171, 12), (1180, 12), (1183, 12), (1184, 12), (1193, 12), (1195, 12), (1198, 12), (1204, 12), (1210, 12), (1215, 12), (1239, 12), (1246, 12), (1254, 12), (1278, 12), (1284, 12), (1300, 12), (1320, 12), (1333, 12), (1347, 12), (1354, 12), (1357, 12), (1358, 12), (1362, 12), (1363, 12), (1365, 12), (1377, 12), (1381, 12), (1386, 12), (1410, 12), (1427, 12), (1437, 12), (1444, 12), (1460, 12), (1483, 12), (1485, 12), (1490, 12), (1514, 12), (1560, 12), (1563, 12), (1564, 12), (1569, 12), (1570, 12), (1574, 12), (1580, 12), (1584, 12), (1586, 12), (1590, 12), (1598, 12), (1607, 12), (1612, 12), (1630, 12), (1667, 12), (1670, 12), (1709, 12), (1725, 12), (1741, 12), (1762, 12), (1764, 12), (1797, 12), (1798, 12), (1815, 12), (1816, 12), (1826, 12), (1845, 12), (1856, 12), (1866, 12), (1872, 12), (1873, 12), (1879, 12), (1889, 12), (1902, 12), (1913, 12), (1922, 12), (1923, 12), (1926, 12), (1934, 12), (1949, 12), (1963, 12), (1982, 12), (1992, 12), (1993, 12), (2022, 12), (2053, 12), (2060, 12), (2063, 12), (2065, 12), (2067, 12), (2098, 12), (2101, 12), (2103, 12), (2117, 12), (2127, 12), (2144, 12), (2159, 12), (2160, 12), (2172, 12), (2179, 12), (2190, 12), (2196, 12), (2206, 12), (2254, 12), (2258, 12), (2278, 12), (2294, 12), (2307, 12), (2324, 12), (2328, 12), (2334, 12), (2339, 12), (2340, 12), (2342, 12), (2344, 12), (2346, 12), (2353, 12), (2380, 12), (2388, 12), (2424, 12), (2430, 12), (2431, 12), (2451, 12), (2465, 12), (2466, 12), (2475, 12), (2489, 12), (2492, 12), (2509, 12), (2514, 12), (2521, 12), (2523, 12), (2549, 12), (2557, 12), (2572, 12), (2596, 12), (2607, 12), (2636, 12), (2705, 12), (2712, 12), (2719, 12), (2727, 12), (2735, 12), (2746, 12), (2750, 12), (2764, 12), (2774, 12), (2776, 12), (2779, 12), (2789, 12), (2825, 12), (2827, 12), (2831, 12), (2834, 12), (2836, 12), (2838, 12), (2848, 12), (2866, 12), (2873, 12), (2875, 12), (2879, 12), (2881, 12), (2888, 12),

(2915, 12), (8, 11), (10, 11), (30, 11), (35, 11), (38, 11), (39, 11), (51, 11), (57, 11), (65, 11), (68, 11), (78, 11), (83, 11), (88, 11), (106, 11), (108, 11), (111, 11), (119, 11), (122, 11), (141, 11), (165, 11), (174, 11), (177, 11), (178, 11), (194, 11), (204, 11), (222, 11), (223, 11), (227, 11), (229, 11), (236, 11), (256, 11), (260, 11), (265, 11), (270, 11), (272, 11), (273, 11), (280, 11), (286, 11), (289, 11), (309, 11), (310, 11), (311, 11), (339, 11), (343, 11), (358, 11), (364, 11), (372, 11), (373, 11), (404, 11), (409, 11), (416, 11), (430, 11), (432, 11), (446, 11), (453, 11), (454, 11), (458, 11), (463, 11), (465, 11), (466, 11), (470, 11), (471, 11), (472, 11), (506, 11), (534, 11), (544, 11), (551, 11), (552, 11), (557, 11), (559, 11), (583, 11), (585, 11), (590, 11), (592, 11), (599, 11), (600, 11), (609, 11), (622, 11), (631, 11), (632, 11), (635, 11), (646, 11), (648, 11), (652, 11), (657, 11), (677, 11), (679, 11), (684, 11), (685, 11), (694, 11), (697, 11), (701, 11), (708, 11), (710, 11), (713, 11), (716, 11), (719, 11), (720, 11), (722, 11), (731, 11), (743, 11), (748, 11), (763, 11), (775, 11), (776, 11), (798, 11), (806, 11), (813, 11), (845, 11), (848, 11), (852, 11), (853, 11), (856, 11), (859, 11), (868, 11), (873, 11), (889, 11), (898, 11), (903, 11), (914, 11), (917, 11), (919, 11), (921, 11), (926, 11), (930, 11), (937, 11), (943, 11), (947, 11), (951, 11), (955, 11), (971, 11), (981, 11), (982, 11), (1009, 11), (1015, 11), (1023, 11), (1033, 11), (1038, 11), (1039, 11), (1044, 11), (1046, 11), (1050, 11), (1070, 11), (1075, 11), (1082, 11), (1089, 11), (1102, 11), (1116, 11), (1125, 11), (1127, 11), (1132, 11), (1133, 11), (1134, 11), (1136, 11), (1144, 11), (1147, 11), (1154, 11), (1155, 11), (1156, 11), (1177, 11), (1187, 11), (1207, 11), (1216, 11), (1218, 11), (1228, 11), (1237, 11), (1240, 11), (1243, 11), (1251, 11), (1257, 11), (1271, 11), (1276, 11), (1280, 11), (1283, 11), (1292, 11), (1307, 11), (1310, 11), (1318, 11), (1326, 11), (1336, 11), (1344, 11), (1345, 11), (1356, 11), (1361, 11), (1364, 11), (1378, 11), (1384, 11), (1407, 11), (1413, 11), (1417, 11), (1421, 11), (1423, 11), (1434, 11), (1435, 11), (1443, 11), (1450, 11), (1462, 11), (1463, 11), (1467, 11), (1468, 11), (1474, 11), (1476, 11), (1480, 11), (1481, 11), (1496, 11), (1504, 11), (1507, 11), (1517, 11), (1527, 11), (1531, 11), (1534, 11), (1548, 11), (1572, 11), (1579, 11), (1593, 11), (1594, 11), (1600, 11), (1625, 11), (1628, 11), (1636, 11), (1640, 11), (1646, 11), (1657, 11), (1660, 11), (1664, 11), (1665, 11), (1668, 11), (1688, 11), (1693, 11), (1698, 11), (1700, 11), (1704, 11), (1708, 11), (1712, 11), (1732, 11), (1744, 11), (1745, 11), (1754, 11), (1758, 11), (1760, 11), (1794, 11), (1795, 11), (1796, 11), (1819, 11), (1821, 11), (1822, 11), (1825, 11), (1841, 11), (1842, 11), (1852, 11), (1857, 11), (1859, 11), (1875, 11), (1876, 11), (1877, 11), (1899, 11), (1904, 11), (1911, 11), (1916, 11), (1931, 11), (1941, 11), (1943, 11), (1947, 11), (1990, 11), (1991, 11), (1997, 11), (2007, 11), (2015, 11), (2020, 11), (2021, 11), (2026, 11), (2027, 11), (2038, 11), (2040, 11), (2043, 11), (2052, 11), (2076, 11), (2087, 11), (2090, 11), (2094, 11), (2107, 11), (2109, 11), (2114, 11), (2121, 11), (2136, 11), (2154, 11), (2167, 11), (2168, 11), (2178, 11), (2180, 11), (2181, 11), (2192, 11), (2200, 11), (2208, 11), (2210, 11), (2212, 11), (2232, 11), (2234, 11), (2238, 11), (2243, 11), (2246, 11), (2250, 11), (2253, 11), (2257, 11), (2262, 11), (2270, 11), (2284, 11), (2286, 11), (2287, 11), (2290, 11), (2301, 11), (2303, 11), (2305, 11), (2313, 11), (2326, 11), (2332, 11), (2343, 11), (2354, 11), (2361, 11), (2370, 11), (2374, 11), (2378, 11),

(2381, 11), (2382, 11), (2386, 11), (2396, 11), (2399, 11), (2402, 11), (2405, 11), (2416, 11), (2425, 11), (2426, 11), (2429, 11), (2442, 11), (2445, 11), (2448, 11), (2452, 11), (2459, 11), (2474, 11), (2484, 11), (2485, 11), (2490, 11), (2495, 11), (2498, 11), (2504, 11), (2506, 11), (2513, 11), (2524, 11), (2535, 11), (2540, 11), (2541, 11), (2545, 11), (2547, 11), (2553, 11), (2561, 11), (2569, 11), (2573, 11), (2577, 11), (2582, 11), (2591, 11), (2608, 11), (2618, 11), (2620, 11), (2623, 11), (2625, 11), (2626, 11), (2630, 11), (2642, 11), (2645, 11), (2646, 11), (2656, 11), (2659, 11), (2677, 11), (2678, 11), (2679, 11), (2683, 11), (2685, 11), (2688, 11), (2697, 11), (2700, 11), (2703, 11), (2707, 11), (2714, 11), (2725, 11), (2736, 11), (2738, 11), (2743, 11), (2773, 11), (2795, 11), (2797, 11), (2801, 11), (2806, 11), (2809, 11), (2811, 11), (2815, 11), (2845, 11), (2846, 11), (2852, 11), (2863, 11), (2878, 11), (2883, 11), (2896, 11), (2897, 11), (2899, 11), (2905, 11), (2908, 11), (2920, 11), (2929, 11), (2937, 11), (12, 10), (17, 10), (49, 10), (56, 10), (74, 10), (85, 10), (91, 10), (115, 10), (142, 10), (145, 10), (152, 10), (154, 10), (169, 10), (179, 10), (191, 10), (207, 10), (208, 10), (220, 10), (234, 10), (258, 10), (267, 10), (279, 10), (284, 10), (301, 10), (308, 10), (315, 10), (333, 10), (341, 10), (346, 10), (371, 10), (374, 10), (375, 10), (391, 10), (403, 10), (415, 10), (417, 10), (423, 10), (425, 10), (431, 10), (433, 10), (442, 10), (491, 10), (500, 10), (502, 10), (513, 10), (523, 10), (538, 10), (541, 10), (573, 10), (581, 10), (621, 10), (624, 10), (625, 10), (628, 10), (654, 10), (662, 10), (668, 10), (689, 10), (690, 10), (700, 10), (723, 10), (725, 10), (734, 10), (745, 10), (788, 10), (793, 10), (795, 10), (802, 10), (815, 10), (819, 10), (825, 10), (828, 10), (838, 10), (839, 10), (841, 10), (923, 10), (925, 10), (938, 10), (964, 10), (967, 10), (975, 10), (989, 10), (990, 10), (996, 10), (1008, 10), (1020, 10), (1026, 10), (1047, 10), (1051, 10), (1055, 10), (1067, 10), (1079, 10), (1099, 10), (1119, 10), (1137, 10), (1138, 10), (1140, 10), (1150, 10), (1170, 10), (1174, 10), (1196, 10), (1203, 10), (1212, 10), (1214, 10), (1217, 10), (1225, 10), (1229, 10), (1245, 10), (1258, 10), (1259, 10), (1261, 10), (1266, 10), (1270, 10), (1294, 10), (1304, 10), (1314, 10), (1327, 10), (1329, 10), (1330, 10), (1341, 10), (1367, 10), (1374, 10), (1394, 10), (1396, 10), (1402, 10), (1406, 10), (1416, 10), (1425, 10), (1430, 10), (1464, 10), (1469, 10), (1470, 10), (1472, 10), (1477, 10), (1488, 10), (1523, 10), (1532, 10), (1553, 10), (1577, 10), (1587, 10), (1605, 10), (1641, 10), (1675, 10), (1676, 10), (1699, 10), (1703, 10), (1706, 10), (1716, 10), (1730, 10), (1739, 10), (1743, 10), (1747, 10), (1748, 10), (1750, 10), (1756, 10), (1778, 10), (1792, 10), (1793, 10), (1809, 10), (1824, 10), (1878, 10), (1881, 10), (1883, 10), (1896, 10), (1897, 10), (1898, 10), (1905, 10), (1906, 10), (1939, 10), (1944, 10), (1956, 10), (1969, 10), (1972, 10), (1975, 10), (1983, 10), (1986, 10), (1998, 10), (2006, 10), (2009, 10), (2036, 10), (2037, 10), (2039, 10), (2051, 10), (2056, 10), (2068, 10), (2084, 10), (2091, 10), (2126, 10), (2177, 10), (2188, 10), (2197, 10), (2216, 10), (2219, 10), (2226, 10), (2233, 10), (2237, 10), (2289, 10), (2293, 10), (2311, 10), (2318, 10), (2336, 10), (2351, 10), (2355, 10), (2363, 10), (2403, 10), (2444, 10), (2454, 10), (2461, 10), (2486, 10), (2508, 10), (2522, 10), (2563, 10), (2566, 10), (2578, 10), (2584, 10), (2586, 10), (2588, 10), (2592, 10), (2593, 10), (2606, 10), (2634, 10), (2649, 10), (2650, 10), (2662, 10), (2667, 10), (2671, 10), (2689, 10), (2695, 10), (2699, 10), (2724, 10), (2737, 10), (2747, 10),

(2760, 10), (2768, 10), (2777, 10), (2805, 10), (2807, 10), (2857, 10), (2859, 10), (2891, 10), (2892, 10), (2901, 10), (2912, 10), (2925, 10), (2935, 10), (2940, 10), (5, 9), (28, 9), (69, 9), (76, 9), (96, 9), (136, 9), (149, 9), (181, 9), (190, 9), (206, 9), (239, 9), (261, 9), (277, 9), (297, 9), (324, 9), (327, 9), (344, 9), (369, 9), (412, 9), (428, 9), (437, 9), (460, 9), (494, 9), (503, 9), (505, 9), (524, 9), (525, 9), (539, 9), (543, 9), (562, 9), (603, 9), (607, 9), (699, 9), (755, 9), (805, 9), (807, 9), (811, 9), (860, 9), (879, 9), (912, 9), (933, 9), (1007, 9), (1052, 9), (1080, 9), (1106, 9), (1113, 9), (1157, 9), (1159, 9), (1169, 9), (1186, 9), (1235, 9), (1260, 9), (1274, 9), (1287, 9), (1293, 9), (1302, 9), (1343, 9), (1348, 9), (1420, 9), (1431, 9), (1541, 9), (1558, 9), (1623, 9), (1638, 9), (1644, 9), (1669, 9), (1673, 9), (1683, 9), (1692, 9), (1749, 9), (1770, 9), (1812, 9), (1814, 9), (1843, 9), (1864, 9), (1865, 9), (1870, 9), (1880, 9), (1936, 9), (1973, 9), (1981, 9), (2013, 9), (2024, 9), (2055, 9), (2057, 9), (2066, 9), (2077, 9), (2093, 9), (2100, 9), (2161, 9), (2187, 9), (2189, 9), (2222, 9), (2235, 9), (2266, 9), (2299, 9), (2335, 9), (2345, 9), (2384, 9), (2411, 9), (2423, 9), (2457, 9), (2462, 9), (2494, 9), (2505, 9), (2510, 9), (2615, 9), (2633, 9), (2652, 9), (2693, 9), (2708, 9), (2741, 9), (2744, 9), (2766, 9), (2769, 9), (2778, 9), (2794, 9), (2813, 9), (2850, 9), (2898, 9), (2900, 9), (2938, 9), (1, 8), (9, 8), (50, 8), (59, 8), (72, 8), (147, 8), (159, 8), (246, 8), (254, 8), (269, 8), (321, 8), (331, 8), (349, 8), (359, 8), (361, 8), (400, 8), (420, 8), (421, 8), (422, 8), (516, 8), (550, 8), (610, 8), (620, 8), (633, 8), (762, 8), (765, 8), (782, 8), (790, 8), (846, 8), (907, 8), (911, 8), (916, 8), (931, 8), (950, 8), (973, 8), (992, 8), (1130, 8), (1192, 8), (1205, 8), (1226, 8), (1273, 8), (1295, 8), (1324, 8), (1387, 8), (1401, 8), (1405, 8), (1432, 8), (1441, 8), (1486, 8), (1499, 8), (1505, 8), (1508, 8), (1639, 8), (1681, 8), (1682, 8), (1695, 8), (1727, 8), (1734, 8), (1738, 8), (1786, 8), (1790, 8), (1799, 8), (1802, 8), (1861, 8), (1888, 8), (1901, 8), (1909, 8), (1932, 8), (1988, 8), (2008, 8), (2035, 8), (2041, 8), (2062, 8), (2071, 8), (2072, 8), (2086, 8), (2120, 8), (2131, 8), (2145, 8), (2153, 8), (2209, 8), (2224, 8), (2259, 8), (2265, 8), (2285, 8), (2297, 8), (2298, 8), (2323, 8), (2352, 8), (2369, 8), (2387, 8), (2390, 8), (2397, 8), (2412, 8), (2418, 8), (2419, 8), (2434, 8), (2443, 8), (2446, 8), (2529, 8), (2531, 8), (2543, 8), (2544, 8), (2548, 8), (2609, 8), (2647, 8), (2675, 8), (2686, 8), (2691, 8), (2709, 8), (2715, 8), (2728, 8), (2780, 8), (2791, 8), (2793, 8), (2821, 8), (2882, 8), (2886, 8), (0, 7), (13, 7), (94, 7), (114, 7), (127, 7), (155, 7), (171, 7), (259, 7), (295, 7), (299, 7), (342, 7), (356, 7), (357, 7), (384, 7), (392, 7), (395, 7), (455, 7), (487, 7), (545, 7), (554, 7), (611, 7), (639, 7), (650, 7), (705, 7), (727, 7), (799, 7), (809, 7), (847, 7), (895, 7), (901, 7), (942, 7), (956, 7), (1010, 7), (1021, 7), (1022, 7), (1031, 7), (1054, 7), (1064, 7), (1107, 7), (1165, 7), (1200, 7), (1238, 7), (1312, 7), (1313, 7), (1331, 7), (1412, 7), (1440, 7), (1456, 7), (1457, 7), (1494, 7), (1503, 7), (1537, 7), (1547, 7), (1571, 7), (1592, 7), (1599, 7), (1614, 7), (1637, 7), (1651, 7), (1677, 7), (1710, 7), (1714, 7), (1769, 7), (1779, 7), (1830, 7), (1835, 7), (1918, 7), (1996, 7), (2004, 7), (2019, 7), (2028, 7), (2134, 7), (2186, 7), (2242, 7), (2319, 7), (2393, 7), (2447, 7), (2525, 7), (2551, 7), (2599, 7), (2613, 7), (2681, 7), (2722, 7), (2759, 7), (2782, 7), (2785, 7), (2808, 7), (2822, 7), (2844, 7), (2849, 7), (2869, 7), (2887, 7), (2926, 7), (2933, 7), (2939, 7), (25, 6), (34,

```
6), (40, 6), (64, 6), (66, 6), (97, 6), (130, 6), (132, 6), (198, 6), (199, 6),
(212, 6), (268, 6), (317, 6), (318, 6), (380, 6), (396, 6), (408, 6), (413, 6),
(439, 6), (440, 6), (444, 6), (456, 6), (464, 6), (469, 6), (476, 6), (482, 6),
(490, 6), (499, 6), (520, 6), (522, 6), (526, 6), (546, 6), (556, 6), (561, 6),
(568, 6), (663, 6), (688, 6), (704, 6), (706, 6), (718, 6), (761, 6), (766, 6),
(781, 6), (783, 6), (785, 6), (786, 6), (824, 6), (826, 6), (834, 6), (876, 6),
(896, 6), (897, 6), (904, 6), (908, 6), (920, 6), (932, 6), (941, 6), (966, 6),
(968, 6), (1000, 6), (1002, 6), (1004, 6), (1053, 6), (1083, 6), (1084, 6),
(1095, 6), (1129, 6), (1152, 6), (1206, 6), (1227, 6), (1236, 6), (1241, 6),
(1298, 6), (1301, 6), (1308, 6), (1309, 6), (1311, 6), (1316, 6), (1334, 6),
(1338, 6), (1370, 6), (1433, 6), (1452, 6), (1466, 6), (1473, 6), (1482, 6),
(1533, 6), (1536, 6), (1538, 6), (1552, 6), (1585, 6), (1629, 6), (1634, 6),
(1647, 6), (1654, 6), (1655, 6), (1680, 6), (1684, 6), (1687, 6), (1701, 6),
(1711, 6), (1715, 6), (1718, 6), (1722, 6), (1733, 6), (1742, 6), (1752, 6),
(1753, 6), (1771, 6), (1783, 6), (1784, 6), (1785, 6), (1789, 6), (1831, 6),
(1869, 6), (1871, 6), (1884, 6), (1900, 6), (1907, 6), (1908, 6), (1914, 6),
(1929, 6), (1933, 6), (1942, 6), (1964, 6), (1965, 6), (1976, 6), (1987, 6),
(2005, 6), (2011, 6), (2029, 6), (2045, 6), (2061, 6), (2081, 6), (2095, 6),
(2099, 6), (2111, 6), (2123, 6), (2124, 6), (2130, 6), (2141, 6), (2146, 6),
(2165, 6), (2198, 6), (2241, 6), (2255, 6), (2267, 6), (2277, 6), (2280, 6),
(2317, 6), (2325, 6), (2329, 6), (2337, 6), (2348, 6), (2357, 6), (2366, 6),
(2415, 6), (2455, 6), (2463, 6), (2500, 6), (2503, 6), (2511, 6), (2526, 6),
(2597, 6), (2635, 6), (2673, 6), (2690, 6), (2692, 6), (2704, 6), (2711, 6),
(2721, 6), (2734, 6), (2823, 6), (2829, 6), (2837, 6), (2865, 6), (2871, 6),
(2874, 6), (2877, 6), (2884, 6), (2889, 6), (2893, 6), (2909, 6), (2934, 6),
(32, 5), (353, 5), (980, 5), (1016, 5), (1072, 5), (1557, 5), (1043, 4), (2231,
4), (2611, 4), (2775, 4), (1694, 3)]
```

[ ]:
```
for key,value in l.items():
  l[key]=(value/(x_mord.shape[0]))*100
```

[ ]:
```
x=sorted(l.items(),key=lambda x:x[1],reverse=True)
```

[ ]:
```
print(x)
```

```
[('MDEN-11', 97.8600543478261), ('MDEN-13', 97.48641304347827), ('MDEN-33',
96.33152173913044), ('MDEN-23', 91.16847826086956), ('MDEC-44',
89.50407608695652), ('MDEN-12', 83.25407608695652), ('MDEN-22',
81.92934782608695), ('MDEC-14', 80.53668478260869), ('MDEC-24', 75.78125),
('MDEC-34', 75.54347826086956), ('MDEO-22', 73.09782608695652), ('MDEC-11',
58.22010869565217), ('MDEO-12', 47.75815217391305), ('MDEC-12',
33.76358695652174), ('MDEC-13', 33.59375), ('AATS8s', 28.43070652173913),
('AATSC8s', 28.43070652173913), ('MATS8s', 28.43070652173913), ('GATS8s',
28.43070652173913), ('AATS7s', 24.69429347826087), ('AATSC7s',
24.69429347826087), ('MATS7s', 24.69429347826087), ('GATS7s',
24.69429347826087), ('AATS6s', 21.773097826086957), ('AATSC6s',
21.773097826086957), ('MATS6s', 21.773097826086957), ('GATS6s',
21.773097826086957), ('BCUTc-1h', 21.195652173913043), ('BCUTc-1l',
```

21.195652173913043), ('SpAbs_A', 21.161684782608695), ('SpMax_A', 21.161684782608695), ('SpDiam_A', 21.161684782608695), ('SpAD_A', 21.161684782608695), ('SpMAD_A', 21.161684782608695), ('LogEE_A', 21.161684782608695), ('VE1_A', 21.161684782608695), ('VE2_A', 21.161684782608695), ('VE3_A', 21.161684782608695), ('VR1_A', 21.161684782608695), ('VR2_A', 21.161684782608695), ('VR3_A', 21.161684782608695), ('BCUTdv-1h', 21.161684782608695), ('BCUTdv-1l', 21.161684782608695), ('BCUTd-1h', 21.161684782608695), ('BCUTd-1l', 21.161684782608695), ('BCUTs-1h', 21.161684782608695), ('BCUTs-1l', 21.161684782608695), ('BCUTZ-1h', 21.161684782608695), ('BCUTZ-1l', 21.161684782608695), ('BCUTm-1h', 21.161684782608695), ('BCUTm-1l', 21.161684782608695), ('BCUTv-1h', 21.161684782608695), ('BCUTv-1l', 21.161684782608695), ('BCUTse-1h', 21.161684782608695), ('BCUTse-1l', 21.161684782608695), ('BCUTpe-1h', 21.161684782608695), ('BCUTpe-1l', 21.161684782608695), ('BCUTare-1h', 21.161684782608695), ('BCUTare-1l', 21.161684782608695), ('BCUTp-1h', 21.161684782608695), ('BCUTp-1l', 21.161684782608695), ('BCUTi-1h', 21.161684782608695), ('BCUTi-1l', 21.161684782608695), ('SpAbs_DzZ', 21.161684782608695), ('SpMax_DzZ', 21.161684782608695), ('SpDiam_DzZ', 21.161684782608695), ('SpAD_DzZ', 21.161684782608695), ('SpMAD_DzZ', 21.161684782608695), ('LogEE_DzZ', 21.161684782608695), ('SM1_DzZ', 21.161684782608695), ('VE1_DzZ', 21.161684782608695), ('VE2_DzZ', 21.161684782608695), ('VE3_DzZ', 21.161684782608695), ('VR1_DzZ', 21.161684782608695), ('VR2_DzZ', 21.161684782608695), ('VR3_DzZ', 21.161684782608695), ('SpAbs_Dzm', 21.161684782608695), ('SpMax_Dzm', 21.161684782608695), ('SpDiam_Dzm', 21.161684782608695), ('SpAD_Dzm', 21.161684782608695), ('SpMAD_Dzm', 21.161684782608695), ('LogEE_Dzm', 21.161684782608695), ('SM1_Dzm', 21.161684782608695), ('VE1_Dzm', 21.161684782608695), ('VE2_Dzm', 21.161684782608695), ('VE3_Dzm', 21.161684782608695), ('VR1_Dzm', 21.161684782608695), ('VR2_Dzm', 21.161684782608695), ('VR3_Dzm', 21.161684782608695), ('SpAbs_Dzv', 21.161684782608695), ('SpMax_Dzv', 21.161684782608695), ('SpDiam_Dzv', 21.161684782608695), ('SpAD_Dzv', 21.161684782608695), ('SpMAD_Dzv', 21.161684782608695), ('LogEE_Dzv', 21.161684782608695), ('SM1_Dzv', 21.161684782608695), ('VE1_Dzv', 21.161684782608695), ('VE2_Dzv', 21.161684782608695), ('VE3_Dzv', 21.161684782608695), ('VR1_Dzv', 21.161684782608695), ('VR2_Dzv', 21.161684782608695), ('VR3_Dzv', 21.161684782608695), ('SpAbs_Dzse', 21.161684782608695), ('SpMax_Dzse', 21.161684782608695), ('SpDiam_Dzse', 21.161684782608695), ('SpAD_Dzse', 21.161684782608695), ('SpMAD_Dzse', 21.161684782608695), ('LogEE_Dzse', 21.161684782608695), ('SM1_Dzse', 21.161684782608695), ('VE1_Dzse', 21.161684782608695), ('VE2_Dzse', 21.161684782608695), ('VE3_Dzse', 21.161684782608695), ('VR1_Dzse', 21.161684782608695), ('VR2_Dzse', 21.161684782608695), ('VR3_Dzse', 21.161684782608695), ('SpAbs_Dzpe', 21.161684782608695), ('SpMax_Dzpe', 21.161684782608695), ('SpDiam_Dzpe', 21.161684782608695), ('SpAD_Dzpe', 21.161684782608695), ('SpMAD_Dzpe', 21.161684782608695), ('LogEE_Dzpe', 21.161684782608695), ('SM1_Dzpe', 21.161684782608695), ('VE1_Dzpe', 21.161684782608695), ('VE2_Dzpe', 21.161684782608695), ('VE3_Dzpe',

21.161684782608695), ('VR1_Dzpe', 21.161684782608695), ('VR2_Dzpe', 21.161684782608695), ('VR3_Dzpe', 21.161684782608695), ('SpAbs_Dzare', 21.161684782608695), ('SpMax_Dzare', 21.161684782608695), ('SpDiam_Dzare', 21.161684782608695), ('SpAD_Dzare', 21.161684782608695), ('SpMAD_Dzare', 21.161684782608695), ('LogEE_Dzare', 21.161684782608695), ('SM1_Dzare', 21.161684782608695), ('VE1_Dzare', 21.161684782608695), ('VE2_Dzare', 21.161684782608695), ('VE3_Dzare', 21.161684782608695), ('VR1_Dzare', 21.161684782608695), ('VR2_Dzare', 21.161684782608695), ('VR3_Dzare', 21.161684782608695), ('SpAbs_Dzp', 21.161684782608695), ('SpMax_Dzp', 21.161684782608695), ('SpDiam_Dzp', 21.161684782608695), ('SpAD_Dzp', 21.161684782608695), ('SpMAD_Dzp', 21.161684782608695), ('LogEE_Dzp', 21.161684782608695), ('SM1_Dzp', 21.161684782608695), ('VE1_Dzp', 21.161684782608695), ('VE2_Dzp', 21.161684782608695), ('VE3_Dzp', 21.161684782608695), ('VR1_Dzp', 21.161684782608695), ('VR2_Dzp', 21.161684782608695), ('VR3_Dzp', 21.161684782608695), ('SpAbs_Dzi', 21.161684782608695), ('SpMax_Dzi', 21.161684782608695), ('SpDiam_Dzi', 21.161684782608695), ('SpAD_Dzi', 21.161684782608695), ('SpMAD_Dzi', 21.161684782608695), ('LogEE_Dzi', 21.161684782608695), ('SM1_Dzi', 21.161684782608695), ('VE1_Dzi', 21.161684782608695), ('VE2_Dzi', 21.161684782608695), ('VE3_Dzi', 21.161684782608695), ('VR1_Dzi', 21.161684782608695), ('VR2_Dzi', 21.161684782608695), ('VR3_Dzi', 21.161684782608695), ('SpAbs_D', 21.161684782608695), ('SpMax_D', 21.161684782608695), ('SpDiam_D', 21.161684782608695), ('SpAD_D', 21.161684782608695), ('SpMAD_D', 21.161684782608695), ('LogEE_D', 21.161684782608695), ('VE1_D', 21.161684782608695), ('VE2_D', 21.161684782608695), ('VE3_D', 21.161684782608695), ('VR1_D', 21.161684782608695), ('VR2_D', 21.161684782608695), ('VR3_D', 21.161684782608695), ('ETA_alpha', 21.161684782608695), ('AETA_alpha', 21.161684782608695), ('ETA_shape_p', 21.161684782608695), ('ETA_shape_y', 21.161684782608695), ('ETA_shape_x', 21.161684782608695), ('ETA_beta', 21.161684782608695), ('AETA_beta', 21.161684782608695), ('ETA_beta_s', 21.161684782608695), ('AETA_beta_s', 21.161684782608695), ('ETA_beta_ns', 21.161684782608695), ('AETA_beta_ns', 21.161684782608695), ('ETA_beta_ns_d', 21.161684782608695), ('AETA_beta_ns_d', 21.161684782608695), ('ETA_eta', 21.161684782608695), ('AETA_eta', 21.161684782608695), ('ETA_eta_L', 21.161684782608695), ('AETA_eta_L', 21.161684782608695), ('ETA_eta_R', 21.161684782608695), ('AETA_eta_R', 21.161684782608695), ('ETA_eta_RL', 21.161684782608695), ('AETA_eta_RL', 21.161684782608695), ('ETA_eta_F', 21.161684782608695), ('AETA_eta_F', 21.161684782608695), ('ETA_eta_FL', 21.161684782608695), ('AETA_eta_FL', 21.161684782608695), ('ETA_eta_B', 21.161684782608695), ('AETA_eta_B', 21.161684782608695), ('ETA_eta_BR', 21.161684782608695), ('AETA_eta_BR', 21.161684782608695), ('ETA_dAlpha_A', 21.161684782608695), ('ETA_dAlpha_B', 21.161684782608695), ('ETA_epsilon_1', 21.161684782608695), ('ETA_epsilon_2', 21.161684782608695), ('ETA_epsilon_3', 21.161684782608695), ('ETA_epsilon_4', 21.161684782608695), ('ETA_epsilon_5', 21.161684782608695), ('ETA_dEpsilon_A', 21.161684782608695), ('ETA_dEpsilon_B', 21.161684782608695), ('ETA_dEpsilon_C', 21.161684782608695), ('ETA_dEpsilon_D', 21.161684782608695), ('ETA_dBeta', 21.161684782608695), ('AETA_dBeta',

21.161684782608695), ('ETA_psi_1', 21.161684782608695), ('ETA_dPsi_A',
21.161684782608695), ('ETA_dPsi_B', 21.161684782608695), ('MID',
21.161684782608695), ('AMID', 21.161684782608695), ('MID_h',
21.161684782608695), ('AMID_h', 21.161684782608695), ('MID_C',
21.161684782608695), ('AMID_C', 21.161684782608695), ('MID_N',
21.161684782608695), ('AMID_N', 21.161684782608695), ('MID_O',
21.161684782608695), ('AMID_O', 21.161684782608695), ('MID_X',
21.161684782608695), ('AMID_X', 21.161684782608695), ('AATS5s',
20.652173913043477), ('AATSC5s', 20.652173913043477), ('MATS5s',
20.652173913043477), ('GATS5s', 20.652173913043477), ('AATS4s',
20.14266304347826), ('AATSC4s', 20.14266304347826), ('MATS4s',
20.14266304347826), ('GATS4s', 20.14266304347826), ('AATS3s',
20.040760869565215), ('AATSC3s', 20.040760869565215), ('MATS3s',
20.040760869565215), ('GATS3s', 20.040760869565215), ('ATS0s',
19.938858695652172), ('ATS1s', 19.938858695652172), ('ATS2s',
19.938858695652172), ('ATS3s', 19.938858695652172), ('ATS4s',
19.938858695652172), ('ATS5s', 19.938858695652172), ('ATS6s',
19.938858695652172), ('ATS7s', 19.938858695652172), ('ATS8s',
19.938858695652172), ('AATS0s', 19.938858695652172), ('AATS1s',
19.938858695652172), ('AATS2s', 19.938858695652172), ('ATSC0s',
19.938858695652172), ('ATSC1s', 19.938858695652172), ('ATSC2s',
19.938858695652172), ('ATSC3s', 19.938858695652172), ('ATSC4s',
19.938858695652172), ('ATSC5s', 19.938858695652172), ('ATSC6s',
19.938858695652172), ('ATSC7s', 19.938858695652172), ('ATSC8s',
19.938858695652172), ('AATSC0s', 19.938858695652172), ('AATSC1s',
19.938858695652172), ('AATSC2s', 19.938858695652172), ('MATS1s',
19.938858695652172), ('MATS2s', 19.938858695652172), ('GATS1s',
19.938858695652172), ('GATS2s', 19.938858695652172), ('Xp-0d',
19.938858695652172), ('AXp-0d', 19.938858695652172), ('mZagreb1',
19.938858695652172), ('MDEO-11', 18.376358695652172), ('Vabc',
17.32336956521739), ('Xp-0dv', 17.05163043478261), ('AXp-0dv',
17.05163043478261), ('AATSC8c', 16.338315217391305), ('MATS8c',
16.338315217391305), ('GATS8c', 16.338315217391305), ('AATS8dv',
16.27038043478261), ('AATS8d', 16.27038043478261), ('AATS8Z',
16.27038043478261), ('AATS8m', 16.27038043478261), ('AATS8v',
16.27038043478261), ('AATS8se', 16.27038043478261), ('AATS8pe',
16.27038043478261), ('AATS8are', 16.27038043478261), ('AATS8p',
16.27038043478261), ('AATS8i', 16.27038043478261), ('AATSC8dv',
16.27038043478261), ('AATSC8d', 16.27038043478261), ('AATSC8Z',
16.27038043478261), ('AATSC8m', 16.27038043478261), ('AATSC8v',
16.27038043478261), ('AATSC8se', 16.27038043478261), ('AATSC8pe',
16.27038043478261), ('AATSC8are', 16.27038043478261), ('AATSC8p',
16.27038043478261), ('AATSC8i', 16.27038043478261), ('MATS8dv',
16.27038043478261), ('MATS8d', 16.27038043478261), ('MATS8Z',
16.27038043478261), ('MATS8m', 16.27038043478261), ('MATS8v',
16.27038043478261), ('MATS8se', 16.27038043478261), ('MATS8pe',
16.27038043478261), ('MATS8are', 16.27038043478261), ('MATS8p',
16.27038043478261), ('MATS8i', 16.27038043478261), ('GATS8dv',

16.27038043478261), ('GATS8d', 16.27038043478261), ('GATS8Z',
16.27038043478261), ('GATS8m', 16.27038043478261), ('GATS8v',
16.27038043478261), ('GATS8se', 16.27038043478261), ('GATS8pe',
16.27038043478261), ('GATS8are', 16.27038043478261), ('GATS8p',
16.27038043478261), ('GATS8i', 16.27038043478261), ('MDEC-33',
8.457880434782608), ('AXp-7d', 7.914402173913043), ('AXp-7dv',
7.914402173913043), ('MDEC-22', 7.574728260869565), ('AATSC7c',
7.133152173913043), ('MATS7c', 7.133152173913043), ('GATS7c',
7.133152173913043), ('AATS7dv', 7.065217391304348), ('AATS7d',
7.065217391304348), ('AATS7Z', 7.065217391304348), ('AATS7m',
7.065217391304348), ('AATS7v', 7.065217391304348), ('AATS7se',
7.065217391304348), ('AATS7pe', 7.065217391304348), ('AATS7are',
7.065217391304348), ('AATS7p', 7.065217391304348), ('AATS7i',
7.065217391304348), ('AATSC7dv', 7.065217391304348), ('AATSC7d',
7.065217391304348), ('AATSC7Z', 7.065217391304348), ('AATSC7m',
7.065217391304348), ('AATSC7v', 7.065217391304348), ('AATSC7se',
7.065217391304348), ('AATSC7pe', 7.065217391304348), ('AATSC7are',
7.065217391304348), ('AATSC7p', 7.065217391304348), ('AATSC7i',
7.065217391304348), ('MATS7dv', 7.065217391304348), ('MATS7d',
7.065217391304348), ('MATS7Z', 7.065217391304348), ('MATS7m',
7.065217391304348), ('MATS7v', 7.065217391304348), ('MATS7se',
7.065217391304348), ('MATS7pe', 7.065217391304348), ('MATS7are',
7.065217391304348), ('MATS7p', 7.065217391304348), ('MATS7i',
7.065217391304348), ('GATS7dv', 7.065217391304348), ('GATS7d',
7.065217391304348), ('GATS7Z', 7.065217391304348), ('GATS7m',
7.065217391304348), ('GATS7v', 7.065217391304348), ('GATS7se',
7.065217391304348), ('GATS7pe', 7.065217391304348), ('GATS7are',
7.065217391304348), ('GATS7p', 7.065217391304348), ('GATS7i',
7.065217391304348), ('MDEC-23', 5.264945652173913), ('AXp-6d',
4.789402173913043), ('AXp-6dv', 4.789402173913043), ('AXp-5d',
2.989130434782609), ('AXp-5dv', 2.989130434782609), ('AATSC6c',
2.921195652173913), ('MATS6c', 2.921195652173913), ('GATS6c',
2.921195652173913), ('AATS6dv', 2.8532608695652173), ('AATS6d',
2.8532608695652173), ('AATS6Z', 2.8532608695652173), ('AATS6m',
2.8532608695652173), ('AATS6v', 2.8532608695652173), ('AATS6se',
2.8532608695652173), ('AATS6pe', 2.8532608695652173), ('AATS6are',
2.8532608695652173), ('AATS6p', 2.8532608695652173), ('AATS6i',
2.8532608695652173), ('AATSC6dv', 2.8532608695652173), ('AATSC6d',
2.8532608695652173), ('AATSC6Z', 2.8532608695652173), ('AATSC6m',
2.8532608695652173), ('AATSC6v', 2.8532608695652173), ('AATSC6se',
2.8532608695652173), ('AATSC6pe', 2.8532608695652173), ('AATSC6are',
2.8532608695652173), ('AATSC6p', 2.8532608695652173), ('AATSC6i',
2.8532608695652173), ('MATS6dv', 2.8532608695652173), ('MATS6d',
2.8532608695652173), ('MATS6Z', 2.8532608695652173), ('MATS6m',
2.8532608695652173), ('MATS6v', 2.8532608695652173), ('MATS6se',
2.8532608695652173), ('MATS6pe', 2.8532608695652173), ('MATS6are',
2.8532608695652173), ('MATS6p', 2.8532608695652173), ('MATS6i',
2.8532608695652173), ('GATS6dv', 2.8532608695652173), ('GATS6d',

2.8532608695652173), ('GATS6Z', 2.8532608695652173), ('GATS6m', 2.8532608695652173), ('GATS6v', 2.8532608695652173), ('GATS6se', 2.8532608695652173), ('GATS6pe', 2.8532608695652173), ('GATS6are', 2.8532608695652173), ('GATS6p', 2.8532608695652173), ('GATS6i', 2.8532608695652173), ('AXp-4d', 1.8342391304347827), ('AXp-4dv', 1.8342391304347827), ('AATSC5c', 1.528532608695652), ('MATS5c', 1.528532608695652), ('GATS5c', 1.528532608695652), ('AATS5dv', 1.4605978260869565), ('AATS5d', 1.4605978260869565), ('AATS5Z', 1.4605978260869565), ('AATS5m', 1.4605978260869565), ('AATS5v', 1.4605978260869565), ('AATS5se', 1.4605978260869565), ('AATS5pe', 1.4605978260869565), ('AATS5are', 1.4605978260869565), ('AATS5p', 1.4605978260869565), ('AATS5i', 1.4605978260869565), ('AATSC5dv', 1.4605978260869565), ('AATSC5d', 1.4605978260869565), ('AATSC5Z', 1.4605978260869565), ('AATSC5m', 1.4605978260869565), ('AATSC5v', 1.4605978260869565), ('AATSC5se', 1.4605978260869565), ('AATSC5pe', 1.4605978260869565), ('AATSC5are', 1.4605978260869565), ('AATSC5p', 1.4605978260869565), ('AATSC5i', 1.4605978260869565), ('MATS5dv', 1.4605978260869565), ('MATS5d', 1.4605978260869565), ('MATS5Z', 1.4605978260869565), ('MATS5m', 1.4605978260869565), ('MATS5v', 1.4605978260869565), ('MATS5se', 1.4605978260869565), ('MATS5pe', 1.4605978260869565), ('MATS5are', 1.4605978260869565), ('MATS5p', 1.4605978260869565), ('MATS5i', 1.4605978260869565), ('GATS5dv', 1.4605978260869565), ('GATS5d', 1.4605978260869565), ('GATS5Z', 1.4605978260869565), ('GATS5m', 1.4605978260869565), ('GATS5v', 1.4605978260869565), ('GATS5se', 1.4605978260869565), ('GATS5pe', 1.4605978260869565), ('GATS5are', 1.4605978260869565), ('GATS5p', 1.4605978260869565), ('GATS5i', 1.4605978260869565), ('AXp-3d', 1.0869565217391304), ('AXp-3dv', 1.0869565217391304), ('Kier3', 1.0869565217391304), ('AATSC4c', 0.9171195652173914), ('MATS4c', 0.9171195652173914), ('GATS4c', 0.9171195652173914), ('AATS4dv', 0.8491847826086956), ('AATS4d', 0.8491847826086956), ('AATS4Z', 0.8491847826086956), ('AATS4m', 0.8491847826086956), ('AATS4v', 0.8491847826086956), ('AATS4se', 0.8491847826086956), ('AATS4pe', 0.8491847826086956), ('AATS4are', 0.8491847826086956), ('AATS4p', 0.8491847826086956), ('AATS4i', 0.8491847826086956), ('AATSC4dv', 0.8491847826086956), ('AATSC4d', 0.8491847826086956), ('AATSC4Z', 0.8491847826086956), ('AATSC4m', 0.8491847826086956), ('AATSC4v', 0.8491847826086956), ('AATSC4se', 0.8491847826086956), ('AATSC4pe', 0.8491847826086956), ('AATSC4are', 0.8491847826086956), ('AATSC4p', 0.8491847826086956), ('AATSC4i', 0.8491847826086956), ('MATS4dv', 0.8491847826086956), ('MATS4d', 0.8491847826086956), ('MATS4Z', 0.8491847826086956), ('MATS4m', 0.8491847826086956), ('MATS4v', 0.8491847826086956), ('MATS4se', 0.8491847826086956), ('MATS4pe', 0.8491847826086956), ('MATS4are', 0.8491847826086956), ('MATS4p', 0.8491847826086956), ('MATS4i', 0.8491847826086956), ('GATS4dv', 0.8491847826086956), ('GATS4d', 0.8491847826086956), ('GATS4Z', 0.8491847826086956), ('GATS4m', 0.8491847826086956), ('GATS4v', 0.8491847826086956), ('GATS4se', 0.8491847826086956), ('GATS4pe',

0.8491847826086956), ('GATS4are', 0.8491847826086956), ('GATS4p',
0.8491847826086956), ('GATS4i', 0.8491847826086956), ('AATSC3c', 0.78125),
('MATS3c', 0.78125), ('GATS3c', 0.78125), ('HybRatio', 0.78125), ('MATS3d',
0.7472826086956522), ('GATS3d', 0.7472826086956522), ('AATS3dv',
0.7133152173913043), ('AATS3d', 0.7133152173913043), ('AATS3Z',
0.7133152173913043), ('AATS3m', 0.7133152173913043), ('AATS3v',
0.7133152173913043), ('AATS3se', 0.7133152173913043), ('AATS3pe',
0.7133152173913043), ('AATS3are', 0.7133152173913043), ('AATS3p',
0.7133152173913043), ('AATS3i', 0.7133152173913043), ('AATSC3dv',
0.7133152173913043), ('AATSC3d', 0.7133152173913043), ('AATSC3Z',
0.7133152173913043), ('AATSC3m', 0.7133152173913043), ('AATSC3v',
0.7133152173913043), ('AATSC3se', 0.7133152173913043), ('AATSC3pe',
0.7133152173913043), ('AATSC3are', 0.7133152173913043), ('AATSC3p',
0.7133152173913043), ('AATSC3i', 0.7133152173913043), ('MATS3dv',
0.7133152173913043), ('MATS3Z', 0.7133152173913043), ('MATS3m',
0.7133152173913043), ('MATS3v', 0.7133152173913043), ('MATS3se',
0.7133152173913043), ('MATS3pe', 0.7133152173913043), ('MATS3are',
0.7133152173913043), ('MATS3p', 0.7133152173913043), ('MATS3i',
0.7133152173913043), ('GATS3dv', 0.7133152173913043), ('GATS3Z',
0.7133152173913043), ('GATS3m', 0.7133152173913043), ('GATS3v',
0.7133152173913043), ('GATS3se', 0.7133152173913043), ('GATS3pe',
0.7133152173913043), ('GATS3are', 0.7133152173913043), ('GATS3p',
0.7133152173913043), ('GATS3i', 0.7133152173913043), ('AXp-2d',
0.5095108695652174), ('AXp-2dv', 0.5095108695652174), ('Kier2',
0.5095108695652174), ('AATSC2c', 0.4755434782608696), ('MATS2c',
0.4755434782608696), ('MATS2d', 0.4755434782608696), ('GATS2c',
0.4755434782608696), ('GATS2d', 0.4755434782608696), ('AATSC1c',
0.44157608695652173), ('MATS1c', 0.44157608695652173), ('MATS1d',
0.44157608695652173), ('GATS1c', 0.44157608695652173), ('GATS1d',
0.44157608695652173), ('AXp-1d', 0.44157608695652173), ('AXp-1dv',
0.44157608695652173), ('Kier1', 0.44157608695652173), ('RotRatio',
0.44157608695652173), ('VAdjMat', 0.44157608695652173), ('AATS2dv',
0.4076086956521739), ('AATS2d', 0.4076086956521739), ('AATS2Z',
0.4076086956521739), ('AATS2m', 0.4076086956521739), ('AATS2v',
0.4076086956521739), ('AATS2se', 0.4076086956521739), ('AATS2pe',
0.4076086956521739), ('AATS2are', 0.4076086956521739), ('AATS2p',
0.4076086956521739), ('AATS2i', 0.4076086956521739), ('AATSC2dv',
0.4076086956521739), ('AATSC2d', 0.4076086956521739), ('AATSC2Z',
0.4076086956521739), ('AATSC2m', 0.4076086956521739), ('AATSC2v',
0.4076086956521739), ('AATSC2se', 0.4076086956521739), ('AATSC2pe',
0.4076086956521739), ('AATSC2are', 0.4076086956521739), ('AATSC2p',
0.4076086956521739), ('AATSC2i', 0.4076086956521739), ('MATS2dv',
0.4076086956521739), ('MATS2Z', 0.4076086956521739), ('MATS2m',
0.4076086956521739), ('MATS2v', 0.4076086956521739), ('MATS2se',
0.4076086956521739), ('MATS2pe', 0.4076086956521739), ('MATS2are',
0.4076086956521739), ('MATS2p', 0.4076086956521739), ('MATS2i',
0.4076086956521739), ('GATS2dv', 0.4076086956521739), ('GATS2Z',
0.4076086956521739), ('GATS2m', 0.4076086956521739), ('GATS2v',

0.4076086956521739), ('GATS2se', 0.4076086956521739), ('GATS2pe', 0.4076086956521739), ('GATS2are', 0.4076086956521739), ('GATS2p', 0.4076086956521739), ('GATS2i', 0.4076086956521739), ('AATS1dv', 0.3736413043478261), ('AATS1d', 0.3736413043478261), ('AATS1Z', 0.3736413043478261), ('AATS1m', 0.3736413043478261), ('AATS1v', 0.3736413043478261), ('AATS1se', 0.3736413043478261), ('AATS1pe', 0.3736413043478261), ('AATS1are', 0.3736413043478261), ('AATS1p', 0.3736413043478261), ('AATS1i', 0.3736413043478261), ('AATSC1dv', 0.3736413043478261), ('AATSC1d', 0.3736413043478261), ('AATSC1Z', 0.3736413043478261), ('AATSC1m', 0.3736413043478261), ('AATSC1v', 0.3736413043478261), ('AATSC1se', 0.3736413043478261), ('AATSC1pe', 0.3736413043478261), ('AATSC1are', 0.3736413043478261), ('AATSC1p', 0.3736413043478261), ('AATSC1i', 0.3736413043478261), ('MATS1dv', 0.3736413043478261), ('MATS1Z', 0.3736413043478261), ('MATS1m', 0.3736413043478261), ('MATS1v', 0.3736413043478261), ('MATS1se', 0.3736413043478261), ('MATS1pe', 0.3736413043478261), ('MATS1are', 0.3736413043478261), ('MATS1p', 0.3736413043478261), ('MATS1i', 0.3736413043478261), ('GATS1dv', 0.3736413043478261), ('GATS1Z', 0.3736413043478261), ('GATS1m', 0.3736413043478261), ('GATS1v', 0.3736413043478261), ('GATS1se', 0.3736413043478261), ('GATS1pe', 0.3736413043478261), ('GATS1are', 0.3736413043478261), ('GATS1p', 0.3736413043478261), ('GATS1i', 0.3736413043478261), ('BIC0', 0.3736413043478261), ('BIC1', 0.3736413043478261), ('BIC2', 0.3736413043478261), ('BIC3', 0.3736413043478261), ('BIC4', 0.3736413043478261), ('BIC5', 0.3736413043478261), ('ATSC0c', 0.06793478260869565), ('ATSC1c', 0.06793478260869565), ('ATSC2c', 0.06793478260869565), ('ATSC3c', 0.06793478260869565), ('ATSC4c', 0.06793478260869565), ('ATSC5c', 0.06793478260869565), ('ATSC6c', 0.06793478260869565), ('ATSC7c', 0.06793478260869565), ('ATSC8c', 0.06793478260869565), ('AATSC0c', 0.06793478260869565), ('RNCG', 0.06793478260869565), ('RPCG', 0.06793478260869565), ('nAcid', 0.0), ('nBase', 0.0), ('nAromAtom', 0.0), ('nAromBond', 0.0), ('nAtom', 0.0), ('nHeavyAtom', 0.0), ('nSpiro', 0.0), ('nBridgehead', 0.0), ('nHetero', 0.0), ('nH', 0.0), ('nB', 0.0), ('nC', 0.0), ('nN', 0.0), ('nO', 0.0), ('nS', 0.0), ('nP', 0.0), ('nF', 0.0), ('nCl', 0.0), ('nBr', 0.0), ('nI', 0.0), ('nX', 0.0), ('ATS0dv', 0.0), ('ATS1dv', 0.0), ('ATS2dv', 0.0), ('ATS3dv', 0.0), ('ATS4dv', 0.0), ('ATS5dv', 0.0), ('ATS6dv', 0.0), ('ATS7dv', 0.0), ('ATS8dv', 0.0), ('ATS0d', 0.0), ('ATS1d', 0.0), ('ATS2d', 0.0), ('ATS3d', 0.0), ('ATS4d', 0.0), ('ATS5d', 0.0), ('ATS6d', 0.0), ('ATS7d', 0.0), ('ATS8d', 0.0), ('ATS0Z', 0.0), ('ATS1Z', 0.0), ('ATS2Z', 0.0), ('ATS3Z', 0.0), ('ATS4Z', 0.0), ('ATS5Z', 0.0), ('ATS6Z', 0.0), ('ATS7Z', 0.0), ('ATS8Z', 0.0), ('ATS0m', 0.0), ('ATS1m', 0.0), ('ATS2m', 0.0), ('ATS3m', 0.0), ('ATS4m', 0.0), ('ATS5m', 0.0), ('ATS6m', 0.0), ('ATS7m', 0.0), ('ATS8m', 0.0), ('ATS0v', 0.0), ('ATS1v', 0.0), ('ATS2v', 0.0), ('ATS3v', 0.0), ('ATS4v', 0.0), ('ATS5v', 0.0), ('ATS6v', 0.0), ('ATS7v', 0.0), ('ATS8v', 0.0), ('ATS0se', 0.0), ('ATS1se', 0.0), ('ATS2se', 0.0), ('ATS3se', 0.0), ('ATS4se', 0.0), ('ATS5se', 0.0), ('ATS6se', 0.0), ('ATS7se', 0.0), ('ATS8se', 0.0), ('ATS0pe', 0.0), ('ATS1pe', 0.0), ('ATS2pe', 0.0), ('ATS3pe', 0.0), ('ATS4pe', 0.0), ('ATS5pe', 0.0), ('ATS6pe', 0.0), ('ATS7pe', 0.0), ('ATS8pe', 0.0), ('ATS0are', 0.0),

('ATS1are', 0.0), ('ATS2are', 0.0), ('ATS3are', 0.0), ('ATS4are', 0.0), ('ATS5are', 0.0), ('ATS6are', 0.0), ('ATS7are', 0.0), ('ATS8are', 0.0), ('ATS0p', 0.0), ('ATS1p', 0.0), ('ATS2p', 0.0), ('ATS3p', 0.0), ('ATS4p', 0.0), ('ATS5p', 0.0), ('ATS6p', 0.0), ('ATS7p', 0.0), ('ATS8p', 0.0), ('ATS0i', 0.0), ('ATS1i', 0.0), ('ATS2i', 0.0), ('ATS3i', 0.0), ('ATS4i', 0.0), ('ATS5i', 0.0), ('ATS6i', 0.0), ('ATS7i', 0.0), ('ATS8i', 0.0), ('AATS0dv', 0.0), ('AATS0d', 0.0), ('AATS0Z', 0.0), ('AATS0m', 0.0), ('AATS0v', 0.0), ('AATS0se', 0.0), ('AATS0pe', 0.0), ('AATS0are', 0.0), ('AATS0p', 0.0), ('AATS0i', 0.0), ('ATSC0dv', 0.0), ('ATSC1dv', 0.0), ('ATSC2dv', 0.0), ('ATSC3dv', 0.0), ('ATSC4dv', 0.0), ('ATSC5dv', 0.0), ('ATSC6dv', 0.0), ('ATSC7dv', 0.0), ('ATSC8dv', 0.0), ('ATSC0d', 0.0), ('ATSC1d', 0.0), ('ATSC2d', 0.0), ('ATSC3d', 0.0), ('ATSC4d', 0.0), ('ATSC5d', 0.0), ('ATSC6d', 0.0), ('ATSC7d', 0.0), ('ATSC8d', 0.0), ('ATSC0Z', 0.0), ('ATSC1Z', 0.0), ('ATSC2Z', 0.0), ('ATSC3Z', 0.0), ('ATSC4Z', 0.0), ('ATSC5Z', 0.0), ('ATSC6Z', 0.0), ('ATSC7Z', 0.0), ('ATSC8Z', 0.0), ('ATSC0m', 0.0), ('ATSC1m', 0.0), ('ATSC2m', 0.0), ('ATSC3m', 0.0), ('ATSC4m', 0.0), ('ATSC5m', 0.0), ('ATSC6m', 0.0), ('ATSC7m', 0.0), ('ATSC8m', 0.0), ('ATSC0v', 0.0), ('ATSC1v', 0.0), ('ATSC2v', 0.0), ('ATSC3v', 0.0), ('ATSC4v', 0.0), ('ATSC5v', 0.0), ('ATSC6v', 0.0), ('ATSC7v', 0.0), ('ATSC8v', 0.0), ('ATSC0se', 0.0), ('ATSC1se', 0.0), ('ATSC2se', 0.0), ('ATSC3se', 0.0), ('ATSC4se', 0.0), ('ATSC5se', 0.0), ('ATSC6se', 0.0), ('ATSC7se', 0.0), ('ATSC8se', 0.0), ('ATSC0pe', 0.0), ('ATSC1pe', 0.0), ('ATSC2pe', 0.0), ('ATSC3pe', 0.0), ('ATSC4pe', 0.0), ('ATSC5pe', 0.0), ('ATSC6pe', 0.0), ('ATSC7pe', 0.0), ('ATSC8pe', 0.0), ('ATSC0are', 0.0), ('ATSC1are', 0.0), ('ATSC2are', 0.0), ('ATSC3are', 0.0), ('ATSC4are', 0.0), ('ATSC5are', 0.0), ('ATSC6are', 0.0), ('ATSC7are', 0.0), ('ATSC8are', 0.0), ('ATSC0p', 0.0), ('ATSC1p', 0.0), ('ATSC2p', 0.0), ('ATSC3p', 0.0), ('ATSC4p', 0.0), ('ATSC5p', 0.0), ('ATSC6p', 0.0), ('ATSC7p', 0.0), ('ATSC8p', 0.0), ('ATSC0i', 0.0), ('ATSC1i', 0.0), ('ATSC2i', 0.0), ('ATSC3i', 0.0), ('ATSC4i', 0.0), ('ATSC5i', 0.0), ('ATSC6i', 0.0), ('ATSC7i', 0.0), ('ATSC8i', 0.0), ('AATSC0dv', 0.0), ('AATSC0d', 0.0), ('AATSC0Z', 0.0), ('AATSC0m', 0.0), ('AATSC0v', 0.0), ('AATSC0se', 0.0), ('AATSC0pe', 0.0), ('AATSC0are', 0.0), ('AATSC0p', 0.0), ('AATSC0i', 0.0), ('BalabanJ', 0.0), ('BertzCT', 0.0), ('nBonds', 0.0), ('nBondsO', 0.0), ('nBondsS', 0.0), ('nBondsD', 0.0), ('nBondsT', 0.0), ('nBondsA', 0.0), ('nBondsM', 0.0), ('nBondsKS', 0.0), ('nBondsKD', 0.0), ('C1SP1', 0.0), ('C2SP1', 0.0), ('C1SP2', 0.0), ('C2SP2', 0.0), ('C3SP2', 0.0), ('C1SP3', 0.0), ('C2SP3', 0.0), ('C3SP3', 0.0), ('C4SP3', 0.0), ('FCSP3', 0.0), ('Xch-3d', 0.0), ('Xch-4d', 0.0), ('Xch-5d', 0.0), ('Xch-6d', 0.0), ('Xch-7d', 0.0), ('Xch-3dv', 0.0), ('Xch-4dv', 0.0), ('Xch-5dv', 0.0), ('Xch-6dv', 0.0), ('Xch-7dv', 0.0), ('Xc-3d', 0.0), ('Xc-4d', 0.0), ('Xc-5d', 0.0), ('Xc-6d', 0.0), ('Xc-3dv', 0.0), ('Xc-4dv', 0.0), ('Xc-5dv', 0.0), ('Xc-6dv', 0.0), ('Xpc-4d', 0.0), ('Xpc-5d', 0.0), ('Xpc-6d', 0.0), ('Xpc-4dv', 0.0), ('Xpc-5dv', 0.0), ('Xpc-6dv', 0.0), ('Xp-1d', 0.0), ('Xp-2d', 0.0), ('Xp-3d', 0.0), ('Xp-4d', 0.0), ('Xp-5d', 0.0), ('Xp-6d', 0.0), ('Xp-7d', 0.0), ('Xp-1dv', 0.0), ('Xp-2dv', 0.0), ('Xp-3dv', 0.0), ('Xp-4dv', 0.0), ('Xp-5dv', 0.0), ('Xp-6dv', 0.0), ('Xp-7dv', 0.0), ('SZ', 0.0), ('Sm', 0.0), ('Sv', 0.0), ('Sse', 0.0), ('Spe', 0.0), ('Sare', 0.0), ('Sp', 0.0), ('Si', 0.0), ('MZ', 0.0), ('Mm', 0.0), ('Mv', 0.0), ('Mse', 0.0), ('Mpe', 0.0), ('Mare', 0.0), ('Mp', 0.0), ('Mi', 0.0), ('ECIndex', 0.0), ('fragCpx', 0.0),

('fMF', 0.0), ('nHBAcc', 0.0), ('nHBDon', 0.0), ('IC0', 0.0), ('IC1', 0.0),
('IC2', 0.0), ('IC3', 0.0), ('IC4', 0.0), ('IC5', 0.0), ('TIC0', 0.0), ('TIC1',
0.0), ('TIC2', 0.0), ('TIC3', 0.0), ('TIC4', 0.0), ('TIC5', 0.0), ('SIC0', 0.0),
('SIC1', 0.0), ('SIC2', 0.0), ('SIC3', 0.0), ('SIC4', 0.0), ('SIC5', 0.0),
('CIC0', 0.0), ('CIC1', 0.0), ('CIC2', 0.0), ('CIC3', 0.0), ('CIC4', 0.0),
('CIC5', 0.0), ('MIC0', 0.0), ('MIC1', 0.0), ('MIC2', 0.0), ('MIC3', 0.0),
('MIC4', 0.0), ('MIC5', 0.0), ('ZMIC0', 0.0), ('ZMIC1', 0.0), ('ZMIC2', 0.0),
('ZMIC3', 0.0), ('ZMIC4', 0.0), ('ZMIC5', 0.0), ('Lipinski', 0.0),
('GhoseFilter', 0.0), ('FilterItLogS', 0.0), ('VMcGowan', 0.0), ('LabuteASA',
0.0), ('PEOE_VSA1', 0.0), ('PEOE_VSA2', 0.0), ('PEOE_VSA3', 0.0), ('PEOE_VSA4',
0.0), ('PEOE_VSA5', 0.0), ('PEOE_VSA6', 0.0), ('PEOE_VSA7', 0.0), ('PEOE_VSA8',
0.0), ('PEOE_VSA9', 0.0), ('PEOE_VSA10', 0.0), ('PEOE_VSA11', 0.0),
('PEOE_VSA12', 0.0), ('PEOE_VSA13', 0.0), ('SMR_VSA1', 0.0), ('SMR_VSA2', 0.0),
('SMR_VSA3', 0.0), ('SMR_VSA4', 0.0), ('SMR_VSA5', 0.0), ('SMR_VSA6', 0.0),
('SMR_VSA7', 0.0), ('SMR_VSA8', 0.0), ('SMR_VSA9', 0.0), ('SlogP_VSA1', 0.0),
('SlogP_VSA2', 0.0), ('SlogP_VSA3', 0.0), ('SlogP_VSA4', 0.0), ('SlogP_VSA5',
0.0), ('SlogP_VSA6', 0.0), ('SlogP_VSA7', 0.0), ('SlogP_VSA8', 0.0),
('SlogP_VSA9', 0.0), ('SlogP_VSA10', 0.0), ('SlogP_VSA11', 0.0), ('MPC2', 0.0),
('MPC3', 0.0), ('MPC4', 0.0), ('MPC5', 0.0), ('MPC6', 0.0), ('MPC7', 0.0),
('MPC8', 0.0), ('MPC9', 0.0), ('MPC10', 0.0), ('TMPC10', 0.0), ('piPC1', 0.0),
('piPC2', 0.0), ('piPC3', 0.0), ('piPC4', 0.0), ('piPC5', 0.0), ('piPC6', 0.0),
('piPC7', 0.0), ('piPC8', 0.0), ('piPC9', 0.0), ('piPC10', 0.0), ('TpiPC10',
0.0), ('apol', 0.0), ('bpol', 0.0), ('nRing', 0.0), ('n3Ring', 0.0), ('n4Ring',
0.0), ('n5Ring', 0.0), ('n6Ring', 0.0), ('n7Ring', 0.0), ('n8Ring', 0.0),
('n9Ring', 0.0), ('n10Ring', 0.0), ('n11Ring', 0.0), ('n12Ring', 0.0),
('nG12Ring', 0.0), ('nHRing', 0.0), ('n3HRing', 0.0), ('n4HRing', 0.0),
('n5HRing', 0.0), ('n6HRing', 0.0), ('n7HRing', 0.0), ('n8HRing', 0.0),
('n9HRing', 0.0), ('n10HRing', 0.0), ('n11HRing', 0.0), ('n12HRing', 0.0),
('nG12HRing', 0.0), ('naRing', 0.0), ('n3aRing', 0.0), ('n4aRing', 0.0),
('n5aRing', 0.0), ('n6aRing', 0.0), ('n7aRing', 0.0), ('n8aRing', 0.0),
('n9aRing', 0.0), ('n10aRing', 0.0), ('n11aRing', 0.0), ('n12aRing', 0.0),
('nG12aRing', 0.0), ('naHRing', 0.0), ('n3aHRing', 0.0), ('n4aHRing', 0.0),
('n5aHRing', 0.0), ('n6aHRing', 0.0), ('n7aHRing', 0.0), ('n8aHRing', 0.0),
('n9aHRing', 0.0), ('n10aHRing', 0.0), ('n11aHRing', 0.0), ('n12aHRing', 0.0),
('nG12aHRing', 0.0), ('nARing', 0.0), ('n3ARing', 0.0), ('n4ARing', 0.0),
('n5ARing', 0.0), ('n6ARing', 0.0), ('n7ARing', 0.0), ('n8ARing', 0.0),
('n9ARing', 0.0), ('n10ARing', 0.0), ('n11ARing', 0.0), ('n12ARing', 0.0),
('nG12ARing', 0.0), ('nAHRing', 0.0), ('n3AHRing', 0.0), ('n4AHRing', 0.0),
('n5AHRing', 0.0), ('n6AHRing', 0.0), ('n7AHRing', 0.0), ('n8AHRing', 0.0),
('n9AHRing', 0.0), ('n10AHRing', 0.0), ('n11AHRing', 0.0), ('n12AHRing', 0.0),
('nG12AHRing', 0.0), ('nFRing', 0.0), ('n4FRing', 0.0), ('n5FRing', 0.0),
('n6FRing', 0.0), ('n7FRing', 0.0), ('n8FRing', 0.0), ('n9FRing', 0.0),
('n10FRing', 0.0), ('n11FRing', 0.0), ('n12FRing', 0.0), ('nG12FRing', 0.0),
('nFHRing', 0.0), ('n4FHRing', 0.0), ('n5FHRing', 0.0), ('n6FHRing', 0.0),
('n7FHRing', 0.0), ('n8FHRing', 0.0), ('n9FHRing', 0.0), ('n10FHRing', 0.0),
('n11FHRing', 0.0), ('n12FHRing', 0.0), ('nG12FHRing', 0.0), ('nFaRing', 0.0),
('n4FaRing', 0.0), ('n5FaRing', 0.0), ('n6FaRing', 0.0), ('n7FaRing', 0.0),
('n8FaRing', 0.0), ('n9FaRing', 0.0), ('n10FaRing', 0.0), ('n11FaRing', 0.0),

```
('n12FaRing', 0.0), ('nG12FaRing', 0.0), ('nFaHRing', 0.0), ('n4FaHRing', 0.0),
('n5FaHRing', 0.0), ('n6FaHRing', 0.0), ('n7FaHRing', 0.0), ('n8FaHRing', 0.0),
('n9FaHRing', 0.0), ('n10FaHRing', 0.0), ('n11FaHRing', 0.0), ('n12FaHRing',
0.0), ('nG12FaHRing', 0.0), ('nFARing', 0.0), ('n4FARing', 0.0), ('n5FARing',
0.0), ('n6FARing', 0.0), ('n7FARing', 0.0), ('n8FARing', 0.0), ('n9FARing',
0.0), ('n10FARing', 0.0), ('n11FARing', 0.0), ('n12FARing', 0.0), ('nG12FARing',
0.0), ('nFAHRing', 0.0), ('n4FAHRing', 0.0), ('n5FAHRing', 0.0), ('n6FAHRing',
0.0), ('n7FAHRing', 0.0), ('n8FAHRing', 0.0), ('n9FAHRing', 0.0), ('n10FAHRing',
0.0), ('n11FAHRing', 0.0), ('n12FAHRing', 0.0), ('nG12FAHRing', 0.0), ('nRot',
0.0), ('SLogP', 0.0), ('SMR', 0.0), ('TopoPSA(NO)', 0.0), ('TopoPSA', 0.0),
('GGI1', 0.0), ('GGI2', 0.0), ('GGI3', 0.0), ('GGI4', 0.0), ('GGI5', 0.0),
('GGI6', 0.0), ('GGI7', 0.0), ('GGI8', 0.0), ('GGI9', 0.0), ('GGI10', 0.0),
('JGI1', 0.0), ('JGI2', 0.0), ('JGI3', 0.0), ('JGI4', 0.0), ('JGI5', 0.0),
('JGI6', 0.0), ('JGI7', 0.0), ('JGI8', 0.0), ('JGI9', 0.0), ('JGI10', 0.0),
('JGT10', 0.0), ('Diameter', 0.0), ('Radius', 0.0), ('TopoShapeIndex', 0.0),
('PetitjeanIndex', 0.0), ('MWC01', 0.0), ('MWC02', 0.0), ('MWC03', 0.0),
('MWC04', 0.0), ('MWC05', 0.0), ('MWC06', 0.0), ('MWC07', 0.0), ('MWC08', 0.0),
('MWC09', 0.0), ('MWC10', 0.0), ('TMWC10', 0.0), ('SRW02', 0.0), ('SRW03', 0.0),
('SRW04', 0.0), ('SRW05', 0.0), ('SRW06', 0.0), ('SRW07', 0.0), ('SRW08', 0.0),
('SRW09', 0.0), ('SRW10', 0.0), ('TSRW10', 0.0), ('MW', 0.0), ('AMW', 0.0),
('WPath', 0.0), ('WPol', 0.0), ('Zagreb1', 0.0), ('Zagreb2', 0.0), ('mZagreb2',
0.0)]
```

[ ]:
```
drop_columns=[i[0] for i in x if i[1]>40]#We'll drop columns which have more␣
↪than 40 percent missing values
```

[ ]:
```
print(drop_columns,)
```

```
['MDEN-11', 'MDEN-13', 'MDEN-33', 'MDEN-23', 'MDEC-44', 'MDEN-12', 'MDEN-22',
'MDEC-14', 'MDEC-24', 'MDEC-34', 'MDEO-22', 'MDEC-11', 'MDEO-12']
```

[ ]:
```
x_mord.drop(labels=drop_columns,axis=1,inplace=True)
```

[ ]:
```
list_of_columns=x_mord.columns
```

[ ]:
```
x_mord.info()
```

```
<class 'pandas.core.frame.DataFrame'>
RangeIndex: 2944 entries, 0 to 2943
Columns: 1249 entries, nAcid to MDEO-11
dtypes: float64(1249)
memory usage: 28.1 MB
```

[ ]:
```
from sklearn.preprocessing import StandardScaler
from sklearn.impute import KNNImputer
from sklearn.decomposition import PCA

def pre_process(x_train,x_test):
```

```
  imp=KNNImputer(missing_values=np.nan)
  x_train=imp.fit_transform(x_train)
  x_test=imp.transform(x_test)

  '''scale=StandardScaler()
  x_train=scale.fit_transform(x_train)
  x_test=scale.transform(x_test)

  pca=PCA(n_components=0.9999)
  x_train=pca.fit_transform(x_train)
  x_test=pca.transform(x_test)
  print("No of components", pca.n_components_)'''
  return x_train,x_test
```

[ ]:
```
#We choose Random forest as our model.The choice for model is based on similar␣
↪challanges in cheminformatics
#where the convention is to use Random forests for predicting molecular␣
↪property.They are also very powerful
#in capturing non-linear relationships and robust to outliers.

from sklearn.ensemble import RandomForestClassifier
from sklearn.model_selection import cross_val_predict
from sklearn.model_selection import cross_validate
import statistics
from sklearn.metrics import␣
↪f1_score,precision_score,recall_score,jaccard_score,multilabel_confusion_matrix
#Utility function to get model performance
def eval_train(model,x,y):

  #splitting the data
  x_train,y_train,x_test,y_test=iterative_split(x,y,test_size=0.2)

  x_train,x_test=pre_process(x_train,x_test)

  #Fitting our model
  model.fit(x_train,y_train)

  #Getting predictions on training set
 ␣
↪#y_train_prediction=cross_val_predict(model,x_train,y_train,cv=MultilabelStratifiedKFold(n_

  #Building a confusion matrix
  #mcf_matrix=multilabel_confusion_matrix(y_train,y_train_prediction)
  #print(mcf_matrix)
  #Evaluating model performance on training data
 ␣
↪train_score=cross_validate(model,x_train,y_train,scoring=["f1_micro","precision_micro","rec
```

```
  print(statistics.mean(train_score["test_f1_micro"]))
  print(statistics.mean(train_score["test_precision_micro"]))
  print(statistics.mean(train_score["test_recall_micro"]))

  #Evaluating on test data
  y_pred=model.predict(x_test)
  print("f1_score {}".format(f1_score(y_test,y_pred,average="micro")))
  print("precision_score {}".
 ↪format(precision_score(y_test,y_pred,average="micro")))
  print("recall_score {}".format(recall_score(y_test,y_pred,average="micro")))
  print("jaccard_score {}".format(jaccard_score(y_test,y_pred,average="micro")))
```

```
[ ]: from sklearn.ensemble import RandomForestClassifier
     def get_rf():
       rf= RandomForestClassifier(random_state=42,class_weight="balanced",n_jobs=-1)
       return rf
```

```
[ ]: from skmultilearn.problem_transform import BinaryRelevance
     from sklearn.ensemble import RandomForestClassifier
     def get_bin_rel():
       classifier = BinaryRelevance(
           classifier =␣
      ↪RandomForestClassifier(random_state=42,class_weight="balanced",n_jobs=-1))
       return classifier
```

```
[ ]: from skmultilearn.problem_transform import ClassifierChain
     from sklearn.ensemble import RandomForestClassifier

     def get_cls_chain():
         classifier = ClassifierChain(
             RandomForestClassifier(random_state=42, n_jobs=-1,␣
      ↪class_weight="balanced"),
             # require_dense=[False, True]
         )
         return classifier
```

```
[ ]: from sklearn.ensemble import RandomForestClassifier
     from sklearn.model_selection import cross_val_predict
     from sklearn.model_selection import cross_validate
     from sklearn.metrics import␣
      ↪f1_score,precision_score,recall_score,jaccard_score,multilabel_confusion_matrix
     #Utility function to get model performance
     def cross_val(model,x,y):

       #Preprocessing
       imp=KNNImputer(missing_values=np.nan)
       x=imp.fit_transform(x)
```

```
    #Evaluating model performance on training data
␣
↪cross_score=cross_validate(model,x,y,scoring=["f1_micro","precision_micro","recall_micro","

    return cross_score
```

[ ]:
```
model=get_rf()
eval_train(model,x_mord,Y)
```

```
0.5313701368214249
0.7513871868082512
0.4129401334816375
f1_score 0.5774548863370049
precision_score 0.761433868974042
recall_score 0.4650811627029068
jaccard_score 0.4059308072487644
```

[ ]:
```
model=get_rf()
arr=cross_val(model,x_mord,Y)
print(arr)

print(f"f1_score {sum(arr['test_f1_micro'])/5}")
print(f"precision_score {sum(arr['test_precision_micro'])/5}")
print(f"recall_score {sum(arr['test_recall_micro'])/5}")
```

```
{'fit_time': array([30.17161131, 29.96637917, 31.06865907, 28.73971725,
28.75052238]), 'score_time': array([0.43619823, 0.34393239, 0.37017393,
0.35580873, 0.34336162]), 'test_f1_micro': array([0.70430622, 0.70949185,
0.68019324, 0.70381232, 0.6993144 ]), 'test_precision_micro': array([0.80701754,
0.81497797, 0.79101124, 0.82758621, 0.82448037]), 'test_recall_micro':
array([0.62478778, 0.62818336, 0.59661017, 0.6122449 , 0.60714286]),
'test_jaccard_micro': array([0.54357459, 0.54977712, 0.51537335, 0.54298643,
0.5376506 ])}
f1_score 0.6994236043214046
precision_score 0.8130146659589101
recall_score 0.6137938124229577
```

[ ]:
```
pip install -U yellowbrick
```

```
Requirement already satisfied: yellowbrick in /usr/local/lib/python3.10/dist-
packages (1.5)
Requirement already satisfied: matplotlib!=3.0.0,>=2.0.2 in
/usr/local/lib/python3.10/dist-packages (from yellowbrick) (3.8.0)
Requirement already satisfied: scipy>=1.0.0 in /usr/local/lib/python3.10/dist-
packages (from yellowbrick) (1.13.1)
Requirement already satisfied: scikit-learn>=1.0.0 in
/usr/local/lib/python3.10/dist-packages (from yellowbrick) (1.5.2)
```

```
Requirement already satisfied: numpy>=1.16.0 in /usr/local/lib/python3.10/dist-
packages (from yellowbrick) (1.26.4)
Requirement already satisfied: cycler>=0.10.0 in /usr/local/lib/python3.10/dist-
packages (from yellowbrick) (0.12.1)
Requirement already satisfied: contourpy>=1.0.1 in
/usr/local/lib/python3.10/dist-packages (from
matplotlib!=3.0.0,>=2.0.2->yellowbrick) (1.3.1)
Requirement already satisfied: fonttools>=4.22.0 in
/usr/local/lib/python3.10/dist-packages (from
matplotlib!=3.0.0,>=2.0.2->yellowbrick) (4.55.0)
Requirement already satisfied: kiwisolver>=1.0.1 in
/usr/local/lib/python3.10/dist-packages (from
matplotlib!=3.0.0,>=2.0.2->yellowbrick) (1.4.7)
Requirement already satisfied: packaging>=20.0 in
/usr/local/lib/python3.10/dist-packages (from
matplotlib!=3.0.0,>=2.0.2->yellowbrick) (24.2)
Requirement already satisfied: pillow>=6.2.0 in /usr/local/lib/python3.10/dist-
packages (from matplotlib!=3.0.0,>=2.0.2->yellowbrick) (11.0.0)
Requirement already satisfied: pyparsing>=2.3.1 in
/usr/local/lib/python3.10/dist-packages (from
matplotlib!=3.0.0,>=2.0.2->yellowbrick) (3.2.0)
Requirement already satisfied: python-dateutil>=2.7 in
/usr/local/lib/python3.10/dist-packages (from
matplotlib!=3.0.0,>=2.0.2->yellowbrick) (2.8.2)
Requirement already satisfied: joblib>=1.2.0 in /usr/local/lib/python3.10/dist-
packages (from scikit-learn>=1.0.0->yellowbrick) (1.4.2)
Requirement already satisfied: threadpoolctl>=3.1.0 in
/usr/local/lib/python3.10/dist-packages (from scikit-learn>=1.0.0->yellowbrick)
(3.5.0)
Requirement already satisfied: six>=1.5 in /usr/local/lib/python3.10/dist-
packages (from python-dateutil>=2.7->matplotlib!=3.0.0,>=2.0.2->yellowbrick)
(1.16.0)
```

[ ]:
```
import yellowbrick
from yellowbrick.model_selection import FeatureImportances
plt.rcParams['figure.figsize'] = (12,8)
plt.style.use("ggplot")

viz = FeatureImportances(model,topn=30,labels=list_of_columns)

viz.fit(x_train, y_train)
viz.show()
```

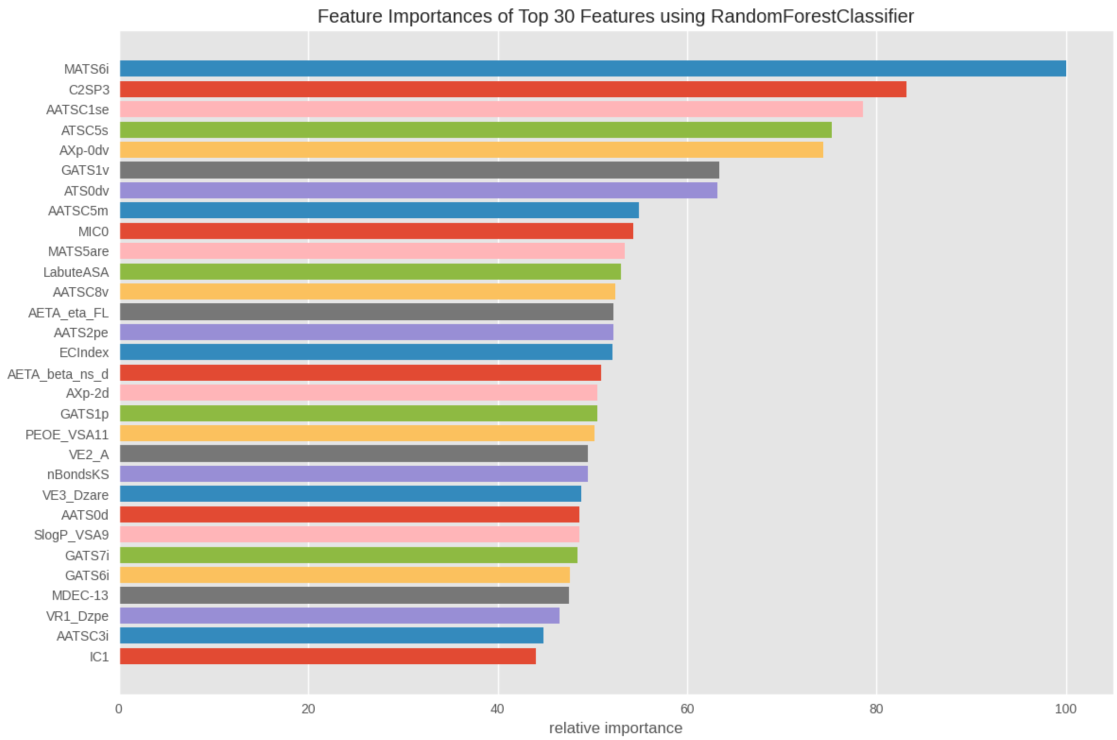


[ ]: <Axes: title={'center': 'Feature Importances of Top 30 Features using RandomForestClassifier'}, xlabel='relative importance'>

[ ]:
```
model=get_bin_rel()
eval_train(model,x_mord,Y)
```

```
/usr/local/lib/python3.10/dist-
packages/sklearn/model_selection/_validation.py:1000: UserWarning: Scoring
failed. The score on this train-test partition for these parameters will be set
to nan. Details:
Traceback (most recent call last):
  File "/usr/local/lib/python3.10/dist-packages/sklearn/metrics/_scorer.py",
line 139, in __call__
    score = scorer._score(
  File "/usr/local/lib/python3.10/dist-packages/sklearn/metrics/_scorer.py",
line 371, in _score
    y_pred = method_caller(
  File "/usr/local/lib/python3.10/dist-packages/sklearn/metrics/_scorer.py",
line 89, in _cached_call
    result, _ = _get_response_values(
  File "/usr/local/lib/python3.10/dist-packages/sklearn/utils/_response.py",
line 199, in _get_response_values
    classes = estimator.classes_
AttributeError: 'BinaryRelevance' object has no attribute 'classes_'
```

```
  warnings.warn(
/usr/local/lib/python3.10/dist-
packages/sklearn/model_selection/_validation.py:1000: UserWarning: Scoring
failed. The score on this train-test partition for these parameters will be set
to nan. Details:
Traceback (most recent call last):
  File "/usr/local/lib/python3.10/dist-packages/sklearn/metrics/_scorer.py",
line 139, in __call__
    score = scorer._score(
  File "/usr/local/lib/python3.10/dist-packages/sklearn/metrics/_scorer.py",
line 371, in _score
    y_pred = method_caller(
  File "/usr/local/lib/python3.10/dist-packages/sklearn/metrics/_scorer.py",
line 89, in _cached_call
    result, _ = _get_response_values(
  File "/usr/local/lib/python3.10/dist-packages/sklearn/utils/_response.py",
line 199, in _get_response_values
    classes = estimator.classes_
AttributeError: 'BinaryRelevance' object has no attribute 'classes_'

  warnings.warn(
/usr/local/lib/python3.10/dist-
packages/sklearn/model_selection/_validation.py:1000: UserWarning: Scoring
failed. The score on this train-test partition for these parameters will be set
to nan. Details:
Traceback (most recent call last):
  File "/usr/local/lib/python3.10/dist-packages/sklearn/metrics/_scorer.py",
line 139, in __call__
    score = scorer._score(
  File "/usr/local/lib/python3.10/dist-packages/sklearn/metrics/_scorer.py",
line 371, in _score
    y_pred = method_caller(
  File "/usr/local/lib/python3.10/dist-packages/sklearn/metrics/_scorer.py",
line 89, in _cached_call
    result, _ = _get_response_values(
  File "/usr/local/lib/python3.10/dist-packages/sklearn/utils/_response.py",
line 199, in _get_response_values
    classes = estimator.classes_
AttributeError: 'BinaryRelevance' object has no attribute 'classes_'

  warnings.warn(
/usr/local/lib/python3.10/dist-
packages/sklearn/model_selection/_validation.py:1000: UserWarning: Scoring
failed. The score on this train-test partition for these parameters will be set
to nan. Details:
Traceback (most recent call last):
  File "/usr/local/lib/python3.10/dist-packages/sklearn/metrics/_scorer.py",
```

```
line 139, in __call__
    score = scorer._score(
  File "/usr/local/lib/python3.10/dist-packages/sklearn/metrics/_scorer.py",
line 371, in _score
    y_pred = method_caller(
  File "/usr/local/lib/python3.10/dist-packages/sklearn/metrics/_scorer.py",
line 89, in _cached_call
    result, _ = _get_response_values(
  File "/usr/local/lib/python3.10/dist-packages/sklearn/utils/_response.py",
line 199, in _get_response_values
    classes = estimator.classes_
AttributeError: 'BinaryRelevance' object has no attribute 'classes_'

  warnings.warn(
/usr/local/lib/python3.10/dist-
packages/sklearn/model_selection/_validation.py:1000: UserWarning: Scoring
failed. The score on this train-test partition for these parameters will be set
to nan. Details:
Traceback (most recent call last):
  File "/usr/local/lib/python3.10/dist-packages/sklearn/metrics/_scorer.py",
line 139, in __call__
    score = scorer._score(
  File "/usr/local/lib/python3.10/dist-packages/sklearn/metrics/_scorer.py",
line 371, in _score
    y_pred = method_caller(
  File "/usr/local/lib/python3.10/dist-packages/sklearn/metrics/_scorer.py",
line 89, in _cached_call
    result, _ = _get_response_values(
  File "/usr/local/lib/python3.10/dist-packages/sklearn/utils/_response.py",
line 199, in _get_response_values
    classes = estimator.classes_
AttributeError: 'BinaryRelevance' object has no attribute 'classes_'

  warnings.warn(
```

```
nan
nan
nan
f1_score 0.6184239733629301
precision_score 0.7505387931034483
recall_score 0.5258588146470367
jaccard_score 0.4476221079691517
```

[ ]:
```
model=get_bin_rel()
arr=cross_val(model,x_mord,Y)
print(arr)

print(f"f1_score {sum(arr['test_f1_micro'])/5}")
```

```
print(f"precision_score {sum(arr['test_precision_micro'])/5}")
print(f"recall_score {sum(arr['test_recall_micro'])/5}")
```

```
{'fit_time': array([34.61546087, 33.78824902, 29.69570017, 30.00454283,
30.11443686]), 'score_time': array([0.00071812, 0.0011158 , 0.00068688,
0.00078344, 0.00072837]), 'test_f1_micro': array([nan, nan, nan, nan, nan]),
'test_precision_micro': array([nan, nan, nan, nan, nan]), 'test_recall_micro':
array([nan, nan, nan, nan, nan]), 'test_jaccard_micro': array([nan, nan, nan,
nan, nan])}
f1_score nan
precision_score nan
recall_score nan
```

[ ]:
```
print(f"x_mord shape: {x_mord.shape}")
print(f"Y shape: {Y.shape}")
```

```
x_mord shape: (2944, 1249)
Y shape: (2944, 44)
```

[ ]:
```
print(f"Length of vocabt: {len(vocabt)}")
```

```
Length of vocabt: 44
```

[ ]:
```
model=get_cls_chain()
eval_train(model,x_mord,Y)
```

```
/usr/local/lib/python3.10/dist-
packages/sklearn/model_selection/_validation.py:1000: UserWarning: Scoring
failed. The score on this train-test partition for these parameters will be set
to nan. Details:
Traceback (most recent call last):
  File "/usr/local/lib/python3.10/dist-packages/sklearn/metrics/_scorer.py",
line 139, in __call__
    score = scorer._score(
  File "/usr/local/lib/python3.10/dist-packages/sklearn/metrics/_scorer.py",
line 371, in _score
    y_pred = method_caller(
  File "/usr/local/lib/python3.10/dist-packages/sklearn/metrics/_scorer.py",
line 89, in _cached_call
    result, _ = _get_response_values(
  File "/usr/local/lib/python3.10/dist-packages/sklearn/utils/_response.py",
line 199, in _get_response_values
    classes = estimator.classes_
AttributeError: 'ClassifierChain' object has no attribute 'classes_'

  warnings.warn(
/usr/local/lib/python3.10/dist-
packages/sklearn/model_selection/_validation.py:1000: UserWarning: Scoring
```

```
failed. The score on this train-test partition for these parameters will be set
to nan. Details:
Traceback (most recent call last):
  File "/usr/local/lib/python3.10/dist-packages/sklearn/metrics/_scorer.py",
line 139, in __call__
    score = scorer._score(
  File "/usr/local/lib/python3.10/dist-packages/sklearn/metrics/_scorer.py",
line 371, in _score
    y_pred = method_caller(
  File "/usr/local/lib/python3.10/dist-packages/sklearn/metrics/_scorer.py",
line 89, in _cached_call
    result, _ = _get_response_values(
  File "/usr/local/lib/python3.10/dist-packages/sklearn/utils/_response.py",
line 199, in _get_response_values
    classes = estimator.classes_
AttributeError: 'ClassifierChain' object has no attribute 'classes_'

  warnings.warn(
/usr/local/lib/python3.10/dist-
packages/sklearn/model_selection/_validation.py:1000: UserWarning: Scoring
failed. The score on this train-test partition for these parameters will be set
to nan. Details:
Traceback (most recent call last):
  File "/usr/local/lib/python3.10/dist-packages/sklearn/metrics/_scorer.py",
line 139, in __call__
    score = scorer._score(
  File "/usr/local/lib/python3.10/dist-packages/sklearn/metrics/_scorer.py",
line 371, in _score
    y_pred = method_caller(
  File "/usr/local/lib/python3.10/dist-packages/sklearn/metrics/_scorer.py",
line 89, in _cached_call
    result, _ = _get_response_values(
  File "/usr/local/lib/python3.10/dist-packages/sklearn/utils/_response.py",
line 199, in _get_response_values
    classes = estimator.classes_
AttributeError: 'ClassifierChain' object has no attribute 'classes_'

  warnings.warn(
/usr/local/lib/python3.10/dist-
packages/sklearn/model_selection/_validation.py:1000: UserWarning: Scoring
failed. The score on this train-test partition for these parameters will be set
to nan. Details:
Traceback (most recent call last):
  File "/usr/local/lib/python3.10/dist-packages/sklearn/metrics/_scorer.py",
line 139, in __call__
    score = scorer._score(
  File "/usr/local/lib/python3.10/dist-packages/sklearn/metrics/_scorer.py",
line 371, in _score
```

```
    y_pred = method_caller(
  File "/usr/local/lib/python3.10/dist-packages/sklearn/metrics/_scorer.py",
line 89, in _cached_call
    result, _ = _get_response_values(
  File "/usr/local/lib/python3.10/dist-packages/sklearn/utils/_response.py",
line 199, in _get_response_values
    classes = estimator.classes_
AttributeError: 'ClassifierChain' object has no attribute 'classes_'

  warnings.warn(
/usr/local/lib/python3.10/dist-
packages/sklearn/model_selection/_validation.py:1000: UserWarning: Scoring
failed. The score on this train-test partition for these parameters will be set
to nan. Details:
Traceback (most recent call last):
  File "/usr/local/lib/python3.10/dist-packages/sklearn/metrics/_scorer.py",
line 139, in __call__
    score = scorer._score(
  File "/usr/local/lib/python3.10/dist-packages/sklearn/metrics/_scorer.py",
line 371, in _score
    y_pred = method_caller(
  File "/usr/local/lib/python3.10/dist-packages/sklearn/metrics/_scorer.py",
line 89, in _cached_call
    result, _ = _get_response_values(
  File "/usr/local/lib/python3.10/dist-packages/sklearn/utils/_response.py",
line 199, in _get_response_values
    classes = estimator.classes_
AttributeError: 'ClassifierChain' object has no attribute 'classes_'

  warnings.warn(
```

```
nan
nan
nan
f1_score 0.6153169014084507
precision_score 0.7377308707124011
recall_score 0.5277463193657984
jaccard_score 0.4443738080101716
```

[ ]:

```
model=get_cls_chain()
arr=cross_val(model,x_mord,Y)
print(arr)

print(f"f1_score {sum(arr['test_f1_micro'])/5}")
print(f"precision_score {sum(arr['test_precision_micro'])/5}")
print(f"recall_score {sum(arr['test_recall_micro'])/5}")
```

```
{'fit_time': array([28.04308105, 28.24094152, 31.43649507, 29.13527417,
```

```
29.29067779]), 'score_time': array([0.85899425, 0.81674933, 0.69341254,
0.70193982, 0.68509459]), 'test_f1_micro': array([0.69483568, 0.71916509,
0.71153846, 0.71132075, 0.72328244]), 'test_precision_micro': array([0.77731092,
0.81505376, 0.82039911, 0.80042463, 0.82391304]), 'test_recall_micro':
array([0.62818336, 0.6434635 , 0.62818336, 0.64006791, 0.64455782]),
'test_jaccard_micro': array([0.5323741 , 0.56148148, 0.55223881, 0.55197657,
0.56651719])}
f1_score 0.7120284850287405
precision_score 0.8074202945642031
recall_score 0.6368911911114191
```

[ ]:
```
from sklearn.multioutput import ClassifierChain
chains =␣
 ↪[ClassifierChain(RandomForestClassifier(class_weight="balanced",n_jobs=-1),␣
 ↪order='random', random_state=i)
          for i in range(10)]
x_train, y_train, x_test, y_test = iterative_split(x_mord, Y, test_size=0.2)

# print("x_train shape:", x_train.shape)
# print("y_train shape:", y_train.shape)
# print("x_test shape:", x_test.shape)
# print("y_test shape:", y_test.shape)


x_train,x_test=pre_process(x_train,x_test)
for chain in chains:
    chain.fit(x_train, y_train)

Y_pred_chains = np.array([chain.predict(x_test) for chain in
                          chains])
Y_pred_ensemble = (Y_pred_chains.mean(axis=0) > 0.5).astype(int)


# print("Y_pred_ensemble shape:", Y_pred_ensemble.shape)

ensemble_f1_score = f1_score(y_test, Y_pred_ensemble, average='micro')

ensemble_precision_score = precision_score(y_test, Y_pred_ensemble,␣
 ↪average='micro')
ensemble_recall_score = recall_score(y_test, Y_pred_ensemble, average='micro')
ensemble_jaccard_score = jaccard_score(y_test, Y_pred_ensemble, average='micro')

print("f1_score {}".format(ensemble_f1_score))
print("precision_score {}".format(ensemble_precision_score))
print("recall_score {}".format(ensemble_recall_score))
print("jaccard_score {}".format(ensemble_jaccard_score))
```

```
f1_score 0.6232785428698356
```

```
precision_score 0.7571505666486779
recall_score 0.5296338240845602
jaccard_score 0.4527266860277509
```

[ ]:
```
model=get_rf()
eval_train(model,x_morg,Y)
```

```
0.612131271969342
0.7571947927551376
0.5141342983390174
f1_score 0.6298245614035087
precision_score 0.7514390371533228
recall_score 0.542091355228388
jaccard_score 0.45966709346991036
```

[ ]:
```
model=get_rf()
arr=cross_val(model,x_morg,Y)
print(arr)

print(f"f1_score {sum(arr['test_f1_micro'])/5}")
print(f"precision_score {sum(arr['test_precision_micro'])/5}")
print(f"recall_score {sum(arr['test_recall_micro'])/5}")
```

```
{'fit_time': array([1.50530553, 1.51132512, 1.45346522, 2.02018237,
2.53782368]), 'score_time': array([0.08371329, 0.08627248, 0.0820899 , 0.1057179
, 0.08202124]), 'test_f1_micro': array([0.75070028, 0.70931326, 0.7247191 ,
0.71588785, 0.72676056]), 'test_precision_micro': array([0.8340249 , 0.79368421,
0.80793319, 0.7962578 , 0.81302521]), 'test_recall_micro': array([0.68251273,
0.64115646, 0.65704584, 0.65025467, 0.65704584]), 'test_jaccard_micro':
array([0.60089686, 0.54956268, 0.56828194, 0.55749636, 0.57079646])}
```

[ ]:
```
model=get_bin_rel()
eval_train(model,x_morg.astype(float),Y)
```

```
/usr/local/lib/python3.10/dist-
packages/sklearn/model_selection/_validation.py:1000: UserWarning: Scoring
failed. The score on this train-test partition for these parameters will be set
to nan. Details:
Traceback (most recent call last):
  File "/usr/local/lib/python3.10/dist-packages/sklearn/metrics/_scorer.py",
line 139, in __call__
    score = scorer._score(
  File "/usr/local/lib/python3.10/dist-packages/sklearn/metrics/_scorer.py",
line 371, in _score
    y_pred = method_caller(
  File "/usr/local/lib/python3.10/dist-packages/sklearn/metrics/_scorer.py",
line 89, in _cached_call
    result, _ = _get_response_values(
```

```
  File "/usr/local/lib/python3.10/dist-packages/sklearn/utils/_response.py",
line 199, in _get_response_values
    classes = estimator.classes_
AttributeError: 'BinaryRelevance' object has no attribute 'classes_'

  warnings.warn(
/usr/local/lib/python3.10/dist-
packages/sklearn/model_selection/_validation.py:1000: UserWarning: Scoring
failed. The score on this train-test partition for these parameters will be set
to nan. Details:
Traceback (most recent call last):
  File "/usr/local/lib/python3.10/dist-packages/sklearn/metrics/_scorer.py",
line 139, in __call__
    score = scorer._score(
  File "/usr/local/lib/python3.10/dist-packages/sklearn/metrics/_scorer.py",
line 371, in _score
    y_pred = method_caller(
  File "/usr/local/lib/python3.10/dist-packages/sklearn/metrics/_scorer.py",
line 89, in _cached_call
    result, _ = _get_response_values(
  File "/usr/local/lib/python3.10/dist-packages/sklearn/utils/_response.py",
line 199, in _get_response_values
    classes = estimator.classes_
AttributeError: 'BinaryRelevance' object has no attribute 'classes_'

  warnings.warn(
/usr/local/lib/python3.10/dist-
packages/sklearn/model_selection/_validation.py:1000: UserWarning: Scoring
failed. The score on this train-test partition for these parameters will be set
to nan. Details:
Traceback (most recent call last):
  File "/usr/local/lib/python3.10/dist-packages/sklearn/metrics/_scorer.py",
line 139, in __call__
    score = scorer._score(
  File "/usr/local/lib/python3.10/dist-packages/sklearn/metrics/_scorer.py",
line 371, in _score
    y_pred = method_caller(
  File "/usr/local/lib/python3.10/dist-packages/sklearn/metrics/_scorer.py",
line 89, in _cached_call
    result, _ = _get_response_values(
  File "/usr/local/lib/python3.10/dist-packages/sklearn/utils/_response.py",
line 199, in _get_response_values
    classes = estimator.classes_
AttributeError: 'BinaryRelevance' object has no attribute 'classes_'

  warnings.warn(
/usr/local/lib/python3.10/dist-
packages/sklearn/model_selection/_validation.py:1000: UserWarning: Scoring
```

```
failed. The score on this train-test partition for these parameters will be set
to nan. Details:
Traceback (most recent call last):
  File "/usr/local/lib/python3.10/dist-packages/sklearn/metrics/_scorer.py",
line 139, in __call__
    score = scorer._score(
  File "/usr/local/lib/python3.10/dist-packages/sklearn/metrics/_scorer.py",
line 371, in _score
    y_pred = method_caller(
  File "/usr/local/lib/python3.10/dist-packages/sklearn/metrics/_scorer.py",
line 89, in _cached_call
    result, _ = _get_response_values(
  File "/usr/local/lib/python3.10/dist-packages/sklearn/utils/_response.py",
line 199, in _get_response_values
    classes = estimator.classes_
AttributeError: 'BinaryRelevance' object has no attribute 'classes_'

  warnings.warn(
/usr/local/lib/python3.10/dist-
packages/sklearn/model_selection/_validation.py:1000: UserWarning: Scoring
failed. The score on this train-test partition for these parameters will be set
to nan. Details:
Traceback (most recent call last):
  File "/usr/local/lib/python3.10/dist-packages/sklearn/metrics/_scorer.py",
line 139, in __call__
    score = scorer._score(
  File "/usr/local/lib/python3.10/dist-packages/sklearn/metrics/_scorer.py",
line 371, in _score
    y_pred = method_caller(
  File "/usr/local/lib/python3.10/dist-packages/sklearn/metrics/_scorer.py",
line 89, in _cached_call
    result, _ = _get_response_values(
  File "/usr/local/lib/python3.10/dist-packages/sklearn/utils/_response.py",
line 199, in _get_response_values
    classes = estimator.classes_
AttributeError: 'BinaryRelevance' object has no attribute 'classes_'

  warnings.warn(
```

```
nan
nan
nan
f1_score 0.6334626994394136
precision_score 0.738562091503268
recall_score 0.5545488863722159
jaccard_score 0.4635531713474282
```

```
[ ]: model=get_bin_rel()
     arr=cross_val(model,x_morg.astype(float),Y)
     print(arr)

     print(f"f1_score {sum(arr['test_f1_micro'])/5}")
     print(f"precision_score {sum(arr['test_precision_micro'])/5}")
     print(f"recall_score {sum(arr['test_recall_micro'])/5}")
```

```
{'fit_time': array([6.2628088 , 7.97275376, 6.39946747, 7.99715376,
6.27549767]), 'score_time': array([0.00069427, 0.00065422, 0.00067377,
0.00067091, 0.00062108]), 'test_f1_micro': array([nan, nan, nan, nan, nan]),
'test_precision_micro': array([nan, nan, nan, nan, nan]), 'test_recall_micro':
array([nan, nan, nan, nan, nan]), 'test_jaccard_micro': array([nan, nan, nan,
nan, nan])}
f1_score nan
precision_score nan
recall_score nan
```

```
[ ]: model=get_cls_chain()
     eval_train(model,x_morg.astype(float),Y)
```

```
0.6313497081300928
0.740843137254902
0.5503959894572694
f1_score 0.6474820143884892
precision_score 0.7366393837265286
recall_score 0.5775764439411099
jaccard_score 0.4787234042553192
```

```
[ ]: model=get_cls_chain()
     arr=cross_val(model,x_morg.astype(float),Y)
     print(arr)

     print(f"f1_score {sum(arr['test_f1_micro'])/5}")
     print(f"precision_score {sum(arr['test_precision_micro'])/5}")
     print(f"recall_score {sum(arr['test_recall_micro'])/5}")
```

```
{'fit_time': array([7.62174249, 6.39267039, 7.68412423, 6.09819579, 6.9811542
]), 'score_time': array([0.68084145, 0.90557384, 0.69500971, 0.8486712 ,
0.71326947]), 'test_f1_micro': array([0.75795053, 0.71403197, 0.72924188,
0.73291925, 0.71721677]), 'test_precision_micro': array([0.79005525, 0.74860335,
0.77842004, 0.76623377, 0.7556391 ]), 'test_recall_micro': array([0.72835314,
0.68251273, 0.68590832, 0.70238095, 0.68251273]), 'test_jaccard_micro':
array([0.61024182, 0.55524862, 0.57386364, 0.57843137, 0.55910987])}
f1_score 0.7302720808542532
precision_score 0.7677903006175729
recall_score 0.6963335758751719
```

[ ]:
```
model=get_rf()
eval_train(model,x_path,Y)
```

```
0.5670709534153858
0.7592911879142976
0.4531785992539685
f1_score 0.5647716682199441
precision_score 0.7376749847839318
recall_score 0.4575311438278596
jaccard_score 0.3935064935064935
```

[ ]:
```
model=get_rf()
arr=cross_val(model,x_path,Y)
print(arr)

print(f"f1_score {sum(arr['test_f1_micro'])/5}")
print(f"precision_score {sum(arr['test_precision_micro'])/5}")
print(f"recall_score {sum(arr['test_recall_micro'])/5}")
```

```
{'fit_time': array([1.71445632, 1.70234466, 1.63118196, 2.996171  ,
1.88761044]), 'score_time': array([0.08056498, 0.07936549, 0.10582972, 0.1433413
, 0.08719301]), 'test_f1_micro': array([0.67621777, 0.70554493, 0.70943396,
0.68768473, 0.70781099]), 'test_precision_micro': array([0.77292576, 0.80743982,
0.79830149, 0.81924883, 0.81737194]), 'test_recall_micro': array([0.60101868,
0.62648557, 0.63837012, 0.59252971, 0.62414966]), 'test_jaccard_micro':
array([0.51082251, 0.5450517 , 0.5497076 , 0.52402402, 0.54776119])}
f1_score 0.6973384765398645
precision_score 0.8030575678534577
recall_score 0.6165107469133664
```

[ ]:
```
get_cls_chain()
```

[ ]:
```
ClassifierChain(base_estimator=RandomForestClassifier(class_weight='balanced',
                                                      n_jobs=-1,
                                                      random_state=42))
```

[ ]:
```
model=get_bin_rel()
eval_train(model,x_path,Y)
```

```
/usr/local/lib/python3.10/dist-
packages/sklearn/model_selection/_validation.py:1000: UserWarning: Scoring
failed. The score on this train-test partition for these parameters will be set
to nan. Details:
Traceback (most recent call last):
  File "/usr/local/lib/python3.10/dist-packages/sklearn/metrics/_scorer.py",
line 139, in __call__
    score = scorer._score(
  File "/usr/local/lib/python3.10/dist-packages/sklearn/metrics/_scorer.py",
```

```
line 371, in _score
    y_pred = method_caller(
  File "/usr/local/lib/python3.10/dist-packages/sklearn/metrics/_scorer.py",
line 89, in _cached_call
    result, _ = _get_response_values(
  File "/usr/local/lib/python3.10/dist-packages/sklearn/utils/_response.py",
line 199, in _get_response_values
    classes = estimator.classes_
AttributeError: 'BinaryRelevance' object has no attribute 'classes_'

  warnings.warn(
/usr/local/lib/python3.10/dist-
packages/sklearn/model_selection/_validation.py:1000: UserWarning: Scoring
failed. The score on this train-test partition for these parameters will be set
to nan. Details:
Traceback (most recent call last):
  File "/usr/local/lib/python3.10/dist-packages/sklearn/metrics/_scorer.py",
line 139, in __call__
    score = scorer._score(
  File "/usr/local/lib/python3.10/dist-packages/sklearn/metrics/_scorer.py",
line 371, in _score
    y_pred = method_caller(
  File "/usr/local/lib/python3.10/dist-packages/sklearn/metrics/_scorer.py",
line 89, in _cached_call
    result, _ = _get_response_values(
  File "/usr/local/lib/python3.10/dist-packages/sklearn/utils/_response.py",
line 199, in _get_response_values
    classes = estimator.classes_
AttributeError: 'BinaryRelevance' object has no attribute 'classes_'

  warnings.warn(
/usr/local/lib/python3.10/dist-
packages/sklearn/model_selection/_validation.py:1000: UserWarning: Scoring
failed. The score on this train-test partition for these parameters will be set
to nan. Details:
Traceback (most recent call last):
  File "/usr/local/lib/python3.10/dist-packages/sklearn/metrics/_scorer.py",
line 139, in __call__
    score = scorer._score(
  File "/usr/local/lib/python3.10/dist-packages/sklearn/metrics/_scorer.py",
line 371, in _score
    y_pred = method_caller(
  File "/usr/local/lib/python3.10/dist-packages/sklearn/metrics/_scorer.py",
line 89, in _cached_call
    result, _ = _get_response_values(
  File "/usr/local/lib/python3.10/dist-packages/sklearn/utils/_response.py",
line 199, in _get_response_values
    classes = estimator.classes_
```

```
AttributeError: 'BinaryRelevance' object has no attribute 'classes_'

  warnings.warn(
/usr/local/lib/python3.10/dist-
packages/sklearn/model_selection/_validation.py:1000: UserWarning: Scoring
failed. The score on this train-test partition for these parameters will be set
to nan. Details:
Traceback (most recent call last):
  File "/usr/local/lib/python3.10/dist-packages/sklearn/metrics/_scorer.py",
line 139, in __call__
    score = scorer._score(
  File "/usr/local/lib/python3.10/dist-packages/sklearn/metrics/_scorer.py",
line 371, in _score
    y_pred = method_caller(
  File "/usr/local/lib/python3.10/dist-packages/sklearn/metrics/_scorer.py",
line 89, in _cached_call
    result, _ = _get_response_values(
  File "/usr/local/lib/python3.10/dist-packages/sklearn/utils/_response.py",
line 199, in _get_response_values
    classes = estimator.classes_
AttributeError: 'BinaryRelevance' object has no attribute 'classes_'

  warnings.warn(
/usr/local/lib/python3.10/dist-
packages/sklearn/model_selection/_validation.py:1000: UserWarning: Scoring
failed. The score on this train-test partition for these parameters will be set
to nan. Details:
Traceback (most recent call last):
  File "/usr/local/lib/python3.10/dist-packages/sklearn/metrics/_scorer.py",
line 139, in __call__
    score = scorer._score(
  File "/usr/local/lib/python3.10/dist-packages/sklearn/metrics/_scorer.py",
line 371, in _score
    y_pred = method_caller(
  File "/usr/local/lib/python3.10/dist-packages/sklearn/metrics/_scorer.py",
line 89, in _cached_call
    result, _ = _get_response_values(
  File "/usr/local/lib/python3.10/dist-packages/sklearn/utils/_response.py",
line 199, in _get_response_values
    classes = estimator.classes_
AttributeError: 'BinaryRelevance' object has no attribute 'classes_'

  warnings.warn(

nan
nan
nan
f1_score 0.5944148936170213
```

```
precision_score 0.7198067632850241
recall_score 0.5062287655719139
jaccard_score 0.4228949858088931
```

[ ]:
```
model=get_bin_rel()
arr=cross_val(model,x_path,Y)
print(arr)

print(f"f1_score {sum(arr['test_f1_micro'])/5}")
print(f"precision_score {sum(arr['test_precision_micro'])/5}")
print(f"recall_score {sum(arr['test_recall_micro'])/5}")
```

```
{'fit_time': array([6.04961085, 7.62805319, 5.90413451, 7.47657442,
5.93639684]), 'score_time': array([0.00058913, 0.00062728, 0.00071645,
0.00061798, 0.00098372]), 'test_f1_micro': array([nan, nan, nan, nan, nan]),
'test_precision_micro': array([nan, nan, nan, nan, nan]), 'test_recall_micro':
array([nan, nan, nan, nan, nan]), 'test_jaccard_micro': array([nan, nan, nan,
nan, nan])}
f1_score nan
precision_score nan
recall_score nan
```

[ ]:
```
model=get_cls_chain()
eval_train(model,x_path,Y)
```

```
0.6010628833263475
0.7319958643280379
0.5106827727808885
f1_score 0.5941738937069158
precision_score 0.7229437229437229
recall_score 0.5043412608531521
jaccard_score 0.4226510597912053
```

[ ]:
```
model=get_cls_chain()
arr=cross_val(model,x_path,Y)
print(arr)

print(f"f1_score {sum(arr['test_f1_micro'])/5}")
print(f"precision_score {sum(arr['test_precision_micro'])/5}")
print(f"recall_score {sum(arr['test_recall_micro'])/5}")
```

```
{'fit_time': array([7.21085453, 5.7931354 , 7.29663515, 5.76852274,
7.42309117]), 'score_time': array([0.73366022, 0.71091199, 0.67717552,
0.67349124, 0.72294569]), 'test_f1_micro': array([0.72694064, 0.71912168,
0.6953271 , 0.70919325, 0.70082342]), 'test_precision_micro': array([0.78656126,
0.77821782, 0.77338877, 0.79245283, 0.75992063]), 'test_recall_micro':
array([0.67572156, 0.66836735, 0.63157895, 0.6417657 , 0.65025467]),
'test_jaccard_micro': array([0.57101865, 0.56142857, 0.53295129, 0.5494186 ,
```

```
0.53943662])}
f1_score 0.7102812186133521
precision_score 0.77810826502048
recall_score 0.6535376459582135
```

[ ]:
```
from sklearn.ensemble import RandomForestClassifier
from sklearn.model_selection import cross_val_predict
from sklearn.model_selection import cross_validate
from sklearn.metrics import␣
↪f1_score,precision_score,recall_score,jaccard_score,multilabel_confusion_matrix
#Utility function to get model performance
def eval_train(model,x,y):

  #splitting the data
  x_train,y_train,x_test,y_test=iterative_split(x,y,test_size=0.2)

  x_train,x_test=pre_process(x_train,x_test)

  #Fitting our model
  model.fit(x_train,y_train)

  #Evaluating on test data
  y_pred=model.predict(x_test)
  print("f1_score {}".format(f1_score(y_test,y_pred,average="micro")))
  print("precision_score {}".
↪format(precision_score(y_test,y_pred,average="micro")))
  print("recall_score {}".format(recall_score(y_test,y_pred,average="micro")))
  print("jaccard_score {}".format(jaccard_score(y_test,y_pred,average="micro")))

  return multilabel_confusion_matrix(y_test,y_pred)
```

[ ]:
```
model=get_rf()
cfmatrix = eval_train(model,x_morg,Y)
```

```
f1_score 0.6298245614035087
precision_score 0.7514390371533228
recall_score 0.542091355228388
jaccard_score 0.45966709346991036
```

[ ]:
```
cfmatrix.shape
```

[ ]:
```
(44, 2, 2)
```

[ ]:
```
cfmatrix
```

[ ]:
```
array([[[2358,    1],
        [   0,    0]],
```

```
[[2358,    0],
 [   1,    0]],

[[2349,    0],
 [   9,    1]],

[[2345,    0],
 [  14,    0]],

[[2358,    0],
 [   1,    0]],

[[1164,  248],
 [ 261,  686]],

[[2347,    2],
 [   7,    3]],

[[2349,    0],
 [  10,    0]],

[[2358,    0],
 [   1,    0]],

[[2352,    0],
 [   7,    0]],

[[2358,    0],
 [   1,    0]],

[[2354,    1],
 [   4,    0]],

[[2358,    0],
 [   1,    0]],

[[2349,    0],
 [   9,    1]],

[[2358,    0],
 [   1,    0]],

[[2358,    0],
 [   1,    0]],

[[2353,    0],
```

```
 [   5,    1]],

[[2353,    0],
 [   6,    0]],

[[2356,    0],
 [   3,    0]],

[[2358,    0],
 [   1,    0]],

[[2356,    0],
 [   2,    1]],

[[2358,    0],
 [   1,    0]],

[[2353,    3],
 [   3,    0]],

[[2351,    1],
 [   7,    0]],

[[2344,    1],
 [  12,    2]],

[[2356,    0],
 [   3,    0]],

[[2057,   47],
 [ 149,  106]],

[[2344,    1],
 [  13,    1]],

[[2337,    1],
 [  17,    4]],

[[2355,    4],
 [   0,    0]],

[[2296,    1],
 [  60,    2]],

[[2358,    0],
 [   1,    0]],
```

```
        [[2353,    0],
         [   6,    0]],

        [[2352,    0],
         [   6,    1]],

        [[2313,    0],
         [  42,    4]],

        [[2351,    2],
         [   6,    0]],

        [[2357,    0],
         [   2,    0]],

        [[1340,  140],
         [ 319,  560]],

        [[2356,    0],
         [   3,    0]],

        [[2175,   15],
         [ 158,   11]],

        [[2276,    1],
         [  30,   52]],

        [[2340,    4],
         [  15,    0]],

        [[2347,    2],
         [  10,    0]],

        [[2354,    0],
         [   5,    0]]])
```

[ ]:
```
df.shape
```

[ ]: (2944, 10)

[ ]:
```
label_per=pd.
 ↪DataFrame(index=tastes,columns=["Precision_score","Recall_score","F1_score","Percentage_of_
```

[ ]:
```
label_per
```

[ ]:
```
                   Precision_score Recall_score F1_score Percentage_of_samples
acid                           NaN          NaN      NaN                   NaN
```

```
acrid                    NaN        NaN      NaN                   NaN
and tingling             NaN        NaN      NaN                   NaN
astringent               NaN        NaN      NaN                   NaN
barely sweet             NaN        NaN      NaN                   NaN
bitter                   NaN        NaN      NaN                   NaN
burning                  NaN        NaN      NaN                   NaN
cooling                  NaN        NaN      NaN                   NaN
entirely bitter          NaN        NaN      NaN                   NaN
extremely bitter         NaN        NaN      NaN                   NaN
faint bitter             NaN        NaN      NaN                   NaN
faintly bitter           NaN        NaN      NaN                   NaN
feebly bitter            NaN        NaN      NaN                   NaN
heating                  NaN        NaN      NaN                   NaN
highly bitter            NaN        NaN      NaN                   NaN
highly sweet             NaN        NaN      NaN                   NaN
hot burning              NaN        NaN      NaN                   NaN
intensely bitter         NaN        NaN      NaN                   NaN
intensely sweet          NaN        NaN      NaN                   NaN
lacking sweet            NaN        NaN      NaN                   NaN
less sweet               NaN        NaN      NaN                   NaN
like fresh walnut        NaN        NaN      NaN                   NaN
low sweet                NaN        NaN      NaN                   NaN
moderately bitter        NaN        NaN      NaN                   NaN
neutral                  NaN        NaN      NaN                   NaN
non-bitter               NaN        NaN      NaN                   NaN
non-sweet                NaN        NaN      NaN                   NaN
pungent                  NaN        NaN      NaN                   NaN
salty                    NaN        NaN      NaN                   NaN
scratchy                 NaN        NaN      NaN                   NaN
slightly bitter          NaN        NaN      NaN                   NaN
slightly burning         NaN        NaN      NaN                   NaN
slightly sweet           NaN        NaN      NaN                   NaN
somewhat bitter          NaN        NaN      NaN                   NaN
sour                     NaN        NaN      NaN                   NaN
strongly bitter          NaN        NaN      NaN                   NaN
sulphurous               NaN        NaN      NaN                   NaN
sweet                    NaN        NaN      NaN                   NaN
sweetish                 NaN        NaN      NaN                   NaN
tasteless                NaN        NaN      NaN                   NaN
umami                    NaN        NaN      NaN                   NaN
very bitter              NaN        NaN      NaN                   NaN
very sweet               NaN        NaN      NaN                   NaN
weak umami               NaN        NaN      NaN                   NaN
```

[ ]:
```
label_per["Percentage_of_samples"]=[((x[1]/df.shape[0])*100) for x in␣
↪sorted(taste_count.items())]
```

[ ]:
```
f1=[]
pre=[]
rec=[]
for lab in cfmatrix:
  p=r=f=0
  if(lab[1][1]!=0):
    p=lab[1][1]/(lab[1][1]+lab[0][1])
    r=lab[1][1]/(lab[1][1]+lab[1][0])
    f=(p*r*2)/(p+r)
  f1.append(f)
  pre.append(p)
  rec.append(r)
```

[ ]:
```
label_per["Precision_score"]=pre
label_per["Recall_score"]=rec
label_per["F1_score"]=f1
```

[ ]:
```
label_per
```

[ ]:

| | Precision_score | Recall_score | F1_score | \ |
|---|---|---|---|---|
| acid | 0.000000 | 0.000000 | 0.000000 | |
| acrid | 0.000000 | 0.000000 | 0.000000 | |
| and tingling | 1.000000 | 0.100000 | 0.181818 | |
| astringent | 0.000000 | 0.000000 | 0.000000 | |
| barely sweet | 0.000000 | 0.000000 | 0.000000 | |
| bitter | 0.734475 | 0.724393 | 0.729399 | |
| burning | 0.600000 | 0.300000 | 0.400000 | |
| cooling | 0.000000 | 0.000000 | 0.000000 | |
| entirely bitter | 0.000000 | 0.000000 | 0.000000 | |
| extremely bitter | 0.000000 | 0.000000 | 0.000000 | |
| faint bitter | 0.000000 | 0.000000 | 0.000000 | |
| faintly bitter | 0.000000 | 0.000000 | 0.000000 | |
| feebly bitter | 0.000000 | 0.000000 | 0.000000 | |
| heating | 1.000000 | 0.100000 | 0.181818 | |
| highly bitter | 0.000000 | 0.000000 | 0.000000 | |
| highly sweet | 0.000000 | 0.000000 | 0.000000 | |
| hot burning | 1.000000 | 0.166667 | 0.285714 | |
| intensely bitter | 0.000000 | 0.000000 | 0.000000 | |
| intensely sweet | 0.000000 | 0.000000 | 0.000000 | |
| lacking sweet | 0.000000 | 0.000000 | 0.000000 | |
| less sweet | 1.000000 | 0.333333 | 0.500000 | |
| like fresh walnut | 0.000000 | 0.000000 | 0.000000 | |
| low sweet | 0.000000 | 0.000000 | 0.000000 | |
| moderately bitter | 0.000000 | 0.000000 | 0.000000 | |
| neutral | 0.666667 | 0.142857 | 0.235294 | |
| non-bitter | 0.000000 | 0.000000 | 0.000000 | |
| non-sweet | 0.692810 | 0.415686 | 0.519608 | |

| | | | |
|---|---|---|---|
| pungent | 0.500000 | 0.071429 | 0.125000 |
| salty | 0.800000 | 0.190476 | 0.307692 |
| scratchy | 0.000000 | 0.000000 | 0.000000 |
| slightly bitter | 0.666667 | 0.032258 | 0.061538 |
| slightly burning | 0.000000 | 0.000000 | 0.000000 |
| slightly sweet | 0.000000 | 0.000000 | 0.000000 |
| somewhat bitter | 1.000000 | 0.142857 | 0.250000 |
| sour | 1.000000 | 0.086957 | 0.160000 |
| strongly bitter | 0.000000 | 0.000000 | 0.000000 |
| sulphurous | 0.000000 | 0.000000 | 0.000000 |
| sweet | 0.800000 | 0.637088 | 0.709310 |
| sweetish | 0.000000 | 0.000000 | 0.000000 |
| tasteless | 0.423077 | 0.065089 | 0.112821 |
| umami | 0.981132 | 0.634146 | 0.770370 |
| very bitter | 0.000000 | 0.000000 | 0.000000 |
| very sweet | 0.000000 | 0.000000 | 0.000000 |
| weak umami | 0.000000 | 0.000000 | 0.000000 |

| | Percentage_of_samples |
|---|---|
| acid | 0.033967 |
| acrid | 0.033967 |
| and tingling | 0.407609 |
| astringent | 0.577446 |
| barely sweet | 0.033967 |
| bitter | 40.489130 |
| burning | 0.407609 |
| cooling | 0.407609 |
| entirely bitter | 0.033967 |
| extremely bitter | 0.305707 |
| faint bitter | 0.033967 |
| faintly bitter | 0.169837 |
| feebly bitter | 0.033967 |
| heating | 0.407609 |
| highly bitter | 0.033967 |
| highly sweet | 0.033967 |
| hot burning | 0.271739 |
| intensely bitter | 0.271739 |
| intensely sweet | 0.135870 |
| lacking sweet | 0.033967 |
| less sweet | 0.135870 |
| like fresh walnut | 0.033967 |
| low sweet | 0.135870 |
| moderately bitter | 0.305707 |
| neutral | 0.577446 |
| non-bitter | 0.135870 |
| non-sweet | 10.903533 |
| pungent | 0.577446 |

```
salty                          0.883152
scratchy                       0.033967
slightly bitter                2.615489
slightly burning               0.033967
slightly sweet                 0.237772
somewhat bitter                0.305707
sour                           1.936141
strongly bitter                0.271739
sulphurous                     0.067935
sweet                         37.941576
sweetish                       0.135870
tasteless                      7.167120
umami                          3.532609
very bitter                    0.645380
very sweet                     0.407609
weak umami                     0.203804
```

[ ]:
```
label_per.sort_values(by="Percentage_of_samples").head(15)
```

[ ]:
```
                   Precision_score  Recall_score  F1_score  \
acid                           0.0           0.0       0.0
slightly burning               0.0           0.0       0.0
scratchy                       0.0           0.0       0.0
lacking sweet                  0.0           0.0       0.0
highly sweet                   0.0           0.0       0.0
highly bitter                  0.0           0.0       0.0
feebly bitter                  0.0           0.0       0.0
faint bitter                   0.0           0.0       0.0
like fresh walnut              0.0           0.0       0.0
barely sweet                   0.0           0.0       0.0
acrid                          0.0           0.0       0.0
entirely bitter                0.0           0.0       0.0
sulphurous                     0.0           0.0       0.0
sweetish                       0.0           0.0       0.0
intensely sweet                0.0           0.0       0.0

                   Percentage_of_samples
acid                            0.033967
slightly burning                0.033967
scratchy                        0.033967
lacking sweet                   0.033967
highly sweet                    0.033967
highly bitter                   0.033967
feebly bitter                   0.033967
faint bitter                    0.033967
like fresh walnut               0.033967
barely sweet                    0.033967
```

```
acrid                              0.033967
entirely bitter                    0.033967
sulphurous                         0.067935
sweetish                           0.135870
intensely sweet                    0.135870
```

[ ]:
```
label_per["Percentage_of_samples"].max()
```

[ ]:
```
40.48913043478261
```

[ ]:
```
lb=label_per.drop(labels=label_per[label_per["F1_score"]==0].index,axis=0).
 ↪sort_values(by="F1_score")
```

[ ]:
```
lb[:15]
```

[ ]:
```
                 Precision_score  Recall_score  F1_score  \
slightly bitter         0.666667      0.032258  0.061538
tasteless               0.423077      0.065089  0.112821
pungent                 0.500000      0.071429  0.125000
sour                    1.000000      0.086957  0.160000
and tingling            1.000000      0.100000  0.181818
heating                 1.000000      0.100000  0.181818
neutral                 0.666667      0.142857  0.235294
somewhat bitter         1.000000      0.142857  0.250000
hot burning             1.000000      0.166667  0.285714
salty                   0.800000      0.190476  0.307692
burning                 0.600000      0.300000  0.400000
less sweet              1.000000      0.333333  0.500000
non-sweet               0.692810      0.415686  0.519608
sweet                   0.800000      0.637088  0.709310
bitter                  0.734475      0.724393  0.729399

                 Percentage_of_samples
slightly bitter               2.615489
tasteless                     7.167120
pungent                       0.577446
sour                          1.936141
and tingling                  0.407609
heating                       0.407609
neutral                       0.577446
somewhat bitter               0.305707
hot burning                   0.271739
salty                         0.883152
burning                       0.407609
less sweet                    0.135870
non-sweet                    10.903533
sweet                        37.941576
```

```
bitter                          40.489130
```

[ ]:
```
lb[:15]["Percentage_of_samples"].mean()
```

[ ]:
```
7.001811594202899
```

[ ]:
```
lb[-15:].sort_values(by="F1_score",ascending=False)
```

[ ]:
```
                 Precision_score  Recall_score  F1_score  \
umami                   0.981132      0.634146  0.770370
bitter                  0.734475      0.724393  0.729399
sweet                   0.800000      0.637088  0.709310
non-sweet               0.692810      0.415686  0.519608
less sweet              1.000000      0.333333  0.500000
burning                 0.600000      0.300000  0.400000
salty                   0.800000      0.190476  0.307692
hot burning             1.000000      0.166667  0.285714
somewhat bitter         1.000000      0.142857  0.250000
neutral                 0.666667      0.142857  0.235294
and tingling            1.000000      0.100000  0.181818
heating                 1.000000      0.100000  0.181818
sour                    1.000000      0.086957  0.160000
pungent                 0.500000      0.071429  0.125000
tasteless               0.423077      0.065089  0.112821

                 Percentage_of_samples
umami                         3.532609
bitter                       40.489130
sweet                        37.941576
non-sweet                    10.903533
less sweet                    0.135870
burning                       0.407609
salty                         0.883152
hot burning                   0.271739
somewhat bitter               0.305707
neutral                       0.577446
and tingling                  0.407609
heating                       0.407609
sour                          1.936141
pungent                       0.577446
tasteless                     7.167120
```

[ ]:
```
lb[-15:].sort_values(by="F1_score",ascending=False)["Percentage_of_samples"].
 ↪mean()
```

[ ]:
```
7.062952898550723
```

[ ]:
```
import numpy as np
import matplotlib.pyplot as plt
from sklearn.datasets import fetch_openml
from sklearn.multioutput import ClassifierChain
from sklearn.model_selection import train_test_split
from sklearn.multiclass import OneVsRestClassifier
from sklearn.metrics import jaccard_score
from sklearn.linear_model import LogisticRegression

from sklearn.ensemble import RandomForestClassifier
from sklearn.model_selection import cross_val_predict
from sklearn.model_selection import cross_validate
from sklearn.metrics import␣
↪f1_score,precision_score,recall_score,jaccard_score,multilabel_confusion_matrix

x_train,y_train,x_test,y_test=iterative_split(x_mord,Y,test_size=0.2)

x_train,x_test=pre_process(x_train,x_test)

base_lr = get_rf()
# Fit an ensemble of logistic regression classifier chains and take the
# take the average prediction of all the chains.
chains = [ClassifierChain(base_lr, order='random', random_state=i)
          for i in range(10)]
for chain in chains:
    chain.fit(x_train, y_train)

Y_pred_chains = np.array([chain.predict(x_test) for chain in
                          chains])

y_pred= Y_pred_chains.mean(axis=0)

# Convert continuous predictions to binary using a threshold
y_pred_binary = (y_pred >= 0.5).astype(int)

# Calculate metrics
print("f1_score {}".format(f1_score(y_test, y_pred_binary, average="micro")))
print("precision_score {}".format(precision_score(y_test, y_pred_binary,␣
↪average="micro")))
print("recall_score {}".format(recall_score(y_test, y_pred_binary,␣
↪average="micro")))
print("jaccard_score {}".format(jaccard_score(y_test, y_pred_binary,␣
↪average="micro")))
```

```
f1_score 0.625137091467427
precision_score 0.7460732984293194
recall_score 0.5379388448471121
```

```
jaccard_score 0.45469049138481177
```

[ ]:
```
Y_pred_chains.shape
```

[ ]:
```
(10, 2359, 44)
```

[ ]:
```
y_pred_t=y_pred
```

[ ]:
```
y_pred_temp=y_pred_t
```

[ ]:
```
for i in range(y_pred_temp.shape[0]):
  for j in range(y_pred_temp.shape[1]):
    if (y_pred_temp[i][j]>0.5):
      y_pred_temp[i][j]=1
    else:
      y_pred_temp[i][j]=0
```

[ ]:
```
y_pred_temp[3]
```

[ ]:
```
array([0., 0., 0., 0., 0., 1., 0., 0., 0., 0., 0., 0., 0., 0., 0., 0., 0.,
       0., 0., 0., 0., 0., 0., 0., 0., 0., 0., 0., 0., 0., 0., 0., 0., 0.,
       0., 0., 0., 0., 0., 0., 0., 0., 0., 0.])
```

[ ]:
```
print("f1_score {}".format(f1_score(y_test,y_pred_temp,average="micro")))
print("precision_score {}".
  ↪format(precision_score(y_test,y_pred_temp,average="micro")))
print("recall_score {}".
  ↪format(recall_score(y_test,y_pred_temp,average="micro")))
print("jaccard_score {}".
  ↪format(jaccard_score(y_test,y_pred_temp,average="micro")))
```

```
f1_score 0.6163101604278075
precision_score 0.7520391517128875
recall_score 0.522083805209513
jaccard_score 0.4454106280193237
```

[ ]:
```
from sklearn.ensemble import RandomForestClassifier
from sklearn.model_selection import cross_val_predict
from sklearn.model_selection import cross_validate
from sklearn.metrics import␣
  ↪f1_score,precision_score,recall_score,jaccard_score,multilabel_confusion_matrix
#Utility function to get model performance
def cross_val(model,x,y):

  #Preprocessing
  imp=KNNImputer(missing_values=np.nan)
  x=imp.fit_transform(x)
```

```
    #Evaluating model performance on training data
␣
↪cross_score=cross_validate(model,x,y,scoring=["f1_micro","precision_micro","recall_micro","

    return cross_score
```

[ ]:
```
model=get_rf()
arr=cross_val(model,x_mord,Y)
print(arr)
```

```
{'fit_time': array([43.59005284, 43.69837213, 42.07647347, 41.49449229,
41.99584389]), 'score_time': array([0.13768053, 0.13763547, 0.12987995,
0.13827991, 0.13846612]), 'test_f1_micro': array([0.69157255, 0.66319444,
0.66903915, 0.66094421, 0.6602954 ]), 'test_precision_micro': array([0.8122449 ,
0.77484787, 0.81034483, 0.77464789, 0.7755102 ]), 'test_recall_micro':
array([0.602118  , 0.57966616, 0.56969697, 0.57634731, 0.57488654]),
'test_jaccard_micro': array([0.52855246, 0.4961039 , 0.5026738 , 0.49358974,
0.49286641])}
```

[ ]:
```
sum(arr['test_f1_micro'])/5
```

[ ]: 0.6690091474562879

[ ]:
```
model=get_bin_rel()
arr=cross_val(model,x_mord,Y)
print(arr)
```

```
/usr/local/lib/python3.10/dist-
packages/sklearn/model_selection/_validation.py:1000: UserWarning: Scoring
failed. The score on this train-test partition for these parameters will be set
to nan. Details:
Traceback (most recent call last):
  File "/usr/local/lib/python3.10/dist-packages/sklearn/metrics/_scorer.py",
line 139, in __call__
    score = scorer._score(
  File "/usr/local/lib/python3.10/dist-packages/sklearn/metrics/_scorer.py",
line 371, in _score
    y_pred = method_caller(
  File "/usr/local/lib/python3.10/dist-packages/sklearn/metrics/_scorer.py",
line 89, in _cached_call
    result, _ = _get_response_values(
  File "/usr/local/lib/python3.10/dist-packages/sklearn/utils/_response.py",
line 199, in _get_response_values
    classes = estimator.classes_
AttributeError: 'BinaryRelevance' object has no attribute 'classes_'
```

```
  warnings.warn(
/usr/local/lib/python3.10/dist-
packages/sklearn/model_selection/_validation.py:1000: UserWarning: Scoring
failed. The score on this train-test partition for these parameters will be set
to nan. Details:
Traceback (most recent call last):
  File "/usr/local/lib/python3.10/dist-packages/sklearn/metrics/_scorer.py",
line 139, in __call__
    score = scorer._score(
  File "/usr/local/lib/python3.10/dist-packages/sklearn/metrics/_scorer.py",
line 371, in _score
    y_pred = method_caller(
  File "/usr/local/lib/python3.10/dist-packages/sklearn/metrics/_scorer.py",
line 89, in _cached_call
    result, _ = _get_response_values(
  File "/usr/local/lib/python3.10/dist-packages/sklearn/utils/_response.py",
line 199, in _get_response_values
    classes = estimator.classes_
AttributeError: 'BinaryRelevance' object has no attribute 'classes_'

  warnings.warn(
/usr/local/lib/python3.10/dist-
packages/sklearn/model_selection/_validation.py:1000: UserWarning: Scoring
failed. The score on this train-test partition for these parameters will be set
to nan. Details:
Traceback (most recent call last):
  File "/usr/local/lib/python3.10/dist-packages/sklearn/metrics/_scorer.py",
line 139, in __call__
    score = scorer._score(
  File "/usr/local/lib/python3.10/dist-packages/sklearn/metrics/_scorer.py",
line 371, in _score
    y_pred = method_caller(
  File "/usr/local/lib/python3.10/dist-packages/sklearn/metrics/_scorer.py",
line 89, in _cached_call
    result, _ = _get_response_values(
  File "/usr/local/lib/python3.10/dist-packages/sklearn/utils/_response.py",
line 199, in _get_response_values
    classes = estimator.classes_
AttributeError: 'BinaryRelevance' object has no attribute 'classes_'

  warnings.warn(
/usr/local/lib/python3.10/dist-
packages/sklearn/model_selection/_validation.py:1000: UserWarning: Scoring
failed. The score on this train-test partition for these parameters will be set
to nan. Details:
Traceback (most recent call last):
  File "/usr/local/lib/python3.10/dist-packages/sklearn/metrics/_scorer.py",
line 139, in __call__
```

```
    score = scorer._score(
  File "/usr/local/lib/python3.10/dist-packages/sklearn/metrics/_scorer.py",
line 371, in _score
    y_pred = method_caller(
  File "/usr/local/lib/python3.10/dist-packages/sklearn/metrics/_scorer.py",
line 89, in _cached_call
    result, _ = _get_response_values(
  File "/usr/local/lib/python3.10/dist-packages/sklearn/utils/_response.py",
line 199, in _get_response_values
    classes = estimator.classes_
AttributeError: 'BinaryRelevance' object has no attribute 'classes_'

  warnings.warn(
```

```
{'fit_time': array([59.69230747, 62.71601367, 64.92409706, 59.50623989,
64.77622724]), 'score_time': array([0.00095844, 0.00090122, 0.00115108,
0.00083876, 0.00095892]), 'test_f1_micro': array([nan, nan, nan, nan, nan]),
'test_precision_micro': array([nan, nan, nan, nan, nan]), 'test_recall_micro':
array([nan, nan, nan, nan, nan]), 'test_jaccard_micro': array([nan, nan, nan,
nan, nan])}
```

```
/usr/local/lib/python3.10/dist-
packages/sklearn/model_selection/_validation.py:1000: UserWarning: Scoring
failed. The score on this train-test partition for these parameters will be set
to nan. Details:
Traceback (most recent call last):
  File "/usr/local/lib/python3.10/dist-packages/sklearn/metrics/_scorer.py",
line 139, in __call__
    score = scorer._score(
  File "/usr/local/lib/python3.10/dist-packages/sklearn/metrics/_scorer.py",
line 371, in _score
    y_pred = method_caller(
  File "/usr/local/lib/python3.10/dist-packages/sklearn/metrics/_scorer.py",
line 89, in _cached_call
    result, _ = _get_response_values(
  File "/usr/local/lib/python3.10/dist-packages/sklearn/utils/_response.py",
line 199, in _get_response_values
    classes = estimator.classes_
AttributeError: 'BinaryRelevance' object has no attribute 'classes_'

  warnings.warn(
```

[ ]:
```
# Importing necessary libraries
import pandas as pd
import matplotlib.pyplot as plt
import seaborn as sns
from sklearn.preprocessing import LabelEncoder
from sklearn.model_selection import train_test_split
```

```
from sklearn.ensemble import RandomForestClassifier
from sklearn.metrics import accuracy_score, classification_report
from sklearn.metrics import accuracy_score, classification_report,␣
 ↪confusion_matrix

import warnings
warnings.filterwarnings('ignore')
```

[ ]:
```
data.head()
```

[ ]:
```
     ID                          Name PubChem CID   CAS number  \
0  0001                (-)-Haematoxylin      320930     517-28-2
1  0002       (+)-4 -hydroxyhernandulcin     126862  145385-64-4
2  0003  (+)-Dihydroquercetin 3-acetate      442540   78834-97-6
3  0004                (+)-Haematoxylin      442514     517-28-2
4  0005              (±)-chiro-inositol         892     643-12-9

                             canonical SMILES    Taste Class taste  \
0         Oc1cc2c(cc1O)C1c3ccc(c(c3OCC1(O)C2)O)O  [sweet]   sweetness
1              CC(C)=CCCC(C)(O)C1CC(O)C(=CC1=O)C  [sweet]   sweetness
2  CC(=O)OC1C(Oc2cc(cc(c2C1=O)O)O)c1ccc(c(c1)O)O  [sweet]   sweetness
3         Oc1cc2c(cc1O)C1c3ccc(c(c3OCC1(O)C2)O)O  [sweet]   sweetness
4                       OC1C(O)C(O)C(O)C(O)C1O  [sweet]   sweetness

                             Reference_(cod)/[pp]
0            Arnoldi1995_((-)-1); Bassoli2001_(39)
1  Kinghorn1998_(2); Kinghorn2002_(7); Kinghorn20…
2  Bouysset2020_(175); Kinghorn2002_(26); Shallen…
3  Arnoldi1995_((+)-1); Arnoldi1996_(12); Bassoli…
4                          Shallenberger1993_[149]
```

[ ]:
```
data.info()
```

```
<class 'pandas.core.frame.DataFrame'>
Index: 2944 entries, 0 to 2943
Data columns (total 8 columns):
 #   Column                Non-Null Count  Dtype
---  ------                --------------  -----
 0   ID                    2944 non-null   object
 1   Name                  2944 non-null   object
 2   PubChem CID           2944 non-null   object
 3   CAS number            2944 non-null   object
 4   canonical SMILES      2944 non-null   object
 5   Taste                 2944 non-null   object
 6   Class taste           2944 non-null   object
 7   Reference_(cod)/[pp]  2944 non-null   object
dtypes: object(8)
```

```
memory usage: 271.5+ KB
```

[ ]:
```
print("\nDistribution of tastes:")
print(data['Class taste'].value_counts())
```

```
Distribution of tastes:
Class taste
bitterness        1183
sweetness          977
non-sweetness      233
tastelessness      203
multitaste         113
umaminess           98
miscellaneous       87
sourness            38
saltiness           12
Name: count, dtype: int64
```

[ ]:
```
from matplotlib import pyplot as plt
import seaborn as sns
data.groupby('Class taste').size().plot(kind='barh', color=sns.palettes.
  ↪mpl_palette('Dark2'))
plt.gca().spines[['top', 'right',]].set_visible(False)
```

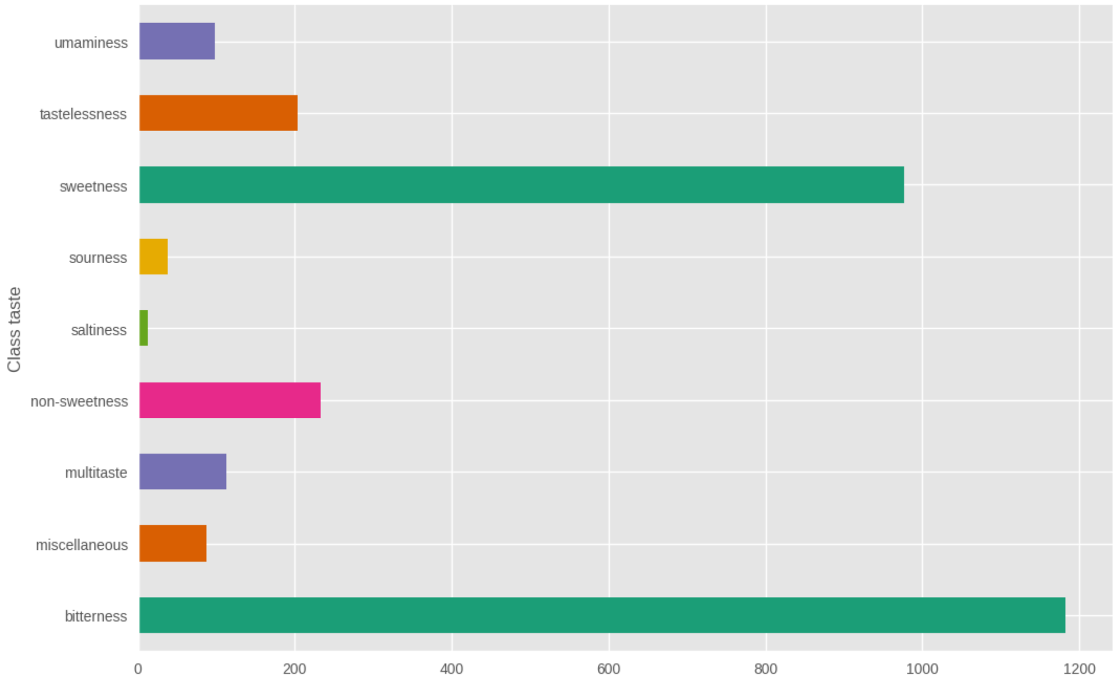

[ ]:
```
plt.figure(figsize=(20, 12))
plt.subplot(2, 2, 1)
plt.xticks(rotation='vertical')
sns.histplot(data['Class taste'], kde=True, bins=20, color='skyblue')
plt.title('Taste Distribution')
```

[ ]:
```
Text(0.5, 1.0, 'Taste Distribution')
```

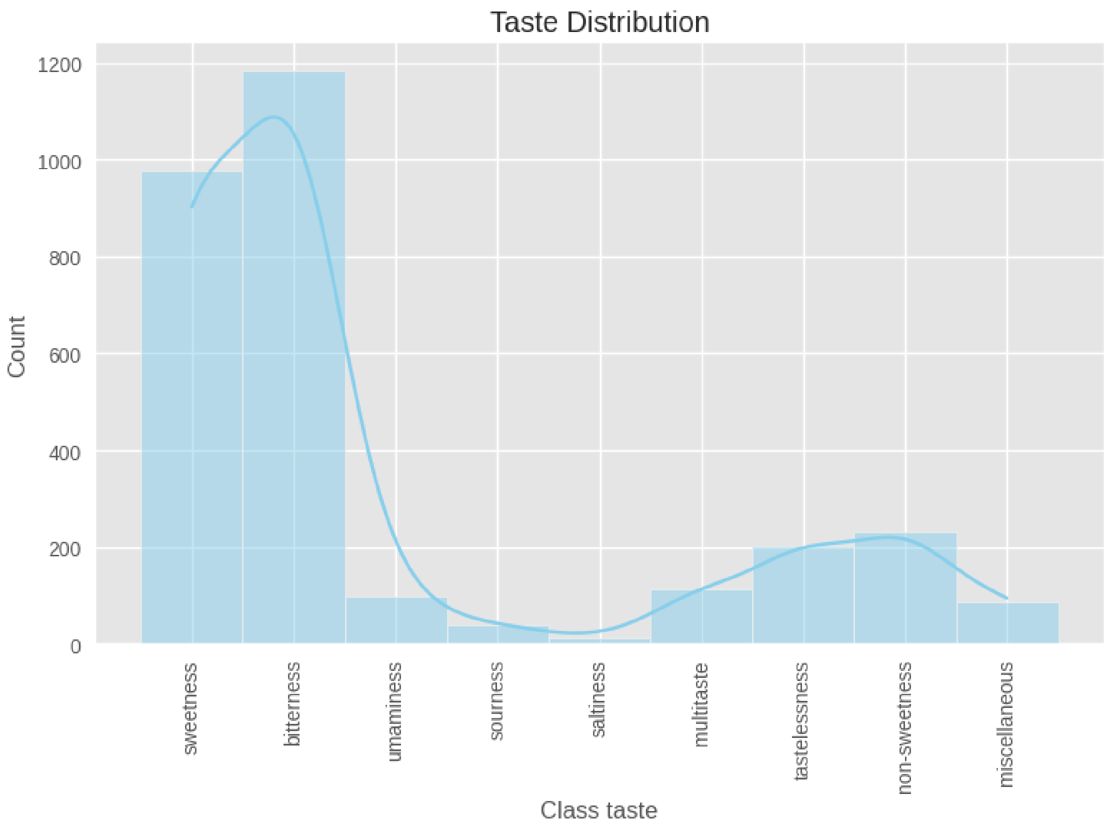


[ ]:
```
# Split the dataset into features (X) and target (y)
X = df.drop('Class taste', axis=1)
y = df['Class taste']
```

[ ]:
```
# Encode the target variable using LabelEncoder
label_encoder = LabelEncoder()
y = label_encoder.fit_transform(y)
```

[ ]:
```
df.head()
```

[ ]:
```
     ID                                        Name PubChem CID  \
0  0566                          Compound (R)-(+)-9           *
1  1221                                   Artemisin       65030
```

```
2   1678                                        L-Leucine       6106
3   1578                                   Hexethal sodium    23690440
4  2832*  Sodium N-[3-chloro-5-(trifluoromethyl)phenyl]s…          *

  CAS number                          canonical SMILES  \
0          *        COc1ccc(cc1O)C1OC2CC3CCC2(CS1)C3(C)C
1   481-05-0       CC1C2C(O)CC3(C)C=CC(=O)C(=C3C2OC1=O)C
2    61-90-5                              CC(C)CC(N)C(O)=O
3   144-00-3       [Na+].CCCCCCC1(CC)C(=O)NC(=NC1=O)[O-]
4          *  [Na+].[O-]S(=O)(=O)Nc1cc(cc(c1)C(F)(F)F)Cl

                               Taste    Class taste  \
0                            [sweet]      sweetness
1                           [bitter]     bitterness
2  [bitter, slightly bitter, non-sweet]     bitterness
3                           [bitter]     bitterness
4                        [non-sweet]  non-sweetness

                            Reference_(cod)/[pp]  SMILES length  \
0                              Bassoli2000_((+)-9)             36
1                           Dagan-Wiener2019_(463)             37
2  Belitz2009_[35]; Dagan-Wiener2019_(751); Glase…            16
3                           Dagan-Wiener2019_(587)             37
4                              Spillane2009b_(51A)             42

   Taste count
0            1
1            1
2            3
3            1
4            1
```

[ ]:

```python
# Assuming y contains your 'Class taste' labels
Y_onehot = pd.get_dummies(df['Class taste'], prefix='', prefix_sep='')
Y_onehot = Y_onehot.reindex(sorted(Y_onehot.columns), axis=1)  # Sort the␣
↪columns alphabetically

# Show the result
print(Y_onehot.head())
```

```
   bitterness  miscellaneous  multitaste  non-sweetness  saltiness  sourness  \
0       False          False       False          False      False     False
1        True          False       False          False      False     False
2        True          False       False          False      False     False
3        True          False       False          False      False     False
4       False          False       False           True      False     False
```

```
   sweetness  tastelessness  umaminess
0       True          False      False
1      False          False      False
2      False          False      False
3      False          False      False
4      False          False      False
```

[ ]:
```python
from sklearn.preprocessing import LabelEncoder, OneHotEncoder

# Use LabelEncoder
label_encoder = LabelEncoder()
y_encoded = label_encoder.fit_transform(df['Class taste'])

# Reshape to make it compatible with OneHotEncoder
y_encoded = y_encoded.reshape(-1, 1)

# Apply OneHotEncoder with the correct argument
onehot_encoder = OneHotEncoder(sparse_output=False)
Y_onehot = onehot_encoder.fit_transform(y_encoded)

# Convert to DataFrame and name columns after the unique classes
Y = pd.DataFrame(Y_onehot, columns=label_encoder.classes_)

# Show the result
Y.head(100)
```

[ ]:
```
    bitterness  miscellaneous  multitaste  non-sweetness  saltiness  sourness  \
0          0.0            0.0         0.0            0.0        0.0       0.0
1          1.0            0.0         0.0            0.0        0.0       0.0
2          1.0            0.0         0.0            0.0        0.0       0.0
3          1.0            0.0         0.0            0.0        0.0       0.0
4          0.0            0.0         0.0            1.0        0.0       0.0
..         …              …           …              …          …         …
95         1.0            0.0         0.0            0.0        0.0       0.0
96         1.0            0.0         0.0            0.0        0.0       0.0
97         0.0            0.0         0.0            0.0        0.0       0.0
98         0.0            0.0         0.0            0.0        0.0       0.0
99         1.0            0.0         0.0            0.0        0.0       0.0

    sweetness  tastelessness  umaminess
0         1.0            0.0        0.0
1         0.0            0.0        0.0
2         0.0            0.0        0.0
3         0.0            0.0        0.0
4         0.0            0.0        0.0
..        …              …          …
95        0.0            0.0        0.0
```

```
96        0.0            0.0        0.0
97        1.0            0.0        0.0
98        1.0            0.0        0.0
99        0.0            0.0        0.0

[100 rows x 9 columns]
```

[ ]:
```python
classes=label_encoder.classes_
print(classes)
```

```
['bitterness' 'miscellaneous' 'multitaste' 'non-sweetness' 'saltiness'
 'sourness' 'sweetness' 'tastelessness' 'umaminess']
```

[ ]:
```python
model=get_rf()
eval_train(model,x_mord,Y)
```

```
f1_score 0.6207621550591327
precision_score 0.8139214334941419
recall_score 0.5016992353440951
jaccard_score 0.4500762195121951
```

[ ]:
```
array([[[1272,  136],
        [ 297,  649]],

       [[2282,    2],
        [  69,    1]],

       [[2263,    1],
        [  88,    2]],

       [[2155,   13],
        [ 160,   26]],

       [[2344,    0],
        [  10,    0]],

       [[2324,    0],
        [  30,    0]],

       [[1477,   95],
        [ 342,  440]],

       [[2169,   23],
        [ 143,   19]],

       [[2276,    0],
        [  34,   44]]])
```

[ ]:
```
model=get_bin_rel()
eval_train(model,x_mord,Y)
```

```
f1_score 0.6251276813074566
precision_score 0.7836107554417413
recall_score 0.5199660152931181
jaccard_score 0.45468053491827637
```

[ ]:
```
array([[[1246,  162],
        [ 272,  674]],

       [[2279,    5],
        [  70,    0]],

       [[2259,    5],
        [  87,    3]],

       [[2142,   26],
        [ 154,   32]],

       [[2342,    2],
        [   8,    2]],

       [[2324,    0],
        [  30,    0]],

       [[1463,  109],
        [ 331,  451]],

       [[2164,   28],
        [ 142,   20]],

       [[2275,    1],
        [  36,   42]]])
```

[ ]:
```
model=get_cls_chain()
eval_train(model,x_mord,Y)
```

```
f1_score 0.6320040383644624
precision_score 0.7786069651741293
recall_score 0.5318606627017842
jaccard_score 0.4619926199261993
```

[ ]:
```
array([[[1246,  162],
        [ 272,  674]],

       [[2281,    3],
```

```
        [  70,    0]],

       [[2263,    1],
        [  87,    3]],

       [[2138,   30],
        [ 153,   33]],

       [[2342,    2],
        [   8,    2]],

       [[2324,    0],
        [  30,    0]],

       [[1450,  122],
        [ 308,  474]],

       [[2157,   35],
        [ 139,   23]],

       [[2275,    1],
        [  35,   43]]])
```

[ ]:
```
model=get_rf()
eval_train(model,x_morg,Y)
```

```
f1_score 0.6563421828908554
precision_score 0.778879813302217
recall_score 0.5671197960917587
jaccard_score 0.4884742041712404
```

[ ]:
```
array([[[1209,  199],
        [ 215,  731]],

       [[2283,    1],
        [  68,    2]],

       [[2260,    4],
        [  84,    6]],

       [[2139,   29],
        [ 131,   55]],

       [[2344,    0],
        [  10,    0]],

       [[2324,    0],
```

```
        [  29,    1]],

       [[1458,  114],
        [ 311,  471]],

       [[2160,   32],
        [ 142,   20]],

       [[2276,    0],
        [  29,   49]]])
```

[ ]:
```
model=get_bin_rel()
eval_train(model,x_morg.astype(float),Y)
```

```
f1_score 0.6595330739299611
precision_score 0.7713310580204779
recall_score 0.5760407816482583
jaccard_score 0.49201741654571846
```

[ ]:
```
array([[[1216,  192],
        [ 233,  713]],

       [[2276,    8],
        [  69,    1]],

       [[2257,    7],
        [  83,    7]],

       [[2122,   46],
        [ 118,   68]],

       [[2344,    0],
        [  10,    0]],

       [[2324,    0],
        [  29,    1]],

       [[1461,  111],
        [ 291,  491]],

       [[2154,   38],
        [ 136,   26]],

       [[2276,    0],
        [  29,   49]]])
```

[ ]:
```
model=get_cls_chain()
eval_train(model,x_morg.astype(float),Y)
```

```
f1_score 0.6665095451331605
precision_score 0.7485442032821599
recall_score 0.6006796941376381
jaccard_score 0.49982325910215625
```

[ ]:
```
array([[[1216,  192],
        [ 233,  713]],

       [[2277,    7],
        [  67,    3]],

       [[2254,   10],
        [  83,    7]],

       [[2120,   48],
        [ 109,   77]],

       [[2344,    0],
        [  10,    0]],

       [[2324,    0],
        [  29,    1]],

       [[1408,  164],
        [ 263,  519]],

       [[2138,   54],
        [ 117,   45]],

       [[2276,    0],
        [  29,   49]]])
```

[ ]:
```
model=get_rf()
eval_train(model,x_path,Y)
```

```
f1_score 0.6015706806282722
precision_score 0.7837653478854024
recall_score 0.4881053525913339
jaccard_score 0.4301759640584051
```

[ ]:
```
array([[[1293,  115],
        [ 386,  560]],

       [[2277,    7],
```

```
        [  63,    7]],

       [[2256,    8],
        [  81,    9]],

       [[2145,   23],
        [ 150,   36]],

       [[2324,   20],
        [   7,    3]],

       [[2321,    3],
        [  28,    2]],

       [[1461,  111],
        [ 317,  465]],

       [[2165,   27],
        [ 140,   22]],

       [[2273,    3],
        [  33,   45]]])
```

[ ]:
```
model=get_bin_rel()
eval_train(model,x_path,Y)
```

```
f1_score 0.6304239401496259
precision_score 0.7632850241545893
recall_score 0.5369583687340697
jaccard_score 0.46030589949016754
```

[ ]:
```
array([[[1237,  171],
        [ 290,  656]],

       [[2268,   16],
        [  64,    6]],

       [[2250,   14],
        [  78,   12]],

       [[2140,   28],
        [ 149,   37]],

       [[2326,   18],
        [   7,    3]],

       [[2322,    2],
```

```
         [  26,    4]],

        [[1455,  117],
         [ 297,  485]],

        [[2170,   22],
         [ 145,   17]],

        [[2272,    4],
         [  34,   44]]])
```

[ ]:
```
model=get_cls_chain()
eval_train(model,x_path,Y)
```

```
f1_score 0.6363636363636364
precision_score 0.7661483253588517
recall_score 0.5441801189464741
jaccard_score 0.4666666666666667
```

[ ]:
```
array([[[1237,  171],
         [ 290,  656]],

        [[2270,   14],
         [  63,    7]],

        [[2251,   13],
         [  80,   10]],

        [[2141,   27],
         [ 145,   41]],

        [[2341,    3],
         [  10,    0]],

        [[2322,    2],
         [  26,    4]],

        [[1439,  133],
         [ 285,  497]],

        [[2167,   25],
         [ 140,   22]],

        [[2273,    3],
         [  34,   44]]])
```

[ ]:
```
#This is a random split
␣
↪X_train,X_test,Y_train,Y_test=train_test_split(morgan_fing(df),Y,random_state=42,test_size=
↪2)
```

[ ]:
```
print("y_train{}  ".format(y_train.shape) +"\n"+"y_test{}   ".format(y_test.
↪shape)+"\n"+"x_train{}  ".format(x_train.shape)+"\n"+"x_test{}  ".
↪format(x_test.shape))
```

```
y_train(585, 44)
y_test(2359, 44)
x_train(585, 1249)
x_test(2359, 1249)
```

[ ]:
```
# Initialize the Random Forest classifier
rf_classifier = RandomForestClassifier(random_state=42)
```

[ ]:
```
# Train the classifier on the training data
rf_classifier.fit(X_train, Y_train)
```

[ ]: RandomForestClassifier(random_state=42)

[ ]:
```
# Predict the classes for test data
Y_pred = rf_classifier.predict(X_test)
```

[ ]:
```
# Calculate accuracy
accuracy = accuracy_score(Y_test, Y_pred)
print("Accuracy:", accuracy)
```

```
Accuracy: 0.6689303904923599
```

[ ]:
```
# Classification report
print("\nClassification Report:")
print(classification_report(Y_test, Y_pred, target_names=label_encoder.
↪classes_))
```

```
Classification Report:
               precision    recall  f1-score   support

   bitterness       0.84      0.88      0.86       228
miscellaneous       0.67      0.17      0.27        12
   multitaste       0.00      0.00      0.00        28
non-sweetness       0.53      0.55      0.54        44
    saltiness       1.00      0.33      0.50         3
     sourness       0.20      0.25      0.22         4
    sweetness       0.87      0.68      0.77       222
tastelessness       0.64      0.24      0.35        38
```

```
   umaminess       1.00      0.50      0.67        10

   micro avg       0.81      0.67      0.73       589
   macro avg       0.64      0.40      0.46       589
weighted avg       0.77      0.67      0.70       589
 samples avg       0.67      0.67      0.67       589
```

[ ]:
```python
from sklearn.metrics import multilabel_confusion_matrix

# Compute confusion matrix for multi-label classification
confusion_matrices = multilabel_confusion_matrix(Y_test, Y_pred)

# Print the confusion matrices for each label
for idx, label in enumerate(onehot_encoder.categories_[0]):
    print(f"\nConfusion matrix for label '{label}':")
    print(confusion_matrices[idx])
```

```
Confusion matrix for label '0':
[[323  38]
 [ 28 200]]

Confusion matrix for label '1':
[[576   1]
 [ 10   2]]

Confusion matrix for label '2':
[[558   3]
 [ 28   0]]

Confusion matrix for label '3':
[[524  21]
 [ 20  24]]

Confusion matrix for label '4':
[[586   0]
 [  2   1]]

Confusion matrix for label '5':
[[581   4]
 [  3   1]]

Confusion matrix for label '6':
[[344  23]
 [ 70 152]]

Confusion matrix for label '7':
```

```
[[546   5]
 [ 29   9]]

Confusion matrix for label '8':
[[579   0]
 [  5   5]]
```

[ ]:
```
# Plot confusion matrices for all labels
for idx, label in enumerate(onehot_encoder.categories_[0]):
    plt.figure(figsize=(4, 3))
    sns.set(font_scale=1.4)
    sns.heatmap(confusion_matrices[idx], annot=True, fmt="d", cmap="Blues",␣
  ↪cbar=False)
    plt.xlabel('Predicted')
    plt.ylabel('Actual')
    plt.title(f"{label_encoder.classes_[idx]}")
    plt.show()
    print(" ")
    print(" ")
    print(" ")
    print(" ")
```

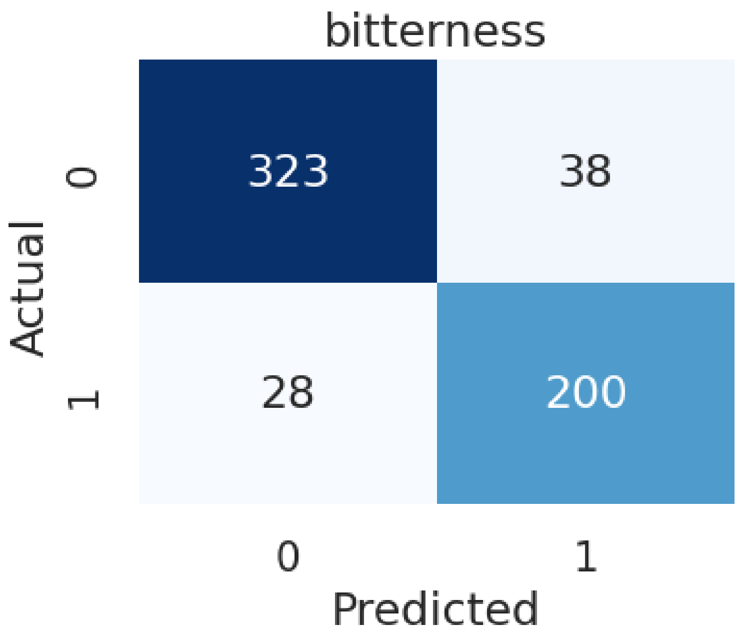

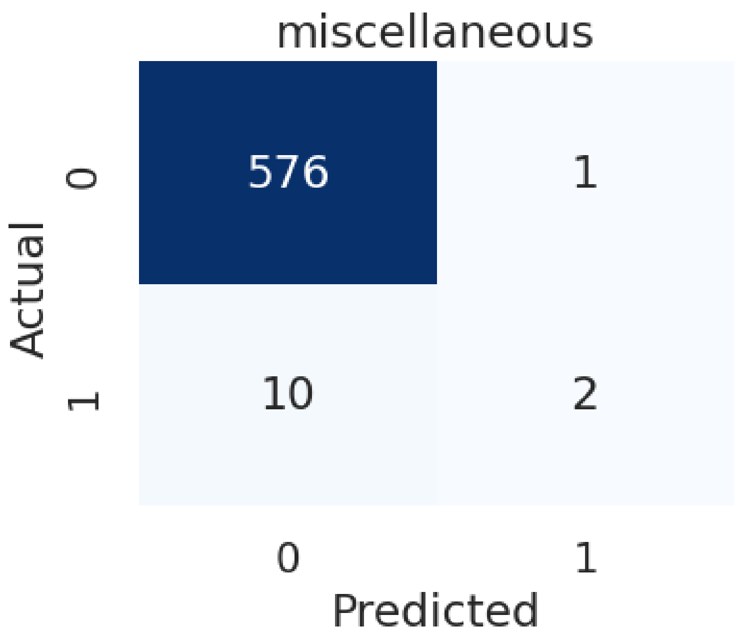
miscellaneous
576
1
10
2
0
1
Actual
Predicted

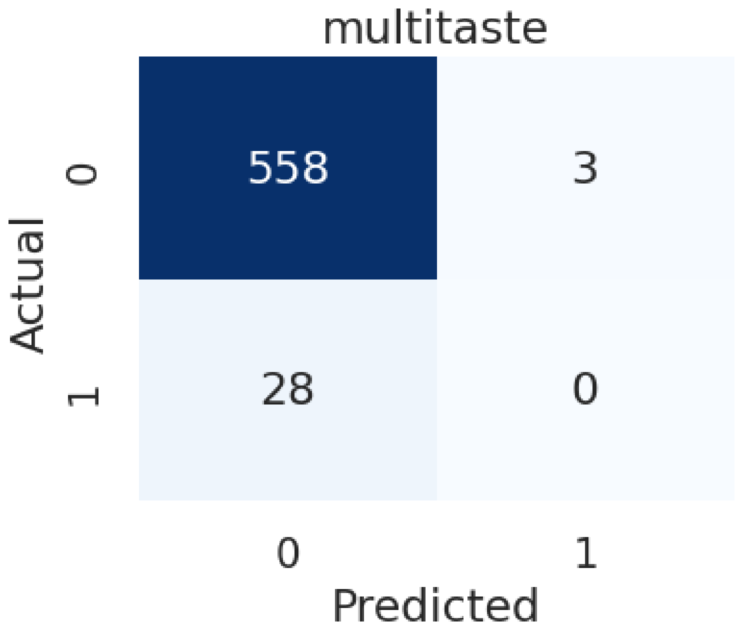
multitaste
558
3
28
0
0
1
Actual
Predicted

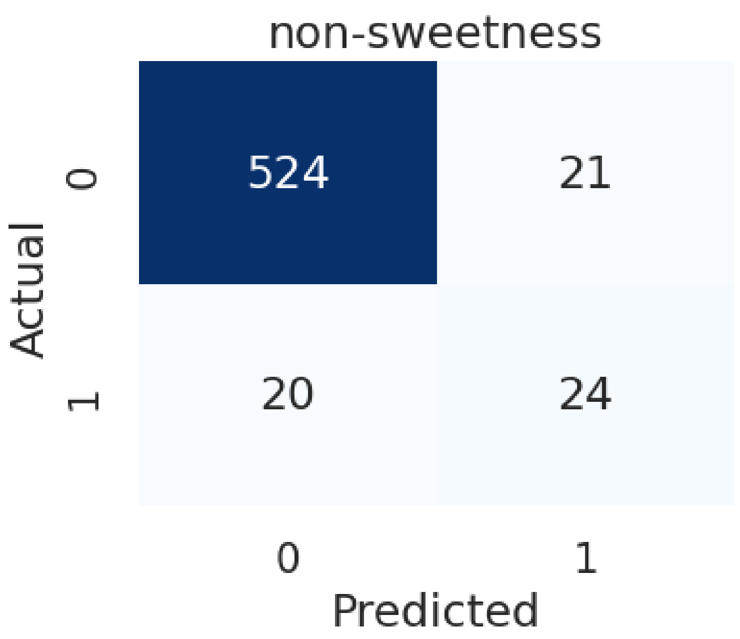
non-sweetness
524
21
20
24
0
1
Actual
0
1
Predicted

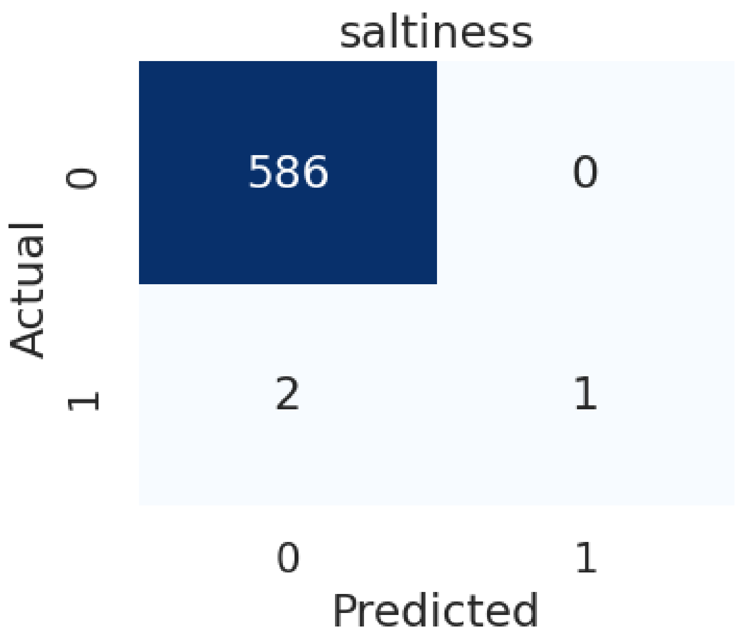
saltiness
586
0
2
1
Actual
0
1
0
1
Predicted

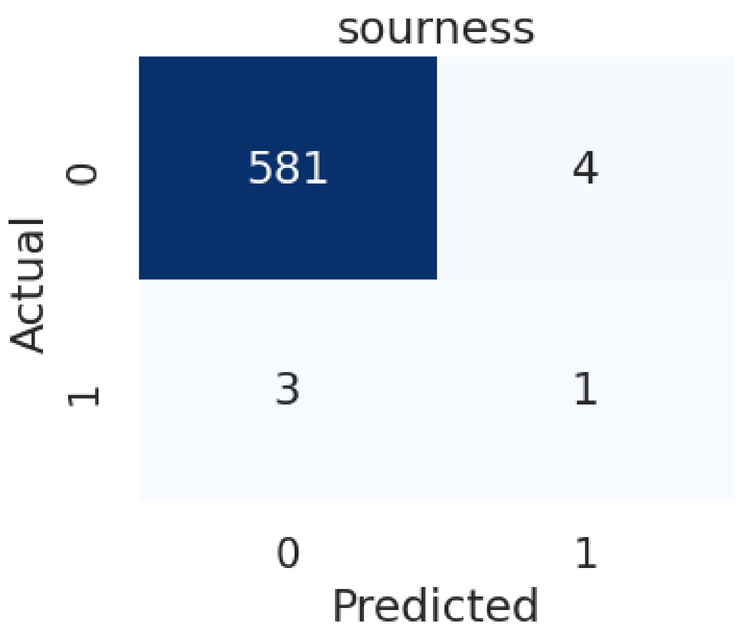
sourness
581
4
3
1
Actual
0
1
0
1
Predicted

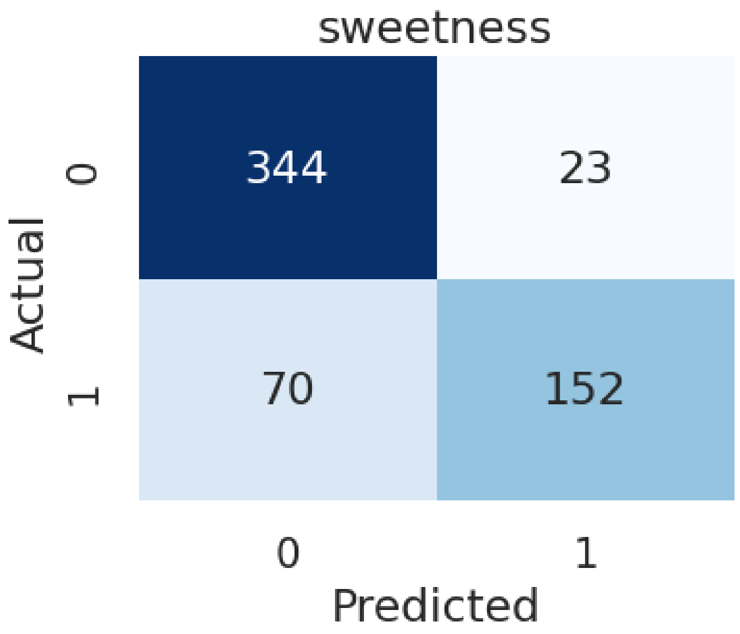

sweetness
Actual
0
1
344
23
70
152
0
1
Predicted

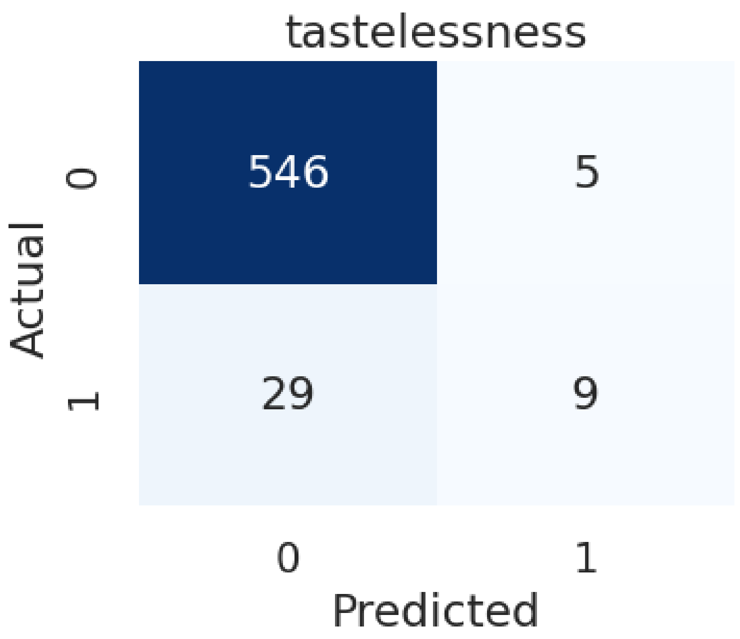
tastelessness
546
5
29
9
0
1
Actual
0
1
Predicted

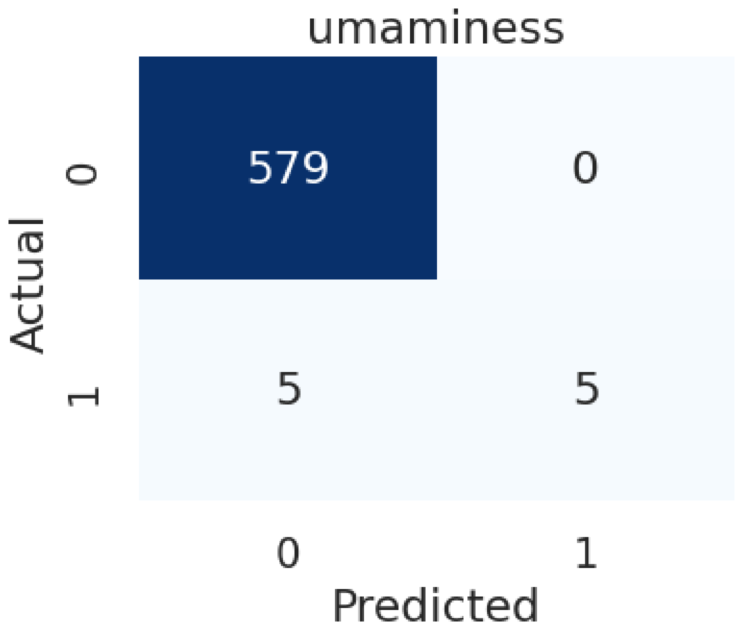
umaminess
579
0
5
5
0
1
Actual
0
1
Predicted

175
